# Human protein complex interfaces reveal candidate assembly roles for initiator methionine excision

**DOI:** 10.64898/2026.08.31.748389

**Authors:** Yie-Hwa Chang

## Abstract

Initiator methionine excision is an early step in protein maturation, but where it might influence protein complex assembly remains poorly understood. This study asked which human protein N-terminal positions engage assembly partners and whether their interface environments point to specific roles for methionine removal. A survey of 7,246 deposited human biological assemblies compared 1,123 eligible proteins. N-terminal positions in MetAP-permissive residue-2 classes were less often interface-engaged than those in nonpermissive classes (37.5% versus 47.5%; adjusted odds ratio 0.65). Nevertheless, 261 permissive-class proteins showed substantial engagement in at least one modeled assembly, including 175 candidates retained after filtering for observed terminal boundaries. Partner-resolved comparisons of proteasome subunits PSMA7, PSMA6 and PSMA4 revealed distinct neighboring-subunit, assembly-chaperone and multi-partner environments. Thus, the possible importance of methionine removal depends not only on sequence, but also on which partner an N-terminal position encounters in a particular assembly context. This structural atlas identifies candidates for direct tests of how verified terminal processing affects partner engagement, complex assembly and function.

## Introduction

Removal of the initiator methionine (iMet) from a nascent polypeptide is among the earliest and most widespread covalent modifications in the cell. In the eukaryotic cytosol, two distinct methionine aminopeptidases carry out the reaction; in yeast, deletion of either gene is tolerated but deletion of both is lethal [1]. iMet excision is generally favored when residue 2 has a small, uncharged side chain—alanine, cysteine, glycine, proline, serine, threonine, or valine—although cleavage efficiency also depends on downstream sequence context, enzyme availability, and competing N-terminal modifications. Quantitative N-terminomics has supported both this specificity rule and the division of labor between the two enzymes in human cells [2], and structural work has shown that the reaction is executed within an organized co-translational processing machine [3,4].

Several consequences of the reaction are well established. The identity of the mature first residue determines whether the N-terminal acetyltransferase NatA can act, and excision creates the free α-amino group that N-myristoyltransferase and the N-terminal methyltransferases require, so three further modification branches are conditional on it. The mature first residue is read by the N-degron pathways: retention of methionine in front of a bulky hydrophobic residue creates a Met-Φ degron recognized in metazoa by the UBR-box machinery [5], and such degrons can be shielded by assembly into a complex [6]. In specific proteins, the mature terminus is itself a functional element, most conspicuously in the α subunits of the 20S proteasome, whose N-terminal tails form the gate over the substrate channel [7–9].

The processing rule is well described, but a different question remains: where could removal of a single initiator methionine matter for protein complex assembly? An N-terminal position that faces an assembly partner may have a different interaction environment from the same position in another structural context. The present study therefore asks how often positions in MetAP-permissive residue-2 classes engage human protein interfaces, which partners they approach, and which examples offer tractable tests of a processing-dependent assembly hypothesis. Answering these questions requires separating sequence class, the first observed coordinate and independently measured terminal chemistry: none alone establishes the chemical state of the structural preparation.

This study maps N-terminal sequence positions across human biological assemblies, compares interface engagement by residue-2 class and tests the effect of requiring an observed coordinate boundary. The analysis finds a population-level tendency toward lower engagement in MetAP-permissive classes while identifying a substantial subset of engaged candidates. Partner-resolved proteasome comparisons then pose three different assembly questions: whether the PSMA7 position engages different neighboring subunits across contexts, whether a PSMA6 position encounters an assembly chaperone, and how the PSMA4 position is packed among multiple partners. Together, the census and contact maps show where to test whether a verified change in terminal processing alters a particular interaction, assembly outcome or function.

## Results

### A structural census over deposited human assemblies

Human UniProt processing annotations were mapped to deposited structures using SIFTS [10–12]. Canonical residue 2 was selected when methionine removal was annotated and residue 1 otherwise. The target space comprised 4,087 accessions, 19,064 multi-chain PDB entries and 101,894 chain-level targets covering the selected position. Entries were processed until distinct-protein coverage saturated. Maximum burial remains sampling-dependent because additional structures can reveal more buried states (Supplementary Fig. 1); the selected positions are not uniformly validated biological termini.

The dataset contains 22,291 chain observations of 1,191 proteins in 7,246 biological assemblies (median nine assembly chains; maximum 435). Excluding 48 signal-peptide, propeptide or transit-peptide cases and 20 immunoglobulin/T-cell receptor constant-region entries leaves 1,123 proteins and 19,967 observations: 696 MetAP-permissive and 427 nonpermissive proteins. Within this eligible cohort, 388 have removal annotations, including 21 specifying alternate processing; 735 lack a removal feature and have MET/MSE at the selected position. Absence of the feature does not establish retention, although 108 of these 735 separately carry an N-acetylmethionine annotation (Supplementary Data 1 and 2).

Burial of the selected sequence position is common in this eligible structural corpus. Interface engagement is defined as loss of at least 25% of the residue’s isolated-chain solvent-accessible surface upon assembly; this conventional threshold is accompanied by continuous distributions. In the original model-based analysis, 464 of 1,123 proteins (41.3%) meet this criterion, and engagement increases with assembly size (odds ratio 1.37 per e-fold increase in chain count, p = 6.7 × 10⁻¹⁰). These measurements establish positional interface engagement in deposited models; they do not by themselves establish the processing state or chemical boundary of the corresponding biological protein.

### MetAP-permissive classes engage interfaces less often

Termini in the seven canonical small, uncharged residue-2 classes that generally permit MetAP cleavage are less engaged at interfaces than termini in nonpermissive classes (Fig. 1a). MetAP-permissive proteins have a median burial of 1.4% and a mean burial of 25.3%, against a median of 20.1% and a mean of 32.6% for the nonpermissive class (two-sided Mann–Whitney p = 5.9 × 10⁻⁴). Dichotomizing, 37.5% of MetAP-permissive termini reach 25% burial against 47.5% of nonpermissive termini (two-sided Fisher odds ratio 0.662, p = 9.5 × 10⁻⁴; at the 20% threshold, odds ratio 0.674, p = 0.0016). The finite minimum measured-residue N distance is larger in the MetAP-permissive class (median 6.75 Å versus 5.70 Å; two-sided Mann–Whitney p = 0.004983; 692 permissive and 426 nonpermissive proteins). Five nonfinite exported values are excluded from this distance comparison, not from the burial analysis (Supplementary Data 4).

**Figure 1.**
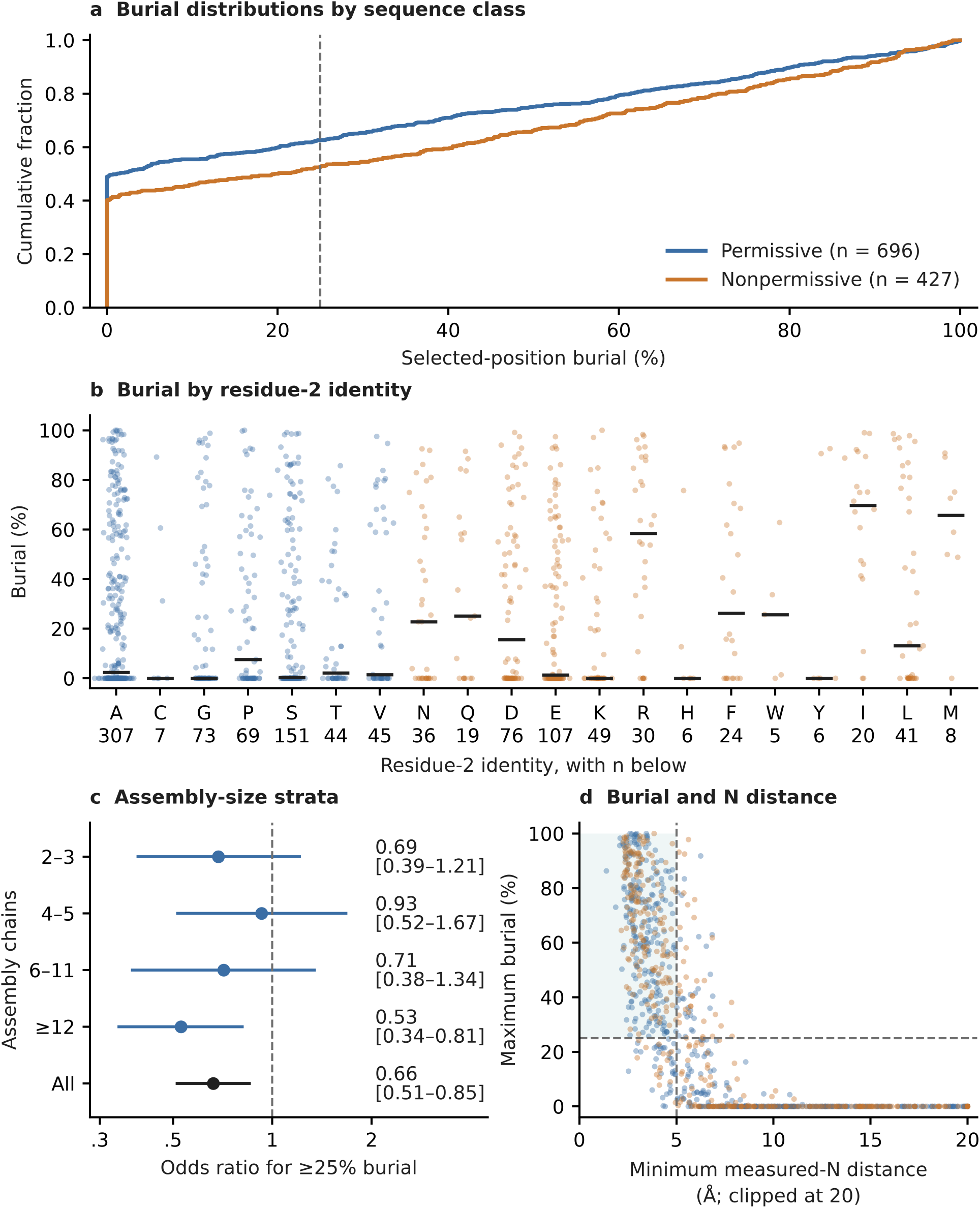
Interface engagement of selected N-terminal sequence positions in biological assemblies. a, Empirical cumulative distributions of burial for 696 MetAP-permissive and 427 MetAP-nonpermissive residue-2 sequence classes. Curves show the proportion of proteins with burial at or below each value; no binning or smoothing is applied. Below the 25% threshold lie 62.5% of permissive and 52.5% of nonpermissive selected positions. Burial is the percentage of the selected residue’s isolated-chain solvent-accessible surface lost on assembly. b, Distributions for all 20 residue-2 identities, with sample sizes below the axis and medians as black bars. Blue denotes Ala/Cys/Gly/Pro/Ser/Thr/Val; orange denotes nonpermissive identities. No identity is excluded from the primary analysis; the Cys, His, Trp and Tyr groups have fewer than ten proteins and are descriptive. c, Stratum-specific and pooled unadjusted odds ratios for burial ≥25%, with exact conditional 95% confidence intervals. The pooled point is not the log-chain-count-adjusted logistic estimate, which is reported in the Results. Odds ratios below one indicate lower engagement odds in the permissive class. d, Maximum burial against the minimum measured-residue N distance across observations; the two summaries need not come from the same structure. Four permissive and one nonpermissive protein with nonfinite minimum distances are omitted from this panel only, leaving 692 and 426 points. Distances are clipped at 20 Å for display. The shaded quadrant requires burial ≥25% and minimum distance ≤5 Å and contains 231 permissive and 170 nonpermissive proteins; the burial-only permissive set contains 261. Original extraction values are retained; first-observed sensitivities are reported separately. A measured N atom is not necessarily a free terminal amino group.

Sensitivities addressing deposition multiplicity, modeled MET/MSE and alternative class or aggregation rules retained odds ratios below one (Supplementary Table 1; Supplementary Data 5). Removing nonfirst observations and reselecting maximum burial retained 845 proteins: 175/463 permissive and 178/382 nonpermissive proteins were engaged (two-sided Fisher odds ratio 0.696, p = 0.0116). Adjustment for log chain count gave an odds ratio of 0.669 (95% confidence interval 0.506–0.886, p = 0.00503). Excluding PSMA4, PSMA6, PSMA7 and Ran alone also retained the direction (odds ratio 0.652, p = 0.000724). These are association sensitivities, not processing-state validation.

Reassigning Cys to the nonpermissive class or adding Asn/Gln, Asp/Asn or Asp/Asn/Gln to the permissive class gave odds ratios of 0.663–0.731 (two-sided Fisher p = 0.000974–0.023274). Median collapse, largest-assembly selection and heteromeric-maximum restriction gave odds ratios of 0.554, 0.663 and 0.625 (p = 0.0000177, 0.001518 and 0.001846; Supplementary Table 1; Supplementary Data 4–5). These exploratory sensitivities are not independent replications. Ala/Cys/Gly/Pro/Ser/Thr/Val remains the primary class definition.

Because burial increases with assembly size, the comparison was stratified (Fig. 1c). Point estimates were below unity in all four strata, strongest in assemblies of at least twelve chains (47.5% versus 63.1% at 25% burial; odds ratio 0.528, p = 0.0029) and undetectable in the smallest assemblies (odds ratio 0.686, p = 0.17). Across all 1,123 proteins, adjustment for log chain count gave an odds ratio of 0.652 (95% confidence interval 0.509–0.837, p = 7.6 × 10⁻⁴), and a class-by-size interaction did not improve fit (p = 0.13): the association is size-adjusted and most visible in the largest machines.

The association identifies sequence classes with different interface environments. Separating contributions from processing, correlated residue-2 chemistry and structural sampling requires verified proteoforms and controlled perturbations.

### A model-based candidate set and its boundary-qualified subset

Despite the distribution shift, 261 of 696 MetAP-permissive proteins reach 25% burial, including 175 retained after nonfirst-observation filtering and maximum reselection (Supplementary Data 3). These sets nominate interfaces for direct tests. Loss of eligibility under the stricter coordinate criterion does not disprove physiological terminal contact.

Of the 261 candidates, 147 are measured at the removal-annotation-selected residue and 114 at modeled methionine (109 MET; five MSE). A removal-annotated residue can be internal in an engineered or unprocessed construct, while missing modeled Met can reflect unresolved density. Coordinate identity and biological terminal chemistry therefore remain separate evidence layers.

### Group-level associations depend on membership and boundary eligibility

The original prefix map provides descriptive summaries of annotation-selected positions (Fig. 2). A reconstructed exploratory screen compared each group with the remaining MetAP-permissive proteins and corrected across the twelve displayed groups plus a histone-symbol group. In the original 696-protein permissive background, the proteasome/assembly group contained 12 engaged proteins among 15 eligible members (odds ratio 6.94, two-sided Fisher p = 0.000879, Benjamini–Hochberg q = 0.0114), and the histone-symbol group contained 9 among 11 (odds ratio 7.73, p = 0.00332, q = 0.0216). CCT/TRiC, OXPHOS and spliceosomal prefix groups were nominal but did not pass this thirteen-test correction (q = 0.0563 each). After filtering nonfirst observations and reselecting maxima in the 463-protein permissive background, the proteasome/assembly result weakened to 7/10 (odds ratio 3.96, p = 0.0466, q = 0.202); the histone-symbol group retained 9/9 (p = 0.000138, q = 0.00180). All thirteen comparisons are reported in Supplementary Table 3 and Supplementary Data 9.

**Figure 2.**
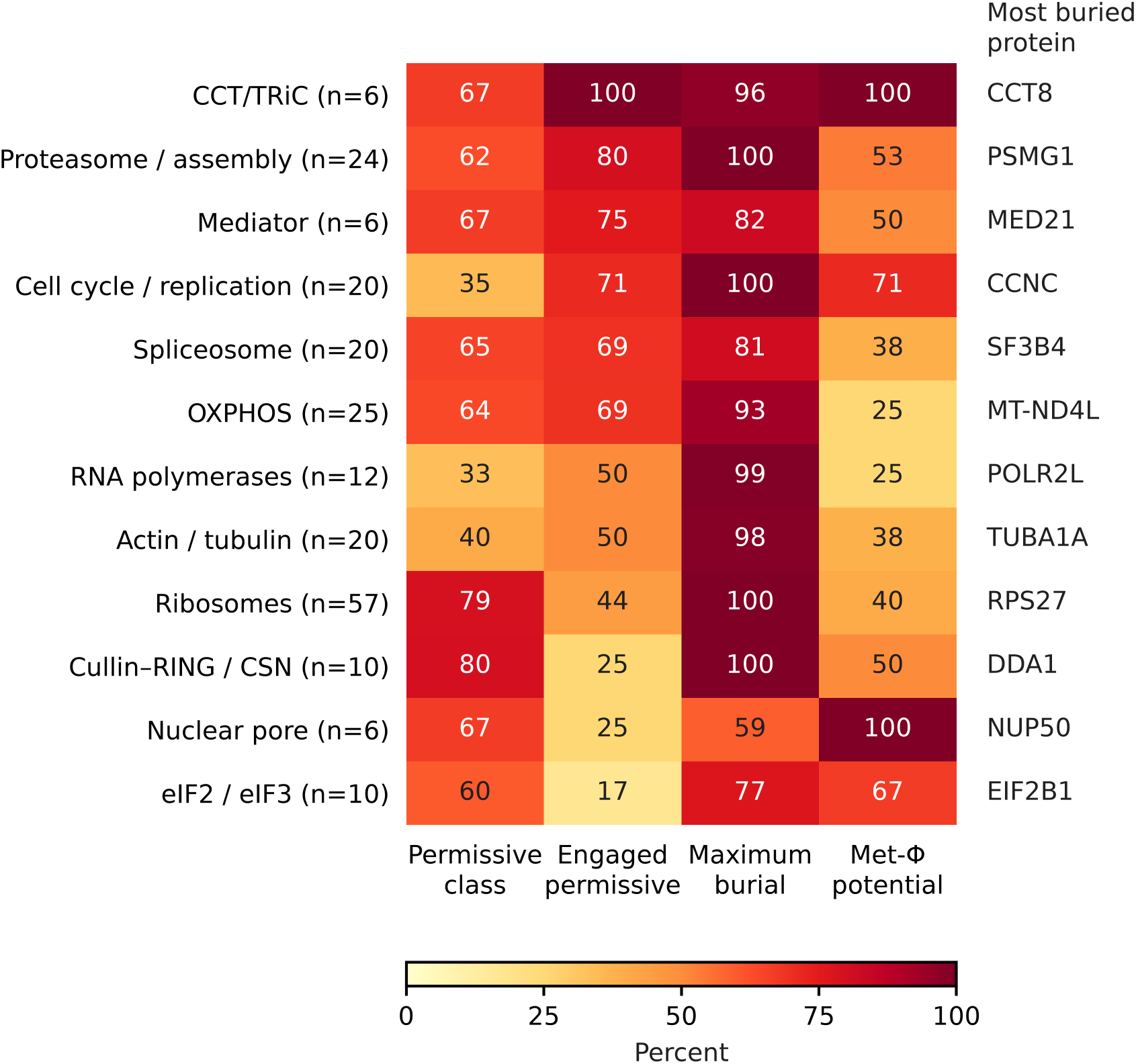
Descriptive positional burial and sequence-based degron potential in prefix-defined groups. Rows are gene-prefix groups with at least four scored proteins in the original 1,123-protein cohort, not curated complex memberships. Columns give the proportion in the MetAP-permissive sequence class, the proportion of that class with selected-position burial ≥25%, the maximum burial in the row, and the proportion of the permissive class meeting the sequence rule for potential Met-Φ recognition upon retention. Sequence class and degron potential are predictions, not sample-specific processing or degradation measurements. Rightmost labels identify the most buried protein. The displayed values retain the original extraction and are descriptive; no enrichment p or q value is encoded. The thirteen-group inferential family adds histones to the twelve displayed groups. Prefix tests, curated-membership tests and first-observed sensitivities are reported separately in Supplementary Tables 3–4 and Supplementary Data 9, with their own denominators and correction families.

A separate screen used manually curated Complex Portal membership rather than gene prefixes. Across 64 complexes with at least four scored permissive members, the original 26S proteasome comparison (CPX-5993) gave 10/13 engaged members (odds ratio 5.74, exact conditional 95% confidence interval 1.46–32.66, p = 0.00628, BH q = 0.0309). After first-observed filtering it gave 7/10 (odds ratio 3.96, p = 0.0466, q = 0.481), and no complex passed catalogue-wide BH correction. Collapsing identical tested accession sets to 42 comparisons changed the original 26S q to 0.0507; dependence-conservative BY correction also did not support a 26S discovery (q = 0.147). Thus, the original enrichment observation is sensitive to coordinate eligibility and multiplicity treatment. Overlapping catalogue records are not independent confirmations, and curated membership does not identify the assembly producing a protein’s maximum burial. Full memberships and outcomes are provided in Supplementary Data 9, with key comparisons in Supplementary Table 4.

### Recurrence within selected observations versus biological persistence

Among the 261 candidates, 126 were engaged in ≥75% of extracted observations, 80 in 25% to <75%, and 55 in <25%; 105 combined burial and measured-residue N distance ≤5 Å in ≥75% of observations (Supplementary Data 6). These counts describe recurrence within the original extraction, not constitutive physiological contacts. Recurrence has not been recalculated for boundary-qualified observations or alternative terminal residues.

PDB-level collapsing correlated with the original engagement fractions (Spearman ρ = 0.925), preserved categories for 221/261 candidates and gave leave-one-entry-out stability for 149/193 multiply represented candidates. These sampling checks neither recover omitted terminal states nor correct internal-residue assignments.

PSMA4, PSMA6 and PSMA7 were engaged in every Met1-selected observation, yet independent top-down mass spectrometry reports methionine removal and N-acetyl-serine for P25789, P60900 and O14818, respectively [13]. This evidence is not preparation-matched. The frozen structures also contain Ser2-starting models excluded by Met1 selection: unengaged PSMA6 in 4R3O and 6KWY and weakly engaged PSMA4 in 8TM4. Thus, complete recurrence within the selected observations does not establish methionine retention or constitutive engagement across terminal states. CCT/TRiC and other recurrent candidates similarly require boundary qualification.

Ran illustrates a construct boundary. In 7MO1/A, the prefix is Ser0–Met1–Ala2–Ala3, with 1.331 Å between Met1 carbonyl C and Ala2 N. Ala2 burial (66.8%) and interchain N distance (2.72 Å) reproduce, but the measured N is internal. The contrasting 3GJ0 observation is also nonfirst, leaving no eligible Ran observation in the strict sensitivity. These are construct-dependent contacts, not a matched processed-versus-retained comparison. Testing effects on transport-factor binding, nucleotide exchange or turnover requires preparations with verified boundaries and acetylation states.

Ribosomal, spliceosomal, Mediator and OXPHOS recurrence summaries likewise guide structural inspection within the extracted observation set (Supplementary Data 6). They measure protein-chain contacts, exclude RNA/DNA occlusion and do not establish native chemistry; repeated chains and entries are not independent biological samples.

### CCT/TRiC candidates require boundary-qualified interpretation

In the original extraction, four permissive-class CCT/TRiC members cross the engagement threshold: CCT8, CCT2, CCT6A and CCT5 have maximum burial values of 96.4%, 86.1%, 83.0% and 75.8%, respectively. The curated CCT comparison is nominal in the original extraction (4/4; p = 0.0195; q = 0.0734 across 64 eligible curated complexes). Boundary filtering retains CCT8, CCT2 and CCT5 at 96.4%, 71.3% and 75.8%, whereas CCT6A falls to 0.0%; the group result becomes 3/4 (p = 0.154; catalogue-wide q = 0.481). Although the original comparison passes correction within the smaller five-question focused family (q = 0.0325), that selected family does not override the broader-screen result or boundary sensitivity. These findings motivate candidate-specific tests, not a confirmed CCT-wide processing dependence (Supplementary Table 4; Supplementary Data 9).

### Histone-tail recognition and spliceosomal candidates

Histone-associated positions form a mechanistically distinct cluster. The histone-symbol screen contains nine engaged records among eleven eligible permissive proteins (odds ratio 7.73, p = 0.00332, thirteen-test BH q = 0.0216). The boundary-qualified result is 9/9 (BH q = 0.00180; Benjamini–Yekutieli q = 0.00571). This stronger numerical result reflects retention of nine engaged records and loss of two unengaged records from eligibility, not independent replication. Several selected maxima come from short N-terminal peptides bound to recognition proteins: H3(1–15) on the ATRX ADD domain (PDB 3QLA) [14], an H3 peptide on LSD2/KDM1B (4FWF) [15] and an H4/H2A peptide on NatD (4U9W) [16]. These examples concern motif recognition rather than burial within an intact nucleosome. The curated nucleosome-membership union is a different set, with 5/7 engaged originally and 5/5 after filtering; membership does not establish that the selected structures depict nucleosomes. Neither first-observed status nor positional burial proves native chemistry, and homologous histone variants are not independent biological or evolutionary replications. The supported corpus association therefore motivates tests of terminal-motif recognition, not a demonstrated nucleosome-assembly vulnerability (Supplementary Tables 3–4).

The broader curated union of seventeen spliceosomal complexes does not show enrichment: 11 of 22 eligible permissive members are engaged in the original extraction (odds ratio 1.70, p = 0.264), compared with 4 of 13 after boundary filtering (odds ratio 0.725, p = 0.774). The prefix grouping has different membership and gives 9/13 originally and 4/7 after filtering; it cannot substitute for the curated union. Individual candidates remain useful: SNRNP70 retains 85.7% burial, whereas SF3B4, SF3B5 and PRPF19 lack eligible observations under the first-observed sensitivity and require boundary validation. These observations support targeted inspection, not spliceosome-wide enrichment (Supplementary Tables 3–4; Supplementary Data 9).

### Oxidative phosphorylation requires a mitochondrial branch

Oxidative phosphorylation complexes require an explicit processing distinction. Nuclear-encoded respiratory-chain subunits fall under the cytosolic MetAP1/MetAP2 framework before import, whereas the eleven mitochondrially encoded proteins in the census are synthesized with formylmethionine and processed by mitochondrial deformylation and MetAP1D [17], so they are not direct tests of cytosolic MetAP specificity. Excluding them retains the direction of the proteome-wide result (odds ratio 0.663, p = 0.000971). The curated union of respiratory complexes I–V contains 11 engaged proteins among 16 eligible permissive members under both extraction schemes. After boundary filtering, its odds ratio is 3.80 (p = 0.0155), with BH q = 0.0389 across five focused questions but q = 0.101 in the thirteen-prefix analysis. Five of six mitochondrially encoded OXPHOS members and six of ten nuclear-encoded members are engaged; neither subgroup passes correction across the two encoding comparisons. An exact one-sided enrichment test conditional on assembly-size and deposition-count strata gives boundary p = 0.0379 and five-question q = 0.0947. OXPHOS is therefore retained as an exploratory compartment- and assembly-context lead, not evidence of general cytosolic MetAP dependence (Supplementary Table 4; Supplementary Data 9).

### Proteasome terminal evidence and structural context define distinct biological questions

The proteasome provides a biological case study in which processing evidence and structural context can be examined without treating them as interchangeable (Fig. 3). Independent top-down measurements report removed, N-acetyl-serine proteoforms for PSMA4, PSMA6 and PSMA7 [13], whereas the audited coordinates include both Met1-starting and Ser2-starting models. These observations establish different kinds of evidence: the proteoform measurements identify terminal chemistry in an independent preparation, and the coordinate measurements describe positional geometry in specific assemblies. Neither makes the structural samples a matched processing experiment.

**Figure 3.**
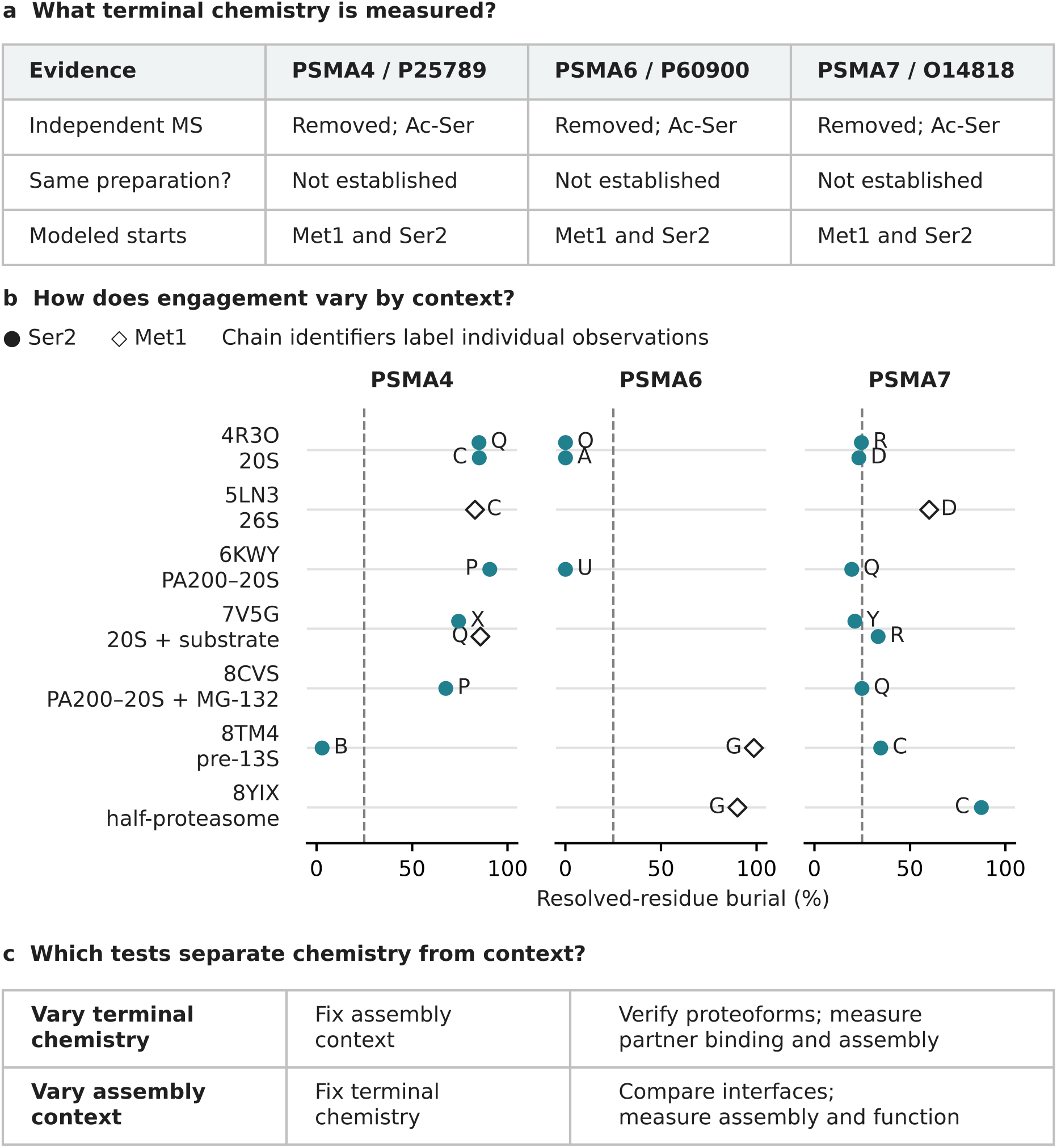
Evidence-aware comparison of proteasome N-terminal positions across deposited structural contexts. a, Independent top-down mass-spectrometric evidence reports initiator-methionine removal and N-acetyl-serine for PSMA4/P25789, PSMA6/P60900 and PSMA7/O14818 [13]. These measurements were not made on the plotted structural preparations; accession-matched evidence is distinguished from sample-matched validation. b, All 22 unchanged-model, first-observed Met1 or Ser2 observations from the completed focal audit are shown across seven PDB entries, with chain labels beside points. Filled circles indicate Ser2 without modeled upstream amino acids; open diamonds indicate modeled Met1. The common 0–100% scale shows full-assembly burial of resolved target-residue atoms. The dashed line marks the descriptive ≥25% threshold, not a biological cutoff. Blanks indicate unavailable qualifying measurements, not zero burial. Repeated chains and entries are not independent biological replicates. Deposited entry contexts do not define an ordered maturation trajectory. Missing modeled Met does not establish excision, and first-observed Ser2 does not establish an acetylated mature terminus. c, Proposed, unperformed experiments distinguish terminal-chemistry effects at fixed assembly context from context effects at fixed terminal chemistry. No static Met-deletion model, pooled processing-group test or disease outcome is included. Five side chains are incomplete: OG is absent from PSMA6 Ser2 in 4R3O/A and 4R3O/O, PSMA4 Ser2 in 8TM4/B and PSMA7 Ser2 in 8TM4/C; CG, SD and CE are absent from PSMA6 Met1 in 8TM4/G. No atoms were rebuilt. Figure 4 resolves partners for the same observations; source measurements and code are in Supplementary Data 7.

The comparison includes all 22 first-observed Met1 or Ser2 measurements for these three proteins in the completed focal geometry audit across seven structural entries, including repeated chains rather than only selected extremes. Partner-resolved analysis extended this comparison to the surrounding chains (Fig. 4; reproduced in Supplementary Fig. 3; source measurements in Supplementary Data 7): all 32 target–partner chain pairs containing a heavy-atom approach of ≤4 Å had unique accession assignments in the frozen chain mapping. Full-assembly burial and nearest-contact distances reproduced the preceding geometry measurements to numerical precision. Pair-only burial measures occlusion of the resolved target-residue atoms with one partner chain present, normalized to their surface area in the isolated target chain; values are nonadditive and are not fractions of total assembly burial. Met1 and Ser2 measurements remain distinct, and neither chain copies nor structural entries were treated as independent biological replicates. Internal positions such as PSMA2 Ala2 in 8YIX and PSMA3 author-numbered Ser1 in 6KWY remain excluded from this first-observed-position comparison.

**Figure 4.**
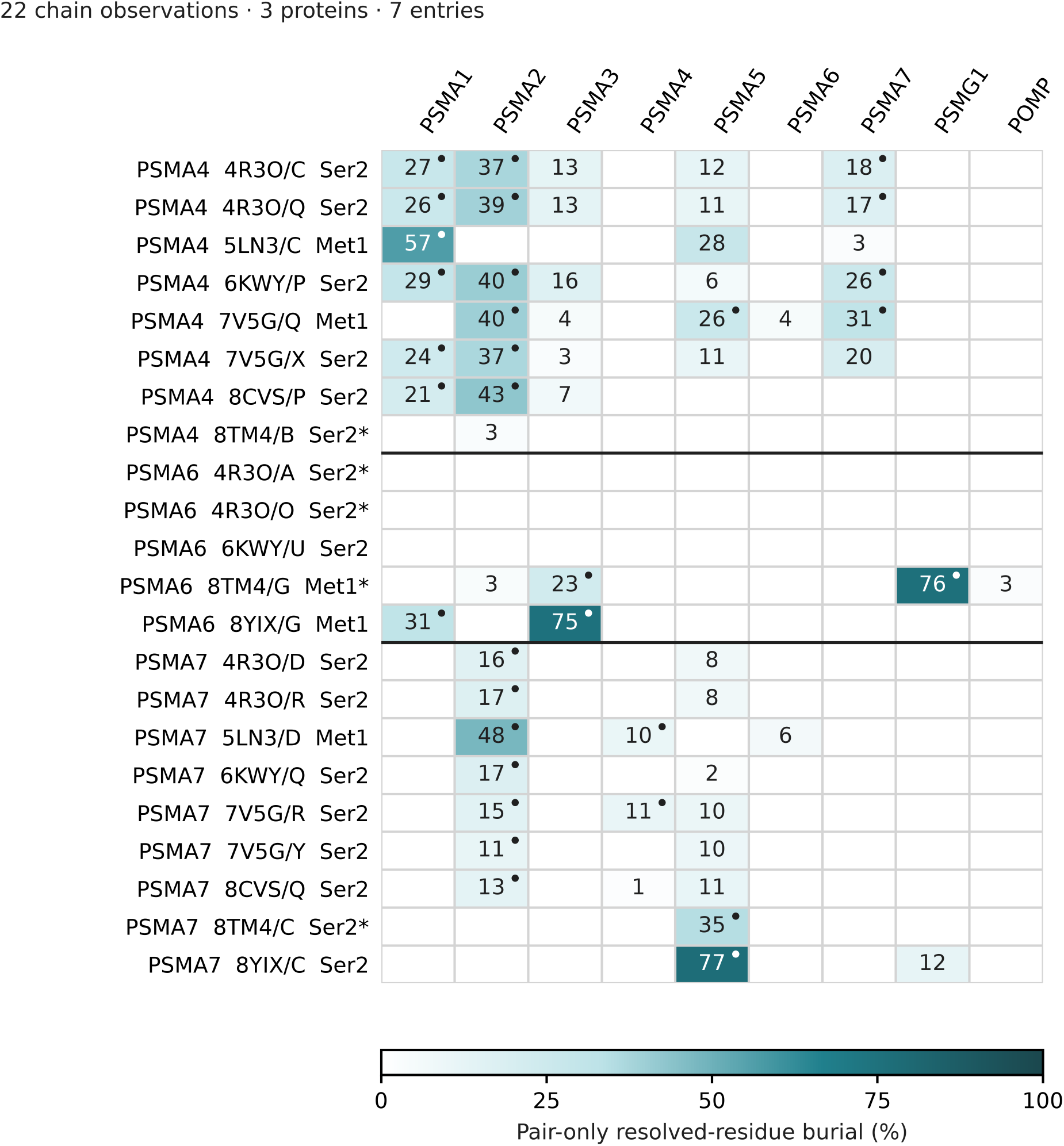
Partner-resolved geometry of selected PSMA4, PSMA6 and PSMA7 N-terminal positions. Rows show 22 first-modeled Met1 or Ser2 observations from seven entries; columns identify partner genes. Shading and values ≥1% report burial of resolved target-residue atoms with one partner chain present. Dots mark heavy-atom approaches ≤4 Å. Pair-only burial uses target-residue SASA in the isolated target chain as denominator; values are nonadditive, not fractions of total assembly burial. The largest chain-specific value is displayed per partner gene; no row contains multiple informative chains of the same gene at the displayed thresholds. Blanks mean <1% burial or no eligible partner, not absence of interaction. Pair SASA was calculated for partners within 8 Å; more distant pairs are marked not calculated in Supplementary Data 7. The color scale is 0–100%. Asterisks mark missing OG in PSMA6 Ser2 (4R3O/A, 4R3O/O), PSMA4 Ser2 (8TM4/B) and PSMA7 Ser2 (8TM4/C), and missing CG, SD and CE in PSMA6 Met1 (8TM4/G). No atoms were rebuilt. Met1 and Ser2 are distinct positions, not sample-validated processing states. Proximity does not establish bonds, energies or function; chains and entries are not independent biological replicates. PSMA7 Ser2 occupies PSMA2-facing or PSMA5-facing environments, PSMA6 includes a PSMG1-facing Met1 backbone, and PSMA4 illustrates multi-partner packing. These comparisons do not establish temporal exchange, processing causality or disease mechanisms. Supplementary Data 7 supplies measurements, contacts, backbone sensitivities, chain assignments, code and frozen inputs. Main Fig. 4 and Supplementary Fig. 3 show the same map, not independent analyses. This selected comparison is distinct from group-level tests in Supplementary Tables 3–4 and Supplementary Data 9 and does not independently confirm proteasome-wide enrichment.

PSMA7 provided a same-position comparison across structural contexts. Among eight Ser2-starting observations, PSMA2 was the nearest partner and the largest pair-only occluder in six observations from four entries, whereas PSMA5 occupied these roles in the pre-13S and half-proteasome entries. The contrast persisted after excluding the pre-13S Ser2 model, which lacked OG. In the complete Ser2 models, PA200–20S PSMA7 in 6KWY/Q approached PSMA2/O at 2.92 Å, with 17.3% pair-only burial and 19.4% total burial, whereas half-proteasome PSMA7 in 8YIX/C approached PSMA5/D at 3.27 Å, with 77.2% pair-only burial and 87.4% total burial. Common-backbone measurements also distinguished these environments: the respective partners produced 28.9% and 79.1% pair-only backbone burial. These observations identify different partner environments at the same modeled Ser2 position, without establishing a temporal transition or a processing-dependent mechanism.

PSMA6 revealed a candidate chaperone-associated environment. In the pre-13S model 8TM4/G, the modeled Met1 backbone approached PSMG1/c at 3.27 Å, and PSMG1 alone reproduced its 98.2% backbone burial. In the half-proteasome model 8YIX/G, the complete Met1 instead contacted PSMA3 and PSMA1, with PSMA3 supplying the nearest approach and largest pair-only occlusion. The pre-13S Met1 lacks CG, SD and CE; consequently, its 98.5% total target-residue burial refers only to the five resolved atoms and cannot establish burial of a complete methionine side chain. All three Ser2-starting PSMA6 observations had zero interchain burial and no other protein atom within 5 Å, but only the 6KWY/U observation contained a complete Ser2 side chain. This selected contrast does not establish a general difference between processed and unprocessed PSMA6.

PSMA4 illustrated multi-partner packing. Five complete Ser2 observations showed 67.6–90.7% total burial, with PSMA2 providing the largest individual pair-only occlusion, 37.0–43.0%, whereas the nearest partner could differ. In 6KWY/P, PSMA7 supplied the shortest approach, but PSMA2 produced the largest pair-only burial. The pre-13S PSMA4 Ser2 position in 8TM4/B showed only 2.8% resolved-atom burial and no contact within 4 Å; its nearest partner was PSMA2 at 4.39 Å. Although this model lacked OG, low backbone burial of 4.1% supported a qualitatively less occluded modeled backbone environment. No chaperone approached this focal position within 4 Å. Together, the partner-resolved comparisons connect heterogeneous N-terminal geometry to specific protein partners, while keeping chemical-state, assembly-context and model-completeness questions separate.

### The nuclear transport system as a contrasting case

Nuclear transport provides contrasting environments (Fig. 5). Nonpermissive SEH1L buries modeled Met1 by 37.1–38.0% at 3.85–3.86 Å in 304-chain 5A9Q. Permissive NUP188 Ala2 is unburied and 40.5–46.9 Å from another protein chain in 208-chain 5IJO, whereas nonpermissive NUP93 has 7.6–7.7% burial at 6.68 Å. NUP50 instead buries its annotation-selected alanine by 59.1% at 4.88 Å in its importin-α-binding element, identifying a regulatory rather than scaffold interface.

**Figure 5.**
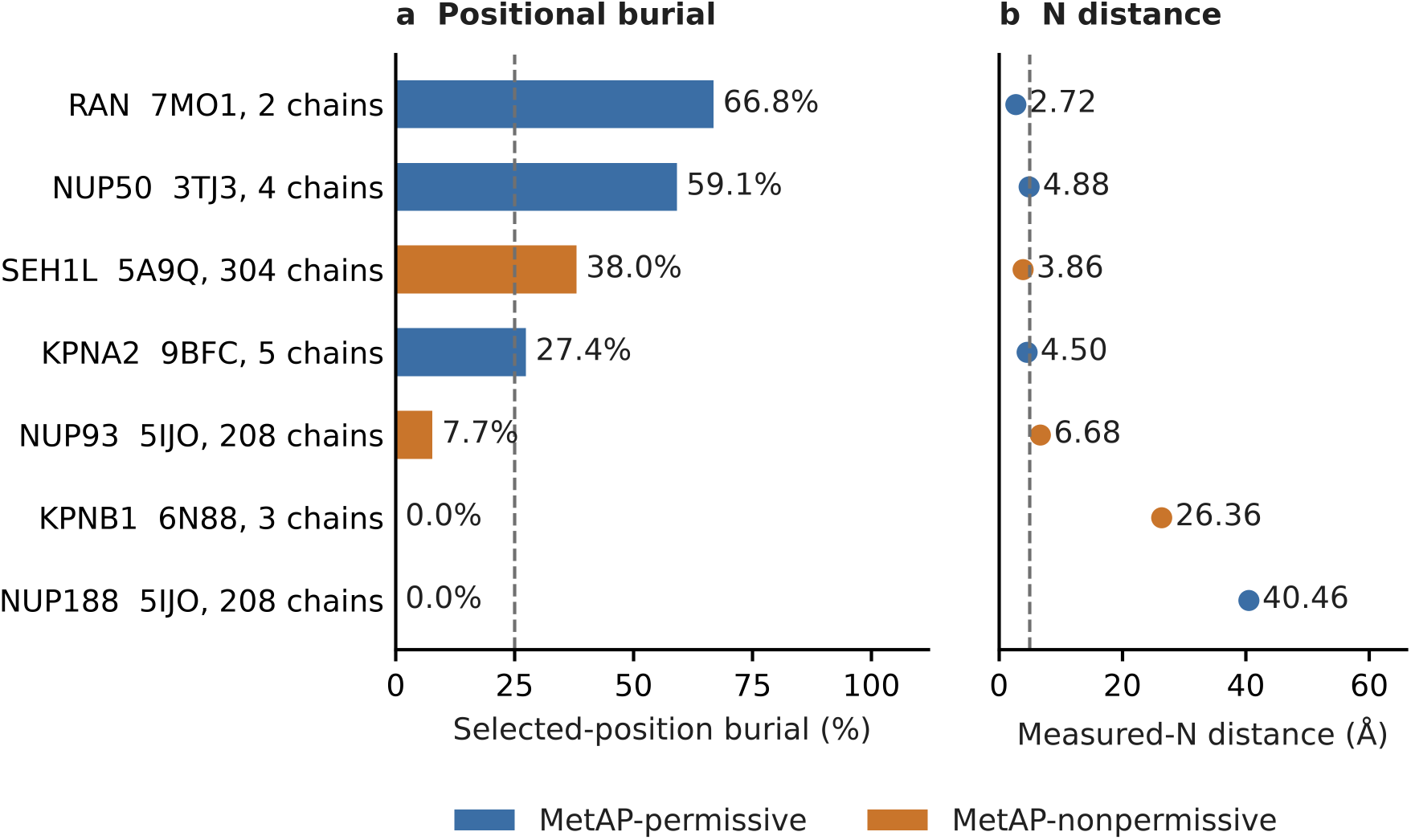
Geometry of selected N-terminal sequence positions in nuclear transport proteins. a, Burial of the annotation-selected residue for nuclear-pore-associated proteins and soluble transport factors in biological assemblies, with entry and chain count. b, Distance from that residue’s N atom to the nearest protein atom in another chain. Each row uses the highest-burial observation for that protein in the indicated entry; burial and distance come from that same observation. Blue and orange indicate permissive and nonpermissive residue-2 classes, respectively. Dashed lines mark descriptive thresholds of 25% burial and 5 Å; values are individual structural measurements, not replicate means. These measurements do not establish that the N atom is a free terminal amino group. In particular, Ran Ala2 in 7MO1 is internal to modeled Ser0–Met1–Ala2 and represents construct-dependent geometry.

The soluble-carrier observations further distinguish positional geometry from processing-dependent function. Importin-β has exposed modeled Met1 positions, whereas selected KPNA2 and Ran positions engage partner interfaces in the original extraction. However, Ran’s Ala2 in 7MO1 is internal to an engineered extension, so this comparison does not establish a native processed-terminal contact or predict a specific transport phenotype. Matched-proteoform experiments should measure processing and acetylation directly and separate altered geometry from changes in modification, partner binding and turnover. Co-translational assembly of nucleoporins provides additional context [18], but does not establish the chemical state of the individual deposited termini.

### Conditional-degron potential and interface engagement

MetAP-permissive termini whose retention would create a Met-Φ degron are not less interface-engaged than the remainder: among 696 MetAP-permissive proteins, the 352 degron-forming candidates have a mean burial of 26.3% and 38.6% at or above 25%, versus 24.4% and 36.3% for the other 344 (odds ratio 1.10, 95% confidence interval 0.81–1.50; Fisher p = 0.58; Mann– Whitney p = 0.61), and the class-by-degron interaction is null after assembly-size adjustment (likelihood-ratio p = 0.89). Interface engagement and conditional-degron potential should therefore be treated as separable candidate mechanisms.

### Downstream-modification annotations identify contrasting interface environments

Initiator-methionine removal can enable subsequent modification of the exposed residue. N-myristoyltransferase requires a free N-terminal glycine, and the N-terminal methyltransferases NTMT1 and NTMT2 recognize the Xaa-Pro-Lys/Arg motif [19]. Grouping the 696 permissive-class proteins by accession-level annotations and sequence motifs identifies contrasting interface environments (Supplementary Fig. 2); these flags do not establish modification in a structural preparation.

Among 18 proteins with annotated N-myristoylglycine, median burial was 59.1% and 55.6% reached at least 25% burial; the median was 1.55% among the 390 proteins without any of the three display flags (Supplementary Fig. 2). Within the 73 proteins with annotation-selected glycine, the 17 myristoylation-annotated proteins had a median burial of 70.0% versus 0.0% for the other 56 (Mann–Whitney p = 0.0017), an engagement odds ratio of 5.24 (Fisher p = 0.0059) and a shorter median measured-residue N distance (3.42 versus 8.74 Å, p = 0.0028). Annotation bias or docking of acylated termini could contribute. Four of six strict-motif candidates reached the burial threshold (Fisher p = 0.20), compared with five of seven under the relaxed Lys/Arg rule (p = 0.11). These exploratory patterns do not establish methylation effects. Annotated N-terminal acetylation showed no positional association (median 0.0% versus 2.55%, p = 0.67).

## Discussion

Where might initiator methionine excision influence human protein complex assembly? This census connects that question to identifiable N-terminal positions, partners and assembly contexts. MetAP-permissive classes are less often interface-engaged overall, yet 175 permissive-class proteins retain substantial engagement after observed-boundary filtering. The biological opportunity lies in this heterogeneity: particular interfaces, rather than every permissive terminus, may be sensitive to processing. State dependence here describes differences among deposited contexts, not a measured temporal transition.

PSMA7, PSMA6 and PSMA4 translate this candidate set into distinct questions about neighboring-subunit contacts, assembly-chaperone association and multi-partner packing (Figs. 3–4). Candidate priority should integrate terminal evidence, model quality and informative structural contrasts. The experimental endpoint is a change in a specified partner interaction linked to assembly or function, with terminal chemistry verified in the relevant preparation.

Does PSMA7 terminal chemistry influence its compatibility with different neighboring subunits? Complete Ser2 models occupy PSMA2-facing and PSMA5-facing environments, providing a same-position comparison. Testing assembly contexts at verified, constant terminal chemistry, followed by terminal perturbation within a fixed context, would separate these variables. Measurements of partner association, assembly and catalytic output, with abundance and protein-integrity controls, could then identify a processing-sensitive interface.

Does terminal processing influence PSMA6 association with PSMG1 versus neighboring α subunits? The PSMG1-facing Met1 backbone and the α-subunit-facing half-proteasome configuration define a specific comparison. Verified terminal proteoforms, partner occupancy and assembly-intermediate measurements could test whether processing changes either interaction. The incomplete pre-13S Met1 side chain makes the backbone contrast the starting evidence, rather than a demonstrated chaperone handoff.

Can a defined terminal perturbation redistribute PSMA4 contacts across the α ring? Its nearest partner need not contribute the greatest surface occlusion. The contrast between low pre-13S backbone occlusion and extensively packed Ser2 environments motivates measuring several partner contacts together. Atom completeness and local density should be assessed alongside these nonadditive geometric measurements before testing their relationship to assembly.

Which other recognition or assembly events warrant direct tests? The boundary-qualified histone-symbol association points to terminal-motif recognition in peptide-binding contexts. CCT and OXPHOS nominate subunit- and compartment-specific questions, while proteasome enrichment depends on boundary and multiplicity choices and the curated spliceosomal union is not enriched (Supplementary Tables 3–4). Thus, candidate-specific interfaces can remain informative without a complex-wide association; neither level alone establishes processing dependence.

When during biogenesis could these interfaces become processing-sensitive? Co-translational assembly provides a relevant time window [20,21]. Independent proteasome work places α-subunit N-terminal regions at chaperone and neighboring-subunit interfaces and shows that regional deletion can impair maturation [22]. Matched-proteoform experiments could determine whether retention of one methionine or altered acetylation affects these events, separating terminal chemistry from the broader effects of regional deletion.

These questions also motivate separate tests in other processing systems. Escherichia coli map essentiality [23], bacterial processing measurements [24], a MetAP-linked photosystem phenotype [25] and HslV terminal-threonine function [26,27] provide precedents, not evidence for a universal assembly-block mechanism. The eleven mitochondrially encoded proteins likewise require a separate processing framework; their exclusion preserves the primary human association.

Could a verified processing-dependent assembly defect alter proteostasis or drug response? PSMG2/PAC2 variants associated with impaired proteasome assembly and activity in CANDLE/PRAAS4 [28] and exosomal PSMA3/PSMA3-AS1 signaling linked to proteasome-inhibitor resistance in multiple myeloma [29] identify relevant functional settings, although neither study tests terminal processing and the latter concerns abundance regulation. Establishing a terminal perturbation-to-interaction-to-assembly connection would justify subsequent inflammatory or drug-response assays with abundance and fitness controls. PSMA3 itself requires boundary validation. Disease effects remain prospective endpoints.

Several evidence distinctions guide these experiments. Sequence class, processing annotation and coordinate identity are separate layers; absent removal annotation does not establish retention. Using sequence predictions as processing measurements would be circular. Correlated residue-2 chemistry also prevents volume and hydropathy models from isolating a processing effect (Supplementary Data 4). Direct proteoform measurements and controlled perturbations are therefore essential.

The atlas also depends on deposited assembly assignments, resolved positions and uneven structural sampling. Maximum burial describes an observed extreme, and minimum N-contact distance may come from another entry. Alternative aggregation rules cannot recover terminal states omitted by the selection rule. RNA, DNA and nonprotein-ligand occlusion are excluded, so protein-interface exposure leaves other interactions unmeasured; the direction of resulting bias remains unknown.

First-observed filtering is a coordinate criterion, not proof of a native terminus: upstream sequence may be unresolved or engineered away, and the residue may be capped. Ran and proteasome examples highlight the need to inspect constructs, internal peptide bonds and alternative mature products. Mapped spans do not establish coordinate completeness. These uncertainties remain explicit evidence flags.

Independent proteoform evidence is not preparation-matched, and five of the 22 focal residues have incomplete side chains. Backbone comparisons cannot recover missing interactions; construct differences and model quality remain alternatives, and original-map fit and local resolution were not validated. Static Met deletion likewise does not reconstruct a relaxed, acetylated terminus.

The principal advance is a partner-specific framework for testing how methionine excision contributes to complex assembly. The proteasome comparisons identify defined interfaces, and the atlas extends candidate selection beyond this system. Verifying terminal chemistry, perturbing a specified target and measuring partner association, assembly intermediates, mature complexes and function could reveal selective processing requirements or explain how some interfaces accommodate both terminal states.

## Methods

### N-terminal annotation

The frozen human UniProtKB table was parsed for initiator-methionine, modified-residue, signal-peptide, propeptide and transit-peptide features, together with sequence, length and gene name. Canonical position 2 was selected when the initiator-methionine feature contained “Removed”; position 1 was selected otherwise. This operational selection rule does not interpret an absent feature as positive evidence of retention. N-terminal acetylation annotations at positions 1 or 2 were retained as a separate evidence dimension, and “Removed; alternate” qualifiers were flagged in the audit rather than treated as evidence of an exclusively processed proteoform. Entries shorter than 30 residues were excluded in the original enumeration. Multiple annotated chain products were flagged for product-level review rather than automatically assigned a single mature proteoform.

### Target enumeration and structure selection

PDB-to-UniProt mappings were taken from the SIFTS pdb_chain_uniprot chain-segment table. Segment endpoints were used to derive candidate author residue numbers; this procedure is not equivalent to validating a complete residue-level alignment. A segment was retained when its UniProt start was at or before the annotation-selected position. The original target space comprised 4,087 accessions, 19,064 multi-chain entries and 101,894 chain targets, and entries were processed in order of their contribution of not-yet-observed accessions until distinct-protein coverage saturated.

### Biological assembly reconstruction

For each entry, the mmCIF file was retrieved from the RCSB Protein Data Bank, and the first annotated biological assembly was generated with gemmi version 0.7.5 [30], applying the deposited symmetry and transformation operators and renaming chains to preserve provenance. Only the first model was used; ligands and waters were removed, so the retained atoms are those of the polymer chains, and selenomethionine was treated as methionine when matching residue identity while keeping its own selenium radius in the surface calculation; hydrogens and non-primary alternate conformations were discarded. Assemblies with fewer than two chains after reconstruction, or exceeding 600,000 atoms, were skipped in the main run; a targeted set of large machines was re-analyzed with the ceiling raised to 1.5 million atoms.

### Identification and boundary audit of the selected residue

The original extraction derived candidate author residue numbers from SIFTS segment offsets and tested offsets in the order 0, +1, −1, +2, −2, accepting the first residue whose identity matched the canonical sequence at the selected position. The first-observed flag compared the accepted residue number with the minimum numbered amino-acid residue in the chain; it is a coordinate flag, not a verified biological boundary. The audit reconciled this arithmetic for all exported observations. Five analyzable observations required a +1 shift, affecting ACTG1, MRPL13 and GSTP1, and remain flagged for full residue-mapping review. For the 72 analyzable observations represented in the 11 frozen conformer-scan assemblies, modeled upstream residues and the distance from the preceding carbonyl C to the selected N were inspected directly. A C–N distance of 1.0–1.8 Å was used as a diagnostic of peptide-bond geometry, not as a substitute for full proteoform validation.

### Geometry

Solvent-accessible surface area was calculated with FreeSASA (version 2.2.1) [31] for the protein atoms of the complete assembly and each chain in isolation, using the radii implemented in the deposited census code: C 1.70, N 1.55, O 1.52, S 1.80, Se 1.80 and P 1.80 Å, with 1.70 Å as the fallback. Burial was 100 × (Aisolated − Aassembly) / Aisolated for the selected residue; nonpositive isolated areas were excluded. Nearest-other-chain distances were computed for any atom of that residue and for its N atom. The latter was interpreted as a backbone-N contact unless terminal position and chemical state were independently supported. Focal recalculations used the already expanded frozen assemblies without a second assembly transformation. Met1 and Ser2 measurements in unchanged models were kept distinct; no static deletion result was substituted for a validated mature-proteoform measurement.

Evidence-aware focal comparison. Figure 3b was assembled from the completed focal geometry audit by retaining unchanged deposited-model measurements of PSMA4, PSMA6 and PSMA7 at Met1 or Ser2 with zero modeled upstream amino-acid residues. All 22 qualifying records were displayed without statistical pooling, including repeated chains from the same entry. Entry context labels were read from the frozen mmCIF structure titles. Independent processing evidence was matched by UniProt accession and was not assigned to individual structural preparations. The displayed full-assembly measurements were retained, and a subsequent partner-resolved analysis extended this fixed subset without complete re-enumeration of proteasome states (Fig. 4; Supplementary Fig. 3; Supplementary Data 7). The comparison remains a selected audit subset, not an exhaustive structural survey.

Partner-resolved geometry. The seven frozen, already-expanded assembly files were verified against their original SHA256 manifest and analyzed without a second assembly transformation. Target and partner identities were assigned from the frozen SIFTS chain mapping and human-protein annotation. Minimum heavy-atom distances were measured for all 540 target–other-protein-chain pairs, with atom-pair counts at 3.5, 4 and 5 Å; 4 Å defined the descriptive contact threshold. Using the same protein-only atom selection and radii as the full-assembly calculation, pair-only burial was 100 × (A₀ − Aₚ)/A₀, where A₀ is the resolved target-residue SASA in its isolated chain and Aₚ is its SASA with one partner chain present. Pair SASA was calculated for 78 pairs approaching within 8 Å; more distant pairs were explicitly marked as not calculated and assigned zero occlusion under this conservative geometric exclusion. Pair-only values are nonadditive because partners can occlude overlapping surface and are not fractions of total assembly burial. No inferential tests were applied to this selected structural subset.

Focal atom-completeness sensitivity. Expected Met and Ser heavy atoms were checked in each target. OG was absent from PSMA6 Ser2 in 4R3O/A and 4R3O/O, PSMA4 Ser2 in 8TM4/B and PSMA7 Ser2 in 8TM4/C; CG, SD and CE were absent from PSMA6 Met1 in 8TM4/G. All four backbone atoms, N, CA, C and O, were present in every target. Backbone-only burial summarized their SASA from the unchanged coordinate calculations, and backbone partner distances were measured separately. No side chains were deleted or reconstructed for this sensitivity analysis. Distances denote geometric proximity, not validated bonds or favorable interaction energies; no original-map fit or local-resolution assessment was performed.

### Downstream modification branches

Lipidation and modified-residue features at positions 1 and 2 were taken from the same UniProtKB retrieval and used to flag annotated N-myristoylglycine and annotated N^α^-methylation. Substrates of the N-terminal methyltransferases were assigned by the strict canonical recognition motif applied to the mature sequence, requiring alanine, proline, serine, or glycine at mature position 1, proline at position 2, and lysine at position 3. NTMT1 and NTMT2, in fact, accept arginine as well as lysine at position 3 — the motif is Xaa-Pro-Lys/Arg, and human CENP-A and Drosophila H2B are the structurally characterized arginine-bearing substrates [19] — and at position 1 they tolerate any residue other than aspartate or glutamate, with a wide range of residues accepted on synthetic peptides in vitro [32,33], so the strict motif is conservative and undercounts the branch; the effect of relaxing it to lysine or arginine is reported in the results. One physiological consequence of the strict form is that CENP-A, the one known methylated protein carrying an arginine at position 3, is excluded by definition, and in any case contributes no analyzable chain because its N-terminal tail is not modeled in deposited nucleosome assemblies. Branch comparisons were confined to proteins classified as MetAP-permissive substrates.

### Classification

Proteins whose mature terminus is generated by signal-peptide, propeptide, or transit-peptide cleavage were excluded because their observed N-termini do not report initiator-methionine processing. Twenty immunoglobulin and T-cell receptor constant-region entries were excluded on the same principle: their deposited N-termini are variable–constant domain boundaries generated by V(D)J recombination rather than translation starts, so an initiator-methionine model does not apply. These entries are flagged in Supplementary Data 1 and are absent from every downstream analysis. The sequence rule classified alanine, cysteine, glycine, proline, serine, threonine and valine at residue 2 as the canonical small, uncharged MetAP-permissive set; all other residue-2 identities were classified as MetAP-nonpermissive for the primary analysis. This classification describes predicted biochemical permissiveness and was not interpreted as proof that iMet was removed in the specific molecule represented by a structure. Met-Φ degron potential on retention was assigned after leucine, phenylalanine, tyrosine, tryptophan, isoleucine, valine or alanine [5], the hydrophobic residue-2 set recognized by the UBR-box N-degron machinery; because the first five of these residues are MetAP-nonpermissive, degron-forming MetAP-permissive proteins are by construction those with alanine or valine at residue 2.

### Statistics and reproducibility

Two-sided Fisher exact and two-sided Mann–Whitney U tests were used for the primary sequence-class comparisons. The primary logistic model used burial ≥25% as outcome, with sequence class and natural-log chain count as predictors; confidence intervals were Wald intervals. Interaction models were compared by likelihood-ratio tests. Alternative thresholds, collapse rules, strata and modification-branch comparisons were exploratory and reported with unadjusted p values, except for the explicitly corrected enrichment families described below. Models containing penultimate side-chain volume quantified confounding between residue chemistry and sequence class. Deposition-multiplicity sensitivity added natural-log PDB-entry count. Compositional robustness excluded each residue-2 identity in turn. Other sensitivities excluded permissive proteins modeled as MET/MSE, required at least two or five PDB entries or two chain observations, considered first-observed status, excluded histones or mitochondrially encoded proteins, or restricted to agreement between sequence class and the exported processing annotation. Alternative sequence classes reassigned cysteine as nonpermissive or added Asn/Gln, Asp/Asn or Asp/Asn/Gln without changing the canonical primary set. The continuous minimum-N-distance comparison excluded five nonfinite exported sentinels, using 692 permissive and 426 nonpermissive proteins. Current-cohort calculations are separated from supplied chemistry-model estimates at exported precision in Supplementary Data 4–5. Analyses used a fixed structural corpus with 1,123 analyzable proteins and 19,967 analyzable chain observations; there were no experimental replicates. Corpus-size sampling used 40 orderings and seed 20250823. The original release, targeted audit and enrichment reconstruction are distinct provenance layers.

### Observed-boundary sensitivity

The audit retained the original 1,123-protein eligibility definition, removed observations whose selected residue was not first observed, and reselected each protein’s maximum burial among remaining observations. Proteins with no eligible observation were omitted from this sensitivity, not classified as unengaged. The original selected record was preserved in maximum-burial ties when still eligible; otherwise original observation-table order broke ties. The retained set contained 845 proteins. Two-sided Fisher tests compared engagement at ≥25% burial, and logistic regression used sequence class and natural-log chain count. The first-observed criterion is a coordinate-boundary sensitivity, not evidence that unresolved upstream sequence is absent or that the N atom is unmodified. Focal recalculations used protein-only atom selection and the original census radii.

### Cross-observation recurrence analysis

The original 261-protein candidate set was recovered by accession from the original observation table, and engagement fractions were calculated using burial ≥25%. Descriptive bins were ≥75%, 25% to <75%, and <25% of extracted observations engaged. Close-contact recurrence additionally used a measured-residue N distance ≤5 Å. PDB-level median collapsing, leave-one-entry-out stability and alternative burial thresholds addressed sampling within this fixed extraction. These outputs describe recurrence of selected modeled positions, not biological persistence across all processing states. They were not recomputed under the boundary-qualified eligibility rule in the present targeted audit.

### Methionine conformer scan and target audit

The legacy scan is not part of the current figure set or inferential results. The audit inspected target selection for all 53 positions in 11 frozen assemblies rather than rerunning its conformer sweep. For controls beginning with MET/MSE, the original code selected the next modeled amino acid and excluded the first Met; otherwise it fell back to the first modeled amino acid. This fallback selects engineered Ser0 instead of Ala2 for Ran in 7MO1. Its frozen result is unscored, and a corrected scan has not been run. Target-selection diagnostics and the frozen implementation are retained as historical provenance in Supplementary Data 8. Explicit canonical-residue mapping and validated upstream-atom exclusion are required before reuse.

### Software

The original repository version-pins the census and legacy conformer-scan environments. The targeted audit used Python 3.14.3, gemmi 0.7.5, NumPy 2.5.3, pandas 3.0.6, SciPy 1.18.1, statsmodels 0.15.0 and FreeSASA 2.2.1. The enrichment reconstruction used the same Python, NumPy, pandas, SciPy and statsmodels versions. Figures were rendered from the supplied source tables using Matplotlib 3.11.2 (https://matplotlib.org/); the current presentation code and its inputs are included in Supplementary Data 8. At audit intake, the six original workbooks matched their frozen-release counterparts; the revised Supplementary Data 1–6 include corrected labels and current-cohort results and are not represented as byte-identical originals. Eleven frozen assembly checksums matched the release manifest, and six previously exported focal measurements reproduced at their reported precision. The enrichment outputs passed 56 numerical/membership checks, including independent enumeration of two-sided Fisher probabilities and verification of BH/BY corrections, and reproduced exactly on rerun. These checks establish input consistency and numerical reproducibility, not native chemistry or independence of homologous proteins. SciPy-based tests follow the algorithms described by Virtanen et al. [34].

### Prefix grouping and reconstructed enrichment

Figure 2 reuses the ordered gene-symbol assignment function in the frozen figure code; twelve groups had at least four scored proteins in the original 1,123-protein cohort. Prefix membership is not equivalent to curated complex membership and can omit or include different proteins. A new thirteen-test family retained those twelve groups and added a histone-symbol group matching HIST or H1/H2/H3/H4-style symbols; the exact regular expression and all accession assignments are supplied in Supplementary Data 9. The original twelve-test correction without histones is a labeled sensitivity only. Enrichment used one record per UniProt accession, a primary burial threshold of 25%, and the table [[member engaged, member unengaged], [nonmember engaged, nonmember unengaged]]. Backgrounds were 696 permissive proteins originally and 463 after first-observed filtering and maximum reselection. Missing eligibility was not counted as non-engagement. These are new exploratory analyses, not recovery of the undocumented historical twelve-curated-comparison analysis.

### Curated memberships and focused questions

The human Complex Portal ComplexTab file was retrieved on 25 September 2026 UTC (https://ftp.ebi.ac.uk/pub/databases/intact/complex/current/complextab/9606.tsv; curation documentation: https://www.ebi.ac.uk/complexportal/documentation). It contained 2,498 manually curated records; separately distributed machine-learning predictions were not used. Manual curation includes inferred evidence and does not imply direct experimental support for every record. Expanded participant lists were mapped to canonical UniProt accessions, preserving suffix normalization and alternative-set membership in an audit. A strict-identifier sensitivity excluded suffix-normalized and alternative-set members. Sixty-four complexes with at least four original-background permissive accessions defined an outcome-independent eligible family, retained for the boundary analysis. A sensitivity collapsed identical original tested accession sets to 42 comparisons.

Five focused exploratory questions used CPX-5993 for 26S proteasome, CPX-6030 for CCT/TRiC, the union of CPX-560/561/577/6123/6151 for OXPHOS I–V, all seventeen recommended names containing “spliceosom” for spliceosomal complexes, and the eight names beginning “Nucleosome” or “CENP-A nucleosome” for core-histone membership. These targets were selected from the biological questions already discussed, not as an independent prospective validation. The focused family does not override the broader screen. Catalogue membership does not determine the structural context supplying an accession’s selected maximum. Nuclear-versus mitochondrial-encoded OXPHOS comparisons were post-result diagnostics corrected across two subgroups; a prefix-histone stratified diagnostic was also labeled separately.

### Multiplicity and sampling sensitivities

Two-sided Fisher tests were accompanied by sample odds ratios and exact conditional odds-ratio confidence intervals. Benjamini–Hochberg q values were calculated separately for each declared family, cohort and threshold; Benjamini–Yekutieli correction was a dependence-conservative sensitivity. Zero member cells can yield infinite sample odds ratios and broad intervals. Focused-family thresholds of 20% and 30% were secondary sensitivities, not substitutes for the 25% endpoint. Exact one-sided upper-tail enrichment tests conditioned on fixed margins within selected-assembly chain-count strata (2–3, 4–5, 6–11 and ≥12) crossed with PDB-entry-count strata (one, two to four and ≥five).

Hypergeometric distributions were convolved exactly; deposition counts in the boundary analysis used eligible observations only. BH correction covered all five focused stratified tests within each cohort. These adjustments address measured sampling covariates, not native chemistry, homolog independence or ascertainment of unresolved termini. Full results and definitions appear in the Supplementary Note, “Reconstructed enrichment”, Supplementary Tables 3–4 and Supplementary Data 9.

### Use of artificial intelligence tools

ChatGPT (OpenAI) and Perplexity Computer (Perplexity AI) were used for manuscript organization, language editing, literature retrieval and synthesis, scientific critique, and assistance in drafting analysis and visualization code; Grammarly was used for language and copy editing. These uses included evaluative and interpretive assistance but did not replace the author’s scholarly judgment. The author independently reviewed and revised all AI-assisted text, verified cited sources, inspected analytical logic and outputs, and retained control of the study design, analyses, interpretation and conclusions. No AI-generated output was accepted as evidence without verification against the underlying data or cited source. The author is solely accountable for the originality, accuracy and integrity of the work. AI tools were not listed as authors.

## Data availability

The corrected datasets and reproducibility materials are publicly available in Zenodo version 1.4.0 (https://doi.org/10.5281/zenodo.22966146; record: https://zenodo.org/records/22966146) [35]. Supplementary Data 1–2 provide the frozen per-protein and per-observation measurements with corrected evidence labels and processing, coordinate-boundary and mapping flags. Supplementary Data 3 separates the original 261 model-selected candidates from the 175 permissive-class candidates retained by the first-observed sensitivity. Supplementary Data 4–5 contain current-cohort statistical and robustness results, with supplied chemistry-model estimates identified separately, and an evidence-aware candidate-to-experiment map. Supplementary Data 6 describes conditional recurrence within the original extraction-selected candidate set.

Supplementary Data 8 supplies audit results, figure source values, current CSV exports, frozen inputs, analysis code and labeled historical provenance in two archives (8a and 8b), which must be extracted into the same directory. Supplementary Data 9 contains the enrichment reconstruction: frozen curated catalogue, memberships and mapping audit, complete test tables, supplementary-table source values, a browsable workbook, code, environment requirements and verification records. Historical release 1.3.0 is retained as provenance, not as a corrected record of all native proteoforms.

Supplementary Data 7. Partner-resolved PSMA4, PSMA6 and PSMA7 geometry for the fixed 22-observation subset. The accompanying ZIP contains observation and partner-chain tables, all atom-pair contacts within 5 Å, common-backbone sensitivities, chain-identity and input-hash checks, both contact-map renderings, reproducible code and seven frozen assembly coordinate files. Pair-only burial is nonadditive; missing focal atoms and excluded pair-SASA calculations are explicitly flagged. This addition does not replace Supplementary Data 1–6 or imply a complete proteasome survey.

## Code availability

The corrected analysis code, frozen inputs, derived tables, figure scripts and verification records are available in Zenodo version 1.4.0, https://doi.org/10.5281/zenodo.22966146 [35]. Partner-resolved geometry, terminal-boundary auditing and enrichment reconstruction are also supplied in Supplementary Data 7, 8 and 9, respectively. Historical release 1.3.0 (https://doi.org/10.5281/zenodo.22847917) preserves the original model-based analysis, including the legacy conformer-scan target-selection limitation, and is not the corrected audit release. The version-family concept DOI is https://doi.org/10.5281/zenodo.22129640.

## Acknowledgements

The author acknowledges the UniProt Consortium, PDBe/SIFTS and the RCSB Protein Data Bank for maintaining the public resources on which this study depends. The figures were prepared by the author from the source data described in Methods.

## Author contributions

Y.-H.C. conceived the study, designed and performed the analyses, interpreted the results, prepared the figures, and wrote the manuscript.

## Additional information

