## Supplementary_Data_7 for "Human protein complex interfaces reveal candidate assembly roles for initiator methionine excision": asm_lib.py

```
import os as _os
BASE = _os.environ.get("NME_DIR") or _os.path.dirname(_os.path.dirname(_os.path.abspath(__file__)))
INPUT_DIR = _os.path.join(BASE, "inputs")
WORK_DIR = _os.environ.get("NME_WORK") or _os.path.join(BASE, "work")
_os.makedirs(WORK_DIR, exist_ok=True)
"""Biological-assembly N-terminal geometry pipeline.

For each PDB entry: download mmCIF, expand biological assembly 1, then for each
target human chain measure (i) the fraction of the mature N-terminal residue's
solvent-accessible surface that is buried by the rest of the assembly and
(ii) the distance from that residue, and from its alpha-amino nitrogen, to the
nearest atom of any other chain in the assembly.
"""
import os, re, gzip, io, urllib.request, collections
import numpy as np
import gemmi, freesasa
from scipy.spatial import cKDTree

freesasa.setVerbosity(freesasa.silent)

AA3 = {'ALA':'A','ARG':'R','ASN':'N','ASP':'D','CYS':'C','GLN':'Q','GLU':'E','GLY':'G','HIS':'H',
       'ILE':'I','LEU':'L','LYS':'K','MET':'M','PHE':'F','PRO':'P','SER':'S','THR':'T','TRP':'W',
       'TYR':'Y','VAL':'V','MSE':'M','SEC':'U','PYL':'O'}

# ProtOr-style united-atom radii used by FreeSASA's default classifier, keyed by element.
RADII = {'C': 1.70, 'N': 1.55, 'O': 1.52, 'S': 1.80, 'SE': 1.80, 'P': 1.80}
DEFAULT_R = 1.70

CIF_DIR = f"{WORK_DIR}/cif"
MAX_ATOMS = 600000


def fetch_cif(pdb, keep=False):
    os.makedirs(CIF_DIR, exist_ok=True)
    path = f"{CIF_DIR}/{pdb}.cif"
    if os.path.exists(path) and os.path.getsize(path) > 500:
        return path
    url = f"https://files.rcsb.org/download/{pdb.upper()}.cif.gz"
    raw = urllib.request.urlopen(url, timeout=90).read()
    txt = gzip.decompress(raw)
    with open(path, "wb") as fh:
        fh.write(txt)
    return path


def load_assembly(path):
    """Return (structure_expanded, orig_name_map) for biological assembly 1.

    orig_name_map maps the expanded chain name back to the author chain name of
    the asymmetric unit, so SIFTS chain identifiers still resolve.
    """
    st = gemmi.read_structure(path)
    st.setup_entities()
    st.remove_alternative_conformations()
    st.remove_hydrogens()
    st.remove_ligands_and_waters()
    orig = {ch.name for ch in st[0]}
    asm_id = None
    for a in st.assemblies:
        asm_id = a.name
        break
    if asm_id is not None:
        try:
            st.transform_to_assembly(asm_id, gemmi.HowToNameCopiedChain.AddNumber)
        except Exception:
            pass
    # map expanded chain name -> original author chain name
    name_map = {}
    for ch in st[0]:
        n = ch.name
        if n in orig:
            name_map[n] = n
        else:
            base = re.sub(r'\d+$', '', n)
            name_map[n] = base if base in orig else n
    return st, name_map


def chain_atom_table(st):
    """Flatten model 0 into arrays. Returns dict with coords, radii, chain idx,
    and a per-chain residue index for SASA bookkeeping."""
    coords, radii, chain_ids, res_keys = [], [], [], []
    chains = []
    for ci, ch in enumerate(st[0]):
        chains.append(ch.name)
        for res in ch:
            if res.name not in AA3:
                continue
            key = (ci, res.seqid.num, res.seqid.icode)
            for at in res:
                el = at.element.name.upper()
                coords.append([at.pos.x, at.pos.y, at.pos.z])
                radii.append(RADII.get(el, DEFAULT_R))
                chain_ids.append(ci)
                res_keys.append(key)
    if not coords:
        return None
    return dict(xyz=np.asarray(coords, dtype=float),
                r=np.asarray(radii, dtype=float),
                cid=np.asarray(chain_ids, dtype=int),
                rkey=res_keys, chains=chains)


def sasa(xyz, radii):
    res = freesasa.calcCoord(xyz.ravel().tolist(), radii.tolist())
    return np.array([res.atomArea(i) for i in range(len(radii))])


def residue_areas(atom_areas, rkey):
    out = collections.defaultdict(float)
    for a, k in zip(atom_areas, rkey):
        out[k] += a
    return out


def analyze_entry(pdb, targets, prot, seqs, segs, cleanup=True):
    """targets: list of dicts with chain/acc for this entry."""
    try:
        path = fetch_cif(pdb)
    except Exception as e:
        return [], f"download_fail:{type(e).__name__}"
    try:
        st, name_map = load_assembly(path)
        T = chain_atom_table(st)
        if T is None:
            return [], "no_atoms"
        if len(T['chains']) < 2:
            return [], "monomeric_assembly"
        if len(T['r']) > MAX_ATOMS:
            return [], "too_big"

        area_cx = residue_areas(sasa(T['xyz'], T['r']), T['rkey'])
        # isolated-chain areas, computed per chain on demand
        iso_cache = {}

        def iso_areas(ci):
            if ci not in iso_cache:
                m = T['cid'] == ci
                iso_cache[ci] = residue_areas(sasa(T['xyz'][m], T['r'][m]),
                                              [k for k, mm in zip(T['rkey'], m) if mm])
            return iso_cache[ci]

        # residues per chain for lookup
        res_by_chain = collections.defaultdict(dict)
        for ci, ch in enumerate(st[0]):
            for res in ch:
                if res.name in AA3 and res.seqid.icode in (' ', '\0', ''):
                    res_by_chain[ci].setdefault(res.seqid.num, res)

        out = []
        by_chain_targets = collections.defaultdict(list)
        for t in targets:
            by_chain_targets[t['chain']].append(t)

        for ci, ch in enumerate(st[0]):
            orig_name = name_map.get(ch.name, ch.name)
            for t in by_chain_targets.get(orig_name, []):
                acc = t['acc']
                p = prot.get(acc)
                useq = seqs.get(acc)
                if not p or not useq:
                    continue
                ms = p['mature_start']
                expect = useq[ms - 1]
                cands = []
                for sp_beg, pdb_beg in segs.get((pdb, orig_name, acc), []):
                    if sp_beg > ms:
                        continue
                    m = re.match(r'^(-?\d+)', str(pdb_beg))
                    if m:
                        cands.append(int(m.group(1)) + (ms - sp_beg))
                hit = None
                rmap = res_by_chain[ci]
                for c in cands:
                    for off in (0, 1, -1, 2, -2):
                        r = rmap.get(c + off)
                        if r is not None and AA3.get(r.name) == expect:
                            hit = c + off
                            break
                    if hit is not None:
                        break
                if hit is None:
                    continue
                res = rmap[hit]
                key = (ci, res.seqid.num, res.seqid.icode)
                ac = area_cx.get(key)
                ai = iso_areas(ci).get(key)
                if ac is None or ai is None or ai <= 0:
                    continue
                buried = 100.0 * (1 - ac / ai)
                # distances to other chains
                other = T['cid'] != ci
                if not other.any():
                    continue
                tree = cKDTree(T['xyz'][other])
                ratoms = np.array([[a.pos.x, a.pos.y, a.pos.z] for a in res])
                d_res = float(tree.query(ratoms)[0].min())
                nat = [a for a in res if a.name == 'N']
                d_N = float(tree.query(np.array([[nat[0].pos.x, nat[0].pos.y, nat[0].pos.z]]))[0][0]) if nat else np.nan
                first_obs = min(rmap) if rmap else None
                out.append(dict(pdb=pdb, asm_chain=ch.name, chain=orig_name, acc=acc,
                                gene=p['gene'], n_chains=len(T['chains']),
                                n_entities=t.get('n_entities', ''),
                                mature_start=ms, init_removed=p['init_removed'],
                                res1=p['res1'], res2=p['res2'], res3=p['res3'],
                                nt_acetyl=p['nt_acetyl'], has_signal=p['has_signal'],
                                has_propep=p['has_propep'], has_transit=p['has_transit'],
                                resname=res.name, authnum=hit,
                                is_first_observed=(hit == first_obs),
                                sasa_complex=round(ac, 1), sasa_isolated=round(ai, 1),
                                buried_pct=round(buried, 1),
                                min_dist_residue=round(d_res, 2),
                                min_dist_alphaamino_N=(round(d_N, 2) if d_N == d_N else ''),
                                n_atoms=int(len(T['r']))))
        return out, "ok"
    except Exception as e:
        return [], f"error:{type(e).__name__}"
    finally:
        if cleanup:
            p = f"{CIF_DIR}/{pdb}.cif"
            if os.path.exists(p):
                os.remove(p)
```
