## Supplementary_Data_7 for "Human protein complex interfaces reveal candidate assembly roles for initiator methionine excision": partner_resolved_analysis.py

```
"""Partner-resolved analysis of the fixed 22-observation Figure 3 subset.
No assembly expansion, terminal modification, coordinate editing or p-value testing.
"""
from pathlib import Path
import os,sys,json,hashlib,platform
import pandas as pd
import numpy as np
import gemmi
from scipy.spatial.distance import cdist

ROOT=Path(__file__).resolve().parent
BASE=ROOT/"paper1_audit/frozen/nterm-census-1.3.0"
OUT=ROOT/"deliverables/paper1_partner_resolved"
OUT.mkdir(parents=True,exist_ok=True)
os.environ["NME_WORK"]=str(OUT/"scratch")
sys.path.insert(0,str(BASE/"code"))
import asm_lib as lib

f=pd.read_csv(ROOT/"deliverables/paper1_evidence_aware/Figure_3_Evidence_Aware_Source_Data.csv")
assert len(f)==22
pdbs=set(f.pdb.str.lower())
uni=pd.read_csv(BASE/"inputs/human_nterm.tsv.gz",sep="\t").fillna("")
gene_lookup=uni.set_index("Entry")["Gene Names (primary)"].to_dict()
segments=[]
for part in pd.read_csv(BASE/"inputs/pdb_chain_uniprot.tsv.gz",sep="\t",comment="#",
                        chunksize=200000,dtype=str):
    hit=part[part.PDB.isin(pdbs)]
    if len(hit):segments.append(hit)
sifts=pd.concat(segments,ignore_index=True)
sifts.to_csv(OUT/"frozen_sifts_segments_used.csv",index=False)
maps=sifts.groupby(["PDB","CHAIN"]).SP_PRIMARY.apply(lambda s:sorted(set(s))).to_dict()
checks=[];chains_out=[];pairs=[];contacts=[];summaries=[];hashes=[]

def category(genes):
    if not genes:return "unresolved"
    categories=set()
    for g in genes:
        if g.startswith("PSMA"):categories.add("20S alpha subunit")
        elif g.startswith("PSMB"):categories.add("20S beta subunit")
        elif g.startswith(("PSMG",)) or g=="POMP":categories.add("assembly chaperone")
        elif g.startswith(("PSMC","PSMD")):categories.add("19S regulatory subunit")
        elif g.startswith("PSME"):categories.add("proteasome activator")
        else:categories.add("other protein")
    return "; ".join(sorted(categories))

for pdb,targets in f.groupby("pdb",sort=True):
    path=BASE/f"inputs/figure5_assemblies/{pdb.lower()}.cif.gz"
    actual_hash=hashlib.sha256(path.read_bytes()).hexdigest()
    expected_hashes={Path(line.split()[1]).name:line.split()[0]
                     for line in (BASE/"inputs/figure5_assemblies/SHA256SUMS.txt").read_text().splitlines() if line.strip()}
    assert actual_hash==expected_hashes[path.name]
    hashes.append({"input":str(path.relative_to(ROOT)),"sha256":actual_hash,"matches_frozen_manifest":True})
    block=gemmi.cif.read(str(path)).sole_block()
    entities=block.get_mmcif_category("_entity.")
    descriptions=dict(zip(entities.get("id",[]),entities.get("pdbx_description",[])))
    ep=block.get_mmcif_category("_entity_poly.")
    chain_entity={}
    for eid,chs in zip(ep.get("entity_id",[]),ep.get("pdbx_strand_id",[])):
        for ch in chs.split(","):chain_entity[ch.strip()]=eid
    res=block.find_value("_em_3d_reconstruction.resolution") or block.find_value("_refine.ls_d_res_high") or ""
    title=gemmi.cif.as_string(block.find_value("_struct.title") or "")
    st=gemmi.read_structure(str(path))
    st.setup_entities();st.remove_alternative_conformations();st.remove_hydrogens();st.remove_ligands_and_waters()
    T=lib.chain_atom_table(st)
    # Match flattening order exactly; atom names/residue identity provide contact provenance.
    atoms=[]
    for ci,ch in enumerate(st[0]):
        for r in ch:
            if r.name not in lib.AA3:continue
            for a in r:
                atoms.append({"ci":ci,"chain":ch.name,"resnum":r.seqid.num,"icode":r.seqid.icode.strip(),
                              "resname":r.name,"atom":a.name,"occupancy":a.occ,"b_iso":a.b_iso})
    assert len(atoms)==len(T["xyz"])
    atoms=pd.DataFrame(atoms)
    chainmeta={}
    for ci,ch in enumerate(st[0]):
        accs=maps.get((pdb.lower(),ch.name),[])
        genes=[gene_lookup.get(a,"") for a in accs]
        genes=[x for x in genes if x]
        eid=chain_entity.get(ch.name,"")
        aa=[r for r in ch if r.name in lib.AA3]
        m={"pdb":pdb,"chain":ch.name,"entity_id":eid,"entity_description":descriptions.get(eid,""),
           "accessions":";".join(accs),"genes":";".join(genes),"partner_class":category(genes),
           "mapping_status":"unique_SIFTS" if len(accs)==1 else "ambiguous_SIFTS" if accs else "no_SIFTS",
           "modeled_residues":len(aa),"modeled_prefix":" ".join(f"{r.name}{r.seqid}" for r in aa[:5]),
           "entry_resolution_A":res,"entry_title":title}
        chainmeta[ci]=m;chains_out.append(m)
    fullarea=lib.sasa(T["xyz"],T["r"])
    iso_cache={}
    for t in targets.itertuples():
        ci=T["chains"].index(t.chain)
        assert chainmeta[ci]["genes"]==t.gene, (pdb,t.chain,chainmeta[ci])
        first=next(r for r in st[0][ci] if r.name in lib.AA3)
        assert first.seqid.num==t.measured_authnum and first.name==t.resname
        idx=np.where((T["cid"]==ci)&(atoms.resnum.values==t.measured_authnum)&
                     (atoms.resname.values==t.resname))[0]
        assert len(idx)>0
        assert len(set(atoms.iloc[idx].icode))==1
        mask=T["cid"]==ci
        if ci not in iso_cache:
            cidx=np.where(mask)[0]
            a=lib.sasa(T["xyz"][mask],T["r"][mask])
            iso_cache[ci]=dict(zip(cidx,a))
        ai=sum(iso_cache[ci][i] for i in idx);ac=fullarea[idx].sum()
        burial=100*(1-ac/ai)
        expected={"SER":{"N","CA","C","O","CB","OG"},"MET":{"N","CA","C","O","CB","CG","SD","CE"}}[t.resname]
        observed=set(atoms.iloc[idx].atom)
        missing=sorted(expected-observed)
        bidx=[i for i in idx if atoms.iloc[i].atom in {"N","CA","C","O"}]
        assert len(bidx)==4
        bai=sum(iso_cache[ci][i] for i in bidx)
        backbone_burial=100*(1-fullarea[bidx].sum()/bai) if bai>0 else np.nan
        nidx=[i for i in idx if atoms.iloc[i].atom=="N"]
        assert len(nidx)==1
        observation=f"{pdb}:{t.chain}:{t.resname}{t.measured_authnum}"
        target_fields={"observation":observation,"gene":t.gene,"pdb":pdb,"target_chain":t.chain,
                       "target_resname":t.resname,"target_authnum":t.measured_authnum,
                       "context":t.entry_context_from_title,"total_buried_pct":burial}
        local=[]
        for cj,ch in enumerate(T["chains"]):
            if cj==ci:continue
            pidx=np.where(T["cid"]==cj)[0]
            if not len(pidx):continue
            D=cdist(T["xyz"][idx],T["xyz"][pidx])
            a,b=np.unravel_index(D.argmin(),D.shape);dist=float(D[a,b])
            dn=float(np.linalg.norm(T["xyz"][pidx]-T["xyz"][nidx[0]],axis=1).min())
            meta=chainmeta[cj]
            closest=atoms.iloc[pidx[b]]
            pair_area=np.nan;pair_pct=0.0
            # Surface occlusion is impossible beyond this conservative geometric bound
            # for the original radii/probe. Mark omitted calculations, do not pretend exact.
            if dist<=8.0:
                keep=(T["cid"]==ci)|(T["cid"]==cj)
                keepidx=np.where(keep)[0]
                sasa=lib.sasa(T["xyz"][keep],T["r"][keep])
                lut=dict(zip(keepidx,sasa))
                pair_area=sum(lut[i] for i in idx)
                pair_pct=100*(1-pair_area/ai)
            row={**target_fields,"partner_chain":ch,"partner_genes":meta["genes"],
                 "partner_accessions":meta["accessions"],"partner_description":meta["entity_description"],
                 "partner_class":meta["partner_class"],"mapping_status":meta["mapping_status"],
                 "min_heavy_atom_distance_A":dist,"target_N_min_distance_A":dn,
                 "target_closest_atom":atoms.iloc[idx[a]].atom,
                 "partner_closest_residue":f"{closest.resname}{closest.resnum}{closest.icode}",
                 "partner_closest_atom":closest.atom,
                 "atom_pairs_le_3_5_A":int((D<=3.5).sum()),
                 "atom_pairs_le_4_A":int((D<=4).sum()),
                 "atom_pairs_le_5_A":int((D<=5).sum()),
                 "partner_residues_le_4_A":len(set((atoms.iloc[pidx[j]].resnum,atoms.iloc[pidx[j]].icode)
                                                       for j in np.where((D<=4).any(axis=0))[0])),
                 "pair_sasa_calculated":bool(dist<=8),"isolated_target_sasa_A2":ai,
                 "target_plus_partner_sasa_A2":pair_area,"pair_only_buried_pct":pair_pct,
                 "target_missing_heavy_atoms":",".join(missing),
                 "backbone_only_total_buried_pct":backbone_burial,
                 "backbone_pair_only_buried_pct":100*(1-sum(lut[i] for i in bidx)/bai) if dist<=8 and bai>0 else 0.0,
                 "backbone_min_distance_A":float(np.linalg.norm(T["xyz"][bidx,None,:]-T["xyz"][None,pidx,:],axis=2).min()),
                 "target_occupancy_min":float(atoms.iloc[idx].occupancy.min()),
                 "target_b_iso_mean":float(atoms.iloc[idx].b_iso.mean()),"entry_resolution_A":res}
            pairs.append(row);local.append(row)
            for ia,ib in zip(*np.where(D<=5)):
                ta=atoms.iloc[idx[ia]];pa=atoms.iloc[pidx[ib]]
                contacts.append({**target_fields,"partner_chain":ch,"partner_genes":meta["genes"],
                                 "partner_accessions":meta["accessions"],"target_atom":ta.atom,
                                 "partner_resname":pa.resname,"partner_authnum":pa.resnum,
                                 "partner_icode":pa.icode,"partner_atom":pa.atom,"distance_A":D[ia,ib]})
        closest=min(local,key=lambda r:r["min_heavy_atom_distance_A"])
        nearestN=min(local,key=lambda r:r["target_N_min_distance_A"])
        dominant=max(local,key=lambda r:r["pair_only_buried_pct"])
        significant=[r for r in local if r["atom_pairs_le_4_A"]>0]
        five=[r for r in local if r["atom_pairs_le_5_A"]>0]
        summary={**target_fields,"target_boundary":"first modeled amino acid; biological chemistry not validated",
                 "nearest_partner":closest["partner_genes"],"nearest_partner_chain":closest["partner_chain"],
                 "nearest_distance_A":closest["min_heavy_atom_distance_A"],
                 "nearest_N_partner":nearestN["partner_genes"],"nearest_N_chain":nearestN["partner_chain"],
                 "nearest_N_distance_A":nearestN["target_N_min_distance_A"],
                 "partners_le_4_A":"; ".join(r["partner_genes"]+":"+r["partner_chain"] for r in significant),
                 "partners_le_5_A":"; ".join(r["partner_genes"]+":"+r["partner_chain"] for r in five),
                 "largest_pair_occlusion_partner":dominant["partner_genes"] if dominant["pair_only_buried_pct"]>.01 else "",
                 "largest_pair_occlusion_chain":dominant["partner_chain"] if dominant["pair_only_buried_pct"]>.01 else "",
                 "largest_pair_only_buried_pct":dominant["pair_only_buried_pct"],
                 "entry_resolution_A":res,"target_atoms":len(idx),
                 "target_missing_heavy_atoms":",".join(missing),
                 "backbone_only_total_buried_pct":backbone_burial,
                 "backbone_partners_le_4_A":"; ".join(r["partner_genes"]+":"+r["partner_chain"] for r in local if r["backbone_min_distance_A"]<=4)}
        summaries.append(summary)
        checks.append({"observation":observation,"burial_error_pp":burial-t.buried_pct,
                       "min_distance_error_A":closest["min_heavy_atom_distance_A"]-t.min_residue_contact_A,
                       "N_distance_error_A":nearestN["target_N_min_distance_A"]-t.target_N_contact_A,
                       "target_gene_matches":chainmeta[ci]["genes"]==t.gene,
                       "max_pair_minus_total_pp":dominant["pair_only_buried_pct"]-burial})
    print("Finished",pdb,flush=True)

frames={"chain_identity_audit":chains_out,"all_partner_measurements":pairs,
        "atom_contacts_within_5A":contacts,"observation_partner_summary":summaries,
        "validation_checks":checks,"input_hashes":hashes}
for name,rows in frames.items():pd.DataFrame(rows).to_csv(OUT/f"{name}.csv",index=False)
c=pd.DataFrame(checks);p=pd.DataFrame(pairs)
assert c.burial_error_pp.abs().max()<1e-8
assert c.min_distance_error_A.abs().max()<1e-8
assert c.N_distance_error_A.abs().max()<1e-8
assert c.max_pair_minus_total_pp.max()<1e-5
contacting=p[p.atom_pairs_le_4_A>0]
assert (contacting.mapping_status=="unique_SIFTS").all()
result={"observations":len(summaries),"proteins":int(f.gene.nunique()),"entries":int(f.pdb.nunique()),
        "partner_chain_pairs":len(p),"contacting_chain_pairs_4A":len(contacting),
        "atom_pairs_5A":len(contacts),"pair_sasa_calculations":int(p.pair_sasa_calculated.sum()),
        "max_burial_error_pp":float(c.burial_error_pp.abs().max()),
        "max_contact_distance_error_A":float(c.min_distance_error_A.abs().max()),
        "all_contacting_partners_uniquely_mapped":True,
        "observations_with_missing_target_heavy_atoms":int(pd.DataFrame(summaries).target_missing_heavy_atoms.ne("").sum()),
        "python":platform.python_version(),"gemmi":gemmi.__version__,
        "scope":"fixed 22-observation Figure 3 subset; no inferential statistics; protein atoms only"}
(OUT/"analysis_summary.json").write_text(json.dumps(result,indent=2))
print(json.dumps(result,indent=2))
print(pd.DataFrame(summaries).to_string(index=False))
```
