## Supplementary_Data_7 for "Human protein complex interfaces reveal candidate assembly roles for initiator methionine excision": plot_partner_contactmap_print.py

```
"""Print-sized rendering of the verified 22-observation partner contact map."""
from pathlib import Path
import pandas as pd
import numpy as np
import matplotlib
matplotlib.use("Agg")
import matplotlib.pyplot as plt
from matplotlib.colors import LinearSegmentedColormap

ROOT=Path(__file__).resolve().parent
DATA=ROOT/"deliverables/paper1_partner_resolved"
OUT=ROOT/"deliverables/paper1_contactmap_integrated"
OUT.mkdir(parents=True,exist_ok=True)
s=pd.read_csv(DATA/"observation_partner_summary.csv").fillna("")
p=pd.read_csv(DATA/"all_partner_measurements.csv").fillna("")
s=s.sort_values(["gene","pdb","target_chain"]).reset_index(drop=True)
genes=["PSMA1","PSMA2","PSMA3","PSMA4","PSMA5","PSMA6","PSMA7","PSMG1","POMP"]
M=np.zeros((len(s),len(genes)));contact=np.zeros_like(M,dtype=bool)
for i,row in s.iterrows():
    for j,g in enumerate(genes):
        z=p[(p.observation==row.observation)&(p.partner_genes==g)]
        if len(z):
            M[i,j]=z.pair_only_buried_pct.max()
            contact[i,j]=(z.atom_pairs_le_4_A>0).any()
expected=pd.read_csv(DATA/"contact_map_pair_burial_values.csv",index_col=0)
assert np.allclose(M,expected.loc[s.observation,genes].values)
plt.rcParams.update({"font.family":"DejaVu Sans","font.size":9,"svg.fonttype":"none"})
fig=plt.figure(figsize=(6.4,7.7),facecolor="white")
fig.text(.02,.977,"Partner-resolved N-terminal engagement",fontsize=12,weight="bold",va="top")
fig.text(.02,.94,"22 chain observations · 3 proteins · 7 structural entries",fontsize=9)
ax=fig.add_axes([.365,.245,.58,.565])
cmap=LinearSegmentedColormap.from_list("pair_burial",["#FFFFFF","#BCE2E7","#20808D","#1B474D"])
ax.imshow(M,cmap=cmap,vmin=0,vmax=100,aspect="auto")
ax.set_xticks(range(len(genes)),genes,rotation=55,ha="left",fontsize=9)
ax.xaxis.tick_top();ax.tick_params(length=0,pad=5)
labels=[f"{r.gene}  {r.pdb}/{r.target_chain}  {r.target_resname.title()}{r.target_authnum}"
        +("*" if r.target_missing_heavy_atoms else "") for r in s.itertuples()]
ax.set_yticks(range(len(s)),labels,fontsize=9)
ax.set_xticks(np.arange(-.5,len(genes),1),minor=True)
ax.set_yticks(np.arange(-.5,len(s),1),minor=True)
ax.grid(which="minor",color="#D4D1CA",lw=.4);ax.tick_params(which="minor",length=0)
for i in range(len(s)):
    for j in range(len(genes)):
        c="white" if M[i,j]>55 else "#28251D"
        if M[i,j]>=1:ax.text(j,i,f"{M[i,j]:.0f}",ha="center",va="center",fontsize=9,color=c)
        if contact[i,j]:ax.plot(j+.32,i-.30,"o",ms=2,color=c)
for y in [7.5,12.5]:ax.axhline(y,color="#28251D",lw=1)
for spine in ax.spines.values():spine.set_visible(False)
notes=[
    "Values: pair-only burial of the target residue (%) with one partner.",
    "Dots: heavy-atom proximity ≤4 Å. Not a validated bond or function.",
    "Pair-only burial is nonadditive; it is not a fraction of total burial.",
    "Blank: <1% burial or no eligible partner; not proof of no interaction.",
    "* Missing OG in four Ser2 models; CG/SD/CE in 8TM4/G Met1.",
    "Met1 and Ser2 are distinct positions; chemistry is not sample-validated."
]
for k,note in enumerate(notes):fig.text(.02,.193-k*.030,note,fontsize=9)
for ext in ["png","svg"]:fig.savefig(OUT/f"Partner_Resolved_Contact_Map_Print.{ext}",dpi=400)
plt.close(fig)
```
