## Supplementary_Data_7 for "Human protein complex interfaces reveal candidate assembly roles for initiator methionine excision": plot_partner_resolved.py

```
from pathlib import Path
import numpy as np,pandas as pd
import matplotlib
matplotlib.use("Agg")
import matplotlib.pyplot as plt
from matplotlib.colors import LinearSegmentedColormap

ROOT=Path(__file__).resolve().parent
OUT=ROOT/"deliverables/paper1_partner_resolved"
s=pd.read_csv(OUT/"observation_partner_summary.csv").fillna("")
p=pd.read_csv(OUT/"all_partner_measurements.csv").fillna("")
s=s.sort_values(["gene","pdb","target_chain"]).reset_index(drop=True)
genes=["PSMA1","PSMA2","PSMA3","PSMA4","PSMA5","PSMA6","PSMA7","PSMG1","POMP"]
M=np.zeros((len(s),len(genes)));touch=np.zeros_like(M,dtype=bool)
for i,row in s.iterrows():
    for j,g in enumerate(genes):
        z=p[(p.observation==row.observation)&(p.partner_genes==g)]
        if len(z):
            M[i,j]=z.pair_only_buried_pct.max()
            touch[i,j]=(z.atom_pairs_le_4_A>0).any()
plt.rcParams.update({"font.family":"DejaVu Sans","svg.fonttype":"none","font.size":12})
fig=plt.figure(figsize=(11,12.4),facecolor="white")
fig.text(.035,.974,"Partner-resolved N-terminal engagement",fontsize=20,fontweight="bold",va="top")
fig.text(.035,.937,"Different structural contexts reveal different contact partners.",fontsize=14)
fig.text(.035,.912,"Selected audit subset: 22 chain observations • 3 proteins • 7 entries",fontsize=12)
ax=fig.add_axes([.38,.205,.57,.635])
cmap=LinearSegmentedColormap.from_list("occlusion",["#FFFFFF","#BCE2E7","#20808D","#1B474D"])
ax.imshow(M,cmap=cmap,vmin=0,vmax=100,aspect="auto")
ax.set_xticks(range(len(genes)),genes,rotation=45,ha="left",fontsize=11.5)
ax.xaxis.tick_top();ax.tick_params(length=0,pad=9)
labels=[]
for _,r in s.iterrows():
    labels.append(f"{r.gene}   {r.pdb}/{r.target_chain}   {r.target_resname.title()}{r.target_authnum}"+("*" if r.target_missing_heavy_atoms else ""))
ax.set_yticks(range(len(s)),labels,fontsize=11.5)
for label in ax.get_yticklabels():label.set_horizontalalignment("right")
ax.set_xticks(np.arange(-.5,len(genes),1),minor=True)
ax.set_yticks(np.arange(-.5,len(s),1),minor=True)
ax.grid(which="minor",color="#D4D1CA",lw=.45)
ax.tick_params(which="minor",length=0)
for i in range(len(s)):
    for j in range(len(genes)):
        if M[i,j]>=1:
            ax.text(j,i,f"{M[i,j]:.0f}",ha="center",va="center",fontsize=12,
                    color="white" if M[i,j]>55 else "#28251D")
        if touch[i,j]:
            ax.plot(j+.32,i-.29,"o",ms=3.3,color="white" if M[i,j]>55 else "#28251D")
for i in [7.5,12.5]:ax.axhline(i,color="#28251D",lw=1.5)
for spine in ax.spines.values():spine.set_visible(False)
fig.text(.66,.157,"Pair-only burial of the target residue (%)",ha="center",fontsize=13)
fig.text(.035,.128,"Numbers: ≥1% burial with that partner chain alone. ●: a heavy-atom contact ≤4 Å.",fontsize=12)
fig.text(.035,.103,"Pair-only burial values are not additive and are not percentages of total burial.",fontsize=12)
fig.text(.035,.078,"Blank cells: <1% pair-only burial or no eligible partner chain. They do not imply no interaction.",fontsize=11.5)
fig.text(.035,.053,"* Incomplete target side chain: missing OG in four Ser2 models; CG/SD/CE in 8TM4/G Met1.",fontsize=11.5)
fig.text(.035,.028,"Met1 and Ser2 are different positions. Contacts are geometric, not demonstrated bonds or functions.",fontsize=11.5)
for ext in ["png","svg"]:fig.savefig(OUT/f"Partner_Resolved_Contact_Map.{ext}",dpi=300)
plt.close(fig)
matrix=pd.DataFrame(M,index=s.observation,columns=genes)
matrix.to_csv(OUT/"contact_map_pair_burial_values.csv")
```
