## Supplementary_Data_7 for "Human protein complex interfaces reveal candidate assembly roles for initiator methionine excision": README.md

# Supplementary Data 7: partner-resolved proteasome geometry

This supplement supports main-text Figure 3d and Supplementary Figure 3. Those panels show the same map, not independent analyses. The fixed selected subset contains 22 PSMA4, PSMA6 and PSMA7 chain observations from seven structural entries; no inferential statistics or exhaustive proteasome survey are implied.

## Contents and units

The measurement tables are in `deliverables/paper1_partner_resolved/`. Distances are in Å, surface areas in Å², and burial measurements are percentages rather than proportions.

- **observation_partner_summary.csv:** One row per focal residue, including total assembly burial, nearest partner, partners within 4 and 5 Å, the largest pair-only occluder, missing target atoms and common-backbone summaries.
- **all_partner_measurements.csv:** All 540 target–other-protein-chain pairs. Includes chain identity, minimum residue/backbone/N distances, atom-pair counts, pair-only burial, backbone-only sensitivity and completeness flags.
- **atom_contacts_within_5A.csv:** All 759 atom-pair approaches within 5 Å, with target and partner atom identities and author residue numbers.
- **chain_identity_audit.csv:** Frozen accession/gene mappings and structural entity descriptions for chain-identity review.
- **contact_map_pair_burial_values.csv:** Display matrix for the contact map. Dots must be recovered from the ≤4 Å contact columns in the partner table, not from a burial threshold.
- **validation_checks.csv:** Per-observation agreement with the preceding full-assembly geometry analysis.
- **input_hashes.csv:** Frozen assembly hashes and manifest-match flags.
- **analysis_summary.json:** Analysis counts, software environment and validation summary.
- **frozen_sifts_segments_used.csv:** The exact frozen SIFTS rows used for the seven entries.

The original large-format contact map and editable SVG are retained with these tables. The manuscript-sized PNG and editable SVG are in `deliverables/paper1_contactmap_integrated/`; their values are checked against the same display matrix.

## Interpretation

Pair-only burial is 100 × (A₀ − Aₚ)/A₀, where A₀ is the resolved target-residue SASA in its isolated chain and Aₚ is the same target SASA with one partner chain present. It is nonadditive across partners, and it is not a fraction of total assembly burial.

Pair SASA was calculated for 78 pairs whose minimum heavy-atom distance was ≤8 Å. More distant pairs have `pair_sasa_calculated=False`, missing target-plus-partner SASA and zero pair-only occlusion under the geometric exclusion. Zero burial does not establish absence of biological interaction. Contacts at 3.5, 4 and 5 Å describe geometry, not validated bonds.

Five focal residues are incomplete. OG is absent from PSMA6 Ser2 in 4R3O/A and 4R3O/O, PSMA4 Ser2 in 8TM4/B and PSMA7 Ser2 in 8TM4/C; CG, SD and CE are absent from PSMA6 Met1 in 8TM4/G. These observations carry asterisks in the map and explicit table flags. Backbone-only values summarize unchanged calculations over N, CA, C and O; no missing side chains were rebuilt.

Met1 and Ser2 are distinct modeled positions, not sample-validated processing states. Independent chemical-state evidence does not establish the proteoform of each structural preparation. Repeated chain copies are not independent biological replicates, and different structural entries are not a demonstrated maturation trajectory.

## Reproduction

Install the dependencies in `requirements.txt` from the archive root, then run:

```sh
python partner_resolved_analysis.py
python plot_partner_resolved.py
python plot_partner_contactmap_print.py
```

The recorded analysis used Python 3.14.3. FreeSASA may require a compiler on platforms without an available binary package.

The seven already-expanded coordinate files are included under `paper1_audit/frozen/nterm-census-1.3.0/inputs/figure5_assemblies/`. Their historical directory name refers to the original frozen analysis, not current Figure 3d numbering. Coordinates are unmodified and verified against the original manifest; do not apply another assembly expansion.

The SIFTS and human-protein annotation inputs are mechanically reduced subsets of the frozen source files, with retained annotation columns preserved. The original geometry library and fixed 22-observation input table are included. No network retrieval is required for these scripts.

The portable bundle was rerun using these reduced inputs, reproducing both the observation summary and all-partner measurement tables byte-for-byte. The print rendering was checked against the original contact-map value matrix. These checks establish numerical reproducibility, not model-density quality or physiological chemistry.
