## Supplementary figures and images for "Human protein complex interfaces reveal candidate assembly roles for initiator methionine excision"

### fig1_census.png

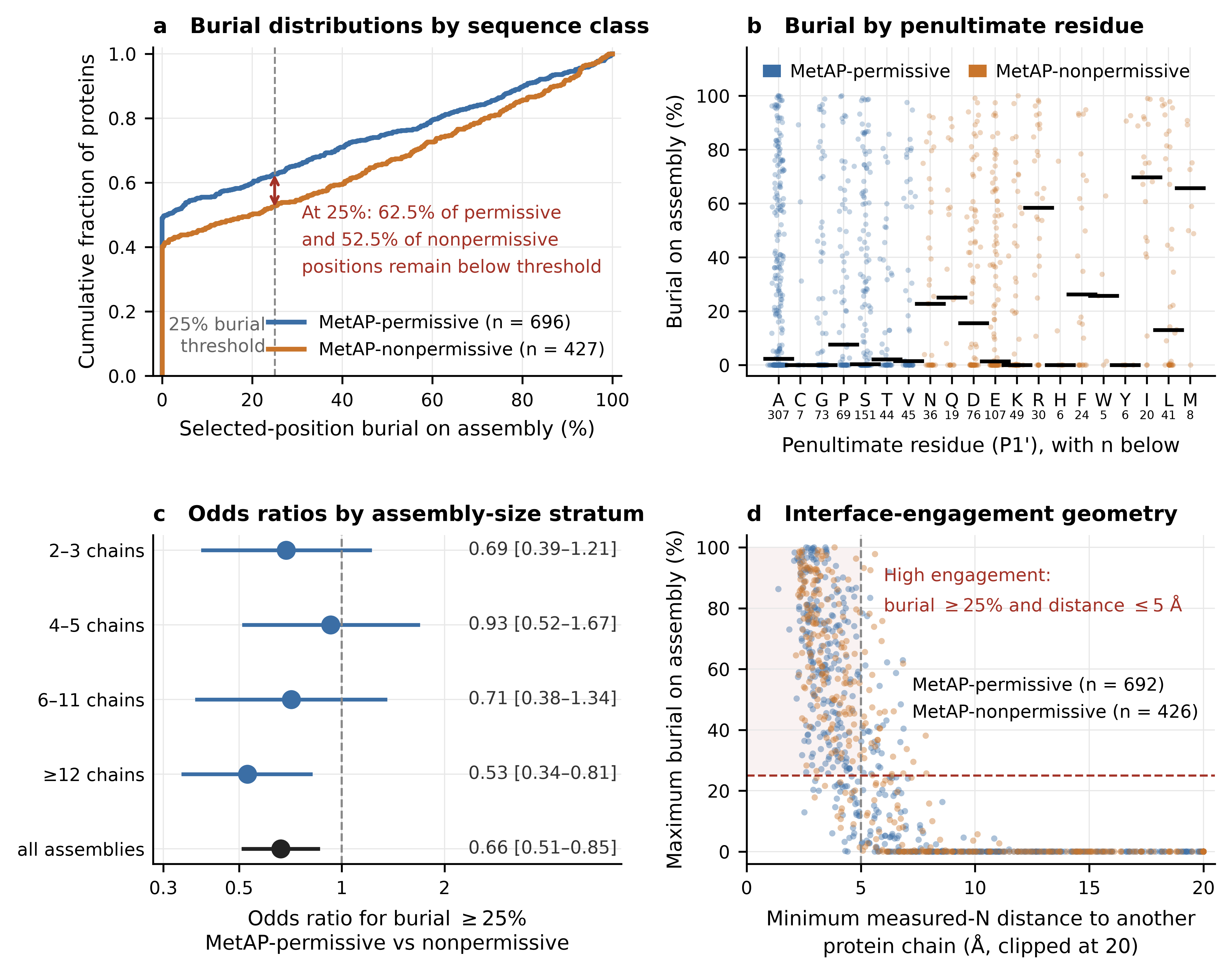

### fig2_machines.png

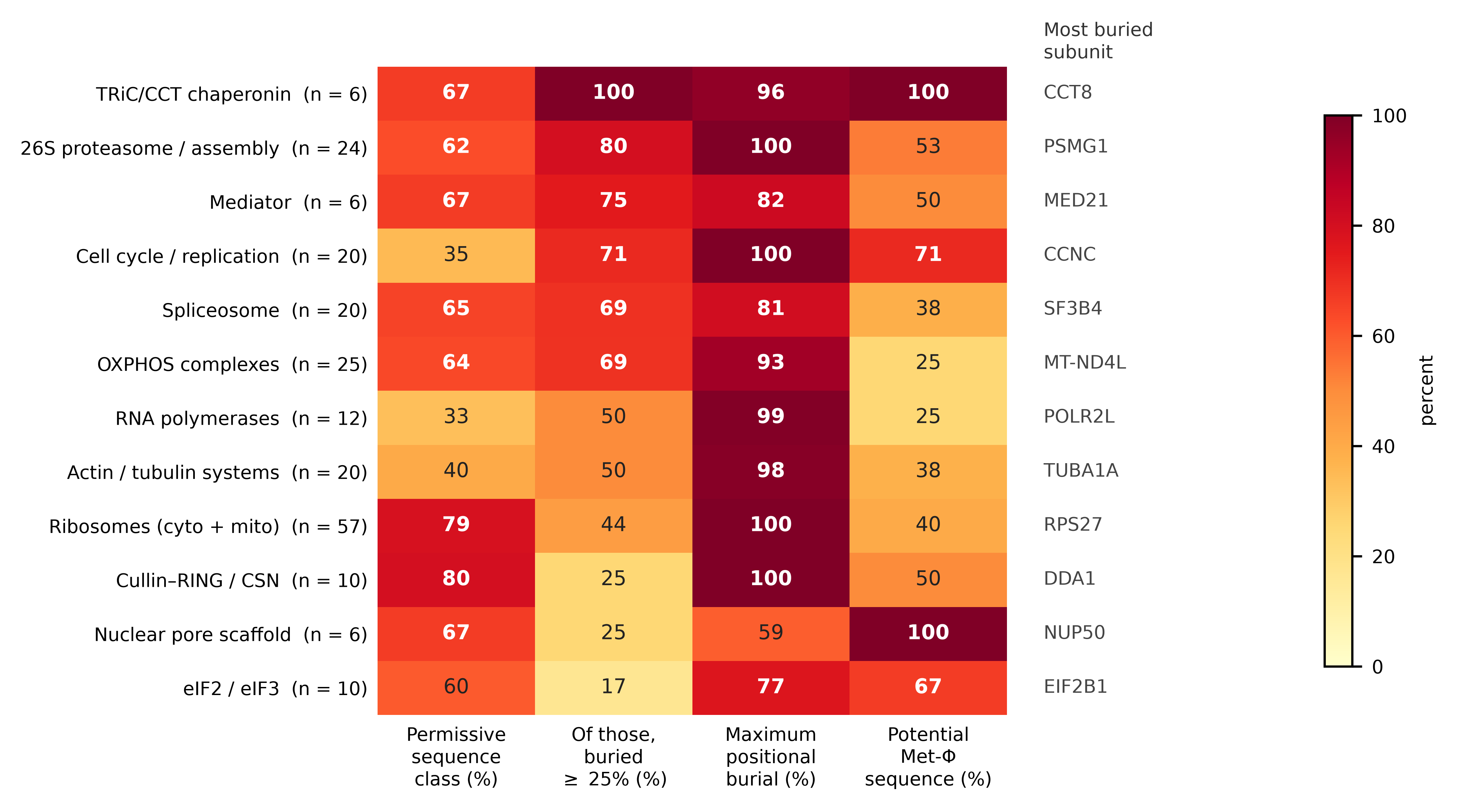

### fig4_transport.png

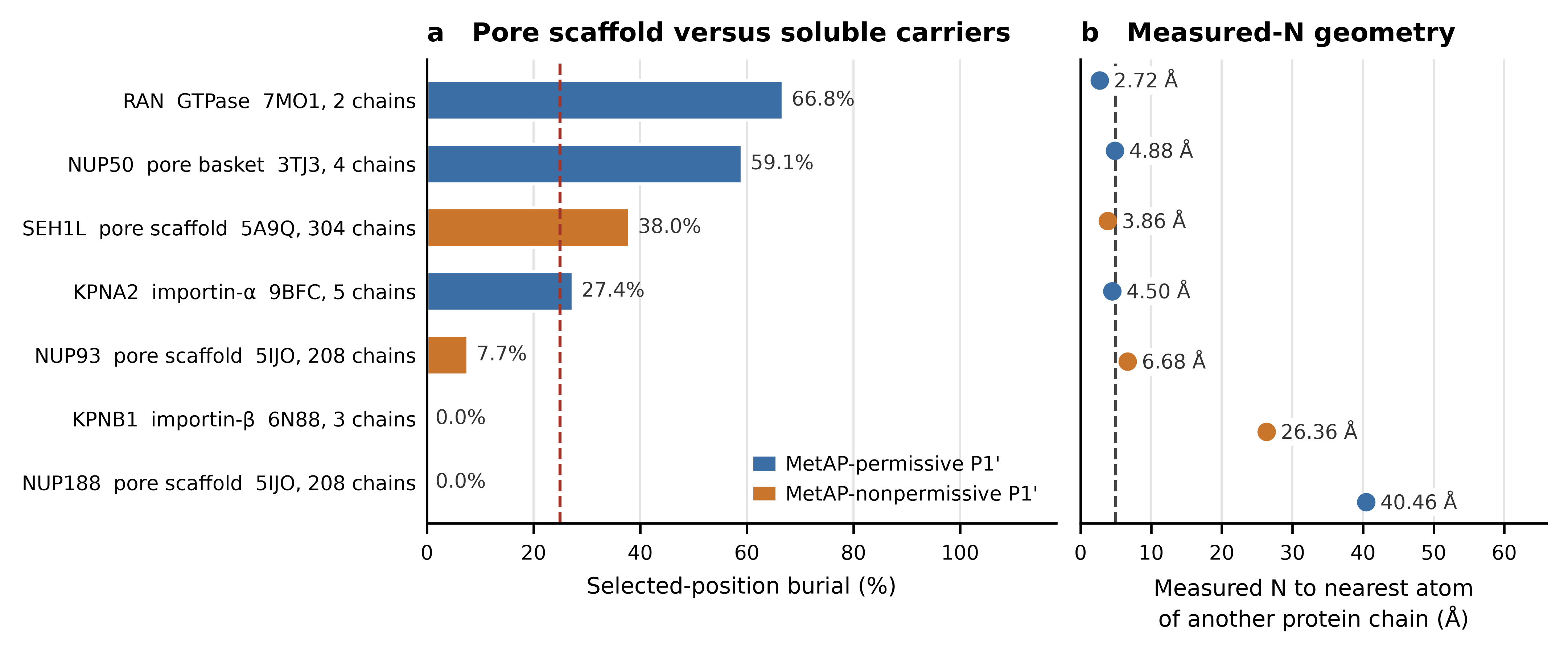

### Partner_Resolved_Contact_Map.png

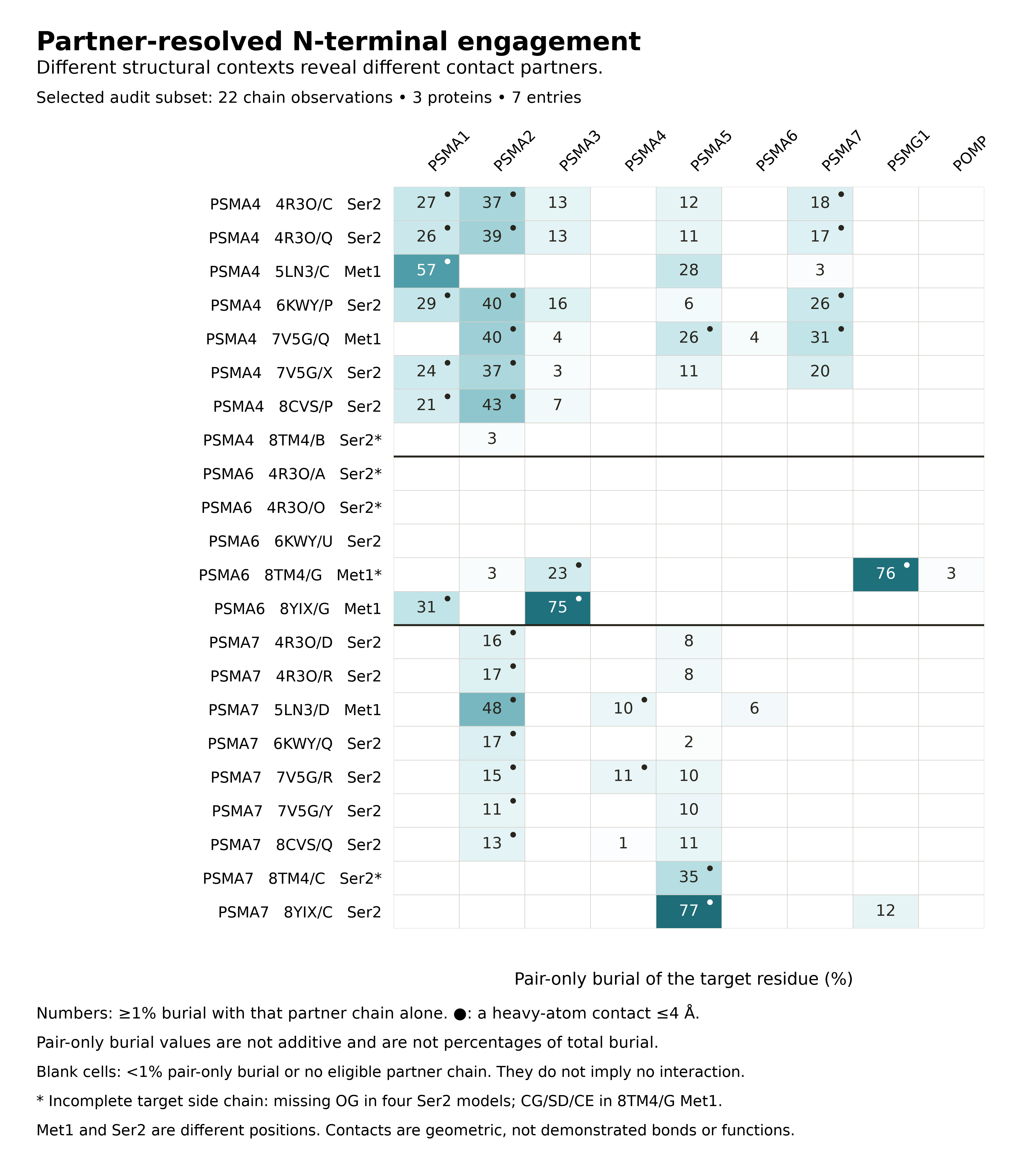

### Partner_Resolved_Contact_Map_Print.png

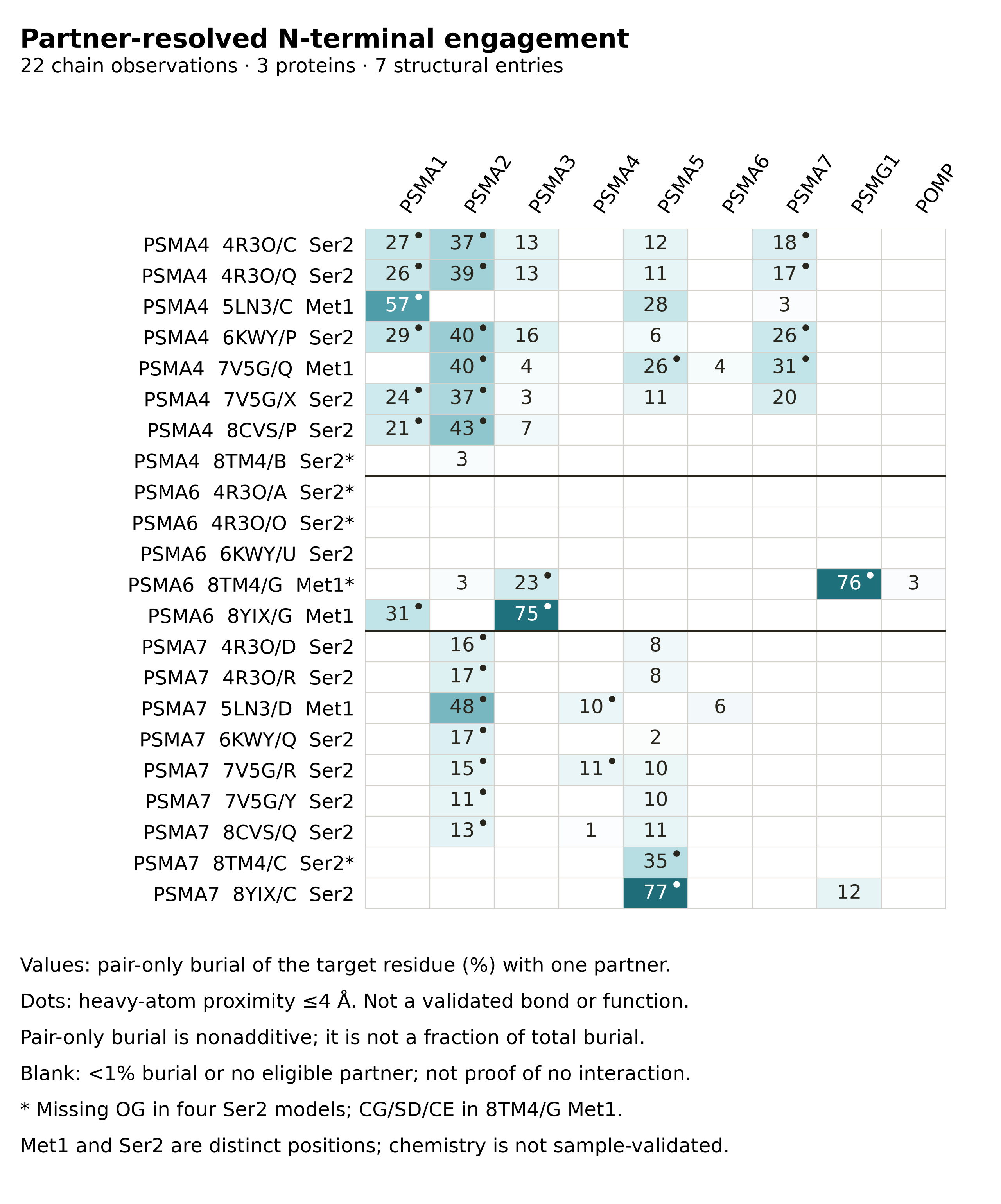

### Supplementary_Figure_1.pdf

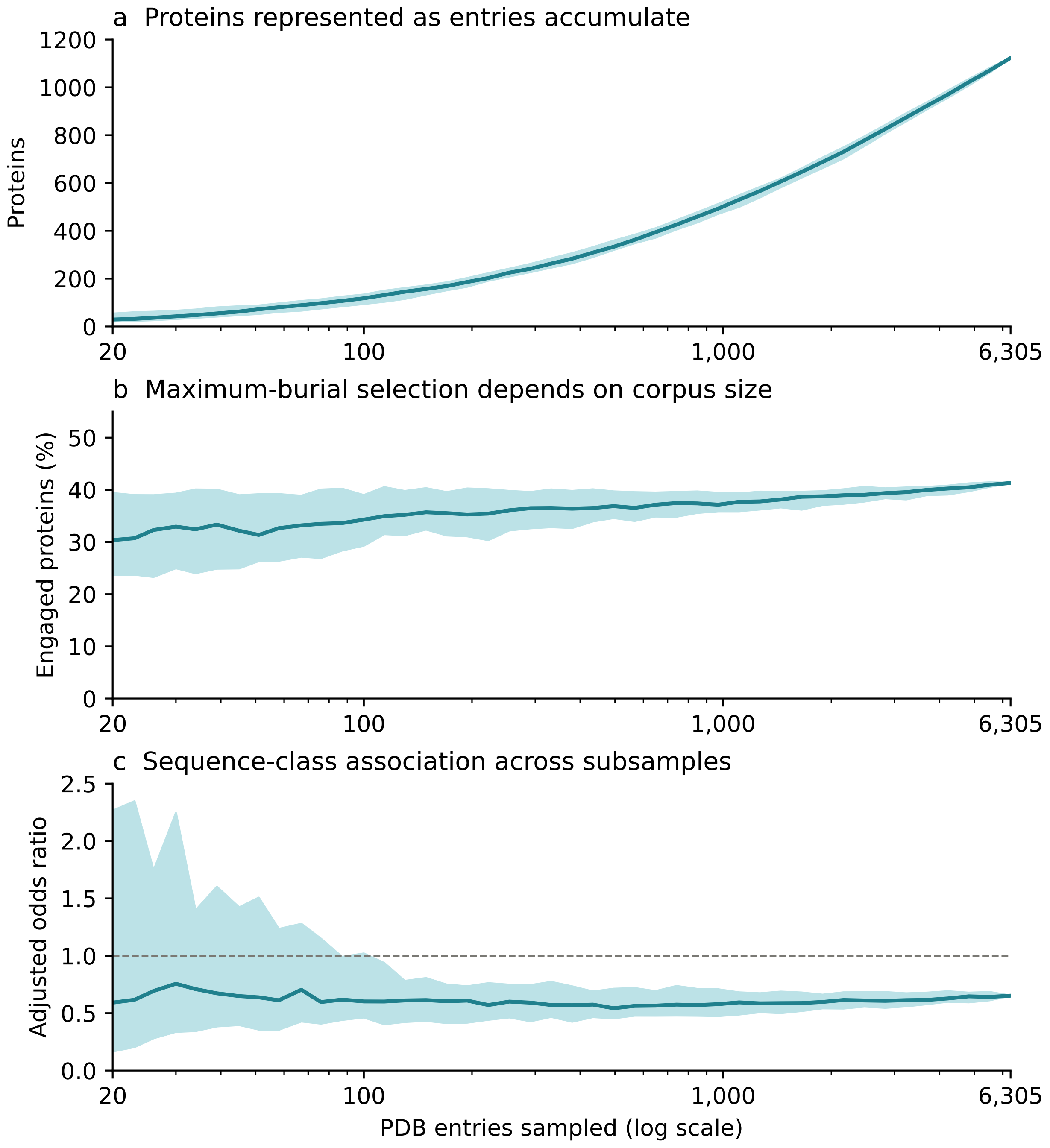

### Supplementary_Figure_1.png

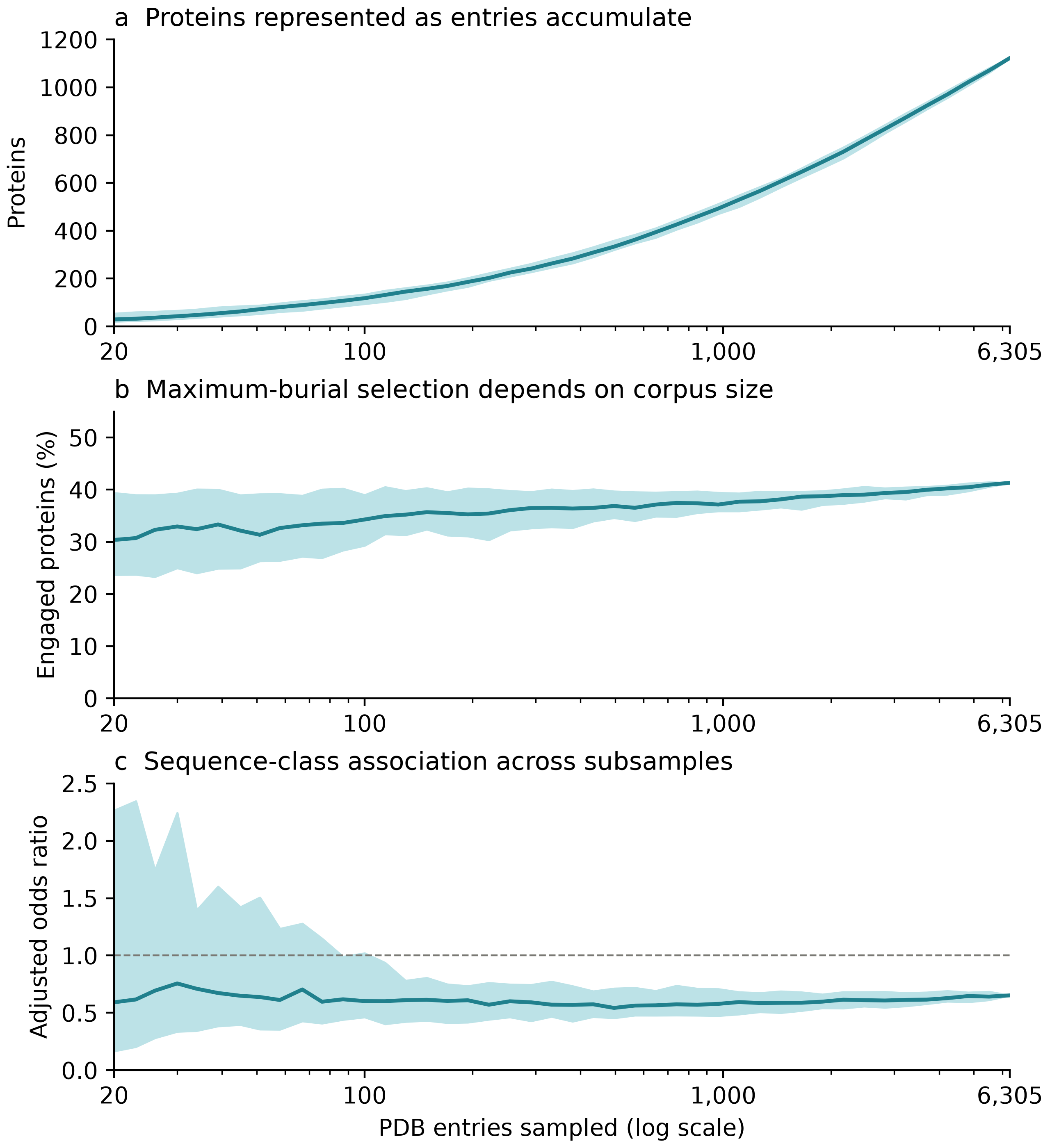

### Supplementary_Figure_2.pdf

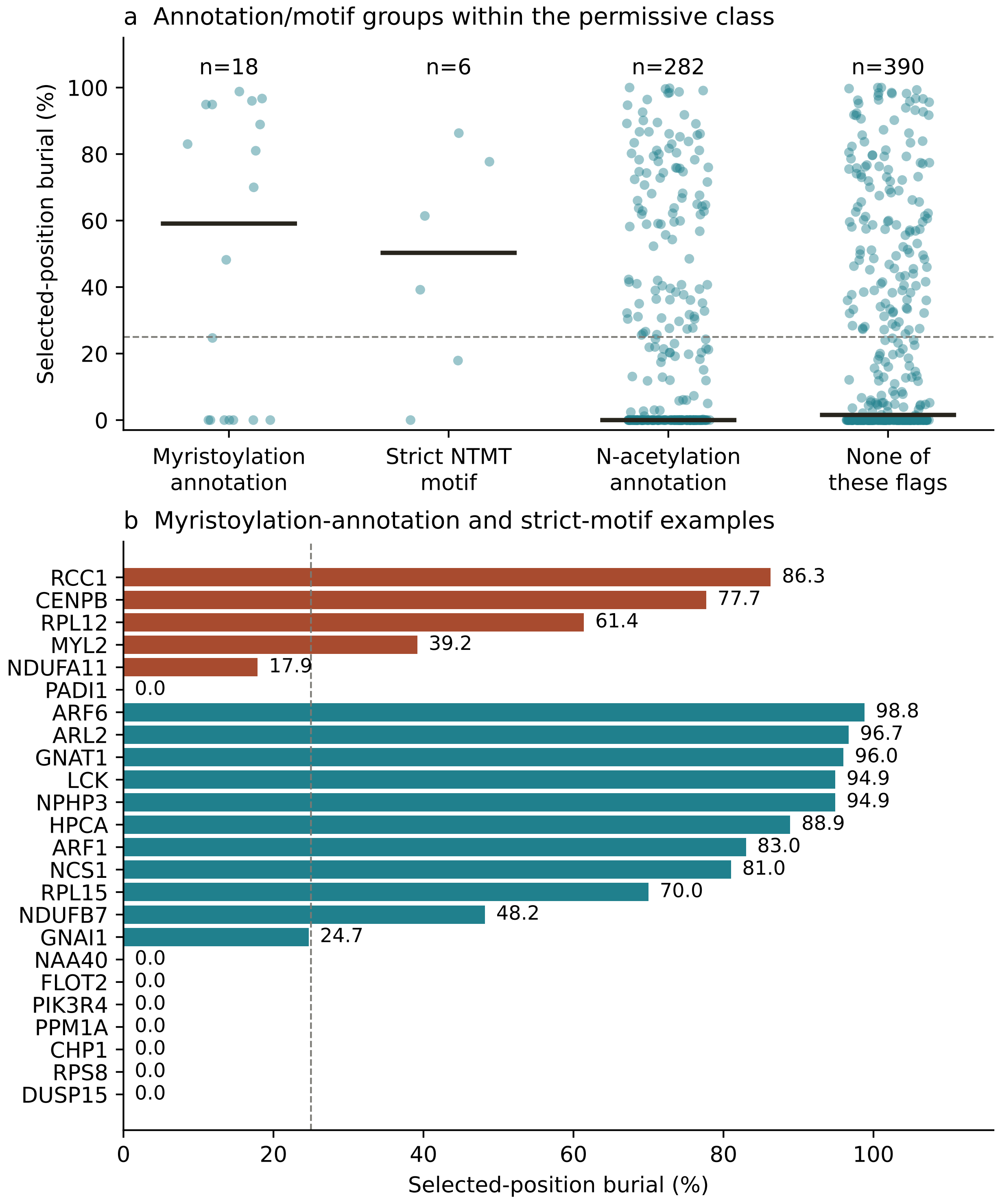

### Supplementary_Figure_2.png

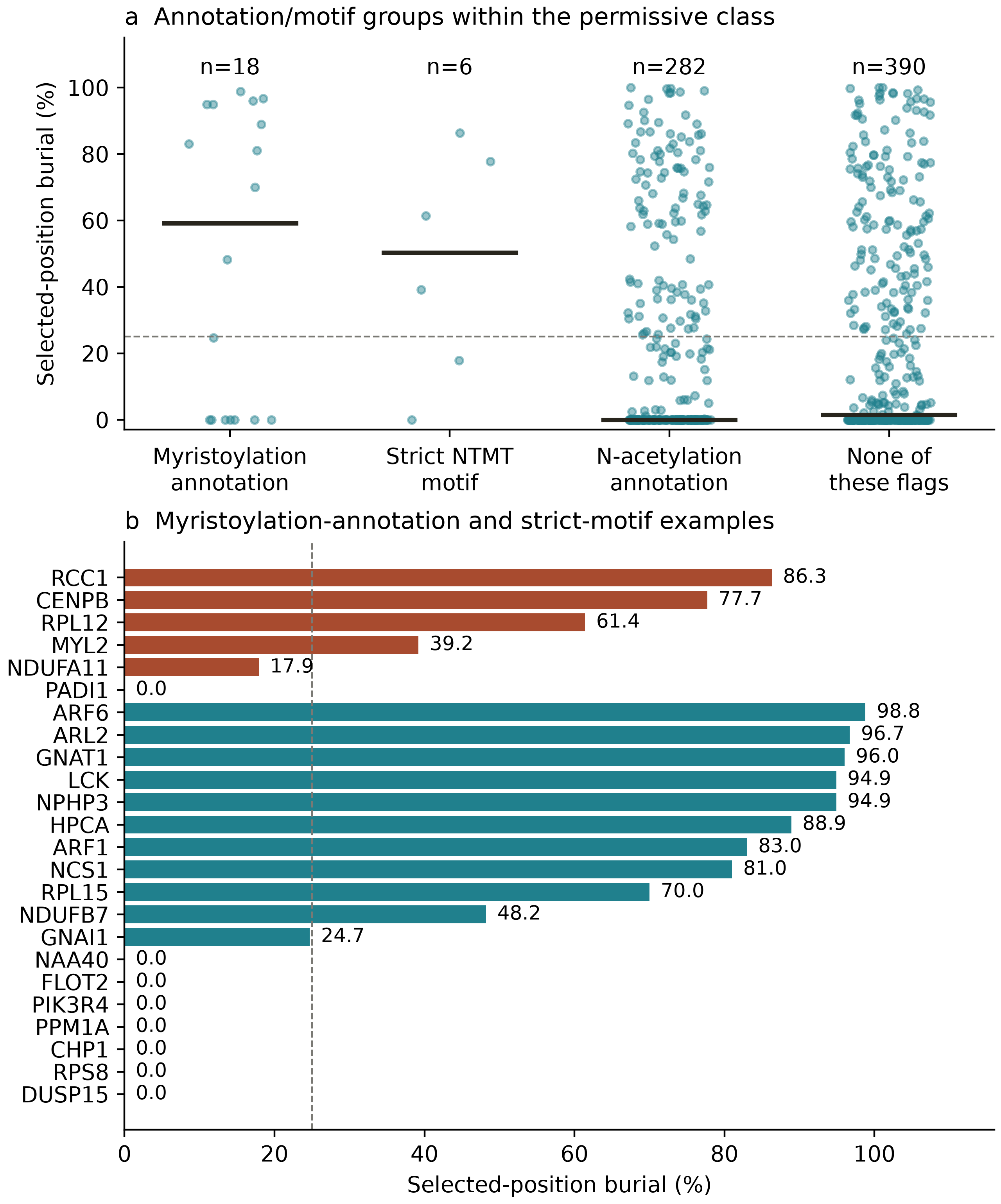
