## Supplementary_Data_8a for "Human protein complex interfaces reveal candidate assembly roles for initiator methionine excision": build_supplement_analysis.py

```
"""Rebuild current-cohort supplement tables and figure source data.

Run from this directory: python build_supplement_analysis.py
Frozen tables are inputs, not newly inferred proteoform assignments.
"""
from pathlib import Path
import json, shutil, os, runpy
import numpy as np
import pandas as pd
from scipy.stats import fisher_exact, mannwhitneyu
import statsmodels.api as sm
import matplotlib
matplotlib.use("Agg")
import matplotlib.pyplot as plt

ROOT=Path(__file__).resolve().parent
LOCAL=ROOT/"inputs"
FROZEN=LOCAL if LOCAL.exists() else ROOT/"paper1_audit/frozen/nterm-census-1.3.0/data"
AUDIT=ROOT/"audit_results" if (ROOT/"audit_results").exists() else ROOT/"paper1_audit/results"
OUT=ROOT/"outputs" if LOCAL.exists() else ROOT/"supplement_build/outputs"
OUT.mkdir(parents=True,exist_ok=True)
p=pd.read_csv(FROZEN/"asm_per_protein_master.csv")
o=pd.read_csv(AUDIT/"observation_processing_mapping_audit.csv")
flags=pd.read_csv(AUDIT/"protein_processing_evidence_flags.csv")
keep=p[~p.cls.str.startswith("excluded")].copy()
ao=o[o.analyzable].copy()
keys=set(zip(p.acc,p.pdb,p.asm_chain))
def reselect(df,largest=False):
    q=df.copy()
    q["_original_selected"]=[(r.acc,r.pdb,r.asm_chain) in keys for r in q.itertuples()]
    q=q.sort_values("_original_selected",ascending=False,kind="stable")
    if largest:
        q=q[q.n_chains==q.groupby("acc").n_chains.transform("max")]
    return q.loc[q.groupby("acc",sort=False).buried_pct.idxmax()].drop(columns="_original_selected")
b=reselect(ao[ao.is_first_observed])
assert len(b)==845
assert (b.res2.isin(list("AGSTCPV"))&(b.buried_pct>=25)).sum()==175

def safe_fields(df):
    q=df.copy()
    if "cls" in q:
        q["cls"]=q.cls.replace({"Met-retained (annotated/observed)":"Modeled MET/MSE; removal not annotated",
            "NME-generated (annotated)":"Initiator removal annotated"})
    if "pred" in q:
        q["pred"]=q.pred.replace({"predicted NME":"MetAP-permissive","predicted Met-retained":"MetAP-nonpermissive"})
    return q.rename(columns={"cls":"annotation_model_category","pred":"sequence_class",
        "mature_start":"annotation_selected_position",
        "min_dist_alphaamino_N":"min_measured_residue_N_distance_A"})
selected=pd.read_csv(AUDIT/"protein_selected_record_audit.csv")
selected=selected.merge(flags[["acc","alternate_removal_annotation","multiple_chain_products_flag","annotated_chain_starts"]],
                        on="acc",how="left",validate="one_to_one")
assert len(selected)==1191
selected["excluded_from_analysis"]=selected.cls.str.startswith("excluded")
selected["boundary_sensitivity_eligible"]=selected.acc.isin(b.acc)
selected["boundary_sensitivity_engaged"]=selected.acc.isin(b.loc[b.buried_pct>=25,"acc"])
selected["boundary_sensitivity_max_burial_pct"]=selected.acc.map(b.set_index("acc").buried_pct)
for name,frame in [("protein_census",selected),("observations",o),("boundary_selected_845",b),
    ("candidates_original_261",selected[(~selected.excluded_from_analysis)&selected.res2.isin(list("AGSTCPV"))&(selected.buried_pct>=25)]),
    ("candidates_boundary_175",b[b.res2.isin(list("AGSTCPV"))&(b.buried_pct>=25)])]:
    safe_fields(frame).to_csv(OUT/(name+".csv"),index=False)

def compare(df,label,threshold=25,small="AGSTCPV"):
    x=df.res2.isin(list(small));y=df.buried_pct>=threshold
    a,b_,c,d=[int(v) for v in [(x&y).sum(),(x&~y).sum(),(~x&y).sum(),(~x&~y).sum()]]
    odds,pv=fisher_exact([[a,b_],[c,d]],alternative="two-sided")
    return dict(analysis=label,threshold_percent=threshold,n=len(df),permissive_n=a+b_,
        permissive_engaged=a,nonpermissive_n=c+d,nonpermissive_engaged=c,odds_ratio=odds,
        fisher_two_sided_p=pv)
rows=[compare(keep,f"Original maximum; threshold {t}%",t) for t in [20,25,30,50]]
summ=[]
for cl,df in keep.groupby(keep.res2.isin(list("AGSTCPV"))):
    dn=df.dN_min.replace([np.inf,-np.inf],np.nan).dropna()
    summ.append(dict(sequence_class="MetAP-permissive" if cl else "MetAP-nonpermissive",
        n=len(df),median_burial_pct=df.buried_pct.median(),mean_burial_pct=df.buried_pct.mean(),
        fraction_engaged=(df.buried_pct>=25).mean(),n_with_finite_N_distance=len(dn),
        median_min_measured_N_distance_A=dn.median()))
pd.DataFrame(summ).to_csv(OUT/"class_summaries.csv",index=False)
tests=[]
for field in ["buried_pct","dN_min"]:
    a=keep.loc[keep.res2.isin(list("AGSTCPV")),field].replace([np.inf,-np.inf],np.nan).dropna()
    z=keep.loc[~keep.res2.isin(list("AGSTCPV")),field].replace([np.inf,-np.inf],np.nan).dropna()
    r=mannwhitneyu(a,z,alternative="two-sided")
    tests.append(dict(test="Mann-Whitney U",field=field,n_permissive=len(a),n_nonpermissive=len(z),
        statistic=r.statistic,two_sided_p=r.pvalue))
pd.DataFrame(tests).to_csv(OUT/"continuous_tests.csv",index=False)
strata=[]
for lo,hi,label in [(1,3,"1-3 chains"),(4,7,"4-7 chains"),(8,11,"8-11 chains"),(12,100000,">=12 chains")]:
    strata.append(compare(keep[keep.n_chains.between(lo,hi)],label))
pd.DataFrame(strata).to_csv(OUT/"assembly_strata.csv",index=False)
alt=[]
for small,label in [("AGSTPV","C reassigned"),("AGSTCPVNQ","N/Q added"),("AGSTCPVDN","D/N added"),("AGSTCPVDNQ","D/N/Q added")]:
    alt.append(compare(keep,label,small=small))
m=keep.copy();m["buried_pct"]=m.buried_median
alt.extend([compare(m,"Median across extracted observations"),compare(reselect(ao,True),"Largest assembly then maximum burial"),
            compare(keep[keep.n_entities>=2],"Selected maximum in heteromeric assembly")])
pd.DataFrame(alt).to_csv(OUT/"alternative_definitions.csv",index=False)
pd.DataFrame(rows).to_csv(OUT/"threshold_tests.csv",index=False)
byres=[]
for aa,g in keep.groupby("res2"):
    byres.append(dict(residue_2=aa,n=len(g),median_burial_pct=g.buried_pct.median(),
        mean_burial_pct=g.buried_pct.mean(),engaged_count=int((g.buried_pct>=25).sum()),
        fraction_engaged=(g.buried_pct>=25).mean()))
pd.DataFrame(byres).to_csv(OUT/"residue2_summary.csv",index=False)
modelrows=[]
for terms in [["permissive","log_chains"],["permissive","log_chains","log_entries"]]:
    X=pd.DataFrame({"permissive":keep.res2.isin(list("AGSTCPV")).astype(float),
                    "log_chains":np.log(keep.n_chains),"log_entries":np.log(keep.n_entries)})
    model=sm.Logit((keep.buried_pct>=25).astype(float),sm.add_constant(X[terms])).fit(disp=False)
    for term in terms:
        ci=model.conf_int().loc[term]
        modelrows.append(dict(model=" + ".join(terms),term=term,n=len(keep),
            odds_ratio=np.exp(model.params[term]),ci_low=np.exp(ci.iloc[0]),ci_high=np.exp(ci.iloc[1]),p=model.pvalues[term]))
pd.DataFrame(modelrows).to_csv(OUT/"current_models.csv",index=False)
loo=[dict(excluded_residue=aa,**compare(keep[keep.res2!=aa],"Leave one residue-2 identity out")) for aa in sorted(keep.res2.unique())]
pd.DataFrame(loo).to_csv(OUT/"leave_one_residue_out.csv",index=False)
extra=[compare(keep[keep.n_entries>=n],f"At least {n} PDB entries") for n in [2,5]]
extra.append(compare(keep[~(keep.res2.isin(list("AGSTCPV"))&keep.resname.isin(["MET","MSE"]))],"Exclude permissive modeled MET/MSE"))
pd.DataFrame(extra).to_csv(OUT/"sampling_sensitivities.csv",index=False)

# Figure S2 retains the existing operational group assignment, not inferred chemistry.
r=pd.DataFrame(json.load(open(FROZEN/"asm_cascade_records.json")))
r=r[r.pred_nme].copy()
r["display_group"]=np.select([r.myr,r.ntmt_motif,r.acetyl],["Myristoylation annotation","Strict NTMT motif","N-terminal acetylation annotation"],default="None of these flags")
assert len(r)==696
r=r.merge(keep[["acc","is_first_observed","resname","asm_chain"]],on="acc",validate="one_to_one")
r=safe_fields(r)
r.to_csv(OUT/"figure_S2_source.csv",index=False)
counts=r.groupby("display_group").agg(n=("acc","size"),median_burial_pct=("buried","median"))
counts.to_csv(OUT/"figure_S2_group_summary.csv")
plt.rcParams.update({"font.family":"DejaVu Sans","font.size":10,"axes.titlesize":11,"svg.fonttype":"none"})
fig,axes=plt.subplots(2,1,figsize=(6.4,7.7),layout="constrained",gridspec_kw={"height_ratios":[1,1.5]})
groups=["Myristoylation annotation","Strict NTMT motif","N-terminal acetylation annotation","None of these flags"]
labels=["Myristoylation\nannotation","Strict NTMT\nmotif","N-acetylation\nannotation","None of\nthese flags"]
rng=np.random.default_rng(7)
for i,g in enumerate(groups):
    v=r.loc[r.display_group==g,"buried"]
    axes[0].scatter(i+rng.uniform(-.19,.19,len(v)),v,s=12,alpha=.45,color="#20808D")
    axes[0].plot([i-.3,i+.3],[v.median()]*2,color="#28251D",lw=2)
    axes[0].text(i,104,f"n={len(v)}",ha="center",fontsize=10)
axes[0].set(xticks=range(4),xticklabels=labels,ylim=(-3,115),ylabel="Selected-position burial (%)")
axes[0].set_title("a  Annotation/motif groups within the permissive class",loc="left")
axes[0].axhline(25,color="#7A7974",ls="--",lw=.8)
cases=pd.concat([r[r.display_group==g].sort_values("buried") for g in groups[:2]])
axes[1].barh(range(len(cases)),cases.buried,color=["#20808D" if g==groups[0] else "#A84B2F" for g in cases.display_group])
axes[1].set(yticks=range(len(cases)),yticklabels=cases.gene,xlim=(0,116),xlabel="Selected-position burial (%)")
for i,v in enumerate(cases.buried):axes[1].text(v+1.5,i,f"{v:.1f}",va="center",fontsize=9)
axes[1].axvline(25,color="#7A7974",ls="--",lw=.8)
axes[1].set_title("b  Myristoylation-annotation and strict-motif examples",loc="left")
for ax in axes:
    for side in ["top","right"]:ax.spines[side].set_visible(False)
for ext in ["png","svg","pdf"]:fig.savefig(OUT/f"Supplementary_Figure_2.{ext}",dpi=400)
plt.close(fig)

# Use the frozen rarefaction implementation, with a new readable rendering.
rare=ROOT/"code/rebuild_supplementary_figure1.py" if LOCAL.exists() else ROOT/"paper1_audit/frozen/nterm-census-1.3.0/code/rebuild_supplementary_figure1.py"
os.environ["NME_WORK"]=str(FROZEN);os.environ["NME_FIGOUT"]=str(OUT)
z=runpy.run_path(str(rare));plt.close("all")
R=z["R"];x=z["checkpoints"]
pd.DataFrame([dict(replicate=i+1,pdb_entries=int(n),protein_count=R[i,j,0],
    percent_engaged=R[i,j,1],adjusted_odds_ratio=R[i,j,2]) for i in range(len(R)) for j,n in enumerate(x)]).to_csv(OUT/"figure_S1_source.csv",index=False)
plt.rcParams.update({"font.size":10,"xtick.labelsize":10,"ytick.labelsize":10,"axes.labelsize":10,"axes.titlesize":11})
fig,axes=plt.subplots(3,1,figsize=(6.4,7.0),layout="constrained")
for k,(title,ylabel) in enumerate([("a  Proteins represented as entries accumulate","Proteins"),("b  Maximum-burial selection depends on corpus size","Engaged proteins (%)"),("c  Sequence-class association across subsamples","Adjusted odds ratio")]):
    axes[k].fill_between(x,z["lo"][:,k],z["hi"][:,k],color="#BCE2E7")
    axes[k].plot(x,z["med"][:,k],color="#20808D",lw=1.7)
    axes[k].set_xscale("log");axes[k].set_ylabel(ylabel);axes[k].set_title(title,loc="left")
    axes[k].set_xlim(20,z["n_pdb"]);axes[k].set_xticks([20,100,1000,6305],["20","100","1,000","6,305"])
    for side in ["top","right"]:axes[k].spines[side].set_visible(False)
axes[0].set_ylim(0,1200);axes[1].set_ylim(0,55);axes[2].set_ylim(0,2.5)
axes[2].axhline(1,color="#7A7974",lw=.8,ls="--")
axes[2].set_xlabel("PDB entries sampled (log scale)")
for ext in ["png","svg","pdf"]:fig.savefig(OUT/f"Supplementary_Figure_1.{ext}",dpi=400)
plt.close(fig)
(OUT/"figS1_rarefaction.png").unlink(missing_ok=True)
summary=dict(proteins=len(p),observations=len(o),analyzable=len(keep),analyzable_observations=len(ao),
    entries=ao.pdb.nunique(),boundary_n=len(b),boundary_candidates=175,original_candidates=261,
    S2=counts.reset_index().to_dict("records"),
    S1_endpoint=[float(v) for v in [z["n_final"],z["pct_final"],z["or_final"]]],
    alternative_definitions=alt,loo_OR_min=min(x["odds_ratio"] for x in loo),
    loo_OR_max=max(x["odds_ratio"] for x in loo),loo_p_max=max(x["fisher_two_sided_p"] for x in loo))
(OUT/"analysis_summary.json").write_text(json.dumps(summary,indent=2))
print(json.dumps(summary,indent=2))
```
