## Supplementary_Data_8a for "Human protein complex interfaces reveal candidate assembly roles for initiator methionine excision": rebuild_supplementary_figure1.py

```
"""Rebuild Supplementary Figure S1 (corpus-size rarefaction) for revision the deposited analysis.

Reads only the deposited derived tables (data/asm_per_protein_master.csv and
data/asm_observations.csv). The v20 script for this figure is not in the deposit, so
the rarefaction is re-derived here from the same tables. On the release-1.3.0 master
table (20 Ig/TCR rows marked excluded) it converges on the three manuscript endpoints exactly
(1123 analyzable proteins over 6305 retained assemblies, 41.3% at or above 25% burial,
size-adjusted odds ratio 0.652); on the 1.2.2 table it gave 1143 / 41.2% / 0.654.

Procedure. Biological assemblies (PDB entries) are shuffled into a random order and
processed one at a time. At log-spaced checkpoints the analysis is rerun on the entries
seen so far: each protein is collapsed to its maximum burial over the entries drawn,
and the three summaries in the panels are recomputed. Because the curve depends on the
draw order, N_REP independent shuffles are run under a fixed seed and the band shows
the 10th-90th percentile across them; the heavy line is the median.

Panel c fits burial >= 25% ~ predicted class + log(number of chains) by iteratively
reweighted least squares and plots exp(coefficient) for the class term, with a Wald
interval. This is the same size-adjusted model reported in the Results.

Authored at 180 mm printed width, so the point sizes below are the point sizes on the
page. Do not widen the figsize without scaling the fonts.

Usage:  NME_WORK=<dir with the derived tables> python rebuild_supplementary_figure1.py
Output: figS1_rarefaction.png
"""
import os as _os
BASE = _os.environ.get("NME_DIR") or _os.path.dirname(_os.path.dirname(_os.path.abspath(__file__)))
WORK_DIR = _os.environ.get("NME_WORK") or _os.path.join(BASE, "work")
OUTDIR = _os.environ.get("NME_FIGOUT") or WORK_DIR
_os.makedirs(OUTDIR, exist_ok=True)

import csv
import numpy as np
import matplotlib
matplotlib.use("Agg")
import matplotlib.pyplot as plt

IN = WORK_DIR + "/"
OUT = OUTDIR + "/"

C_NME = "#3b6ea5"
C_MET = "#c9752b"
GREY = "#555555"
W = 7.09
ANN = 6.6
PANEL = 8.6
AX = 7.4
TICK = 6.6
plt.rcParams["font.size"] = TICK
plt.rcParams["xtick.labelsize"] = TICK
plt.rcParams["ytick.labelsize"] = TICK

N_REP = 40
SEED = 20250823

# ---------------------------------------------------------------- input tables
prot = list(csv.DictReader(open(IN + "asm_per_protein_master.csv")))
keep = {p["acc"]: p for p in prot if not p["cls"].startswith("excluded")}
pred_of = {a: (p["pred"] == "predicted NME") for a, p in keep.items()}

obs = [r for r in csv.DictReader(open(IN + "asm_observations.csv")) if r["acc"] in keep]

accs = sorted(keep)
acc_ix = {a: i for i, a in enumerate(accs)}
pdbs = sorted({r["pdb"] for r in obs})
pdb_ix = {p: i for i, p in enumerate(pdbs)}

o_acc = np.array([acc_ix[r["acc"]] for r in obs], dtype=np.int32)
o_pdb = np.array([pdb_ix[r["pdb"]] for r in obs], dtype=np.int32)
o_bur = np.array([float(r["buried_pct"]) for r in obs])
o_nch = np.array([float(r["n_chains"]) for r in obs])
n_acc, n_pdb = len(accs), len(pdbs)
is_nme = np.array([pred_of[a] for a in accs], dtype=float)

print(f"{n_acc} analyzable proteins, {len(obs)} observations, {n_pdb} biological assemblies")


def adjusted_or(y, x_cls, log_ch):
    """exp(class coefficient) and its Wald interval from a logistic fit."""
    X = np.column_stack([np.ones(len(y)), x_cls, log_ch])
    if len(y) < 12 or len(np.unique(y)) < 2 or len(np.unique(x_cls)) < 2:
        return np.nan, np.nan, np.nan
    b = np.zeros(3)
    for _ in range(60):
        p = 1.0 / (1.0 + np.exp(-X @ b))
        wt = np.clip(p * (1.0 - p), 1e-9, None)
        H = X.T @ (X * wt[:, None]) + 1e-8 * np.eye(3)
        step = np.linalg.solve(H, X.T @ (y - p))
        b = b + np.clip(step, -4, 4)
        if np.max(np.abs(step)) < 1e-9:
            break
    p = 1.0 / (1.0 + np.exp(-X @ b))
    wt = np.clip(p * (1.0 - p), 1e-9, None)
    try:
        se = np.sqrt(np.diag(np.linalg.inv(X.T @ (X * wt[:, None]) + 1e-8 * np.eye(3))))[1]
    except np.linalg.LinAlgError:
        se = np.nan
    if not np.isfinite(b[1]) or abs(b[1]) > 8:
        return np.nan, np.nan, np.nan
    return np.exp(b[1]), np.exp(b[1] - 1.96 * se), np.exp(b[1] + 1.96 * se)


def summarise(mask):
    """Collapse the drawn observations to one row per protein and summarise."""
    bur = np.full(n_acc, -1.0)
    np.maximum.at(bur, o_acc[mask], o_bur[mask])
    seen = bur >= 0.0
    if seen.sum() == 0:
        return 0, np.nan, np.nan
    # chain count of the entry that supplied each protein's maximum burial
    row = np.nonzero(mask)[0]
    a_sel, b_sel, c_sel = o_acc[mask], o_bur[mask], o_nch[mask]
    # ascending burial within protein, ties broken toward the earliest table row, so the
    # final write per protein is the first observation attaining its maximum burial
    order = np.lexsort((-row, b_sel, a_sel))
    nch = np.zeros(n_acc)
    nch[a_sel[order]] = c_sel[order]
    pct = 100.0 * np.mean(bur[seen] >= 25.0)
    orr = adjusted_or((bur[seen] >= 25.0).astype(float), is_nme[seen], np.log(nch[seen]))[0]
    return int(seen.sum()), pct, orr


checkpoints = np.unique(np.round(np.logspace(np.log10(20), np.log10(n_pdb), 44)).astype(int))
rng = np.random.default_rng(SEED)
R = np.full((N_REP, len(checkpoints), 3), np.nan)

for rep in range(N_REP):
    rank = np.empty(n_pdb, dtype=np.int32)
    rank[rng.permutation(n_pdb)] = np.arange(n_pdb)
    o_rank = rank[o_pdb]
    for j, n in enumerate(checkpoints):
        R[rep, j] = summarise(o_rank < n)

med = np.nanmedian(R, axis=0)
lo = np.nanpercentile(R, 10, axis=0)
hi = np.nanpercentile(R, 90, axis=0)

# ---------------------------------------------------------------- final values
n_final, pct_final, or_final = summarise(np.ones(len(obs), dtype=bool))
print(f"endpoints: {n_final} proteins, {pct_final:.2f}% at or above 25% burial, "
      f"size-adjusted OR {or_final:.3f}")

# ---------------------------------------------------------------------- figure
fig, (a0, a1, a2) = plt.subplots(1, 3, figsize=(W, 2.05), layout="constrained")
fig.get_layout_engine().set(wspace=0.07)
x = checkpoints

a0.fill_between(x, lo[:, 0], hi[:, 0], color=C_NME, alpha=0.22, lw=0)
a0.plot(x, med[:, 0], color=C_NME, lw=1.6)
a0.axhline(n_final, color="#777777", lw=0.8, ls=":")
a0.text(22, n_final - 60, f"{n_final} analyzable proteins", fontsize=ANN, color=GREY, va="top")
a0.set_ylabel("distinct proteins recovered", fontsize=AX)
a0.set_title("a   Rarefaction over the corpus", fontsize=PANEL, loc="left")
a0.set_ylim(0, n_final * 1.13)

a1.fill_between(x, lo[:, 1], hi[:, 1], color=C_MET, alpha=0.22, lw=0)
a1.plot(x, med[:, 1], color=C_MET, lw=1.6)
a1.axhline(pct_final, color="#777777", lw=0.8, ls=":")
a1.annotate(f"final value {pct_final:.1f}%", xy=(n_pdb, pct_final), xytext=(150, 47.8),
            fontsize=ANN, color=GREY, ha="left", va="center",
            arrowprops=dict(arrowstyle="-", color=GREY, lw=0.6,
                            shrinkA=0, shrinkB=2, connectionstyle="arc3,rad=0.0"))
a1.text(22, 8.8,
        "rises with corpus size:\nthe maximum collapse is\nupward-biased in the number\nof observations per protein",
        fontsize=ANN, color=C_MET, va="bottom", linespacing=1.5)
a1.set_ylabel("proteins \u2265 25% buried (%)", fontsize=AX)
a1.set_title("b   Burial summary drifts up", fontsize=PANEL, loc="left")
a1.set_ylim(7, 50)

a2.fill_between(x, lo[:, 2], hi[:, 2], color=C_NME, alpha=0.20, lw=0)
a2.plot(x, med[:, 2], color=C_NME, lw=1.6)
a2.axhline(1.0, color="#333333", lw=0.9)
a2.axhline(or_final, color=C_MET, lw=0.9, ls="--")
a2.text(22, 1.03, "no association", fontsize=ANN, color="#333333", va="bottom")
a2.annotate(f"final estimate {or_final:.3f}", xy=(n_pdb, or_final), xytext=(150, 0.36),
            fontsize=ANN, color=C_MET, ha="left", va="center",
            arrowprops=dict(arrowstyle="-", color=C_MET, lw=0.6, shrinkA=0, shrinkB=2))
a2.set_ylabel("size-adjusted odds ratio", fontsize=AX)
a2.set_title("c   Class effect is stable", fontsize=PANEL, loc="left")
a2.set_ylim(0.3, 1.5)

for ax in (a0, a1, a2):
    ax.set_xscale("log")
    ax.set_xlim(20, n_pdb * 1.02)
    ax.set_xlabel("biological assemblies processed", fontsize=AX)
    ax.spines[["top", "right"]].set_visible(False)

fig.savefig(OUT + "figS1_rarefaction.png", dpi=600)
print("wrote " + OUT + "figS1_rarefaction.png")
```
