## Supplementary_Data_8a for "Human protein complex interfaces reveal candidate assembly roles for initiator methionine excision": relabel_main_figures.py

```
"""Rebuild final manuscript Figures 1-4 from the deposited analysis.

Reads only the deposited derived tables (data/asm_per_protein_master.csv and
data/asm_observations.csv). Includes the final all-20-residue panel and the
terminology used by the 25 August 2026 manuscript.

  1. panel letters are lowercase (a, b, c, d) to match the figure legends;
  2. the sub-5 pt explanatory paragraphs baked under Figs 1 and 2 are removed --
     their content lives in the legends, Results and Methods;
  3. every figure is authored at its final printed width (180 mm double column), so
     the point sizes in this file are the point sizes on the page. v20 authored on an
     11.6 in canvas and placed the result at 5.83 in, which halved every label; that
     is why the explanatory paragraphs measured ~4 pt in the accepted PDF.
     The Fig 2 colorbar ticks are also set to the same size as its row labels.

Do not widen the figsize values below without scaling the font sizes to match.

Stratum odds-ratio confidence intervals are exact conditional (Fisher) intervals,
which is what the v20 panel c reports.

Usage:  python rebuild_figures_1_to_4.py  # defaults to the deposited data/ tables
Outputs: fig1_census.png, fig2_machines.png, fig3_alpha_ring.png, fig4_transport.png
"""
import os as _os
BASE = _os.environ.get("NME_DIR") or _os.path.dirname(_os.path.dirname(_os.path.abspath(__file__)))
WORK_DIR = _os.environ.get("NME_WORK") or _os.path.join(BASE, "data")
OUTDIR = _os.environ.get("NME_FIGOUT") or WORK_DIR
_os.makedirs(OUTDIR, exist_ok=True)

import csv, json, collections, re
import numpy as np
import matplotlib
matplotlib.use("Agg")
import matplotlib.pyplot as plt
from matplotlib.patches import Patch
from matplotlib.ticker import NullFormatter, NullLocator
from scipy.stats.contingency import odds_ratio

IN = WORK_DIR + "/"
OUT = OUTDIR + "/"
SMALL = set("AGSTCPV")
PHI = set("LFYWIVA")
C_NME = "#3b6ea5"
C_MET = "#c9752b"
RED = "#a33227"
W = 7.09                       # 180 mm: journal double-column width == placed width
ANN = 7.6                      # enlarged for readability after DOCX placement
PANEL = 9.0                    # compact enough for the two-column panel titles
AX = 8.5                       # axis labels
TICK = 7.6                     # tick labels
plt.rcParams["font.size"] = TICK
plt.rcParams["xtick.labelsize"] = TICK
plt.rcParams["ytick.labelsize"] = TICK

prot = list(csv.DictReader(open(IN + "asm_per_protein_master.csv")))
for p in prot:
    p["b"] = float(p["buried_pct"])
    p["dN"] = float(p["dN_min"]) if p["dN_min"] not in ("", "inf") else np.nan
    p["nch"] = int(p["n_chains"])
# Exclude signal/propeptide/transit rows (48) and immunoglobulin/T-cell receptor
# constant-region entries flagged as unclassified (20); analyses reported in the
# manuscript are computed on the remaining 1,123-protein set.
keep = [p for p in prot if not p["cls"].startswith("excluded") and p["cls"] != "unclassified"]
A = [p for p in keep if p["pred"] == "predicted NME"]
B = [p for p in keep if p["pred"] == "predicted Met-retained"]

# =============================================================== FIGURE 1
fig = plt.figure(figsize=(W, 5.6), layout="constrained")
fig.get_layout_engine().set(hspace=0.10, wspace=0.06)
gs = fig.add_gridspec(2, 2, height_ratios=[1, 1])
ax0, ax1, ax2, ax3 = (fig.add_subplot(gs[0, 0]), fig.add_subplot(gs[0, 1]),
                      fig.add_subplot(gs[1, 0]), fig.add_subplot(gs[1, 1]))

# --- a: cumulative distributions
below = {}
for g, c, lab in ((A, C_NME, "MetAP-permissive"), (B, C_MET, "MetAP-nonpermissive")):
    v = np.sort(np.array([p["b"] for p in g]))
    ax0.step(v, np.arange(1, len(v) + 1) / len(v), color=c, lw=2,
             label=f"{lab} (n = {len(g)})")
    below[lab] = float(np.mean(v < 25))
ax0.set_xlabel("Selected-position burial on assembly (%)", fontsize=AX)
ax0.set_ylabel("Cumulative fraction of proteins", fontsize=AX)
ax0.axvline(25, color="#888888", lw=0.8, ls="--")
ax0.text(23, 0.055, "25% burial\nthreshold", fontsize=ANN, color="#666666",
         va="bottom", ha="right")
ax0.annotate("", xy=(25, below["MetAP-permissive"]), xytext=(25, below["MetAP-nonpermissive"]),
             arrowprops=dict(arrowstyle="<->", color=RED, lw=1.2, shrinkA=0, shrinkB=0))
ax0.text(31, 0.42,
         f"At 25%: {100 * below['MetAP-permissive']:.1f}% of permissive\n"
         f"and {100 * below['MetAP-nonpermissive']:.1f}% of nonpermissive\n"
         "positions remain below threshold",
         fontsize=ANN, color=RED, va="center", linespacing=1.5)
ax0.legend(fontsize=ANN, frameon=False, loc="lower right")
ax0.set_title("a   Burial distributions by sequence class", loc="left",
              fontsize=PANEL, fontweight="bold")
ax0.set_xlim(-2, 102); ax0.set_ylim(0, 1.02)

# --- b: burial by P1' residue
order = [(aa, aa) for aa in "ACGPSTVNQDEKRHFWYILM"]
rng = np.random.default_rng(0)
xs, meds, ns, cols, labs = [], [], [], [], []
for aa, nm in order:
    g = [p["b"] for p in keep if p["res2"] == aa]
    xs.append(len(xs)); meds.append(float(np.median(g))); ns.append(len(g))
    cols.append(C_NME if aa in SMALL else C_MET); labs.append(nm)
    v = np.array(g)
    ax1.scatter(rng.normal(xs[-1], 0.13, len(v)), v, s=5, alpha=0.30,
                color=cols[-1], edgecolors="none")
ax1.scatter(xs, meds, marker="_", s=170, color="black", zorder=4, linewidths=1.6)
ax1.set_xticks(xs)
# The residue letter and its n were a single two-line tick label, so one font size
# governed both and the n row collided at this tick pitch. They are now drawn
# separately: letters as tick labels, counts as smaller annotations beneath.
ax1.set_xticklabels(labs, fontsize=6.8)
for _x, _n in zip(xs, ns):
    ax1.annotate(str(_n), xy=(_x, 0), xycoords=("data", "axes fraction"),
                 xytext=(0, -14.5), textcoords="offset points",
                 ha="center", va="top", fontsize=4.9, annotation_clip=False)
ax1.set_ylabel("Burial on assembly (%)", fontsize=AX)
# labelpad clears the separately drawn n row, which the tick labels no longer occupy
ax1.set_xlabel("Penultimate residue (P1'), with n below", fontsize=AX, labelpad=11)
ax1.set_title("b   Burial by penultimate residue", loc="left",
              fontsize=PANEL, fontweight="bold")
ax1.legend(handles=[Patch(facecolor=C_NME, label="MetAP-permissive"),
                    Patch(facecolor=C_MET, label="MetAP-nonpermissive")],
           fontsize=ANN, frameon=False, loc="upper left", ncol=2, handlelength=1.0,
           columnspacing=1.0, handletextpad=0.5)
ax1.set_ylim(-4, 118)

# --- c: stratified odds ratios, exact conditional intervals
def orci(a, b):
    ka, na = sum(1 for x in a if x["b"] >= 25), len(a)
    kb, nb = sum(1 for x in b if x["b"] >= 25), len(b)
    r = odds_ratio([[ka, na - ka], [kb, nb - kb]], kind="conditional")
    ci = r.confidence_interval(0.95)
    return r.statistic, ci.low, ci.high, na, nb

bands = [(2, 3, "2\u20133 chains"), (4, 5, "4\u20135 chains"), (6, 11, "6\u201311 chains"),
         (12, 10 ** 6, "\u226512 chains"), (2, 10 ** 6, "all assemblies")]
rows = [(lab,) + orci([p for p in A if lo <= p["nch"] <= hi],
                      [p for p in B if lo <= p["nch"] <= hi]) for lo, hi, lab in bands]
y = np.arange(len(rows))[::-1]
for yy, (lab, o, l, u, na, nb) in zip(y, rows):
    c = "#222222" if lab == "all assemblies" else C_NME
    ax2.plot([l, u], [yy, yy], color=c, lw=1.6)
    ax2.plot([o], [yy], "o", color=c, ms=7)
    ax2.text(2.35, yy, f"{o:.2f} [{l:.2f}\u2013{u:.2f}]", fontsize=ANN, va="center", color="#333333")
ax2.axvline(1, color="#888888", lw=0.9, ls="--")
ax2.set_yticks(y)
ax2.set_yticklabels([r[0] for r in rows], fontsize=ANN)
ax2.set_xscale("log"); ax2.set_xlim(0.28, 6.6)
ax2.set_xticks([0.3, 0.5, 1, 2]); ax2.set_xticklabels(["0.3", "0.5", "1", "2"], fontsize=TICK)
ax2.xaxis.set_minor_formatter(NullFormatter()); ax2.xaxis.set_minor_locator(NullLocator())
ax2.set_xlabel("Odds ratio for burial $\\geq$25%\nMetAP-permissive vs nonpermissive", fontsize=AX)
ax2.set_title("c   Odds ratios by assembly-size stratum", loc="left",
              fontsize=PANEL, fontweight="bold")
ax2.tick_params(axis="y", length=0)

# --- d: burial vs alpha-amino contact distance
nA = sum(1 for p in A if p["b"] >= 25 and np.isfinite(p["dN"]) and p["dN"] <= 5)
nB = sum(1 for p in B if p["b"] >= 25 and np.isfinite(p["dN"]) and p["dN"] <= 5)
for g, c, lab in ((A, C_NME, "MetAP-permissive"), (B, C_MET, "MetAP-nonpermissive")):
    g2 = [p for p in g if np.isfinite(p["dN"])]
    ax3.scatter(np.clip([p["dN"] for p in g2], 0, 20), [p["b"] for p in g2],
                s=7, alpha=0.42, color=c, edgecolors="none", label=f"{lab} (n = {len(g2)})")
ax3.axhline(25, color=RED, lw=0.9, ls="--")
ax3.axvline(5, color="#888888", lw=0.9, ls="--")
ax3.add_patch(plt.Rectangle((0, 25), 5, 75, facecolor=RED, alpha=0.06, zorder=0))
ax3.text(6.0, 95, "High engagement:\n"
                   "burial $\\geq$25% and distance $\\leq$5 \u00c5",
         fontsize=ANN, color=RED, va="top", linespacing=1.5)
ax3.set_xlabel("Minimum measured-N distance to another\nprotein chain (\u00c5, clipped at 20)", fontsize=AX)
ax3.set_ylabel("Maximum burial on assembly (%)", fontsize=AX)
ax3.set_title("d   Interface-engagement geometry", loc="left",
              fontsize=PANEL, fontweight="bold")
ax3.legend(fontsize=ANN, frameon=False, loc="center right")
ax3.set_ylim(-4, 104); ax3.set_xlim(0, 20.5)

for ax in (ax0, ax1, ax2, ax3):
    for s in ("top", "right"):
        ax.spines[s].set_visible(False)
    ax.grid(color="#e8e8e8", lw=0.5)
    ax.set_axisbelow(True)
    ax.tick_params(labelsize=TICK)
# The adjusted-model result is stated in the legend and main text; omitting a
# duplicate footer here avoids crowding the enlarged lower-panel axis labels.
fig.savefig(OUT + "fig1_census.png", dpi=600, facecolor="white", bbox_inches="tight")
fig.savefig(OUT + "fig1_census.svg", facecolor="white", bbox_inches="tight")
plt.close(fig)
print("wrote fig1_census.png")

# Restore the established sizes for Figures 2-4. Figure 1 alone uses larger
# typography because its four-panel image is placed narrower in the manuscript.
ANN = 6.6
PANEL = 8.6
AX = 7.4
TICK = 6.6
plt.rcParams["font.size"] = TICK
plt.rcParams["xtick.labelsize"] = TICK
plt.rcParams["ytick.labelsize"] = TICK

# =============================================================== FIGURE 2  machines
def machine(g):
    g = g.split(";")[0]
    if re.match(r"PSM[ABCDEFG]", g): return "26S proteasome / assembly"
    if re.match(r"CCT\d|TCP1", g): return "TRiC/CCT chaperonin"
    if re.match(r"MED\d", g): return "Mediator"
    if re.match(r"SF3[AB]|SNRP|PRPF|LSM|EFTUD|SNU|SART|BUD31|CDC5L", g): return "Spliceosome"
    if re.match(r"COPS|CUL\d|DDB1|DDA1|RBX1|SKP1|ELOB|ELOC|FBXW", g): return "Cullin\u2013RING / CSN"
    if re.match(r"NDUF|UQCR|COX\d|SDH[ABCD]|ATP5|MT-", g): return "OXPHOS complexes"
    if re.match(r"RPL\d|RPS\d|RPLP|FAU|RPSA|MRPL|MRPS", g): return "Ribosomes (cyto + mito)"
    if re.match(r"NUP\d|SEH1L|SEC13|AAAS|RAE1|NDC1|POM121|TPR$", g): return "Nuclear pore scaffold"
    if re.match(r"POLR[123]", g): return "RNA polymerases"
    if re.match(r"EIF3|EIF2", g): return "eIF2 / eIF3"
    if re.match(r"ACT[BGA]|TUB[ABG]|ARPC|ACTR", g): return "Actin / tubulin systems"
    if re.match(r"ANAPC|CDC2|CDK\d|CCN[A-Z]|MCM\d|GINS", g): return "Cell cycle / replication"
    if re.match(r"KPN[AB]|TNPO|RAN$|RANBP|IPO\d|XPO\d", g): return "Soluble transport receptors"
    return None

by = collections.defaultdict(list)
for p in keep:
    m = machine(p["gene"])
    if m:
        by[m].append(p)
recs = []
for m, rs in by.items():
    if len(rs) < 4:
        continue
    nme = [r for r in rs if r["pred"] == "predicted NME"]
    risk = [r for r in nme if r["b"] >= 25]
    top = max(rs, key=lambda r: r["b"])
    deg = [r for r in nme if r["res2"] in PHI]
    recs.append((m, len(rs), 100 * len(nme) / len(rs),
                 100 * len(risk) / len(nme) if nme else np.nan,
                 top["b"], 100 * len(deg) / len(nme) if nme else np.nan,
                 top["gene"].split(";")[0]))
recs.sort(key=lambda t: -(t[3] if t[3] == t[3] else -1))
cols = ["Permissive\nsequence\nclass (%)", "Of those,\nburied\n$\\geq$ 25% (%)",
        "Maximum\npositional\nburial (%)", "Potential\nMet-$\\Phi$\nsequence (%)"]
M = np.array([[r[2], r[3], r[4], r[5]] for r in recs], dtype=float)
fig, ax = plt.subplots(figsize=(W, 3.9), layout="constrained")
im = ax.imshow(M, cmap="YlOrRd", vmin=0, vmax=100, aspect="auto")
ax.set_xticks(range(len(cols))); ax.set_xticklabels(cols, fontsize=ANN)
ax.set_yticks(range(len(recs)))
ax.set_yticklabels([f"{r[0]}  (n = {r[1]})" for r in recs], fontsize=ANN)
for i in range(M.shape[0]):
    for j in range(M.shape[1]):
        v = M[i, j]
        if v != v:
            ax.text(j, i, "\u2013", ha="center", va="center", fontsize=7.2, color="#666666")
        else:
            ax.text(j, i, f"{v:.0f}", ha="center", va="center", fontsize=7.2,
                    color="white" if v > 62 else "#222222",
                    fontweight="bold" if v > 62 else "normal")
for i, r in enumerate(recs):
    ax.text(M.shape[1] - 0.34, i, f"  {r[6]}", ha="left", va="center", fontsize=ANN, color="#444444")
ax.set_xlim(-0.5, M.shape[1] + 1.4)
ax.text(M.shape[1] - 0.34, -0.95, "  Most buried\n  subunit", ha="left", va="center",
        fontsize=ANN, color="#333333")
ax.tick_params(length=0)
for s in ax.spines.values():
    s.set_visible(False)
cb = fig.colorbar(im, ax=ax, fraction=0.030, pad=0.02, shrink=0.85)
cb.set_label("percent", fontsize=ANN); cb.ax.tick_params(labelsize=ANN)
fig.savefig(OUT + "fig2_machines.png", dpi=600, facecolor="white", bbox_inches="tight")
fig.savefig(OUT + "fig2_machines.svg", facecolor="white", bbox_inches="tight")
plt.close(fig)
print("wrote fig2_machines.png")
json.dump([{k: (v if v == v else None) for k, v in
            zip(["machine", "n", "pct_nme", "pct_risk_of_nme", "max_buried", "pct_degron", "top_gene"], r)}
           for r in recs], open(OUT + "machine_table.json", "w"), indent=1)

# =============================================================== FIGURES 3 and 4
obs = list(csv.DictReader(open(IN + "asm_observations.csv")))

def pick(pdb, gene):
    v = [o for o in obs if o["pdb"] == pdb and o["gene"].split(";")[0] == gene]
    return max(v, key=lambda o: float(o["buried_pct"])) if v else None

def best(gene):
    v = [o for o in obs if o["gene"].split(";")[0] == gene]
    return max(v, key=lambda o: float(o["buried_pct"]))["pdb"] if v else None

CMAP = {"excised": C_NME, "refractory": C_MET, "propeptide": "#6b6b6b", "retained": "#8fb3d9"}

ring = [("PSMA6 (\u03b11)", best("PSMA6"), "PSMA6", "retained"),
        ("PSMA2 (\u03b12)", best("PSMA2"), "PSMA2", "excised"),
        ("PSMA4 (\u03b13)", best("PSMA4"), "PSMA4", "retained"),
        ("PSMA7 (\u03b14)", best("PSMA7"), "PSMA7", "retained"),
        ("PSMA5 (\u03b15)", best("PSMA5"), "PSMA5", "refractory"),
        ("PSMA1 (\u03b16)", best("PSMA1"), "PSMA1", "refractory"),
        ("PSMA3 (\u03b17)", best("PSMA3"), "PSMA3", "excised"),
        ("PSMA3 (\u03b17)", "4r3o", "PSMA3", "excised"),
        ("PSMB3 (\u03b23)", best("PSMB3"), "PSMB3", "excised"),
        ("PSMB5 (\u03b25)", "8yix", "PSMB5", "propeptide"),
        ("PSMB7 (\u03b22)", "8yix", "PSMB7", "propeptide")]
recs = []
for lab, pdb, gene, kind in ring:
    o = pick(pdb, gene)
    if o:
        recs.append((lab, float(o["buried_pct"]), float(o["min_dist_alphaamino_N"]),
                     kind, pdb, o["resname"] + str(o["authnum"]), o["res2"]))
recs.sort(key=lambda r: -r[1])
fig, (a1, a2) = plt.subplots(1, 2, figsize=(W, 3.9), layout="constrained",
                             gridspec_kw=dict(width_ratios=[1.35, 1]))
fig.get_layout_engine().set(wspace=0.01)
y = np.arange(len(recs))[::-1]
cols = [CMAP[r[3]] for r in recs]
a1.barh(y, [r[1] for r in recs], color=cols, height=0.62, edgecolor="white")
for yy, r in zip(y, recs):
    a1.text(r[1] + 1.6, yy, f"{r[1]:.1f}%", va="center", fontsize=ANN, color="#333333")
a1.set_yticks(y)
a1.set_yticklabels([f"{r[0]}  {r[5]}, P1'={r[6]}  {r[4].upper()}" for r in recs], fontsize=ANN)
a1.set_xlim(0, 118)
a1.axvline(25, color=RED, ls="--", lw=1)
a1.set_xlabel("Burial of residue 1 in the assembly (%)", fontsize=AX)
a1.set_title("a   Depth of each terminus in the 20S particle", loc="left",
             fontsize=PANEL, fontweight="bold")
a2.scatter([r[2] for r in recs], y, color=cols, s=26, zorder=3)
for yy, r in zip(y, recs):
    a2.text(r[2] + 0.55, yy, f"{r[2]:.2f} \u00c5", va="center", fontsize=ANN, color="#333333",
            bbox=dict(facecolor="white", edgecolor="none", pad=0.8))
a2.axvline(5, color="#444444", ls="--", lw=1)
a2.set_yticks(y); a2.set_yticklabels([])
a2.set_xlim(0, 15)
a2.set_xlabel("\u03b1-amino N to nearest atom\nof another subunit (\u00c5)", fontsize=AX)
a2.set_title("b   \u03b1-amino contact geometry", loc="left",
             fontsize=PANEL, fontweight="bold")
for ax in (a1, a2):
    for s in ("top", "right"):
        ax.spines[s].set_visible(False)
    ax.tick_params(axis="y", length=0)
    ax.grid(axis="x", color="#e4e4e4", lw=0.7)
    ax.set_axisbelow(True)
fig.legend(handles=[Patch(facecolor=C_NME, label="MetAP-permissive P1', mature residue modelled"),
                    Patch(facecolor="#8fb3d9", label="MetAP-permissive P1', Met modelled"),
                    Patch(facecolor=C_MET, label="MetAP-nonpermissive P1'"),
                    Patch(facecolor="#6b6b6b", label="propeptide-bearing pro-form (excluded from statistics)")],
           fontsize=ANN, frameon=False, loc="outside lower center",
           handlelength=1.1, handletextpad=0.5, columnspacing=1.6, ncol=2)
fig.savefig(OUT + "fig3_alpha_ring.png", dpi=600, facecolor="white")
plt.close(fig)
print("wrote fig3_alpha_ring.png")

tr = [("KPNB1  importin-\u03b2", "6n88", "KPNB1", "refractory"),
      ("RAN  GTPase", "7mo1", "RAN", "excised"),
      ("KPNA2  importin-\u03b1", "9bfc", "KPNA2", "excised"),
      ("NUP188  pore scaffold", "5ijo", "NUP188", "excised"),
      ("NUP93  pore scaffold", "5ijo", "NUP93", "refractory"),
      ("SEH1L  pore scaffold", "5a9q", "SEH1L", "refractory"),
      ("NUP50  pore basket", "3tj3", "NUP50", "excised")]
recs = []
for lab, pdb, gene, kind in tr:
    o = pick(pdb, gene)
    if o:
        recs.append((lab, float(o["buried_pct"]), float(o["min_dist_alphaamino_N"]),
                     kind, pdb, int(o["n_chains"])))
recs.sort(key=lambda r: -r[1])
fig, (a1, a2) = plt.subplots(1, 2, figsize=(W, 2.9), layout="constrained",
                             gridspec_kw=dict(width_ratios=[1.35, 1]))
fig.get_layout_engine().set(wspace=0.01)
y = np.arange(len(recs))[::-1]
cols = [CMAP[r[3]] for r in recs]
a1.barh(y, [r[1] for r in recs], color=cols, height=0.62, edgecolor="white")
for yy, r in zip(y, recs):
    a1.text(r[1] + 1.6, yy, f"{r[1]:.1f}%", va="center", fontsize=ANN, color="#333333")
a1.set_yticks(y)
a1.set_yticklabels([f"{r[0]}  {r[4].upper()}, {r[5]} chains" for r in recs], fontsize=ANN)
a1.set_xlim(0, 118)
a1.axvline(25, color=RED, ls="--", lw=1)
a1.set_xlabel("Selected-position burial (%)", fontsize=AX)
a1.set_title("a   Pore scaffold versus soluble carriers", loc="left",
             fontsize=PANEL, fontweight="bold")
a2.scatter([min(r[2], 49) for r in recs], y, color=cols, s=26, zorder=3)
for yy, r in zip(y, recs):
    a2.text(min(r[2], 49) + 2.0, yy, f"{r[2]:.2f} \u00c5", va="center", fontsize=ANN, color="#333333",
            bbox=dict(facecolor="white", edgecolor="none", pad=0.8))
a2.axvline(5, color="#444444", ls="--", lw=1)
a2.set_yticks(y); a2.set_yticklabels([])
a2.set_xlim(0, 66)
a2.set_xlabel("Measured N to nearest atom\nof another protein chain (\u00c5)", fontsize=AX)
a2.set_title("b   Measured-N geometry", loc="left",
             fontsize=PANEL, fontweight="bold")
for ax in (a1, a2):
    for s in ("top", "right"):
        ax.spines[s].set_visible(False)
    ax.tick_params(axis="y", length=0)
    ax.grid(axis="x", color="#e4e4e4", lw=0.7)
    ax.set_axisbelow(True)
a1.legend(handles=[Patch(facecolor=C_NME, label="MetAP-permissive P1'"),
                   Patch(facecolor=C_MET, label="MetAP-nonpermissive P1'")],
          fontsize=ANN, frameon=False, loc="lower right", handlelength=1.1, handletextpad=0.5)
fig.savefig(OUT + "fig4_transport.png", dpi=600, facecolor="white", bbox_inches="tight")
fig.savefig(OUT + "fig4_transport.svg", facecolor="white", bbox_inches="tight")
plt.close(fig)
print("wrote fig4_transport.png")

# ---------------------------------------------------------------- audit trail
print("\nvalues annotated in the regenerated artwork")
print("  n analyzable                     %d" % len(keep))
print("  MetAP-permissive / nonpermissive %d / %d" % (len(A), len(B)))
print("  below 25%% burial                  %.1f%% / %.1f%%" % (100 * below["MetAP-permissive"],
                                                                100 * below["MetAP-nonpermissive"]))
print("  shaded quadrant                  %d / %d" % (nA, nB))
print("  dN resolved                      %d / %d" % (sum(1 for p in A if np.isfinite(p["dN"])),
                                                      sum(1 for p in B if np.isfinite(p["dN"]))))
for lab, o, l, u, na, nb in rows:
    print("  OR %-16s %.2f [%.2f-%.2f]  n = %d vs %d" % (lab, o, l, u, na, nb))
```
