## Supplementary_Data_8a for "Human protein complex interfaces reveal candidate assembly roles for initiator methionine excision": frozen_release_README.md

# Reproducibility deposit for the human N-terminal structural census

## Release 1.3.0 (2026-09-19)

Adds the cross-observation state-persistence analysis (data/Supplementary_Data_6_State_persistence.xlsx,
data/state_persistence_analysis.json) and the script that regenerates it
(code/state_persistence_analysis.py; run with --input-dir data). Adds the per-observation
workbook (data/Supplementary_Data_2_Per_observation_records.xlsx) that the script reads.
All 20 immunoglobulin and T-cell receptor constant-region entries are excluded at the
classification step (excluded_from_analysis = True in Supplementary Data 1), giving the
1,123-protein analyzable set and the 261-protein interface-engaged MetAP-permissive candidate
set used throughout the manuscript; Supplementary Data 3 contains exactly these 261 records.
Supplementary Data 4 gains the sheet "Ig-TCR removed primary" with the primary comparison on
n = 1,123 (Fisher odds ratio 0.662, p = 9.5e-4; size-adjusted odds ratio 0.652, p = 7.6e-4);
the pre-existing sheets remain the release-1.2.2 output on n = 1,143. Terminology across the
derived tables is harmonized to MetAP-permissive / MetAP-nonpermissive. Figures 1-5 are the
submitted versions; code/rebuild_figures_1_to_4.py is replaced by the plotting script whose
filter excludes both the excluded and unclassified rows. In data/asm_per_protein_master.csv the
20 Ig/TCR rows are relabelled from "unclassified" to "excluded: Ig/TCR constant region" (no other
field changes), so every deposited script that drops cls "excluded*" now operates on the
1,123-protein set. Supplementary Figure S1 (figures/Supplementary_Figure_S1.png) is regenerated
with the unchanged code/rebuild_supplementary_figure1.py on this relabelled table: 1,123
proteins over 6,305 retained assemblies, endpoints 41.3% at or above 25% burial and
size-adjusted odds ratio 0.652 (1.2.2: 1,143 proteins, 6,653 assemblies, 41.2%, 0.654). The
census inputs, the 22,291 chain-level observations and every value in the 1.2.2 tables are
unchanged.

Version DOI: https://doi.org/10.5281/zenodo.22847917
Concept DOI: https://doi.org/10.5281/zenodo.22129640


Analysis code, input snapshots, derived data and frozen final figures for the manuscript *A
structural census links penultimate-residue class to N-terminal burial in human protein
assemblies*.

The question the pipeline answers: for every human protein whose mature N-terminal residue is
resolved in a deposited multi-chain biological assembly, how buried is that residue by the rest of
the assembly, and how far is its free α-amino nitrogen from the nearest atom of another subunit?
Those two measurements are then compared between MetAP-permissive and MetAP-nonpermissive
penultimate-residue sequence classes. This sequence-class contrast is not treated as a direct
measurement of initiator-methionine removal: residue chemistry and processing class are correlated,
and the deposited analyses do not isolate a causal effect of excision.

## Contents

```
code/         pipeline, in execution order (see below)
inputs/       frozen snapshots of the two public input files
data/         derived tables and full statistical output
figures/      frozen final manuscript figures 1-5 plus legacy analysis figures
requirements.txt
LICENSE       MIT
```

## Reproducing the analysis

The scripts resolve their own locations, so the archive can be unpacked anywhere. Each one sets

```
BASE       = $NME_DIR, or the parent of code/ (the archive root)
INPUT_DIR  = BASE/inputs      frozen input snapshots, read only
WORK_DIR   = $NME_WORK, or BASE/work      everything the pipeline reads or writes, created on demand
```

A fresh run therefore regenerates its intermediates into `work/` and leaves the published tables in
`data/` untouched, so the two can be compared directly.

```bash
python -m venv .venv && source .venv/bin/activate
pip install -r requirements.txt
cd code
python step1_build_targets.py     # enumerate the target space from UniProt + SIFTS
python step2_select_download.py   # order entries by novel-accession yield, fetch mmCIF
python step3_geometry.py          # assemblies, mature-terminus ID, SASA, contact geometry
python step5_full.py              # full run over the ordered entry list
python step7_targeted.py          # re-run large machines with the atom ceiling raised
python step6_stats.py             # classification and all statistical tests
python step8_cascade.py           # downstream modification branches
python fig_asm.py                 # figures 1-4
python fig_cascade.py             # figure 5
```

`step4_stats.py` and `asm_test.py` are the earlier pilot statistics and the pipeline smoke test;
they are included for completeness but are not on the path to the reported numbers.

Structure downloads dominate the runtime. `step3_geometry.py` and `step5_full.py` write a
resumable cache, so an interrupted run can be restarted without refetching.

As a quick check that the archive is intact and the environment is correct, `step1_build_targets.py`
needs only the two frozen inputs, runs in about five seconds, and should print

```
human proteins parsed: 20376
chain segments covering the mature N-terminus: 111604
entries with >=2 chains: 19064 | heteromeric (>1 distinct UniProt): 11324
```

The three enumeration figures quoted in the Methods — 4,087 accessions, 19,064 multi-chain entries
and 101,894 chain-segment targets — are the counts restricted to entries with at least two chains,
recoverable from the `work/targets.json` this step writes.

## Input snapshots

`inputs/` contains the exact input files used, because both upstream sources change continuously
and the reported counts are only reproducible against a fixed snapshot:

| File | Source | Retrieved |
| --- | --- | --- |
| `human_nterm.tsv.gz` | UniProtKB human subset, with initiator-methionine, modified-residue, signal-peptide, propeptide, transit-peptide, sequence, length and gene-name fields | 19 August 2026 |
| `pdb_chain_uniprot.tsv.gz` | SIFTS `pdb_chain_uniprot` residue-level mapping | 19 August 2026 |

mmCIF files are fetched from RCSB at run time and are not redistributed here.

## Derived data

| File | Records | Content |
| --- | --- | --- |
| `asm_observations.csv` | 22,291 | one row per chain-level observation: PDB entry, assembly chain, source chain, accession, annotation state, residue identity, assembled and isolated SASA, burial, minimum residue distance, minimum α-amino nitrogen distance |
| `asm_per_protein_master.csv` | 1,191 | per-protein collapse (maximum-burial rule) with across-assembly summaries |
| `asm_at_risk_termini.csv` | 264 | predicted-excision termini buried ≥25%; no distance criterion |
| `asm_stats_full.txt` | — | complete statistical output, including all three collapse rules |
| `asm_stats_summary.json`, `asm_machine_table.json`, `asm_cascade_records.json` | — | machine-readable forms used by the figure scripts |

Two schema notes for anyone reusing these tables:

- `dN_min` in `asm_per_protein_master.csv` is a minimum taken **across assemblies** and is not
  comparable to a per-structure value. For any claim about a named structure, read
  `asm_observations.csv`.
- To ask whether a chain was modelled with methionine 1 still present, filter on `resname`
  (the observed residue) and not on `res1` (the residue expected from the UniProt sequence).

## Definitions used in classification

- **Excision predicted** when the penultimate residue is A, C, G, P, S, T or V.
- **Met-Φ degron on retention** predicted when the penultimate residue is L, F, Y, W, I, V or A.
- **Burial on assembly** = `100 × (1 − A_assembly / A_isolated)` for the mature N-terminal residue,
  total residue area, FreeSASA with explicit element radii (C 1.70, N 1.55, O 1.52, S and Se 1.80,
  P 1.80 Å; 1.70 Å otherwise).
- **NTMT recognition motif**, strict canonical form: [A/P/S/G] at mature position 1, P at 2, K at 3.
  NTMT1 and NTMT2 also accept R at position 3; the manuscript reports the effect of relaxing the
  motif as a sensitivity test rather than adopting it, and the strict form excludes CENP-A by
  definition.
- **Chains are dropped, not guessed.** A candidate mature-start position is accepted only if the
  observed residue name at that position, or within two positions of it, matches the residue
  expected from the UniProt sequence.

## Machine-level grouping

Figure 4 aggregates subunits by prefix matching on the primary gene symbol rather than by a
curated complex membership list. This is reproducible and free of curator judgement, but a subunit
whose symbol departs from its family convention is missed, so each row's denominator is a lower
bound on complex membership. The patterns, applied in this order, are:

| Row | Pattern on the gene symbol |
| --- | --- |
| 26S proteasome / assembly | `PSM[ABCDEFG]` |
| TRiC/CCT chaperonin | `CCT\d`, `TCP1` |
| Mediator | `MED\d` |
| Spliceosome | `SF3[AB]`, `SNRP`, `PRPF`, `LSM`, `EFTUD`, `SNU`, `SART`, `BUD31`, `CDC5L` |
| Cullin–RING / CSN | `COPS`, `CUL\d`, `DDB1`, `DDA1`, `RBX1`, `SKP1`, `ELOB`, `ELOC`, `FBXW` |
| OXPHOS complexes | `NDUF`, `UQCR`, `COX\d`, `SDH[ABCD]`, `ATP5`, `MT-` |
| Ribosomes (cyto + mito) | `RPL\d`, `RPS\d`, `RPLP`, `FAU`, `RPSA`, `MRPL`, `MRPS` |
| Nuclear pore scaffold | `NUP\d`, `SEH1L`, `SEC13`, `AAAS`, `RAE1`, `NDC1`, `POM121`, `TPR$` |
| RNA polymerases | `POLR[123]` |
| eIF2 / eIF3 | `EIF3`, `EIF2` |
| Actin / tubulin systems | `ACT[BGA]`, `TUB[ABG]`, `ARPC`, `ACTR` |
| Cell cycle / replication | `ANAPC`, `CDC2`, `CDK\d`, `CCN[A-Z]`, `MCM\d`, `GINS` |
| Soluble transport receptors | `KPN[AB]`, `TNPO`, `RAN$`, `RANBP`, `IPO\d`, `XPO\d` |

Patterns are applied with `re.match`, so they anchor at the start of the symbol. Rows are scored
only where at least four member proteins are present, which is why the soluble transport receptors
are defined here but do not appear as a row in Figure 4.

## Figure provenance

`rebuild_figures_1_to_4.py` rebuilds the final content of Figures 1–4 from the canonical `data/` tables by default;
the script includes the all-20-residue Figure 1 panel and the final MetAP-permissive terminology.
`scan_methionine.py` regenerates the 53-position score table from frozen biological-assembly
coordinates, and `rebuild_figure5.py` renders Figure 5 from that recomputed table. The frozen files
`figures/Figure_1.png` through `figures/Figure_5.png` are the exact images embedded in the final
manuscript. Figures 1, 2 and 5 were replaced on 25 August 2026 to match the final manuscript.

## How to cite

This archive:

> Chang, Y.-H. *Reproducibility deposit for “A structural census links penultimate-residue class to
> N-terminal burial in human protein assemblies”* (version 1.3.0). Zenodo (2026).
> https://doi.org/10.5281/zenodo.22847917

To cite whichever release is current rather than this one specifically, use the concept DOI
https://doi.org/10.5281/zenodo.22129640. The preceding releases remain citable: 1.2.2 at
https://doi.org/10.5281/zenodo.22233711 and 1.2.1 at https://doi.org/10.5281/zenodo.22129641.

The accompanying paper:

> Chang, Y.-H. *A structural census links penultimate-residue class to N-terminal burial in human
> protein assemblies*. bioRxiv (2026). https://doi.org/10.64898/2026.08.31.748389 (preprint;
> the manuscript is under consideration at Communications Biology)

Machine-readable metadata for this archive is in `CITATION.cff`, which records the concept DOI, this
release's version DOI `10.5281/zenodo.22847917`, the preceding version DOIs, the release
date 2026-09-19, the ORCID `0000-0002-1583-0627` and the contact address
``. The version DOI could be written inside the archive it identifies
because Zenodo allows a DOI to be reserved before its record is published. The only remaining
placeholder was the companion paper's DOI; the bioRxiv preprint DOI is now given above and the journal DOI will be added when available.

## Licence

Code and derived data are released under the MIT licence. The input snapshots remain under the terms
of their original providers: UniProtKB (CC BY 4.0), SIFTS/PDBe and the Protein Data Bank.


## Figure 5 provenance
The published Figure 5 is generated end to end from 11 frozen RCSB biological-assembly mmCIF files
in `inputs/figure5_assemblies/`, the target manifest `data/fig5_targets.csv`,
`code/scan_methionine.py`, and `code/rebuild_figure5.py`. SHA-256 checksums for every coordinate
file are deposited with the coordinates. The scan evaluates 34,992 conformers at each of 28
observed-Met controls and 25 MetAP-permissive-class tests. Fixed quality-control criteria yield
finite scores for 26 controls and 23 tests; two positions in each group have no conformer passing
the intrachain-overlap criterion and remain transparently unscored. The primary continuous
comparison among scoreable positions buried by at least 50% gives medians of −1.12 Å for controls
and −0.30 Å for tests (17 versus 10 positions; two-sided Mann–Whitney p = 0.00490).

The earlier frozen-score analysis is retained only in `audit/legacy_figure5/` for provenance and
is not used by the manuscript or current figure.

The per-protein table now distinguishes `dN_min` (the minimum across all observations) from
`dN_at_max_burial` (the distance aligned to the PDB entry and chain supplying maximum burial).

## Robustness analyses and candidate map

`code/run_robustness_analyses.py` reads the frozen canonical tables from `data/` by default and produces
three prespecified sensitivity analyses in the same directory (or in `$NME_OUT`): logistic models adjusted for observation multiplicity,
leave-one-residue-out analyses, and exclusion of predicted-excision proteins modeled with retained
MET/MSE. Their machine-readable outputs are `data/multiplicity_models.csv`,
`data/leave_one_residue_out.csv`, and `data/met_mse_exclusion.csv`.

`data/candidate_to_experiment_table.csv` contains the selected proteins, representative PDB
entries, census measurements, mechanistic hypotheses, and proposed follow-up assays reported in
Supplementary Data 5. These files use only the canonical dataset; no legacy census result is
substituted into the analysis.

The exact submitted workbook is also included as
`data/Supplementary_Data_5_Robustness_analyses_and_candidate_map.xlsx`. Its five worksheets cover
the description, observation-multiplicity models, leave-one-residue-out analysis, MET/MSE
exclusion, and candidate-to-experiment map.

## Alignment with the submitted manuscript

The manuscript submitted to Communications Biology (September 2026) uses the same canonical census
(7,246 assemblies, 22,291 chain-level observations, 1,191 proteins) with the 20 immunoglobulin and
T-cell receptor constant-region entries excluded at classification, giving 1,123 analyzable proteins
and 261 interface-engaged MetAP-permissive candidates (release 1.3.0); the 25 August 2026 manuscript
reported 1,143 and 264 on release 1.2.2. Robustness outputs and the fully recomputed Figure 5
statistics are unchanged. It adds fuller biological interpretation
of proteasome, CCT/TRiC, histone-tail recognition, spliceosome, OXPHOS/MetAP1D and downstream
N-terminal modification examples; those interpretations draw on the deposited per-protein,
per-observation, machine-level and candidate-map records and do not introduce a second dataset.
The final text also explicitly separates assembly burial from peptide-recognition geometry and
adds the structural-coverage/chain-length limitation. See `MANUSCRIPT_ALIGNMENT_AUDIT.md` for the
claim-to-file audit.

## Figure 5 reproducibility audit

`FIGURE5_REPRODUCIBILITY_AUDIT.md` records the final pre-submission audit. Figure 5 now passes
end-to-end reproducibility: frozen coordinates and a frozen target manifest generate the deposited
position-level table, statistics and artwork with the deposited scripts. The four unscored
positions are determined by the fixed intrachain-overlap quality-control rule and are retained in
the output table with empty scores rather than removed from the target manifest.

### Quick rebuilds from deposited tables

From the archive root, the following commands require no legacy working-directory names:

```bash
python code/rebuild_figures_1_to_4.py
python code/run_robustness_analyses.py
```

## Version 1.2.1 correction


This release aligns named-structure alpha-amino distances to the same PDB observation and assembly chain that supplies maximum burial, adds that aligned field to the submitted per-protein workbook, and removes legacy working-directory assumptions from the quick rebuild scripts. The census counts and statistical conclusions are unchanged.

## Version 1.2.2 correction

Version 1.2.1 aligned the named-structure alpha-amino distance to the observation that supplies
maximum burial, but added the aligned field to the per-protein workbook only. This release completes
that change everywhere the paired values appear:

- `data/asm_at_risk_termini.csv` gains `dN_at_max_burial`, so the interface-engaged table carries the
  same aligned field as the per-protein master table.
- `code/step6_stats.py` now computes `dN_at_max_burial` inside the per-protein collapse and writes it
  to both output tables, and comments state explicitly that `dN_min` and `dN_median` are
  protein-level statistics that are not tied to the entry named in the `pdb` column.
- Supplementary Data 3 gains the aligned column and the clarified definitions; the at-risk extremes
  sheet of Supplementary Data 4 renames its distance column to name the protein-level minimum
  explicitly and adds the aligned distance beside it.
- `figures/Figure_1.png` is replaced by the corrected export used in the manuscript. In the 1.2.1
  artwork the residue letter and its n were a single two-line tick label in panel b, so one font
  size governed both and the counts row collided at the panel's tick pitch. The counts are now set
  smaller than the letters. Changed pixels are confined to y 1422-1468, x 2640-4162, the counts row
  only; every other pixel of Figure 1 is identical to 1.2.1, and no plotted value moves.
  `code/rebuild_figures_1_to_4.py` and `code/fig_asm.py` now draw the letters as tick labels and the
  counts as separate smaller annotations, so a rerun reproduces the corrected layout.
- `code/step4_stats.py`, the retired pilot script, now carries a banner saying so.
- `CITATION.cff` records release 1.2.2 and the release date, and its `cff-version` field is
  corrected to the CFF schema version 1.2.0, which release 1.2.1 had overwritten with the release
  number.

Worked example of the ambiguity this removes. PSMA6 (P60900) is reported in the manuscript as 98.5%
buried at 3.51 A in 8TM4 chain G1. In the 1.2.1 interface-engaged table the same row read
`pdb = 8tm4`, `buried_pct = 98.5`, `dN_min = 2.52`, inviting the reading that the alpha-amino
nitrogen sits 2.52 A from a partner chain in 8TM4. It does not: 2.52 A is this protein's smallest
distance over all 12 of its observations and occurs in 8YIZ chain U1, where burial is 79.6%. Both
numbers were correct under their own definitions; only their adjacency was misleading. The row now
carries `dN_at_max_burial = 3.51`, matching the text and Figure 3.

No census count, test statistic, odds ratio, p value or figure changes in this release: 7,246
assemblies, 22,291 chain-level observations, 1,191 proteins, 1,143 analyzable proteins and 264
interface-engaged MetAP-permissive candidates, all unchanged. Verified by recomputing the aligned
distance for all 1,191 proteins from `data/asm_observations.csv` and comparing against the published
1.2.1 per-protein workbook: 1,180 numeric values agree, 11 are legitimately blank, no mismatches.
