## Supplementary_Data_8a for "Human protein complex interfaces reveal candidate assembly roles for initiator methionine excision": original_recurrence_analysis.py

```
#!/usr/bin/env python3
"""State-persistence analysis of interface-engaged MetAP-permissive N-termini.

Reproduces Supplementary Data 6 and the state-persistence numbers reported in the
main text from two deposited tables:

  Supplementary Data 2  (per-observation records, one row per assembly chain)
  Supplementary Data 3  (interface-engaged termini, one row per candidate protein)

Usage (run from the submission package directory, or pass --input-dir):

    python 16_State_persistence_analysis_script.py \
        --input-dir . \
        --json-out  state_persistence_analysis.json \
        --xlsx-out  15_Supplementary_Data_6_State_persistence.xlsx

All paths are relative to the current working directory unless given as absolute
paths. No hard-coded machine-specific locations are used.

Terminology: "MetAP-permissive" denotes the sequence class (Ala/Cys/Gly/Pro/Ser/Thr/Val
at residue 2). It describes predicted biochemical permissiveness and is not evidence
that the initiator methionine was removed in any specific structure.
"""
from __future__ import annotations

import argparse
from copy import copy
import json
import math
from pathlib import Path

import pandas as pd

# ---------------------------------------------------------------------------
# Candidate exclusions
# ---------------------------------------------------------------------------
# All 20 immunoglobulin and T-cell receptor constant-region entries are excluded
# upstream (excluded_from_analysis = True in Supplementary Data 1) and are absent
# from Supplementary Data 3, so no candidate-level exclusion is applied here.
# ---------------------------------------------------------------------------

# ---------------------------------------------------------------------------
# Machine grouping by gene-symbol prefix (see Methods, "Machine-level grouping")
# ---------------------------------------------------------------------------
GROUP_RULES = {
    'CCT/TRiC': lambda g: g in {'CCT2', 'CCT3', 'CCT4', 'CCT5', 'CCT6A', 'CCT6B', 'CCT7', 'CCT8', 'TCP1'},
    'Proteasome / assembly': lambda g: g.startswith(('PSMA', 'PSMB', 'PSMC', 'PSMD', 'PSME', 'PSMF', 'PSMG')),
    'Spliceosome': lambda g: g.startswith(('SNR', 'SF3', 'PRPF', 'LSM', 'U2AF', 'CDC5L', 'BCAS2', 'PLRG1', 'CRNKL1')),
    'Ribosome': lambda g: g.startswith(('RPL', 'RPS', 'MRPL', 'MRPS')) or g == 'FAU',
    # Nuclear transport: nucleoporins (NUP*), importins (KPNA*, KPNB1), Ran-cycle
    # components (RAN, RANBP*, RANGAP1) and the explicitly named pore/Ran-cycle
    # proteins NDC1, TPR and RCC1 (the Ran GEF). RCC1 is included deliberately
    # because it is a Ran-cycle regulator, not a pore scaffold component.
    'Nuclear transport': lambda g: g.startswith(('NUP', 'KPNA', 'KPNB', 'RAN')) or g in {'NDC1', 'TPR', 'RCC1'},
    'Mediator': lambda g: g.startswith('MED'),
    'OXPHOS, nuclear encoded': lambda g: g.startswith(('NDUF', 'UQCR', 'COX', 'SDH', 'ATP5')) and not g.startswith('MT-'),
}
GROUP_ORDER = list(GROUP_RULES)

FOCUS_GENES = ['CCT2', 'CCT5', 'CCT6A', 'CCT8', 'PSMA2', 'PSMA3', 'PSMA4', 'PSMA6', 'PSMA7', 'PSMG1', 'RAN', 'RCC1',
               'SF3B4', 'SF3B5', 'SNRNP70', 'SNRPD3', 'SNRPF', 'PRPF19', 'MED21', 'RPL14', 'RPL15', 'RPL18', 'RPS18',
               'RPS27', 'RPS29', 'RPS3', 'RPS4X', 'RPS9', 'RPSA']

OUTPUT_COLUMNS = ['gene', 'acc', 'pdb', 'n_obs', 'n_entries', 'unique_pdb_entries', 'resname', 'buried_pct', 'buried_median',
                  'fraction_observations_engaged', 'pdb_collapsed_engagement_fraction', 'fraction_observations_close',
                  'pdb_collapsed_close_fraction', 'observation_burial_min', 'observation_burial_max', 'dN_min', 'dN_median',
                  'dN_at_max_burial', 'nt_acetyl', 'res2', 'res3', 'state_class', 'pdb_collapsed_state_class',
                  'chain_vs_pdb_class_concordant', 'leave_one_pdb_out_stable', 'persistent_close_contact']

PERSISTENT = 'persistent engagement (>=75%)'
INTERMITTENT = 'intermittent engagement (25-<75%)'
RARE = 'rare-state engagement (<25%)'

DEFINITIONS = {
    'persistent': 'at least 75% of all observations have burial >=25%',
    'intermittent': '25% to less than 75% of observations have burial >=25%',
    'rare_state': 'less than 25% of observations have burial >=25%',
    'persistent_close_contact': 'persistent engagement plus alpha-amino-nitrogen distance <=5 A in at least 75% of observations',
    'pdb_collapsed_fraction': 'fraction of unique PDB entries whose within-entry median burial is >=25%; reduces weighting by repeated symmetry-related chains',
    'leave_one_pdb_out_stable': 'whether the PDB-collapsed state class is unchanged after removal of every single PDB entry; undefined for one-entry candidates',
    'n_obs': 'number of chain-level observations (assembly chains) contributing to the candidate',
    'n_entries': 'number of distinct PDB entries contributing to the candidate (from Supplementary Data 3)',
    'sequence_class': 'MetAP-permissive = Ala/Cys/Gly/Pro/Ser/Thr/Val at residue 2 (predicted permissiveness, not evidence of excision in a given structure)',
}


def gene_tokens(value) -> list[str]:
    return [x.strip().upper() for x in str(value).split(';') if x.strip()]


def in_group(value, group) -> bool:
    return any(GROUP_RULES[group](g) for g in gene_tokens(value))


def classify_fraction(value: float) -> str:
    if value >= 0.75:
        return PERSISTENT
    if value >= 0.25:
        return INTERMITTENT
    return RARE


def clean(x):
    if isinstance(x, float) and (math.isnan(x) or math.isinf(x)):
        return None
    return x.item() if hasattr(x, 'item') else x


def records(frame: pd.DataFrame) -> list[dict]:
    return [{k: clean(v) for k, v in row.items()} for row in frame.to_dict('records')]


def find_file(input_dir: Path, pattern: str) -> Path:
    matches = sorted(input_dir.glob(pattern))
    if not matches:
        raise FileNotFoundError(f'No file matching {pattern!r} in {input_dir}')
    return matches[0]


def run(input_dir: Path) -> dict:
    d3_path = find_file(input_dir, '*Supplementary_Data_3*.xlsx')
    d2_path = find_file(input_dir, '*Supplementary_Data_2*.xlsx')
    # Supplementary Data 3 may carry either the legacy sheet name or the current one.
    sheet_names = pd.ExcelFile(d3_path).sheet_names
    data_sheet = next((s for s in sheet_names if s.lower().startswith(('interface', 'at-risk'))), sheet_names[0])
    df = pd.read_excel(d3_path, sheet_name=data_sheet)
    obs = pd.read_excel(d2_path, sheet_name='Per-observation records')

    n_candidates = len(df)
    obs = obs[obs['acc'].isin(df['acc'])].copy()

    obs_metrics = obs.groupby('acc').agg(
        fraction_observations_engaged=('buried_pct', lambda x: float((x >= 25).mean())),
        fraction_observations_close=('min_dist_alphaamino_N', lambda x: float((x <= 5).mean())),
        observation_burial_min=('buried_pct', 'min'),
        observation_burial_max=('buried_pct', 'max'),
    ).reset_index()

    # One median record per accession and PDB entry limits weighting by duplicate chains.
    pdb_observations = obs.groupby(['acc', 'pdb'], as_index=False).agg(
        pdb_median_burial_percent=('buried_pct', 'median'),
        pdb_median_alphaamino_N_distance=('min_dist_alphaamino_N', 'median'),
        chains_in_entry=('pdb', 'size'),
    )
    pdb_metrics = pdb_observations.groupby('acc').agg(
        unique_pdb_entries=('pdb', 'size'),
        pdb_collapsed_engagement_fraction=('pdb_median_burial_percent', lambda x: float((x >= 25).mean())),
        pdb_collapsed_close_fraction=('pdb_median_alphaamino_N_distance', lambda x: float((x <= 5).mean())),
    ).reset_index()

    df = df.merge(obs_metrics, on='acc', how='left').merge(pdb_metrics, on='acc', how='left')
    df['state_class'] = df['fraction_observations_engaged'].map(classify_fraction)
    df['persistent_engagement'] = df['fraction_observations_engaged'] >= 0.75
    df['persistent_close_contact'] = df['persistent_engagement'] & (df['fraction_observations_close'] >= 0.75)
    df['pdb_collapsed_state_class'] = df['pdb_collapsed_engagement_fraction'].map(classify_fraction)
    df['chain_vs_pdb_class_concordant'] = df['state_class'] == df['pdb_collapsed_state_class']

    def leave_one_pdb_out_stable(accession):
        rows = pdb_observations[pdb_observations['acc'] == accession]
        if len(rows) < 2:
            return None
        full_class = classify_fraction(float((rows['pdb_median_burial_percent'] >= 25).mean()))
        return all(classify_fraction(float((rows.drop(index=i)['pdb_median_burial_percent'] >= 25).mean())) == full_class
                   for i in rows.index)

    df['leave_one_pdb_out_stable'] = df['acc'].map(leave_one_pdb_out_stable)

    raw_props = df['state_class'].value_counts(normalize=True)
    pdb_props = df['pdb_collapsed_state_class'].value_counts(normalize=True)
    observed_agreement = float(df['chain_vs_pdb_class_concordant'].mean())
    expected_agreement = sum(float(raw_props.get(c, 0) * pdb_props.get(c, 0)) for c in set(raw_props.index) | set(pdb_props.index))
    kappa = (observed_agreement - expected_agreement) / (1 - expected_agreement)
    spearman = float(df['fraction_observations_engaged'].corr(df['pdb_collapsed_engagement_fraction'], method='spearman'))

    threshold_rows = []
    for threshold in [20, 25, 30]:
        fractions = pdb_observations.groupby('acc')['pdb_median_burial_percent'].apply(lambda x: float((x >= threshold).mean()))
        classes = fractions.map(classify_fraction)
        threshold_rows.append({'burial_threshold_percent': threshold,
                               'persistent_candidates': int((classes == PERSISTENT).sum()),
                               'intermittent_candidates': int((classes == INTERMITTENT).sum()),
                               'rare_state_candidates': int((classes == RARE).sum())})

    group_rows = []
    for group in GROUP_ORDER:
        sub = df[df['gene'].map(lambda x: in_group(x, group))]
        group_rows.append({'group': group, 'interface_engaged_candidates': int(len(sub)),
                           'persistent_candidates': int(sub['persistent_engagement'].sum()),
                           'persistent_percent': round(100 * sub['persistent_engagement'].mean(), 1) if len(sub) else None,
                           'persistent_close_contact_candidates': int(sub['persistent_close_contact'].sum()),
                           'median_of_maximum_burial_percent': round(float(sub['buried_pct'].median()), 1) if len(sub) else None,
                           'median_cross_observation_burial_percent': round(float(sub['buried_median'].median()), 1) if len(sub) else None})

    n_two = int(df['leave_one_pdb_out_stable'].notna().sum())
    n_stable = int((df['leave_one_pdb_out_stable'] == True).sum())  # noqa: E712
    result = {
        'inputs': {'supplementary_data_3': d3_path.name, 'supplementary_data_2': d2_path.name,
                   'candidates_in_supplementary_data_3': n_candidates},
        'overall': {'interface_engaged_candidates': int(len(df)),
                    'persistent_candidates': int(df['persistent_engagement'].sum()),
                    'persistent_percent': round(100 * df['persistent_engagement'].mean(), 1),
                    'intermittent_candidates': int((df['state_class'] == INTERMITTENT).sum()),
                    'rare_state_candidates': int((df['state_class'] == RARE).sum()),
                    'persistent_close_contact_candidates': int(df['persistent_close_contact'].sum()),
                    'persistent_close_contact_percent': round(100 * df['persistent_close_contact'].mean(), 1),
                    'deeply_persistent_candidates': int((df['buried_median'] >= 50).sum()),
                    'deeply_persistent_percent': round(100 * (df['buried_median'] >= 50).mean(), 1)},
        'robustness': {'chain_vs_pdb_class_concordant_candidates': int(df['chain_vs_pdb_class_concordant'].sum()),
                       'chain_vs_pdb_class_concordance_percent': round(100 * observed_agreement, 1),
                       'cohens_kappa': round(kappa, 3),
                       'spearman_rho': round(spearman, 3),
                       'pdb_collapsed_persistent_candidates': int((df['pdb_collapsed_state_class'] == PERSISTENT).sum()),
                       'pdb_collapsed_intermittent_candidates': int((df['pdb_collapsed_state_class'] == INTERMITTENT).sum()),
                       'pdb_collapsed_rare_state_candidates': int((df['pdb_collapsed_state_class'] == RARE).sum()),
                       'candidates_with_at_least_two_pdb_entries': n_two,
                       'leave_one_pdb_out_stable_candidates': n_stable,
                       'leave_one_pdb_out_stable_percent': round(100 * n_stable / n_two, 1) if n_two else None},
        'threshold_sensitivity': threshold_rows,
        'groups': group_rows,
        'focus_candidates': records(df[df['gene'].astype(str).isin(FOCUS_GENES)][OUTPUT_COLUMNS].sort_values('gene')),
        'all_candidate_robustness': records(df[OUTPUT_COLUMNS].sort_values(['gene', 'acc'])),
        'definitions': DEFINITIONS,
    }
    return result


def write_xlsx(result: dict, path: Path) -> None:
    o, r = result['overall'], result['robustness']
    summary = pd.DataFrame([
        ('Interface-engaged candidates', o['interface_engaged_candidates']),
        ('Persistent engagement (>=75% observations)', o['persistent_candidates']),
        ('Intermittent engagement (25-<75%)', o['intermittent_candidates']),
        ('Rare-state engagement (<25%)', o['rare_state_candidates']),
        ('Persistent engagement plus close contact', o['persistent_close_contact_candidates']),
        ('Persistent engagement percent', o['persistent_percent']),
        ('Persistent close-contact percent', o['persistent_close_contact_percent']),
    ], columns=['metric', 'value'])
    robustness = pd.DataFrame(list(r.items()), columns=['metric', 'value'])
    definitions = pd.DataFrame(list(result['definitions'].items()), columns=['term', 'definition'])
    with pd.ExcelWriter(path, engine='openpyxl') as xw:
        summary.to_excel(xw, sheet_name='Summary', index=False)
        robustness.to_excel(xw, sheet_name='Robustness summary', index=False)
        pd.DataFrame(result['threshold_sensitivity']).to_excel(xw, sheet_name='Threshold sensitivity', index=False)
        pd.DataFrame(result['groups']).to_excel(xw, sheet_name='Machine groups', index=False)
        pd.DataFrame(result['focus_candidates']).to_excel(xw, sheet_name='Selected candidates', index=False)
        pd.DataFrame(result['all_candidate_robustness']).to_excel(xw, sheet_name='All candidate robustness', index=False)
        definitions.to_excel(xw, sheet_name='Definitions', index=False)
        for ws in xw.book.worksheets:
            for col in ws.columns:
                width = max(len(str(c.value)) if c.value is not None else 0 for c in col)
                ws.column_dimensions[col[0].column_letter].width = min(max(12, width + 2), 60)
            for c in ws[1]:
                f = copy(c.font); f.bold = True; c.font = f


def main() -> None:
    ap = argparse.ArgumentParser(description=__doc__, formatter_class=argparse.RawDescriptionHelpFormatter)
    ap.add_argument('--input-dir', type=Path, default=Path('.'),
                    help='directory containing Supplementary Data 2 and 3 (default: current directory)')
    ap.add_argument('--json-out', type=Path, default=Path('state_persistence_analysis.json'))
    ap.add_argument('--xlsx-out', type=Path, default=None,
                    help='optional path for a rebuilt Supplementary Data 6 workbook')
    args = ap.parse_args()

    result = run(args.input_dir)
    args.json_out.write_text(json.dumps(result, indent=2), encoding='utf-8')
    if args.xlsx_out:
        write_xlsx(result, args.xlsx_out)
    print(json.dumps({'inputs': result['inputs'], 'overall': result['overall'], 'robustness': result['robustness'],
                      'threshold_sensitivity': result['threshold_sensitivity'], 'groups': result['groups']}, indent=2))


if __name__ == '__main__':
    main()
```
