## Supplementary_Data_8a for "Human protein complex interfaces reveal candidate assembly roles for initiator methionine excision": audit.py

```
"""Non-destructive targeted audit of the frozen Paper 1 release.
Run: python paper1_audit/audit.py
Outputs are exploratory audit results, not a replacement biological dataset.
"""
from pathlib import Path
import csv, gzip, hashlib, json, re, sys
from collections import defaultdict
import numpy as np
import pandas as pd
from scipy.stats import fisher_exact, mannwhitneyu
import statsmodels.api as sm
import gemmi

ROOT = Path(__file__).resolve().parent
B = ROOT / "frozen/nterm-census-1.3.0"
OUT = ROOT / "results"
OUT.mkdir(exist_ok=True)
p = pd.read_csv(B / "data/asm_per_protein_master.csv")
o = pd.read_csv(B / "data/asm_observations.csv")
u = pd.read_csv(B / "inputs/human_nterm.tsv.gz", sep="\t").fillna("")
uid = u.set_index("Entry").to_dict("index")
keep = p[~p.cls.str.startswith("excluded")].copy()
o["analyzable"] = o.acc.isin(keep.acc)
o["permissive"] = o.res2.isin(list("AGSTCPV"))
needed = set(zip(o.pdb, o.chain, o.acc))
segments = defaultdict(list)
with gzip.open(B / "inputs/pdb_chain_uniprot.tsv.gz", "rt") as f:
    for r in csv.DictReader((x for x in f if not x.startswith("#")), delimiter="\t"):
        key = (r["PDB"].lower(), r["CHAIN"], r["SP_PRIMARY"])
        if key in needed:
            segments[key].append(r)

def mapping(r):
    ss = segments[(r.pdb, r.chain, r.acc)]
    candidates, intervals = [], []
    for s in ss:
        try:
            start, end = int(s["SP_BEG"]), int(s["SP_END"])
            intervals.append((start, end))
            m = re.match(r"^(-?\d+)", s["PDB_BEG"])
            if start <= r.mature_start and m:
                candidates.append(int(m.group(1)) + r.mature_start - start)
        except ValueError:
            pass
    offsets = [int(r.authnum) - x for x in candidates]
    exact = 0 in offsets
    allowed = [x for x in offsets if x in [0, 1, -1, 2, -2]]
    status = "exact_endpoint_offset" if exact else (
        "shifted_1_or_2" if allowed else "unreconciled")
    # Union of SIFTS canonical intervals is a construct-span proxy, not modeled coverage.
    covered = set()
    for lo, hi in intervals:
        covered.update(range(lo, hi + 1))
    length = int(uid[r.acc]["Length"])
    return dict(mapping_status=status, candidate_author_numbers=";".join(map(str,candidates)),
                accepted_offset=(0 if exact else allowed[0]) if allowed else np.nan,
                canonical_length=length, sifts_union_residues=len(covered),
                sifts_span_fraction=len(covered)/length,
                sifts_short_span_flag=len(covered)<30,
                sifts_low_span_flag=len(covered)/length<0.25,
                frozen_init_annotation=uid[r.acc]["Initiator methionine"])

audit = pd.concat([o.reset_index(drop=True),
                  pd.DataFrame([mapping(r) for r in o.itertuples()])], axis=1)
audit.to_csv(OUT / "observation_processing_mapping_audit.csv", index=False)
pa = p.merge(audit[["acc","pdb","asm_chain","mapping_status","accepted_offset",
                   "canonical_length","sifts_union_residues","sifts_span_fraction",
                   "frozen_init_annotation"]], on=["acc","pdb","asm_chain"],
             how="left", validate="one_to_one")
pa.to_csv(OUT / "protein_selected_record_audit.csv", index=False)

def compare(df, label):
    df = df.copy()
    x = df.res2.isin(list("AGSTCPV"))
    y = df.buried_pct >= 25
    a,b,c,d = int((x&y).sum()),int((x&~y).sum()),int((~x&y).sum()),int((~x&~y).sum())
    odds,pv = fisher_exact([[a,b],[c,d]], alternative="two-sided")
    se = np.sqrt(sum(1/v for v in [a,b,c,d])) if all([a,b,c,d]) else np.nan
    row = dict(analysis=label,n=len(df),permissive_n=a+b,permissive_engaged=a,
               nonpermissive_n=c+d,nonpermissive_engaged=c,odds_ratio=odds,
               wald_ci_low=np.exp(np.log(odds)-1.96*se),
               wald_ci_high=np.exp(np.log(odds)+1.96*se),fisher_two_sided_p=pv)
    X = pd.DataFrame({"constant":1., "permissive":x.astype(float),
                      "log_chains":np.log(df.n_chains.clip(lower=1))})
    model = sm.Logit(y.astype(float),X).fit(disp=False)
    ci=model.conf_int().loc["permissive"]
    row.update(adjusted_or=np.exp(model.params["permissive"]),
               adjusted_ci_low=np.exp(ci.iloc[0]),adjusted_ci_high=np.exp(ci.iloc[1]),
               adjusted_p=model.pvalues["permissive"])
    return row

def reselect(df):
    # Preserve the frozen selected row in maximum-burial ties whenever still eligible.
    # Otherwise use original observation-table order, so a filter does not silently
    # change the chain-count covariate merely by changing tie-breaking conventions.
    keys=set(zip(p.acc,p.pdb,p.asm_chain))
    df=df.copy()
    df["_original_selected"]=[(r.acc,r.pdb,r.asm_chain) in keys for r in df.itertuples()]
    df=df.sort_values("_original_selected",ascending=False,kind="stable")
    return df.loc[df.groupby("acc",sort=False).buried_pct.idxmax()].copy()

ao = audit[audit.analyzable].copy()
sens = [compare(keep,"baseline_frozen"),
        compare(keep[keep.is_first_observed],"selected_record_first_observed_only"),
        compare(reselect(ao[ao.is_first_observed]),"reselect_after_removing_nonfirst_observations"),
        compare(keep[~keep.gene.isin(["PSMA4","PSMA6","PSMA7","RAN"])],
                "exclude_three_PSMA_and_RAN"),
        compare(keep[~(keep.res2.isin(list("AGSTCPV"))&keep.resname.isin(["MET","MSE"]))],
                "exclude_permissive_modeled_MET_MSE"),
        compare(reselect(ao[ao.mapping_status=="exact_endpoint_offset"]),
                "reselect_exact_endpoint_offset_only"),
        compare(reselect(ao[ao.is_first_observed & (ao.mapping_status=="exact_endpoint_offset")]),
                "reselect_first_observed_and_exact_endpoint_offset"),
        compare(reselect(ao[~ao.sifts_short_span_flag & ~ao.sifts_low_span_flag]),
                "exploratory_remove_short_or_low_SIFTS_span"),
        compare(reselect(ao[ao.is_first_observed & ~ao.sifts_short_span_flag & ~ao.sifts_low_span_flag]),
                "exploratory_first_observed_and_not_short_low_span")]
pd.DataFrame(sens).to_csv(OUT / "association_sensitivities.csv", index=False)

selected = reselect(ao[ao.is_first_observed])
changes = keep[["acc","gene","buried_pct","pdb","asm_chain","is_first_observed"]].merge(
    selected[["acc","buried_pct","pdb","asm_chain"]],on="acc",how="left",
    suffixes=("_original","_boundary_filtered"))
changes["original_engaged"] = changes.buried_pct_original>=25
changes["new_engaged"] = changes.buried_pct_boundary_filtered>=25
changes["lost_all_observations"] = changes.buried_pct_boundary_filtered.isna()
changes.to_csv(OUT/"boundary_filter_candidate_changes.csv",index=False)
audit[audit.gene.isin(["PSMA4","PSMA6","PSMA7","RAN"])].to_csv(
    OUT/"focal_processing_records.csv",index=False)

# Boundary inspection of every matching census observation in the 11 frozen assemblies.
# These are ALREADY expanded assemblies. Do not transform to assembly again.
AA3 = {'ALA':'A','ARG':'R','ASN':'N','ASP':'D','CYS':'C','GLN':'Q','GLU':'E',
       'GLY':'G','HIS':'H','ILE':'I','LEU':'L','LYS':'K','MET':'M','PHE':'F',
       'PRO':'P','SER':'S','THR':'T','TRP':'W','TYR':'Y','VAL':'V','MSE':'M',
       'SEC':'U','PYL':'O'}
coord_rows=[]
for path in sorted((B/"inputs/figure5_assemblies").glob("*.cif.gz")):
    code=path.name.split(".")[0]
    st=gemmi.read_structure(str(path))
    st.remove_alternative_conformations()
    st.remove_hydrogens()
    chains={ch.name:ch for ch in st[0]}
    entities={sub:e for e in st.entities for sub in e.subchains}
    for r in ao[ao.pdb==code].itertuples():
        chain=chains.get(r.asm_chain,chains.get(r.chain))
        out={"acc":r.acc,"gene":r.gene,"pdb":code,"asm_chain":r.asm_chain,
             "original_chain":r.chain,"target_authnum":r.authnum,
             "is_first_observed_exported":r.is_first_observed}
        if chain is None:
            coord_rows.append({**out,"status":"chain_not_found"});continue
        residues=[z for z in chain if z.name in AA3 and z.seqid.icode in (' ','','\0')]
        matches=[(i,z) for i,z in enumerate(residues) if z.seqid.num==r.authnum]
        if not matches:
            coord_rows.append({**out,"status":"residue_not_found"});continue
        j,z=matches[0]
        entity=entities.get(z.subchain)
        n=next((a for a in z if a.name=="N"),None)
        prev=residues[j-1] if j else None
        carbon=next((a for a in prev if a.name=="C"),None) if prev else None
        distance=n.pos.dist(carbon.pos) if n is not None and carbon is not None else np.nan
        out.update(status="checked",resolved_chain=chain.name,resname=z.name,
                   modeled_aa_residues=len(residues),modeled_upstream_residues=j,
                   target_label_seq=z.label_seq,
                   entity_polymer_length=len(entity.full_sequence) if entity else None,
                   entity_prefix=" ".join(entity.full_sequence[:8]) if entity else "",
                   preceding_residue=f"{prev.name}{prev.seqid}" if prev else "",
                   preceding_C_to_target_N_A=distance,
                   peptide_bond_geometry=bool(1.0<=distance<=1.8),
                   modeled_prefix=" ".join(f"{v.name}{v.seqid}" for v in residues[:6]))
        coord_rows.append(out)
cd=pd.DataFrame(coord_rows)
cd.to_csv(OUT/"available_coordinate_boundary_audit.csv",index=False)

fig=pd.read_csv(B/"data/fig5_records.csv")
figrows=[]
for label,df in [("baseline_position_level",fig),
                 ("exclude_PSMA4_PSMA6_PSMA7_RAN",
                  fig[~fig.gene.isin(["PSMA4","PSMA6","PSMA7","RAN"])])]:
    sub=df[(df.burial>=50)&df.best_foreign_overlap.notna()]
    a=sub.loc[sub.kind=="control","best_foreign_overlap"]
    b=sub.loc[sub.kind=="test","best_foreign_overlap"]
    figrows.append(dict(analysis=label,control_n=len(a),test_n=len(b),
        control_median=a.median(),test_median=b.median(),
        two_sided_MW_p=mannwhitneyu(a,b,alternative="two-sided").pvalue))
pd.DataFrame(figrows).to_csv(OUT/"figure5_exploratory_sensitivity.csv",index=False)

summary={
 "protein_n":len(p),"analyzable_n":len(keep),"observation_n":len(o),
 "analyzable_observation_n":len(ao),
 "analyzable_first_observed_counts":ao.is_first_observed.value_counts().to_dict(),
 "analyzable_mapping_counts":ao.mapping_status.value_counts().to_dict(),
 "selected_analyzable_first_observed_counts":keep.is_first_observed.value_counts().to_dict(),
 "class_counts":keep.cls.value_counts().to_dict(),
 "retained_class_frozen_INIT_annotation_counts":
     pa[pa.cls.str.startswith("Met-retained")].frozen_init_annotation.fillna("").value_counts().to_dict(),
 "nonfirst_unique_proteins":int(ao.loc[~ao.is_first_observed,"acc"].nunique()),
 "all_observations_nonfirst_proteins":int(changes.lost_all_observations.sum()),
 "original_candidates":int(changes.original_engaged.sum()),
 "engaged_lost_all_observations":int((changes.original_engaged&changes.lost_all_observations).sum()),
 "engaged_lost_threshold_with_remaining_observations":
     int((changes.original_engaged&~changes.new_engaged&~changes.lost_all_observations).sum()),
 "original_permissive_candidates":int(((keep.buried_pct>=25)&keep.res2.isin(list("AGSTCPV"))).sum()),
 "boundary_filtered_permissive_candidates":int(((selected.buried_pct>=25)&selected.res2.isin(list("AGSTCPV"))).sum()),
 "sifts_short_observations":int(ao.sifts_short_span_flag.sum()),
 "sifts_low_span_observations":int(ao.sifts_low_span_flag.sum()),
 "coordinate_checked_observations":len(cd),
 "coordinate_checked_proteins":int(cd.acc.nunique()),
 "coordinate_preceding_peptide_bond_count":int(cd.peptide_bond_geometry.fillna(False).sum()),
 "software": {"python":sys.version,"pandas":pd.__version__,"gemmi":gemmi.__version__}
}
(OUT/"summary.json").write_text(json.dumps(summary,indent=2,default=str)+"\n")
hashes=[]
for path in [B/"data/asm_per_protein_master.csv",B/"data/asm_observations.csv",
             B/"inputs/human_nterm.tsv.gz",B/"inputs/pdb_chain_uniprot.tsv.gz",
             B/"data/fig5_records.csv",*sorted((B/"inputs/figure5_assemblies").glob("*.cif.gz"))]:
    hashes.append({"input":str(path.relative_to(B)),"sha256":hashlib.sha256(path.read_bytes()).hexdigest()})
pd.DataFrame(hashes).to_csv(OUT/"input_sha256.csv",index=False)
print(json.dumps(summary,indent=2,default=str))
print(pd.DataFrame(sens).to_string(index=False))
print(cd[cd.gene.isin(["PSMA4","PSMA6","PSMA7","RAN"])].to_string(index=False))
print(pd.DataFrame(figrows).to_string(index=False))
```
