## Supplementary_Data_8a for "Human protein complex interfaces reveal candidate assembly roles for initiator methionine excision": coordinate_followup.py

```
"""Audit Figure 5 residue selection and recalculate focal deposited-model geometry.
No relaxation, acetylation, or biological processing is inferred from static deletions.
"""
from pathlib import Path
import os, sys
import pandas as pd, numpy as np, gemmi
from scipy.spatial import cKDTree

R=Path(__file__).resolve().parent
B=R/"frozen/nterm-census-1.3.0"
O=R/"results"
os.environ["NME_WORK"]=str(R/"scratch")
sys.path.insert(0,str(B/"code"))
import asm_lib as lib
from scan_methionine import _resolve_chain
f=pd.read_csv(B/"data/fig5_targets.csv")
obs=pd.read_csv(B/"data/asm_observations.csv")
selection=[]
geometry=[]

def measure(T, ci, num, area, iso, removed=()):
    key=(ci,num,' ')
    # gemmi normally represents an empty insertion code as a space.
    keys=[k for k in area if k[0]==ci and k[1]==num]
    if not keys:return None
    key=keys[0]
    ai=iso[key]; ac=area[key]
    mask=np.array([k==key for k in T["rkey"]])
    other=T["cid"]!=ci
    tree=cKDTree(T["xyz"][other])
    distance=tree.query(T["xyz"][mask])[0].min()
    return {"buried_pct":100*(1-ac/ai) if ai>0 else np.nan,
            "sasa_isolated":ai,"sasa_complex":ac,"min_residue_contact_A":distance}

for pdb, targets in f.groupby("pdb",sort=True):
    st=gemmi.read_structure(str(B/f"inputs/figure5_assemblies/{pdb.lower()}.cif.gz"))
    for t in targets.itertuples():
        chname=_resolve_chain(st[0],t.chain)
        ch=next(c for c in st[0] if c.name==chname)
        aa=[z for z in ch if z.name in lib.AA3]
        met=None
        if t.kind=="control" and aa[0].name in ("MET","MSE") and len(aa)>1:
            met=aa[0]; target=aa[1]
        else:target=aa[0]
        selection.append(dict(gene=t.gene,pdb=pdb,manifest_chain=t.chain,resolved_chain=chname,
            kind=t.kind,expected_res2=t.res2,scan_selected_residue=target.name,
            scan_selected_authnum=target.seqid.num,scan_selected_label_seq=target.label_seq,
            expected_identity_matches=lib.AA3[target.name]==t.res2,
            excluded_met_residue=str(met.seqid) if met else "",
            first_modeled_residue=f"{aa[0].name}{aa[0].seqid}",
            modeled_prefix=" ".join(f"{z.name}{z.seqid}" for z in aa[:5])))
    focal=targets[targets.gene.isin(["PSMA4","PSMA6","PSMA7","RAN"])]
    if focal.empty:continue
    st.setup_entities();st.remove_alternative_conformations();st.remove_hydrogens()
    st.remove_ligands_and_waters()
    T=lib.chain_atom_table(st)
    areas=lib.residue_areas(lib.sasa(T["xyz"],T["r"]),T["rkey"])
    isos={}
    for t in focal.itertuples():
        chname=_resolve_chain(st[0],t.chain)
        ci=T["chains"].index(chname);ch=st[0][ci]
        if ci not in isos:
            m=T["cid"]==ci
            isos[ci]=lib.residue_areas(lib.sasa(T["xyz"][m],T["r"][m]),
                                     [k for k,v in zip(T["rkey"],m) if v])
        aa=[z for z in ch if z.name in lib.AA3]
        # Focal targets all have canonical numbering, verified against deposited prefixes.
        for num in ([0,1,2] if t.gene=="RAN" else [1,2]):
            z=next((z for z in aa if z.seqid.num==num),None)
            if z is None:continue
            out=dict(gene=t.gene,pdb=pdb,chain=chname,manifest_chain=t.chain,kind=t.kind,
                     measured_authnum=num,resname=z.name,
                     modeled_upstream_residues=aa.index(z),
                     scenario="unchanged_deposited_model")
            out.update(measure(T,ci,num,areas,isos[ci]))
            n=next((a for a in z if a.name=="N"),None)
            tree=cKDTree(T["xyz"][T["cid"]!=ci])
            out["target_N_contact_A"]=tree.query(n.pos.tolist())[0] if n else np.nan
            old=obs[(obs.pdb==pdb.lower())&(obs.chain==chname)&(obs.authnum==num)]
            out["original_census_buried_pct"]=old.iloc[0].buried_pct if len(old) else np.nan
            out["original_census_N_contact_A"]=old.iloc[0].min_dist_alphaamino_N if len(old) else np.nan
            geometry.append(out)
        # Only the four featured census maximum-burial cases get a deletion diagnostic.
        featured={( "PSMA6","8TM4","G1"),("PSMA4","7V5G","Q1"),
                  ("PSMA7","5LN3","D1"),("RAN","7MO1","A1")}
        if (t.gene,pdb,t.chain) in featured:
            removed=[0,1] if t.gene=="RAN" else [1]
            mask=np.array([not(k[0]==ci and k[1] in removed) for k in T["rkey"]])
            TT=dict(xyz=T["xyz"][mask],r=T["r"][mask],cid=T["cid"][mask],
                    rkey=[k for k,v in zip(T["rkey"],mask) if v],chains=T["chains"])
            ar=lib.residue_areas(lib.sasa(TT["xyz"],TT["r"]),TT["rkey"])
            m=TT["cid"]==ci
            iso=lib.residue_areas(lib.sasa(TT["xyz"][m],TT["r"][m]),
                                 [k for k,v in zip(TT["rkey"],m) if v])
            row=dict(gene=t.gene,pdb=pdb,chain=chname,manifest_chain=t.chain,kind=t.kind,
                     measured_authnum=2,resname="ALA" if t.gene=="RAN" else "SER",
                     modeled_upstream_residues=0,
                     scenario="static_upstream_deletion_NO_relaxation_NO_acetylation",
                     target_N_contact_A=geometry[-1]["target_N_contact_A"])
            row.update(measure(TT,ci,2,ar,iso))
            geometry.append(row)
    print("Completed",pdb,flush=True)

sel=pd.DataFrame(selection)
geo=pd.DataFrame(geometry)
sel.to_csv(O/"figure5_residue_selection_audit.csv",index=False)
geo.to_csv(O/"focal_recomputed_geometry.csv",index=False)
print("Scan identity mismatches:\n",sel[~sel.expected_identity_matches].to_string(index=False))
print("Geometry:\n",geo.to_string(index=False))
```
