## Supplementary_Data_8a for "Human protein complex interfaces reveal candidate assembly roles for initiator methionine excision": asm_test.py

```
import os as _os
BASE = _os.environ.get("NME_DIR") or _os.path.dirname(_os.path.dirname(_os.path.abspath(__file__)))
INPUT_DIR = _os.path.join(BASE, "inputs")
WORK_DIR = _os.environ.get("NME_WORK") or _os.path.join(BASE, "work")
_os.makedirs(WORK_DIR, exist_ok=True)
import gzip, csv, json, collections, sys, time
sys.path.insert(0, _os.path.dirname(_os.path.abspath(__file__)))
import asm_lib as L

TEST = ["1qo5", "3tj3", "5a9q", "8sg8", "7emf", "6pav", "4r3o", "1ibr"]
D = json.load(open(f"{WORK_DIR}/targets.json"))
prot = D["prot"]; targets = D["targets"]
sel = set(TEST)
by_entry = collections.defaultdict(list)
for t in targets:
    if t["pdb"] in sel:
        by_entry[t["pdb"]].append(t)
need = {t["acc"] for e in by_entry.values() for t in e}
seqs = {}
with gzip.open(f"{INPUT_DIR}/human_nterm.tsv.gz", "rt") as fh:
    for r in csv.DictReader(fh, delimiter="\t"):
        if r["Entry"] in need:
            seqs[r["Entry"]] = r["Sequence"]
segs = collections.defaultdict(list)
with gzip.open(f"{INPUT_DIR}/pdb_chain_uniprot.tsv.gz", "rt") as fh:
    fh.readline()
    for r in csv.DictReader(fh, delimiter="\t"):
        pdb = r["PDB"].lower()
        if pdb not in sel:
            continue
        try:
            segs[(pdb, r["CHAIN"], r["SP_PRIMARY"])].append((int(r["SP_BEG"]), r["PDB_BEG"]))
        except ValueError:
            pass

for pdb in TEST:
    t0 = time.time()
    rows, status = L.analyze_entry(pdb, by_entry.get(pdb, []), prot, seqs, segs, cleanup=False)
    print(f"== {pdb} {status} rows={len(rows)} {time.time()-t0:.1f}s "
          f"nchains={rows[0]['n_chains'] if rows else '-'} atoms={rows[0]['n_atoms'] if rows else '-'}")
    agg = collections.defaultdict(list)
    for r in rows:
        agg[r["gene"]].append((r["asm_chain"], r["buried_pct"], r["min_dist_alphaamino_N"]))
    for g, v in sorted(agg.items()):
        print("   ", g, v[:8], "n=", len(v))
```
