## Supplementary_Data_8a for "Human protein complex interfaces reveal candidate assembly roles for initiator methionine excision": assemble_figure5_records.py

```
"""Combine the frozen Figure 5 target metadata with raw conformer-scan scores."""
from __future__ import annotations

import argparse
import csv


def key(row: dict[str, str]) -> tuple[str, ...]:
    return tuple(row[name] for name in ("gene", "pdb", "chain", "kind", "res2"))


def main() -> None:
    parser = argparse.ArgumentParser()
    parser.add_argument("--targets", required=True)
    parser.add_argument("--scores", required=True)
    parser.add_argument("--out", required=True)
    args = parser.parse_args()

    targets = list(csv.DictReader(open(args.targets)))
    scores = list(csv.DictReader(open(args.scores)))
    score_map = {key(row): row for row in scores}
    if len(score_map) != len(scores):
        raise ValueError("duplicate score keys")

    fields = ["gene", "pdb", "chain", "kind", "res2", "burial", "burial_res2",
              "burial_met1_observed", "observed_met1_overlap", "best_foreign_overlap",
              "n_self_ok", "n_both_ok", "n_total", "n_chains"]
    output = []
    for target in targets:
        score = score_map[key(target)]
        row = dict(target)
        for name in ("best_foreign_overlap", "n_self_ok", "n_both_ok", "n_total"):
            row[name] = score[name]
        output.append({name: row.get(name, "") for name in fields})

    with open(args.out, "w", newline="") as handle:
        writer = csv.DictWriter(handle, fieldnames=fields)
        writer.writeheader()
        writer.writerows(output)


if __name__ == "__main__":
    main()
```
