## Supplementary_Data_8a for "Human protein complex interfaces reveal candidate assembly roles for initiator methionine excision": fig_asm.py

```
import os as _os
BASE = _os.environ.get("NME_DIR") or _os.path.dirname(_os.path.dirname(_os.path.abspath(__file__)))
INPUT_DIR = _os.path.join(BASE, "inputs")
WORK_DIR = _os.environ.get("NME_WORK") or _os.path.join(BASE, "work")
_os.makedirs(WORK_DIR, exist_ok=True)
"""Figures for the biological-assembly, full-coverage census (paper Figs 1-4)."""
import csv, json, collections, math, re
import numpy as np
import matplotlib
matplotlib.use("Agg")
import matplotlib.pyplot as plt
from matplotlib.patches import Patch

OUT = f"{WORK_DIR}/"
SMALL = set("AGSTCPV")
PHI = set("LFYWIVA")
C_NME = "#3b6ea5"
C_MET = "#c9752b"

prot = list(csv.DictReader(open(OUT + "asm_per_protein_master.csv")))
for p in prot:
    p["b"] = float(p["buried_pct"])
    p["dN"] = float(p["dN_min"]) if p["dN_min"] not in ("", "inf") else np.nan
    p["nch"] = int(p["n_chains"])
keep = [p for p in prot if not p["cls"].startswith("excluded")]
A = [p for p in keep if p["pred"] == "predicted NME"]
B = [p for p in keep if p["pred"] == "predicted Met-retained"]

# --------------------------------------------------------------- FIGURE 1
fig = plt.figure(figsize=(11.6, 8.2))
gs = fig.add_gridspec(2, 2, height_ratios=[1, 1], hspace=0.42, wspace=0.26)
ax0 = fig.add_subplot(gs[0, 0])
ax1 = fig.add_subplot(gs[0, 1])
ax2 = fig.add_subplot(gs[1, 0])
ax3 = fig.add_subplot(gs[1, 1])

# A: cumulative distributions
for g, c, lab in ((A, C_NME, "MetAP-permissive"), (B, C_MET, "MetAP-nonpermissive")):
    v = np.sort(np.array([p["b"] for p in g]))
    ax0.step(v, np.arange(1, len(v) + 1) / len(v), color=c, lw=2,
             label=f"{lab} (n = {len(g)})")
ax0.set_xlabel("Burial of the mature N-terminal residue on assembly (%)", fontsize=9)
ax0.set_ylabel("Cumulative fraction of proteins", fontsize=9)
ax0.axvline(25, color="#888888", lw=0.8, ls="--")
ax0.text(26.5, 0.08, "25% burial", fontsize=7.4, color="#666666")
ax0.legend(fontsize=8.2, frameon=False, loc="lower right")
ax0.set_title("A   Excision-dependent termini are less buried", loc="left",
              fontsize=10, fontweight="bold")
ax0.set_xlim(-2, 102); ax0.set_ylim(0, 1.02)

# B: swarm-style by P1' residue class
order = [("A", "Ala"), ("S", "Ser"), ("G", "Gly"), ("T", "Thr"), ("P", "Pro"), ("V", "Val"),
         ("E", "Glu"), ("K", "Lys"), ("D", "Asp"), ("L", "Leu"), ("F", "Phe"),
         ("R", "Arg"), ("I", "Ile"), ("M", "Met")]
xs, meds, ns, cols, labs = [], [], [], [], []
for i, (aa, nm) in enumerate(order):
    g = [p["b"] for p in keep if p["res2"] == aa]
    if len(g) < 8:
        continue
    xs.append(len(xs))
    meds.append(float(np.median(g)))
    ns.append(len(g))
    cols.append(C_NME if aa in SMALL else C_MET)
    labs.append(nm)
    v = np.array(g)
    ax1.scatter(np.random.normal(xs[-1], 0.13, len(v)), v, s=5, alpha=0.30,
                color=cols[-1], edgecolors="none")
ax1.scatter(xs, meds, marker="_", s=380, color="black", zorder=4, linewidths=1.8)
ax1.set_xticks(xs)
# Letters and counts are drawn separately so the counts can be set smaller than the
# residue letters; as a single two-line tick label they shared one font size and the
# counts row collided. Same change as in rebuild_figures_1_to_4.py.
ax1.set_xticklabels(labs, fontsize=7.2)
for _x, _n in zip(xs, ns):
    ax1.annotate(str(_n), xy=(_x, 0), xycoords=("data", "axes fraction"),
                 xytext=(0, -15), textcoords="offset points",
                 ha="center", va="top", fontsize=5.2, annotation_clip=False)
ax1.set_ylabel("Burial on assembly (%)", fontsize=9)
ax1.set_xlabel("Penultimate (P1') residue, with n below", fontsize=9, labelpad=11)
ax1.set_title("B   Burial tracks P1' identity, not excision", loc="left",
              fontsize=10, fontweight="bold")
ax1.legend(handles=[Patch(facecolor=C_NME, label="MetAP-permissive P1'"),
                    Patch(facecolor=C_MET, label="MetAP-nonpermissive P1'")],
           fontsize=7.8, frameon=False, loc="upper left")
ax1.set_ylim(-4, 108)

# C: odds ratio by assembly size (>=25% burial, NME vs refractory)
def orci(a, b):
    ka, na = sum(1 for x in a if x["b"] >= 25), len(a)
    kb, nb = sum(1 for x in b if x["b"] >= 25), len(b)
    t = np.array([[ka + 0.5, na - ka + 0.5], [kb + 0.5, nb - kb + 0.5]])
    lor = math.log((t[0, 0] * t[1, 1]) / (t[0, 1] * t[1, 0]))
    se = math.sqrt(sum(1 / t.ravel()))
    return math.exp(lor), math.exp(lor - 1.96 * se), math.exp(lor + 1.96 * se), na, nb

bands = [(2, 3, "2–3 chains"), (4, 5, "4–5 chains"), (6, 11, "6–11 chains"),
         (12, 10 ** 6, "\u226512 chains"), (2, 10 ** 6, "all assemblies")]
rows = []
for lo, hi, lab in bands:
    a = [p for p in A if lo <= p["nch"] <= hi]
    b = [p for p in B if lo <= p["nch"] <= hi]
    rows.append((lab,) + orci(a, b))
y = np.arange(len(rows))[::-1]
for yy, (lab, o, l, u, na, nb) in zip(y, rows):
    c = "#222222" if lab == "all assemblies" else C_NME
    ax2.plot([l, u], [yy, yy], color=c, lw=1.6)
    ax2.plot([o], [yy], "o", color=c, ms=7)
    ax2.text(2.45, yy, f"{o:.2f}  [{l:.2f}–{u:.2f}]", fontsize=7.6, va="center", color="#333333")
ax2.axvline(1, color="#888888", lw=0.9, ls="--")
ax2.set_yticks(y)
ax2.set_yticklabels([f"{r[0]}\nn = {r[4]} vs {r[5]}" for r in rows], fontsize=7.8)
ax2.set_xscale("log")
ax2.set_xlim(0.28, 5.6)
ax2.set_xticks([0.3, 0.5, 1, 2])
ax2.set_xticklabels(["0.3", "0.5", "1", "2"], fontsize=8)
from matplotlib.ticker import NullFormatter, NullLocator
ax2.xaxis.set_minor_formatter(NullFormatter())
ax2.xaxis.set_minor_locator(NullLocator())
ax2.set_xlabel("Odds ratio for burial $\\geq$25%, MetAP-permissive vs nonpermissive", fontsize=8.6)
ax2.set_title("C   Depletion holds across assembly sizes", loc="left",
              fontsize=10, fontweight="bold")
ax2.tick_params(axis="y", length=0)
ax2.text(0.0, -0.34, "Logistic model over all 1,143 proteins: OR 0.65 (0.51\u20130.84) for excision-dependence, adjusted for\n"
         "log chain count; burial rises with assembly size (OR 1.37 per e-fold, $p$ = 4 \u00d7 10$^{-10}$); "
         "interaction $p$ = 0.12.",
         transform=ax2.transAxes, fontsize=7.0, color="#555555", va="top", linespacing=1.5)

# D: burial vs distance, at-risk quadrant
gg = [p for p in keep if np.isfinite(p["dN"])]
for g, c, lab in ((A, C_NME, "MetAP-permissive"), (B, C_MET, "MetAP-nonpermissive")):
    g2 = [p for p in g if np.isfinite(p["dN"])]
    ax3.scatter(np.clip([p["dN"] for p in g2], 0, 20), [p["b"] for p in g2],
                s=7, alpha=0.42, color=c, edgecolors="none", label=f"{lab} (n = {len(g2)})")
risk = [p for p in A if p["b"] >= 25]
ax3.axhline(25, color="#a33227", lw=0.9, ls="--")
ax3.axvline(5, color="#888888", lw=0.9, ls="--")
ax3.add_patch(plt.Rectangle((0, 25), 5, 80, facecolor="#a33227", alpha=0.06, zorder=0))
ax3.text(0.4, 106, f"high-engagement quadrant: {len(risk)} MetAP-permissive termini",
         fontsize=7.8, color="#a33227")
ax3.set_xlabel("Distance, \u03b1-amino nitrogen to nearest atom of another subunit "
               "(\u00c5, clipped at 20)", fontsize=8.6)
ax3.set_ylabel("Burial on assembly (%)", fontsize=9)
ax3.set_title("D   Geometry of the at-risk set", loc="left", fontsize=10, fontweight="bold")
ax3.legend(fontsize=7.8, frameon=False, loc="lower right")
ax3.set_ylim(-4, 112); ax3.set_xlim(0, 20.5)
for ax in (ax0, ax1, ax2, ax3):
    for s in ("top", "right"):
        ax.spines[s].set_visible(False)
    ax.grid(color="#e8e8e8", lw=0.6)
    ax.set_axisbelow(True)
fig.savefig(OUT + "asm_fig1_census.png", dpi=300, bbox_inches="tight", facecolor="white")
plt.close(fig)
print("wrote asm_fig1_census.png")

# --------------------------------------------------------------- FIGURE 2  alpha ring
obs = list(csv.DictReader(open(OUT + "asm_observations.csv")))


def pick(pdb, gene):
    v = [o for o in obs if o["pdb"] == pdb and o["gene"].split(";")[0] == gene]
    if not v:
        return None
    return max(v, key=lambda o: float(o["buried_pct"]))


def best(gene):
    v = [o for o in obs if o["gene"].split(";")[0] == gene]
    return max(v, key=lambda o: float(o["buried_pct"]))["pdb"] if v else None


ring = [("PSMA6 (\u03b11)", best("PSMA6"), "PSMA6", "retained"),
        ("PSMA2 (\u03b12)", best("PSMA2"), "PSMA2", "excised"),
        ("PSMA4 (\u03b13)", best("PSMA4"), "PSMA4", "retained"),
        ("PSMA7 (\u03b14)", best("PSMA7"), "PSMA7", "retained"),
        ("PSMA5 (\u03b15)", best("PSMA5"), "PSMA5", "refractory"),
        ("PSMA1 (\u03b16)", best("PSMA1"), "PSMA1", "refractory"),
        ("PSMA3 (\u03b17)", best("PSMA3"), "PSMA3", "excised"),
        ("PSMA3 (\u03b17)", "4r3o", "PSMA3", "excised"),
        ("PSMB3 (\u03b23)", best("PSMB3"), "PSMB3", "excised"),
        ("PSMB5 (\u03b25)", "8yix", "PSMB5", "propeptide"),
        ("PSMB7 (\u03b22)", "8yix", "PSMB7", "propeptide")]
CMAP = {"excised": C_NME, "refractory": C_MET, "propeptide": "#6b6b6b", "retained": "#8fb3d9"}
recs = []
for lab, pdb, gene, kind in ring:
    o = pick(pdb, gene)
    if o:
        recs.append((lab, float(o["buried_pct"]), float(o["min_dist_alphaamino_N"]),
                     kind, pdb, o["resname"] + str(o["authnum"]), o["res2"]))
recs.sort(key=lambda r: -r[1])
fig, (a1, a2) = plt.subplots(1, 2, figsize=(11.4, 5.6), gridspec_kw=dict(wspace=0.06))
y = np.arange(len(recs))[::-1]
cols = [CMAP[r[3]] for r in recs]
a1.barh(y, [r[1] for r in recs], color=cols, height=0.62, edgecolor="white")
for yy, r in zip(y, recs):
    a1.text(r[1] + 1.6, yy, f"{r[1]:.1f}%", va="center", fontsize=8.2, color="#333333")
a1.set_yticks(y)
a1.set_yticklabels([f"{r[0]}   {r[5]}, P1'={r[6]}   {r[4]}" for r in recs], fontsize=8)
a1.set_xlim(0, 112)
a1.axvline(25, color="#a33227", ls="--", lw=1)
a1.set_xlabel("Burial of residue 1 in the assembly (%)", fontsize=8.8)
a1.set_title("A   How deeply each terminus sits in the 20S particle", loc="left",
             fontsize=9.6, fontweight="bold")
a2.scatter([r[2] for r in recs], y, color=cols, s=58, zorder=3)
for yy, r in zip(y, recs):
    a2.text(r[2] + 0.35, yy, f"{r[2]:.2f} \u00c5", va="center", fontsize=8.2, color="#333333")
a2.axvline(5, color="#444444", ls="--", lw=1)
a2.set_yticks(y); a2.set_yticklabels([])
a2.set_xlim(0, 14)
a2.set_xlabel("\u03b1-amino nitrogen to nearest atom of another subunit (\u00c5)", fontsize=8.8)
a2.set_title("B   Contact geometry of the free \u03b1-amino group", loc="left",
             fontsize=9.6, fontweight="bold")
for ax in (a1, a2):
    for s in ("top", "right"):
        ax.spines[s].set_visible(False)
    ax.tick_params(axis="y", length=0)
    ax.grid(axis="x", color="#e4e4e4", lw=0.7)
    ax.set_axisbelow(True)
a1.legend(handles=[Patch(facecolor=C_NME, label="methionine excised (annotated)"),
                   Patch(facecolor="#8fb3d9", label="excision-competent P1', Met modelled"),
                   Patch(facecolor=C_MET, label="MetAP-nonpermissive P1'"),
                   Patch(facecolor="#6b6b6b", label="propeptide-bearing pro-form (excluded from statistics)")],
          fontsize=7.4, frameon=False, loc="upper center", bbox_to_anchor=(0.62, -0.115),
          handlelength=1.2, ncol=2)
fig.savefig(OUT + "asm_fig2_alpha_ring.png", dpi=300, bbox_inches="tight", facecolor="white")
plt.close(fig)
print("wrote asm_fig2_alpha_ring.png")

# --------------------------------------------------------------- FIGURE 3  transport
tr = [("KPNB1  importin-\u03b2", "6n88", "KPNB1", "refractory"),
      ("RAN  GTPase", "7mo1", "RAN", "excised"),
      ("KPNA2  importin-\u03b1", "9bfc", "KPNA2", "excised"),
      ("NUP188  pore scaffold", "5ijo", "NUP188", "excised"),
      ("NUP93  pore scaffold", "5ijo", "NUP93", "refractory"),
      ("SEH1L  pore scaffold", "5a9q", "SEH1L", "refractory"),
      ("NUP50  pore basket", "3tj3", "NUP50", "excised")]
recs = []
for lab, pdb, gene, kind in tr:
    o = pick(pdb, gene)
    if o:
        recs.append((lab, float(o["buried_pct"]), float(o["min_dist_alphaamino_N"]),
                     kind, pdb, int(o["n_chains"])))
recs.sort(key=lambda r: -r[1])
fig, (a1, a2) = plt.subplots(1, 2, figsize=(11.4, 4.3), gridspec_kw=dict(wspace=0.06))
y = np.arange(len(recs))[::-1]
cols = [CMAP[r[3]] for r in recs]
a1.barh(y, [r[1] for r in recs], color=cols, height=0.62, edgecolor="white")
for yy, r in zip(y, recs):
    a1.text(r[1] + 1.6, yy, f"{r[1]:.1f}%", va="center", fontsize=8.2, color="#333333")
a1.set_yticks(y)
a1.set_yticklabels([f"{r[0]}   {r[4]}, {r[5]} chains" for r in recs], fontsize=8)
a1.set_xlim(0, 112)
a1.axvline(25, color="#a33227", ls="--", lw=1)
a1.set_xlabel("Burial of the mature N-terminal residue (%)", fontsize=8.8)
a1.set_title("A   Pore scaffold versus soluble carriers", loc="left",
             fontsize=9.6, fontweight="bold")
a2.scatter([min(r[2], 49) for r in recs], y, color=cols, s=58, zorder=3)
for yy, r in zip(y, recs):
    a2.text(min(r[2], 49) + 1.1, yy, f"{r[2]:.2f} \u00c5", va="center", fontsize=8.2, color="#333333")
a2.axvline(5, color="#444444", ls="--", lw=1)
a2.set_yticks(y); a2.set_yticklabels([])
a2.set_xlim(0, 60)
a2.set_xlabel("\u03b1-amino nitrogen to nearest atom of another subunit (\u00c5)", fontsize=8.8)
a2.set_title("B   Contact geometry of the \u03b1-amino group", loc="left",
             fontsize=9.6, fontweight="bold")
for ax in (a1, a2):
    for s in ("top", "right"):
        ax.spines[s].set_visible(False)
    ax.tick_params(axis="y", length=0)
    ax.grid(axis="x", color="#e4e4e4", lw=0.7)
    ax.set_axisbelow(True)
a1.legend(handles=[Patch(facecolor=C_NME, label="methionine excised (annotated)"),
                   Patch(facecolor=C_MET, label="MetAP-nonpermissive P1'")],
          fontsize=7.6, frameon=False, loc="lower right", handlelength=1.2)
fig.savefig(OUT + "asm_fig3_transport.png", dpi=300, bbox_inches="tight", facecolor="white")
plt.close(fig)
print("wrote asm_fig3_transport.png")

# --------------------------------------------------------------- FIGURE 4  machines
def machine(g):
    g = g.split(";")[0]
    if re.match(r"PSM[ABCDEFG]", g): return "26S proteasome / assembly"
    if re.match(r"CCT\d|TCP1", g): return "TRiC/CCT chaperonin"
    if re.match(r"MED\d", g): return "Mediator"
    if re.match(r"SF3[AB]|SNRP|PRPF|LSM|EFTUD|SNU|SART|BUD31|CDC5L", g): return "Spliceosome"
    if re.match(r"COPS|CUL\d|DDB1|DDA1|RBX1|SKP1|ELOB|ELOC|FBXW", g): return "Cullin\u2013RING / CSN"
    if re.match(r"NDUF|UQCR|COX\d|SDH[ABCD]|ATP5|MT-", g): return "OXPHOS complexes"
    if re.match(r"RPL\d|RPS\d|RPLP|FAU|RPSA|MRPL|MRPS", g): return "Ribosomes (cyto + mito)"
    if re.match(r"NUP\d|SEH1L|SEC13|AAAS|RAE1|NDC1|POM121|TPR$", g): return "Nuclear pore scaffold"
    if re.match(r"POLR[123]", g): return "RNA polymerases"
    if re.match(r"EIF3|EIF2", g): return "eIF2 / eIF3"
    if re.match(r"ACT[BGA]|TUB[ABG]|ARPC|ACTR", g): return "Actin / tubulin systems"
    if re.match(r"ANAPC|CDC2|CDK\d|CCN[A-Z]|MCM\d|GINS", g): return "Cell cycle / replication"
    if re.match(r"KPN[AB]|TNPO|RAN$|RANBP|IPO\d|XPO\d", g): return "Soluble transport receptors"
    return None


by = collections.defaultdict(list)
for p in keep:
    m = machine(p["gene"])
    if m:
        by[m].append(p)
recs = []
for m, rs in by.items():
    if len(rs) < 4:
        continue
    nme = [r for r in rs if r["pred"] == "predicted NME"]
    risk = [r for r in nme if r["b"] >= 25]
    top = max(rs, key=lambda r: r["b"])
    deg = [r for r in nme if r["res2"] in PHI]
    recs.append((m, len(rs), 100 * len(nme) / len(rs),
                 100 * len(risk) / len(nme) if nme else np.nan,
                 top["b"], 100 * len(deg) / len(nme) if nme else np.nan,
                 top["gene"].split(";")[0]))
recs.sort(key=lambda t: -(t[3] if t[3] == t[3] else -1))
cols = ["Excision-dependent\ntermini (%)", "Of those, buried\n$\\geq$25% (%)",
        "Most buried\nterminus (%)", "Degron-forming\nfraction (%)"]
M = np.array([[r[2], r[3], r[4], r[5]] for r in recs], dtype=float)
fig, ax = plt.subplots(figsize=(9.8, 5.6))
im = ax.imshow(M, cmap="YlOrRd", vmin=0, vmax=100, aspect="auto")
ax.set_xticks(range(len(cols))); ax.set_xticklabels(cols, fontsize=8.2)
ax.set_yticks(range(len(recs)))
ax.set_yticklabels([f"{r[0]}  (n = {r[1]})" for r in recs], fontsize=8.5)
for i in range(M.shape[0]):
    for j in range(M.shape[1]):
        v = M[i, j]
        if v != v:
            ax.text(j, i, "\u2013", ha="center", va="center", fontsize=8.4, color="#666666")
        else:
            ax.text(j, i, f"{v:.0f}", ha="center", va="center", fontsize=8.6,
                    color="white" if v > 62 else "#222222",
                    fontweight="bold" if v > 62 else "normal")
for i, r in enumerate(recs):
    ax.text(M.shape[1] - 0.34, i, f"  {r[6]}", ha="left", va="center", fontsize=7.8, color="#444444")
ax.set_xlim(-0.5, M.shape[1] + 1.5)
ax.text(M.shape[1] - 0.34, -0.85, "  Most buried\n  subunit", ha="left", va="center",
        fontsize=8.2, color="#333333")
ax.set_title("Exposure of human macromolecular machines to a failure of\n"
             "N-terminal methionine excision", loc="left", fontsize=10.2,
             fontweight="bold", pad=8)
ax.tick_params(length=0)
for s in ax.spines.values():
    s.set_visible(False)
cb = fig.colorbar(im, ax=ax, fraction=0.028, pad=0.06)
cb.set_label("percent", fontsize=8.2); cb.ax.tick_params(labelsize=7.6)
fig.text(0.005, -0.02,
         "Each row aggregates subunits whose mature N-terminal residue is resolved in at least one multi-chain "
         "biological assembly, so n is smaller than the\ncomplex. Rightmost labels name the most buried terminus in "
         "the row. Degron-forming means P1$'$ $\\in$ {L, F, Y, W, I, V, A}, which yields a Met-$\\Phi$/N-degron\n"
         "if the methionine is retained and unacetylated.",
         fontsize=7.4, color="#555555", ha="left", va="top", linespacing=1.5)
fig.savefig(OUT + "asm_fig4_machines.png", dpi=300, bbox_inches="tight", facecolor="white")
plt.close(fig)
print("wrote asm_fig4_machines.png")
json.dump([{k: (v if v == v else None) for k, v in
            zip(["machine", "n", "pct_nme", "pct_risk_of_nme", "max_buried", "pct_degron", "top_gene"], r)}
           for r in recs], open(OUT + "asm_machine_table.json", "w"), indent=1)
print("wrote asm_machine_table.json")
```
