## Supplementary_Data_8a for "Human protein complex interfaces reveal candidate assembly roles for initiator methionine excision": fig_cascade.py

```
import os as _os
BASE = _os.environ.get("NME_DIR") or _os.path.dirname(_os.path.dirname(_os.path.abspath(__file__)))
INPUT_DIR = _os.path.join(BASE, "inputs")
WORK_DIR = _os.environ.get("NME_WORK") or _os.path.join(BASE, "work")
_os.makedirs(WORK_DIR, exist_ok=True)
"""Figure: downstream Nalpha-modification branches vs interface geometry."""
import json
import numpy as np
import matplotlib
matplotlib.use("Agg")
import matplotlib.pyplot as plt
from matplotlib.patches import Patch

OUT = f"{WORK_DIR}/"
C_NME = "#3b6ea5"
C_MYR = "#7a4b9c"
C_MET = "#1f8a70"
C_GREY = "#9a9a9a"

recs = json.load(open(OUT + "asm_cascade_records.json"))
nme = [r for r in recs if r["pred_nme"]]
myr = [r for r in nme if r["myr"]]
mot = [r for r in nme if r["ntmt_motif"] and not r["myr"]]
ac = [r for r in nme if r["acetyl"] and not r["myr"] and not r["ntmt_motif"]]
oth = [r for r in nme if not r["myr"] and not r["ntmt_motif"] and not r["acetyl"]]

groups = [("myristoylated\nGly$_1$", myr, C_MYR),
          ("NTMT motif\n[A/P/S/G]-P-K", mot, C_MET),
          ("Nt-acetylated\n(annotated)", ac, C_NME),
          ("no downstream\nannotation", oth, C_GREY)]

fig, (a1, a2) = plt.subplots(1, 2, figsize=(12.6, 5.9),
                             gridspec_kw=dict(width_ratios=[1.0, 1.18], wspace=0.28))

# ---- panel A: burial by branch
rng = np.random.default_rng(7)
for i, (lab, g, c) in enumerate(groups):
    b = np.array([r["buried"] for r in g])
    x = i + rng.uniform(-0.20, 0.20, len(b))
    a1.scatter(x, b, s=26 if len(b) < 40 else 8, alpha=0.75 if len(b) < 40 else 0.30,
               color=c, edgecolors="none", zorder=3)
    a1.plot([i - 0.34, i + 0.34], [np.median(b)] * 2, color="black", lw=2.2, zorder=4)
    a1.text(i, 106, f"n = {len(b)}", ha="center", fontsize=8.4, color="#333333")
    a1.text(i, -11.5, f"med. {np.median(b):.1f}%", ha="center", fontsize=8.0, color=c)
a1.axhline(25, color="#a33227", lw=0.9, ls="--")
a1.text(3.42, 27, "25%", fontsize=7.6, color="#a33227", ha="right")
a1.set_xticks(range(len(groups)))
a1.set_xticklabels([g[0] for g in groups], fontsize=8.6)
a1.set_ylim(-16, 112)
a1.set_xlim(-0.6, len(groups) - 0.4)
a1.set_ylabel("burial of residue 1 at inter-subunit interfaces (%)", fontsize=9.4)
a1.set_title("A   Interface engagement by downstream branch", fontsize=10.4,
             loc="left", pad=14)

# ---- panel B: individual cases
cases = sorted(myr, key=lambda r: r["buried"]) + [None] + sorted(mot, key=lambda r: r["buried"])
labels, vals, cols, dns = [], [], [], []
for r in cases:
    if r is None:
        labels.append("")
        vals.append(0.0)
        cols.append("white")
        dns.append(None)
        continue
    tag = "G" if r["myr"] else r["m1"] + r["m2"] + r["m3"]
    labels.append(f"{r['gene']}  ({tag}$_1$)" if r["myr"] else f"{r['gene']}  ({tag})")
    vals.append(r["buried"])
    cols.append(C_MYR if r["myr"] else C_MET)
    dns.append(r["dN"])
y = np.arange(len(vals))
a2.barh(y, vals, color=cols, height=0.66, edgecolor="white")
for yy, v, d in zip(y, vals, dns):
    if d is None:
        continue
    txt = f"{v:.1f}%" + (f"   {d:.2f} Å" if d is not None and d < 30 else "   no contact")
    a2.text(v + 1.6, yy, txt, va="center", fontsize=7.6, color="#333333")
a2.set_yticks(y)
a2.set_yticklabels(labels, fontsize=8.0)
a2.set_xlim(0, 128)
a2.set_xlabel("burial of residue 1 (%)   ·   right-hand value: min. distance from\n"
              "the free $\\alpha$-amino nitrogen to a neighbouring subunit", fontsize=8.6)
a2.set_title("B   Termini committed to a downstream modification", fontsize=10.4,
             loc="left", pad=14)
a2.axvline(25, color="#a33227", lw=0.9, ls="--")
a2.legend(handles=[Patch(facecolor=C_MYR, label="N-myristoylated (annotated)"),
                   Patch(facecolor=C_MET, label="NTMT recognition motif")],
          fontsize=8.2, frameon=False, loc="lower right", bbox_to_anchor=(1.0, -0.005))

for ax in (a1, a2):
    ax.set_axisbelow(True)
    ax.grid(color="#e8e8e8", lw=0.6)
    for s in ("top", "right"):
        ax.spines[s].set_visible(False)

fig.savefig(OUT + "asm_fig5_cascade.png", dpi=300, bbox_inches="tight", facecolor="white")
print("wrote asm_fig5_cascade.png")
```
