## Supplementary_Data_8a for "Human protein complex interfaces reveal candidate assembly roles for initiator methionine excision": rebuild_figure5.py

```
"""Rebuild manuscript Figure 5 from the recomputed methionine conformer scan.

Reads one deposited table, data/fig5_records.csv, which lists the 53 termini scanned
in the exhaustive rotamer sweep. Each row carries the burial of
residue 1 in the assembly, the best-case steric overlap of a methionine modelled at
position 1 (best_foreign_overlap, in angstroms), and, for the 28 control termini at
which methionine 1 is itself resolved, the same statistic recomputed on the deposited
methionine (observed_met1_overlap). The table is generated by scan_methionine.py
from the 11 frozen RCSB biological-assembly mmCIF files and target manifest.

The primary deeply buried comparison contains 17 controls and 10 tests, with
medians -1.12 and -0.30 A and a two-sided Mann-Whitney p value of 0.00490.

Authored at 180 mm printed width (7.09 in), so point sizes in this file are point
sizes on the page. Do not widen the figsize without scaling the fonts.

Usage:  NME_WORK=<dir with fig5_records.csv> python rebuild_figure5.py
Output: fig5_conformer_scan.png
"""
import os as _os
BASE = _os.environ.get("NME_DIR") or _os.path.dirname(_os.path.dirname(_os.path.abspath(__file__)))
WORK_DIR = _os.environ.get("NME_WORK") or _os.path.join(BASE, "work")
OUTDIR = _os.environ.get("NME_FIGOUT") or WORK_DIR
_os.makedirs(OUTDIR, exist_ok=True)

import csv
import numpy as np
import matplotlib
matplotlib.use("Agg")
import matplotlib.pyplot as plt
from matplotlib.patches import Patch
from scipy.stats import mannwhitneyu

IN = WORK_DIR + "/"
OUT = OUTDIR + "/"

C_CTRL = "#3b6ea5"       # blue: Met1 observed
C_TEST = "#c9752b"       # orange: Met1 absent
RED = "#a33227"

W = 7.09
ANN = 6.6
PANEL = 8.6
AX = 7.4
TICK = 6.6
plt.rcParams["font.size"] = TICK
plt.rcParams["xtick.labelsize"] = TICK
plt.rcParams["ytick.labelsize"] = TICK

# ----------------------------------------------------------------- input table
rows = list(csv.DictReader(open(IN + "fig5_records.csv")))
for r in rows:
    r["burial"] = float(r["burial"])
    r["best_foreign_overlap"] = float(r["best_foreign_overlap"]) if r["best_foreign_overlap"] else np.nan
    r["observed_met1_overlap"] = float(r["observed_met1_overlap"]) if r["observed_met1_overlap"] else np.nan
ctrl = [r for r in rows if r["kind"] == "control"]
test = [r for r in rows if r["kind"] == "test"]

# ---------------------------------------------------------------------- panel a
fig = plt.figure(figsize=(W, 4.6), layout="constrained")
fig.get_layout_engine().set(hspace=0.20, wspace=0.14)
gs = fig.add_gridspec(2, 2, height_ratios=[1.30, 1])
a0 = fig.add_subplot(gs[0, :])
a1 = fig.add_subplot(gs[1, 0])
a2 = fig.add_subplot(gs[1, 1])

a0.axhline(0, color="#888888", lw=0.6, ls=":")

for grp, color, marker, label in ((ctrl, C_CTRL, "o", "Met1 observed (26 scoreable of 28)"),
                                  (test, C_TEST, "^", "Met1 absent; modeled in silico (23 scoreable of 25)")):
    xs = [r["burial"] for r in grp if np.isfinite(r["best_foreign_overlap"])]
    ys = [r["best_foreign_overlap"] for r in grp if np.isfinite(r["best_foreign_overlap"])]
    a0.scatter(xs, ys, s=26, color=color, marker=marker, alpha=0.85,
               edgecolor="white", linewidth=0.4, label=label, zorder=3)

a0.set_xlim(-5, 105)
a0.set_ylim(-10.5, 3.2)
a0.set_xlabel("Burial of the N-terminal residue in the assembly (%)", fontsize=AX)
a0.set_ylabel("Best-case steric overlap of a\nmethionine at position 1 (\u00c5)", fontsize=AX)
a0.set_title("a   Descriptive conformer scan across 53 coordinate positions",
             fontsize=PANEL, loc="left")
a0.legend(fontsize=ANN, frameon=False, loc="lower right", handletextpad=0.6)
a0.spines[["top", "right"]].set_visible(False)

# ---------------------------------------------------------------------- panel b
sub = [r for r in rows if r["burial"] >= 50 and np.isfinite(r["best_foreign_overlap"])]
c_b = np.array([r["best_foreign_overlap"] for r in sub if r["kind"] == "control"])
t_b = np.array([r["best_foreign_overlap"] for r in sub if r["kind"] == "test"])
u_stat, p_val = mannwhitneyu(c_b, t_b, alternative="two-sided")

rng = np.random.default_rng(20250823)
def jitter(n, w=0.12):
    return rng.uniform(-w, w, size=n)
a1.scatter(np.zeros_like(c_b) + jitter(len(c_b)), c_b, color=C_CTRL, s=22, alpha=0.75,
           edgecolor="white", linewidth=0.4)
a1.scatter(np.ones_like(t_b) + jitter(len(t_b)), t_b, color=C_TEST, s=22, alpha=0.75,
           edgecolor="white", linewidth=0.4, marker="^")
a1.hlines(np.median(c_b), -0.3, 0.3, color=C_CTRL, lw=2.2)
a1.hlines(np.median(t_b), 0.7, 1.3, color=C_TEST, lw=2.2)
a1.axhline(0, color="#888888", lw=0.5, ls=":")

y_top = 2.15
a1.plot([0, 0, 1, 1], [y_top - 0.35, y_top, y_top, y_top - 0.35], color="#333333", lw=0.8)
a1.text(0.5, y_top + 0.10, f"p = {p_val:.3f}", ha="center", fontsize=ANN, color="#333333")
a1.text(0, y_top + 0.35, f"n = {len(c_b)}", ha="center", fontsize=ANN, color="#333333")
a1.text(1, y_top + 0.35, f"n = {len(t_b)}", ha="center", fontsize=ANN, color="#333333")

a1.set_xticks([0, 1])
a1.set_xticklabels(["Met1\nobserved", "Met1\nabsent"], fontsize=AX)
a1.set_xlim(-0.6, 1.6)
a1.set_ylim(-4.2, 3.0)
a1.set_ylabel("Best-case overlap (\u00c5)", fontsize=AX)
a1.set_title("b   Primary comparison: deeply buried termini (\u2265 50%)", fontsize=PANEL, loc="left")
a1.spines[["top", "right"]].set_visible(False)

# ---------------------------------------------------------------------- panel c
obs = np.array([r["observed_met1_overlap"] for r in ctrl if np.isfinite(r["observed_met1_overlap"])])
reb = np.array([r["best_foreign_overlap"] for r in ctrl if np.isfinite(r["best_foreign_overlap"])])
bins = np.arange(-10.0, 2.001, 0.5)
a2.hist(obs, bins=bins, color=C_CTRL, alpha=0.75, edgecolor="white",
        label=f"observed Met1, as deposited (n = {len(obs)})")
a2.hist(reb, bins=bins, color="#4a6a86", alpha=0.55, edgecolor="white",
        label=f"same Met1, deleted and rebuilt (n = {len(reb)})")
a2.axvline(0, color="#888888", lw=0.5, ls=":")
a2.set_xlim(-10.2, 2.2)
a2.set_xlabel("Steric overlap (\u00c5)", fontsize=AX)
a2.set_ylabel("Control termini", fontsize=AX)
a2.set_title("c   Control-score calibration", fontsize=PANEL, loc="left")
a2.legend(fontsize=ANN, frameon=False, loc="upper left", handletextpad=0.5)
a2.spines[["top", "right"]].set_visible(False)

fig.savefig(OUT + "fig5_conformer_scan.png", dpi=600)
plt.close(fig)

# ---------------------------------------------------------------- audit trail
n_test_finite = sum(1 for r in test if np.isfinite(r["best_foreign_overlap"]))
n_test_all = len(test)
print(f"panel a: {n_test_finite} of {n_test_all} test termini have finite overlap")
print(f"panel b: n_ctrl={len(c_b)}, n_test={len(t_b)}, "
      f"medians {np.median(c_b):.2f} vs {np.median(t_b):.2f}, "
      f"Mann-Whitney two-sided p={p_val:.3f}")
print(f"panel c: observed n={len(obs)}, rebuilt n={len(reb)}")
print(f"wrote {OUT}fig5_conformer_scan.png")
```
