## Supplementary_Data_8a for "Human protein complex interfaces reveal candidate assembly roles for initiator methionine excision": rebuild_supplementary_figure2.py

```
import os as _os
BASE = _os.environ.get("NME_DIR") or _os.path.dirname(_os.path.dirname(_os.path.abspath(__file__)))
INPUT_DIR = _os.path.join(BASE, "inputs")
WORK_DIR = _os.environ.get("NME_WORK") or _os.path.join(BASE, "work")
_os.makedirs(WORK_DIR, exist_ok=True)
"""Supplementary Figure S2 for revision the deposited analysis: downstream N-alpha-modification
branches vs interface geometry.

Authored at the final printed width (180 mm) so the point sizes here are the point
sizes on the page, and panel letters are lowercase to match the figure legend.
Do not widen the figsize without scaling the font sizes to match.
"""
import json
import numpy as np
import matplotlib
matplotlib.use("Agg")
import matplotlib.pyplot as plt
from matplotlib.patches import Patch

OUT = f"{WORK_DIR}/"
C_NME = "#3b6ea5"
C_MYR = "#7a4b9c"
C_MET = "#1f8a70"
C_GREY = "#9a9a9a"

recs = json.load(open(OUT + "asm_cascade_records.json"))
nme = [r for r in recs if r["pred_nme"]]
myr = [r for r in nme if r["myr"]]
mot = [r for r in nme if r["ntmt_motif"] and not r["myr"]]
ac = [r for r in nme if r["acetyl"] and not r["myr"] and not r["ntmt_motif"]]
oth = [r for r in nme if not r["myr"] and not r["ntmt_motif"] and not r["acetyl"]]

groups = [("myristoylated\nGly$_1$", myr, C_MYR),
          ("NTMT motif\n[APSG]-P-K", mot, C_MET),
          ("Nt-acetylated", ac, C_NME),
          ("no\nannotation", oth, C_GREY)]

W = 7.09
ANN = 6.6
PANEL = 8.6
AX = 7.4
plt.rcParams.update({"font.size": ANN, "xtick.labelsize": ANN, "ytick.labelsize": ANN})
fig, (a1, a2) = plt.subplots(1, 2, figsize=(W, 4.1), layout="constrained",
                             gridspec_kw=dict(width_ratios=[1.18, 1.25]))

# ---- panel A: burial by branch
rng = np.random.default_rng(7)
for i, (lab, g, c) in enumerate(groups):
    b = np.array([r["buried"] for r in g])
    x = i + rng.uniform(-0.20, 0.20, len(b))
    a1.scatter(x, b, s=26 if len(b) < 40 else 8, alpha=0.75 if len(b) < 40 else 0.30,
               color=c, edgecolors="none", zorder=3)
    a1.plot([i - 0.34, i + 0.34], [np.median(b)] * 2, color="black", lw=2.2, zorder=4)
    a1.text(i, 106, f"n = {len(b)}", ha="center", fontsize=ANN, color="#333333")
    a1.text(i, -11.5, f"med. {np.median(b):.1f}%", ha="center", fontsize=ANN, color=c)
a1.axhline(25, color="#a33227", lw=0.9, ls="--")
a1.text(3.42, 27, "25%", fontsize=ANN, color="#a33227", ha="right")
a1.set_xticks(range(len(groups)))
a1.set_xticklabels([g[0] for g in groups], fontsize=6.2)
a1.set_ylim(-16, 112)
a1.set_xlim(-0.6, len(groups) - 0.4)
a1.set_ylabel("Burial of residue 1 at inter-subunit\ninterfaces (%)", fontsize=AX)
a1.set_title("a   Engagement by downstream branch", fontsize=PANEL,
             loc="left", fontweight="bold", pad=12)

# ---- panel B: individual cases
cases = sorted(myr, key=lambda r: r["buried"]) + [None] + sorted(mot, key=lambda r: r["buried"])
labels, vals, cols, dns = [], [], [], []
for r in cases:
    if r is None:
        labels.append("")
        vals.append(0.0)
        cols.append("white")
        dns.append(None)
        continue
    tag = "G" if r["myr"] else r["m1"] + r["m2"] + r["m3"]
    labels.append(f"{r['gene']}  ({tag}$_1$)" if r["myr"] else f"{r['gene']}  ({tag})")
    vals.append(r["buried"])
    cols.append(C_MYR if r["myr"] else C_MET)
    dns.append(r["dN"])
y = np.arange(len(vals))
a2.barh(y, vals, color=cols, height=0.66, edgecolor="white")
for yy, v, d in zip(y, vals, dns):
    if d is None:
        continue
    txt = f"{v:.1f}%" + (f"   {d:.2f} Å" if d is not None and d < 30 else "   no contact")
    a2.text(v + 1.6, yy, txt, va="center", fontsize=ANN, color="#333333",
            bbox=dict(facecolor="white", edgecolor="none", pad=0.6))
a2.set_yticks(y)
a2.set_yticklabels(labels, fontsize=ANN)
a2.set_xlim(0, 128)
a2.set_xlabel("Burial of residue 1 (%); right-hand value: minimum\n"
              "distance from the $\\alpha$-amino N to a neighbouring subunit", fontsize=AX)
a2.set_title("b   Termini committed to a modification", fontsize=PANEL,
             loc="left", fontweight="bold", pad=12)
a2.axvline(25, color="#a33227", lw=0.9, ls="--")
a2.legend(handles=[Patch(facecolor=C_MYR, label="N-myristoylated (annotated)"),
                   Patch(facecolor=C_MET, label="NTMT recognition motif")],
          fontsize=ANN, frameon=False, loc="lower right", bbox_to_anchor=(1.0, -0.005),
          handlelength=1.1, handletextpad=0.5)

for ax in (a1, a2):
    ax.set_axisbelow(True)
    ax.grid(color="#e8e8e8", lw=0.6)
    for s in ("top", "right"):
        ax.spines[s].set_visible(False)

OUTDIR = _os.environ.get("NME_FIGOUT") or WORK_DIR
_os.makedirs(OUTDIR, exist_ok=True)
fig.savefig(OUTDIR + "/figS2_cascade.png", dpi=600, facecolor="white")
print("wrote figS2_cascade.png")
```
