## Supplementary_Data_8a for "Human protein complex interfaces reveal candidate assembly roles for initiator methionine excision": run_robustness_analyses.py

```
"""Reproduce the deposited robustness analyses from canonical data tables."""
import os
from pathlib import Path
import pandas as pd, numpy as np
from scipy.stats import fisher_exact, norm
from scipy.optimize import minimize

BASE = Path(os.environ.get("NME_DIR") or Path(__file__).resolve().parents[1])
DATA_DIR = Path(os.environ.get("NME_WORK") or BASE / "data")
OUT_DIR = Path(os.environ.get("NME_OUT") or DATA_DIR)
OUT_DIR.mkdir(parents=True, exist_ok=True)

p=pd.read_csv(DATA_DIR / "asm_per_protein_master.csv")
p=p[~p['cls'].astype(str).str.startswith('excluded')].copy()
p['excision']=(p['pred']=='predicted NME').astype(int)
p['engaged']=(p['buried_pct']>=25).astype(int)

def fit(df, cols):
 X=np.column_stack([np.ones(len(df)),df[cols].to_numpy(float)]); y=df.engaged.to_numpy(float)
 def fun(b):
  z=X@b
  return np.sum(np.logaddexp(0,z)-y*z)
 def jac(b):
  z=X@b; pr=1/(1+np.exp(-np.clip(z,-40,40)))
  return X.T@(pr-y)
 m=minimize(fun,np.zeros(X.shape[1]),jac=jac,method='BFGS')
 b=m.x; z=X@b; pr=1/(1+np.exp(-np.clip(z,-40,40)))
 H=X.T@((pr*(1-pr))[:,None]*X); cov=np.linalg.inv(H); se=np.sqrt(np.diag(cov))
 j=1; zz=b[j]/se[j]; ci=(b[j]-1.96*se[j],b[j]+1.96*se[j])
 return dict(n=len(df),odds_ratio=np.exp(b[j]),ci_low=np.exp(ci[0]),ci_high=np.exp(ci[1]),p=2*norm.sf(abs(zz)))

p['log_chains']=np.log(p.n_chains.clip(lower=1))
p['log_entries']=np.log(p.n_entries.clip(lower=1))
rows=[]
rows.append({'analysis':'class + log chain count',**fit(p,['excision','log_chains'])})
rows.append({'analysis':'class + log chain count + log PDB-entry count',**fit(p,['excision','log_chains','log_entries'])})

loo=[]
for aa in sorted(p.res2.dropna().unique()):
 d=p[p.res2!=aa]
 a=pd.crosstab(d.excision,d.engaged).reindex(index=[1,0],columns=[1,0],fill_value=0)
 OR,pv=fisher_exact([[a.loc[1,1],a.loc[1,0]],[a.loc[0,1],a.loc[0,0]]])
 loo.append({'excluded_residue':aa,'n_excision':int((d.excision==1).sum()),'n_refractory':int((d.excision==0).sum()),'odds_ratio':OR,'fisher_p':pv})

d=p[~((p.excision==1)&(p.resname.isin(['MET','MSE'])))]
a=pd.crosstab(d.excision,d.engaged).reindex(index=[1,0],columns=[1,0],fill_value=0)
OR,pv=fisher_exact([[a.loc[1,1],a.loc[1,0]],[a.loc[0,1],a.loc[0,0]]])
metex={'analysis':'exclude predicted-excision proteins modeled as MET/MSE','n_excision':int((d.excision==1).sum()),'n_refractory':int((d.excision==0).sum()),'odds_ratio':OR,'fisher_p':pv,'engaged_excision':float(d[d.excision==1].engaged.mean()),'engaged_refractory':float(d[d.excision==0].engaged.mean())}

pd.DataFrame(rows).to_csv(OUT_DIR / "multiplicity_models.csv",index=False)
pd.DataFrame(loo).to_csv(OUT_DIR / "leave_one_residue_out.csv",index=False)
pd.DataFrame([metex]).to_csv(OUT_DIR / "met_mse_exclusion.csv",index=False)
print(pd.DataFrame(rows).to_string(index=False))
print(pd.DataFrame(loo).agg({'odds_ratio':['min','max'],'fisher_p':['min','max']}))
print(metex)
```
