## Supplementary_Data_8a for "Human protein complex interfaces reveal candidate assembly roles for initiator methionine excision": scan_methionine.py

```
"""Run the deposited end-to-end Figure 5 methionine conformer scan.

Faithful re-implementation of the algorithm described in the manuscript Methods
(paragraphs 668-676): an initiator methionine is built N-terminal to the mature
residue with idealized geometry (C-N 1.329 A, C-N-CA 121.7 deg, omega 180 deg),
scanned over phi (of the mature residue, which does not exist until the methionine
is present) and psi (of the built methionine) at 10 degree spacing, and over three
canonical rotameric states for each of chi1, chi2, chi3 -- so 36 * 36 * 3 * 3 * 3
= 34,992 conformers per terminus. Van der Waals overlap is scored as
r_i + r_j - t - d_ij over all atom pairs, with t = 0.4 A against atoms of other
chains and t = 0.6 A against the terminus's own chain (intrachain strain is
relievable). Conformers failing the intrachain criterion are discarded; the minimum
interchain overlap among the remainder is retained.

This script performs the geometry using the 11 frozen biological-assembly mmCIF
files whose PDB codes are in data/fig5_targets.csv
(4R3O 5A9Q 5LN3 6KWY 6N88 6REY 7MO1 7V5G 8CVS 8TM4 8YIX). It writes the same
records table (data/fig5_records.csv) that rebuild_figure5.py consumes, so the end-
to end the pipeline is now:

    mmCIF files
        -> scan_methionine.py
        -> fig5_records.csv
        -> rebuild_figure5.py
        -> fig5_conformer_scan.png

The frozen coordinates and their SHA-256 checksums are included in the deposit.
The four positions with no conformer passing the fixed intrachain criterion remain
in the output with an empty best_foreign_overlap field.

Dependencies:  gemmi >= 0.6, numpy.  Freesasa is only used by the burial pipeline
in step3_geometry.py -- not by this script.

Usage:  python scan_methionine.py --cif-dir <dir> --targets <termini.csv> \\
                                  --out data/fig5_records.csv
        where <termini.csv> lists the termini to scan, one per row:
            gene,pdb,chain,kind,res2
        Set kind = "control" if methionine 1 is resolved in the deposited chain
        and "test" if the mature residue is res2. This is the same column layout
        as the deposited fig5_records.csv, minus the numeric columns which this
        script fills in.
"""
from __future__ import annotations
import argparse
import csv
import itertools
import math
import os
import sys
from dataclasses import dataclass

import numpy as np

try:
    import gemmi
except ImportError:  # pragma: no cover
    sys.stderr.write("This script needs gemmi (`pip install gemmi`).\n")
    raise

# ---------------------------------------------------------------- constants
VDW = {"H": 1.20, "C": 1.70, "N": 1.55, "O": 1.52, "S": 1.80,
       "P": 1.80, "F": 1.47, "CL": 1.75, "BR": 1.85, "I": 1.98}
T_SELF = 0.60          # intrachain overlap tolerance (A), per Methods paragraph 672
T_FOREIGN = 0.40       # interchain overlap tolerance (A)
CLASH_CUT = 6.0        # only atom pairs closer than this contribute
# idealized peptide bond that grafts the initiator methionine on
C_N_BOND = 1.329
C_N_CA = math.radians(121.7)
OMEGA = math.pi        # trans
CA_C_N = math.radians(116.2)
N_CA_C = math.radians(111.2)
CA_C = 1.525
N_CA = 1.458
# methionine side-chain internal geometry
CA_CB = 1.530
CB_CG = 1.520
CG_SD = 1.807
SD_CE = 1.791
CB_CG_SD = math.radians(112.7)
CG_SD_CE = math.radians(100.5)
CA_CB_CG = math.radians(114.4)
N_CA_CB = math.radians(110.5)

# 36 phi and 36 psi values at 10 degree spacing over -180..170; three staggered rotamers
# for each of chi1, chi2, chi3.
PHI_GRID = np.deg2rad(np.arange(-180, 180, 10))
PSI_GRID = np.deg2rad(np.arange(-180, 180, 10))
CHI1 = np.deg2rad([-60.0, 60.0, 180.0])
CHI2 = np.deg2rad([-60.0, 60.0, 180.0])
CHI3 = np.deg2rad([-60.0, 60.0, 180.0])
assert len(PHI_GRID) * len(PSI_GRID) * len(CHI1) * len(CHI2) * len(CHI3) == 34_992


# --------------------------------------------------------------- geometry
def place(prev: np.ndarray, mid: np.ndarray, ref: np.ndarray,
          bond: float, angle: float, torsion: float) -> np.ndarray:
    """Place an atom given the previous three anchors, a bond, an angle
    (mid-ref-new) and a torsion (prev-mid-ref-new)."""
    a = prev - mid
    b = ref - mid
    bn = b / np.linalg.norm(b)
    an = a / np.linalg.norm(a)
    # local frame
    n = np.cross(bn, an)
    n /= np.linalg.norm(n)
    m = np.cross(n, bn)
    d = np.array([bond * math.cos(math.pi - angle),
                  bond * math.sin(math.pi - angle) * math.cos(torsion),
                  bond * math.sin(math.pi - angle) * math.sin(torsion)])
    return ref + d[0] * bn + d[1] * m + d[2] * n


def build_methionine(N2: np.ndarray, CA2: np.ndarray, C2: np.ndarray,
                     phi: float, psi: float,
                     chi1: float, chi2: float, chi3: float
                     ) -> np.ndarray:
    """Return coordinates [N, CA, C, O, CB, CG, SD, CE] for the built Met.
    N2/CA2/C2 are the coordinates of the mature residue whose phi is `phi`."""
    # graft: place the new C (of the built methionine) so that phi(new C, N2, CA2, C2) == phi
    # phi of the mature residue = torsion(C_prev, N, CA, C). C_prev is the new methionine's C.
    C_new = place(C2, CA2, N2, C_N_BOND, C_N_CA, phi)
    # CA_new: place so that omega(CA_new, C_new, N2, CA2) is 180.
    CA_new = place(N2, C_new, C2, CA_C, math.pi - CA_C_N, OMEGA)   # dummy, replaced below
    # simpler: build the Met backbone by inverse propagation using standard geometry.
    # place CA_new using omega=180 torsion (CA2 - N2 - C_new - CA_new)
    CA_new = place(CA2, N2, C_new, CA_C, math.pi - CA_C_N, OMEGA)
    # N_new: using psi (of the built Met) = torsion(N_new, CA_new, C_new, N2)
    N_new = place(N2, C_new, CA_new, N_CA, math.pi - N_CA_C, psi)
    # O of the built methionine: on C_new, opposite N2 (idealized).
    O_new = place(CA_new, N_new, C_new, 1.229, math.radians(120.5), math.pi)
    # side chain: chi1 rotates about N-CA-CB-CG
    CB = place(C_new, N_new, CA_new, CA_CB, math.pi - N_CA_CB, math.radians(122.5))
    CG = place(N_new, CA_new, CB, CB_CG, math.pi - CA_CB_CG, chi1)
    SD = place(CA_new, CB, CG, CG_SD, math.pi - CB_CG_SD, chi2)
    CE = place(CB, CG, SD, SD_CE, math.pi - CG_SD_CE, chi3)
    return np.vstack([N_new, CA_new, C_new, O_new, CB, CG, SD, CE])


# --------------------------------------------------------------- scoring
def overlap(built: np.ndarray, built_elems: list[str],
            neighbours: np.ndarray, neigh_elems: list[str],
            tol: float) -> float:
    """Maximum vdW overlap of any built-vs-neighbour atom pair, above zero."""
    if len(neighbours) == 0:
        return -np.inf
    d = np.linalg.norm(built[:, None, :] - neighbours[None, :, :], axis=-1)
    r_i = np.array([VDW.get(e.upper(), 1.7) for e in built_elems])[:, None]
    r_j = np.array([VDW.get(e.upper(), 1.7) for e in neigh_elems])[None, :]
    o = r_i + r_j - tol - d
    mask = d < CLASH_CUT
    return float(o[mask].max()) if mask.any() else -np.inf


# --------------------------------------------------------------- driver
@dataclass
class Terminus:
    gene: str
    pdb: str
    chain: str
    kind: str
    res2: str          # single-letter code of the mature residue at position 2 of unprocessed


def load_targets(path: str) -> list[Terminus]:
    out = []
    for row in csv.DictReader(open(path)):
        out.append(Terminus(row["gene"], row["pdb"], row["chain"],
                            row["kind"], row["res2"]))
    return out


def load_structure(pdb: str, cif_dir: str) -> gemmi.Structure:
    for cand in (os.path.join(cif_dir, f"{pdb.lower()}.cif"),
                 os.path.join(cif_dir, f"{pdb.lower()}.cif.gz"),
                 os.path.join(cif_dir, f"{pdb.upper()}.cif")):
        if os.path.exists(cand):
            return gemmi.read_structure(cand)
    raise FileNotFoundError(pdb)


def _resolve_chain(model: gemmi.Model, want: str) -> str | None:
    """Resolve a terminus chain code (e.g. 'B1', 'P1') to a label_asym_id in the
    deposited model. The trailing digit is the biological-assembly operator index;
    the leading letters are the auth_asym_id, which matches the label_asym_id for
    the original chain (extra letters like 'AA', 'BA' are assembly-generated
    copies). We match on auth chain letters only."""
    names = {ch.name for ch in model}
    # Assembly downloads encode operator copies as AUTH-OP (for example the
    # legacy target code "72" corresponds to auth chain 7, operator 2, stored
    # by gemmi as "7-2"). Prefer an exact name, then this explicit mapping.
    if want in names:
        return want
    if len(want) >= 2 and want[-1].isdigit():
        assembly_name = f"{want[:-1]}-{want[-1]}"
        if assembly_name in names:
            return assembly_name
    letters = want.rstrip("0123456789") or want
    for ch in model:
        if ch.name == letters:
            return ch.name
    for ch in model:
        if ch.name.rstrip("0123456789") == letters:
            return ch.name
    return None


def scan_one(terminus: Terminus, structure: gemmi.Structure) -> dict:
    """Return the row of numeric fields for this terminus."""
    model = structure[0]
    chain_name = _resolve_chain(model, terminus.chain)
    if chain_name is None:
        raise KeyError(f"{terminus.pdb}:{terminus.chain} not in structure")
    aa3 = {"ALA","ARG","ASN","ASP","CYS","GLU","GLN","GLY","HIS","ILE",
           "LEU","LYS","MET","PHE","PRO","SER","THR","TRP","TYR","VAL","MSE"}
    # Mature residue is the position-2 residue of the unprocessed protein. In
    # test structures Met1 is not resolved, so the first amino acid IS position 2.
    # In control structures Met1 is resolved at seqid 1, so the mature residue
    # is the SECOND amino acid. The scan pretends Met1 is absent and asks "can
    # a methionine be modelled at position 1?" - so at controls, atoms of the
    # resolved Met1 must be excluded from both self and other neighbour sets.
    mature = None
    met1_resid = None
    for ch in model:
        if ch.name != chain_name:
            continue
        aas = [r for r in ch if r.name in aa3]
        if not aas:
            break
        if terminus.kind == "control":
            # find and skip Met1 (or MSE) at position 1
            if aas[0].name in ("MET", "MSE") and len(aas) >= 2:
                met1_resid = aas[0].seqid.num
                mature = aas[1]
            else:
                # kind says control but no Met1 - fall back to first residue
                mature = aas[0]
        else:
            mature = aas[0]
        break
    if mature is None:
        raise LookupError(f"no amino acid in {terminus.pdb}:{terminus.chain}")
    N2 = np.array(mature["N"][0].pos.tolist())
    CA2 = np.array(mature["CA"][0].pos.tolist())
    C2 = np.array(mature["C"][0].pos.tolist())
    # self atoms (same chain, excluding the mature residue): intrachain criterion
    self_atoms, self_elems = [], []
    other_atoms, other_elems = [], []
    for ch in model:
        for r in ch:
            for at in r:
                p = at.pos.tolist()
                if ch.name == chain_name and r.seqid.num == mature.seqid.num:
                    continue
                # Exclude the resolved Met1 from the neighbour set at controls;
                # we are asking whether a methionine can be built there, so it
                # cannot clash with itself.
                if ch.name == chain_name and met1_resid is not None \
                        and r.seqid.num == met1_resid:
                    continue
                elem = at.element.name
                if ch.name == chain_name:
                    self_atoms.append(p)
                    self_elems.append(elem)
                else:
                    other_atoms.append(p)
                    other_elems.append(elem)
    self_atoms = np.asarray(self_atoms, float) if self_atoms else np.zeros((0, 3))
    other_atoms = np.asarray(other_atoms, float) if other_atoms else np.zeros((0, 3))
    # Neighbourhood filter: any atom of the built methionine sits within about
    # 8 A of N2. Only atoms within N2-radius + 8 + max(vdW) can ever contribute
    # to a clash. A 15 A cut around N2 is safe.
    NEIGH_CUT = 15.0
    if len(self_atoms):
        keep = np.linalg.norm(self_atoms - N2, axis=1) <= NEIGH_CUT
        self_atoms = self_atoms[keep]
        self_elems = [self_elems[i] for i in np.flatnonzero(keep)]
    if len(other_atoms):
        keep = np.linalg.norm(other_atoms - N2, axis=1) <= NEIGH_CUT
        other_atoms = other_atoms[keep]
        other_elems = [other_elems[i] for i in np.flatnonzero(keep)]
    built_elems = ["N", "C", "C", "O", "C", "C", "S", "C"]
    r_built = np.array([VDW[e] for e in built_elems])                     # (8,)
    r_self = np.array([VDW.get(e.upper(), 1.7) for e in self_elems])      # (Ns,)
    r_other = np.array([VDW.get(e.upper(), 1.7) for e in other_elems])    # (No,)
    self_pair = r_built[:, None] + r_self[None, :] - T_SELF               # (8, Ns)
    other_pair = r_built[:, None] + r_other[None, :] - T_FOREIGN          # (8, No)

    n_self_ok = 0
    n_both_ok = 0
    best_foreign_overlap = np.inf
    for phi, psi, c1, c2, c3 in itertools.product(PHI_GRID, PSI_GRID, CHI1, CHI2, CHI3):
        built = build_methionine(N2, CA2, C2, phi, psi, c1, c2, c3)
        if len(self_atoms):
            d = np.linalg.norm(built[:, None, :] - self_atoms[None, :, :], axis=-1)
            if (self_pair - d).max() > 0:
                continue
        n_self_ok += 1
        if len(other_atoms):
            d2 = np.linalg.norm(built[:, None, :] - other_atoms[None, :, :], axis=-1)
            fo = float((other_pair - d2).max())
        else:
            fo = -np.inf
        if fo <= 0:
            n_both_ok += 1
        if fo < best_foreign_overlap:
            best_foreign_overlap = fo
    # best_foreign_overlap = -inf means "no foreign atoms within reach" (solvent-
    # exposed terminus, no possible interchain clash). Report as a large negative
    # sentinel so downstream stats don't drop the row. Only empty when no conformer
    # passed intrachain, so n_self_ok == 0.
    if n_self_ok == 0:
        best_foreign_overlap_val: float | str = ""
    elif not math.isfinite(best_foreign_overlap):
        best_foreign_overlap_val = -9.99
    else:
        best_foreign_overlap_val = round(best_foreign_overlap, 2)
    return {
        "n_self_ok": n_self_ok,
        "n_both_ok": n_both_ok,
        "n_total": 34_992,
        "best_foreign_overlap": best_foreign_overlap_val,
    }


def main() -> None:
    ap = argparse.ArgumentParser(description=__doc__.splitlines()[0])
    ap.add_argument("--cif-dir", required=True, help="directory of mmCIF files")
    ap.add_argument("--targets", required=True, help="CSV of termini to scan")
    ap.add_argument("--out", required=True, help="output records table")
    ap.add_argument("--start", type=int, default=0, help="first target index (0-based, inclusive)")
    ap.add_argument("--limit", type=int, default=None, help="maximum number of targets to scan")
    ap.add_argument("--append", action="store_true", help="append to existing output; suppress header")
    args = ap.parse_args()

    targets = load_targets(args.targets)
    if args.limit is not None:
        targets = targets[args.start:args.start + args.limit]
    else:
        targets = targets[args.start:]
    struct_cache: dict[str, gemmi.Structure] = {}
    fields = ["gene", "pdb", "chain", "kind", "res2",
              "n_self_ok", "n_both_ok", "n_total", "best_foreign_overlap"]
    mode = "a" if args.append else "w"
    with open(args.out, mode, newline="", buffering=1) as fh:
        writer = csv.DictWriter(fh, fieldnames=fields)
        if not args.append:
            writer.writeheader()
        for t in targets:
            row = {"gene": t.gene, "pdb": t.pdb, "chain": t.chain,
                   "kind": t.kind, "res2": t.res2,
                   "n_self_ok": "", "n_both_ok": "", "n_total": 34_992,
                   "best_foreign_overlap": ""}
            try:
                if t.pdb not in struct_cache:
                    struct_cache[t.pdb] = load_structure(t.pdb, args.cif_dir)
                row.update(scan_one(t, struct_cache[t.pdb]))
            except (KeyError, LookupError, FileNotFoundError) as exc:
                print(t.pdb, t.chain, t.gene, f"SKIP: {exc}", flush=True)
            else:
                print(t.pdb, t.chain, t.gene, row["best_foreign_overlap"], flush=True)
            writer.writerow(row)
            fh.flush()


if __name__ == "__main__":
    main()
```
