## Supplementary_Data_8a for "Human protein complex interfaces reveal candidate assembly roles for initiator methionine excision": step1_build_targets.py

```
import os as _os
BASE = _os.environ.get("NME_DIR") or _os.path.dirname(_os.path.dirname(_os.path.abspath(__file__)))
INPUT_DIR = _os.path.join(BASE, "inputs")
WORK_DIR = _os.environ.get("NME_WORK") or _os.path.join(BASE, "work")
_os.makedirs(WORK_DIR, exist_ok=True)
import gzip, csv, re, json, collections

# ---------- 1. Parse UniProt human N-terminal annotations ----------
prot = {}
with gzip.open(f"{INPUT_DIR}/human_nterm.tsv.gz", "rt") as fh:
    rd = csv.DictReader(fh, delimiter="\t")
    for r in rd:
        seq = r["Sequence"]
        if len(seq) < 30: continue
        acc = r["Entry"]
        init = r["Initiator methionine"] or ""
        modres = r["Modified residue"] or ""
        init_removed = "INIT_MET" in init and "Removed" in init
        # Nt-acetylation annotated at position 1 or 2
        nac = ""
        for m in re.finditer(r"MOD_RES (\d+);([^;]*);?", modres):
            pos = int(m.group(1)); note = m.group(2)
            if pos <= 2 and "acetyl" in note.lower():
                nac = note.split('note="')[-1].strip('"')
        has_sig = bool(r["Signal peptide"].strip())
        has_pro = bool(r["Propeptide"].strip())
        has_tra = bool(r["Transit peptide"].strip())
        mature_start = 2 if init_removed else 1
        prot[acc] = dict(acc=acc, gene=r["Gene Names (primary)"], length=int(r["Length"]),
                         res1=seq[0], res2=seq[1], res3=seq[2],
                         init_removed=init_removed, nt_acetyl=nac,
                         has_signal=has_sig, has_propep=has_pro, has_transit=has_tra,
                         mature_start=mature_start, seq=seq)
print("human proteins parsed:", len(prot))
print("annotated iMet removed:", sum(p['init_removed'] for p in prot.values()))
print("annotated Nt-acetyl:", sum(bool(p['nt_acetyl']) for p in prot.values()))

# ---------- 2. SIFTS PDB chain -> UniProt ----------
rows = []
entry_chains = collections.defaultdict(set)
entry_accs = collections.defaultdict(set)
with gzip.open(f"{INPUT_DIR}/pdb_chain_uniprot.tsv.gz", "rt") as fh:
    fh.readline()  # comment line
    rd = csv.DictReader(fh, delimiter="\t")
    for r in rd:
        pdb = r["PDB"].lower(); ch = r["CHAIN"]; acc = r["SP_PRIMARY"]
        entry_chains[pdb].add(ch)
        entry_accs[pdb].add(acc)
        if acc not in prot: continue
        try:
            sp_beg = int(r["SP_BEG"]); pdb_beg = r["PDB_BEG"]
        except ValueError:
            continue
        rows.append((pdb, ch, acc, sp_beg, pdb_beg, r["PDB_END"], int(r["SP_END"])))
print("SIFTS entries:", len(entry_chains), "| human-mapped chain segments:", len(rows))

# ---------- 3. keep chains whose mapping covers the mature N-terminal residue ----------
targets = []
for pdb, ch, acc, sp_beg, pdb_beg, pdb_end, sp_end in rows:
    p = prot[acc]
    if sp_beg > p["mature_start"]:      # N-terminal residue not in this segment
        continue
    nch = len(entry_chains[pdb]); nacc = len(entry_accs[pdb])
    targets.append(dict(pdb=pdb, chain=ch, acc=acc, sp_beg=sp_beg,
                        mature_start=p["mature_start"], n_chains=nch, n_entities=nacc))
print("chain segments covering the mature N-terminus:", len(targets))
print("distinct proteins with a modeled N-terminus:", len({t['acc'] for t in targets}))

# prioritise: heteromeric complexes first, then more chains
by_entry = collections.defaultdict(list)
for t in targets: by_entry[t["pdb"]].append(t)
entries = sorted(by_entry, key=lambda e: (-(by_entry[e][0]["n_entities"] > 1),
                                          -by_entry[e][0]["n_entities"],
                                          -by_entry[e][0]["n_chains"]))
multi = [e for e in entries if by_entry[e][0]["n_chains"] >= 2]
print("entries with >=2 chains:", len(multi), "| heteromeric (>1 distinct UniProt):",
      len([e for e in multi if by_entry[e][0]["n_entities"] > 1]))

json.dump({"prot": {a: {k: v for k, v in p.items() if k != 'seq'} for a, p in prot.items()},
           "targets": targets, "entry_order": multi},
          open(f"{WORK_DIR}/targets.json", "w"))
print("saved targets.json")
```
