## Supplementary_Data_8a for "Human protein complex interfaces reveal candidate assembly roles for initiator methionine excision": step2_select_download.py

```
import os as _os
BASE = _os.environ.get("NME_DIR") or _os.path.dirname(_os.path.dirname(_os.path.abspath(__file__)))
INPUT_DIR = _os.path.join(BASE, "inputs")
WORK_DIR = _os.environ.get("NME_WORK") or _os.path.join(BASE, "work")
_os.makedirs(WORK_DIR, exist_ok=True)
import json, collections, os, urllib.request, gzip, io
from concurrent.futures import ThreadPoolExecutor

D = json.load(open(f"{WORK_DIR}/targets.json"))
targets, order = D["targets"], D["entry_order"]
by_entry = collections.defaultdict(list)
for t in targets: by_entry[t["pdb"]].append(t)

MAX_CHAINS = 40
hetero = [e for e in order if by_entry[e][0]["n_entities"] > 1 and by_entry[e][0]["n_chains"] <= MAX_CHAINS]
homo   = [e for e in order if by_entry[e][0]["n_entities"] == 1 and by_entry[e][0]["n_chains"] <= MAX_CHAINS]

# greedy set cover on distinct accessions, heteromers first
covered, chosen = set(), []
def greedy(pool, budget):
    global covered
    pool = list(pool)
    while pool and len(chosen) < budget:
        best, gain = None, -1
        # cheap scan: score = new accessions added
        for e in pool[:4000]:
            g = len({t["acc"] for t in by_entry[e]} - covered)
            if g > gain: best, gain = e, g
            if gain >= 8: break
        if best is None or gain <= 0:
            # no more gain from scanned window; drop the window
            pool = pool[4000:]
            if not pool: break
            continue
        chosen.append(best); covered |= {t["acc"] for t in by_entry[best]}
        pool.remove(best)

greedy(hetero, 1400)
greedy(homo, 2200)
print("entries chosen:", len(chosen), "| distinct proteins covered:", len(covered), flush=True)
json.dump(chosen, open(f"{WORK_DIR}/greedy_list.json", "w"))

os.makedirs(f"{WORK_DIR}/pdb", exist_ok=True)
def get(pdb):
    out = f"{WORK_DIR}/pdb/{pdb}.pdb"
    if os.path.exists(out) and os.path.getsize(out) > 1000: return (pdb, "cached")
    try:
        raw = urllib.request.urlopen(f"https://files.rcsb.org/download/{pdb.upper()}.pdb.gz", timeout=60).read()
        txt = gzip.decompress(raw)
        if len(txt) < 2000: return (pdb, "tiny")
        open(out, "wb").write(txt)
        return (pdb, "ok")
    except Exception as e:
        return (pdb, f"fail:{type(e).__name__}")

res = collections.Counter()
with ThreadPoolExecutor(max_workers=8) as ex:
    for i, (pdb, st) in enumerate(ex.map(get, chosen)):
        res[st.split(":")[0]] += 1
        if i % 200 == 0: print(i, dict(res), flush=True)
print("download summary:", dict(res), flush=True)
```
