## Supplementary_Data_8a for "Human protein complex interfaces reveal candidate assembly roles for initiator methionine excision": step3_geometry.py

```
import os as _os
BASE = _os.environ.get("NME_DIR") or _os.path.dirname(_os.path.dirname(_os.path.abspath(__file__)))
INPUT_DIR = _os.path.join(BASE, "inputs")
WORK_DIR = _os.environ.get("NME_WORK") or _os.path.join(BASE, "work")
_os.makedirs(WORK_DIR, exist_ok=True)
import gzip, csv, json, os, re, collections, warnings, sys
import numpy as np, freesasa
from scipy.spatial import cKDTree
warnings.filterwarnings("ignore")
freesasa.setVerbosity(freesasa.silent)

AA3 = {'ALA':'A','ARG':'R','ASN':'N','ASP':'D','CYS':'C','GLN':'Q','GLU':'E','GLY':'G','HIS':'H',
       'ILE':'I','LEU':'L','LYS':'K','MET':'M','PHE':'F','PRO':'P','SER':'S','THR':'T','TRP':'W',
       'TYR':'Y','VAL':'V','MSE':'M','SEC':'U','PYL':'O'}

D = json.load(open(f"{WORK_DIR}/targets.json"))
prot = D["prot"]; targets = D["targets"]
chosen = set(json.load(open(f"{WORK_DIR}/chosen.json")))

# UniProt sequences for the accessions we need
need = {t["acc"] for t in targets if t["pdb"] in chosen}
seqs = {}
with gzip.open(f"{INPUT_DIR}/human_nterm.tsv.gz", "rt") as fh:
    rd = csv.DictReader(fh, delimiter="\t")
    for r in rd:
        if r["Entry"] in need: seqs[r["Entry"]] = r["Sequence"]

# SIFTS segment starts:  (pdb, chain, acc) -> [(sp_beg, pdb_beg)]
segs = collections.defaultdict(list)
with gzip.open(f"{INPUT_DIR}/pdb_chain_uniprot.tsv.gz", "rt") as fh:
    fh.readline(); rd = csv.DictReader(fh, delimiter="\t")
    for r in rd:
        pdb = r["PDB"].lower()
        if pdb not in chosen: continue
        try: segs[(pdb, r["CHAIN"], r["SP_PRIMARY"])].append((int(r["SP_BEG"]), r["PDB_BEG"]))
        except ValueError: pass

by_entry = collections.defaultdict(list)
for t in targets:
    if t["pdb"] in chosen: by_entry[t["pdb"]].append(t)

def clean(path, out):
    """first model, ATOM records, primary altloc, standard/MSE residues only"""
    keep, seen_model = [], False
    with open(path) as fh:
        for ln in fh:
            rec = ln[:6]
            if rec == "MODEL ":
                if seen_model: break
                seen_model = True; continue
            if rec == "ENDMDL": break
            if rec not in ("ATOM  ", "HETATM"): continue
            resn = ln[17:20].strip()
            if rec == "HETATM" and resn != "MSE": continue
            if resn not in AA3: continue
            if ln[16] not in (" ", "A"): continue
            if ln[76:78].strip() == "H": continue
            keep.append("ATOM  " + ln[6:16] + " " + ("MET" if resn == "MSE" else resn).ljust(3) + ln[20:])
    if len(keep) < 100: return None
    open(out, "w").writelines(keep + ["END\n"])
    return len(keep)

def analyze(pdb):
    path = f"{WORK_DIR}/pdb/{pdb}.pdb"
    tmp = f"{WORK_DIR}/tmp_{pdb}.pdb"
    if not os.path.exists(path): return []
    try:
        if clean(path, tmp) is None: return []
        # observed residues + coords per chain
        atoms = collections.defaultdict(list)   # chain -> (authnum, icode, resn, atomname, xyz)
        for ln in open(tmp):
            if ln[:6] != "ATOM  ": continue
            ch = ln[21]; num = int(ln[22:26]); ic = ln[26]
            atoms[ch].append((num, ic, ln[17:20].strip(), ln[12:16].strip(),
                              (float(ln[30:38]), float(ln[38:46]), float(ln[46:54]))))
        if len(atoms) < 2: return []
        allxyz = np.array([a[4] for ch in atoms for a in atoms[ch]])
        chain_lab = np.array([ch for ch in atoms for a in atoms[ch]])

        s = freesasa.Structure(tmp, options={'hetatm': False, 'skip-unknown': True})
        cx = freesasa.calc(s).residueAreas()
        iso = {}
        for st in freesasa.structureArray(tmp, options={'separate-chains': True,
                                                       'hetatm': False, 'skip-unknown': True}):
            lab = st.chainLabel(1)
            iso[lab] = freesasa.calc(st).residueAreas().get(lab, {})

        out = []
        for t in by_entry[pdb]:
            ch, acc = t["chain"], t["acc"]
            if ch not in atoms: continue
            p = prot[acc]; useq = seqs.get(acc)
            if not useq: continue
            ms = p["mature_start"]; expect = useq[ms-1]
            # candidate author number from SIFTS
            cands = []
            for sp_beg, pdb_beg in segs.get((pdb, ch, acc), []):
                if sp_beg > ms: continue
                m = re.match(r"^(-?\d+)", str(pdb_beg))
                if m: cands.append(int(m.group(1)) + (ms - sp_beg))
            resmap = {}
            for num, ic, resn, an, xyz in atoms[ch]:
                if ic == " ": resmap.setdefault(num, resn)
            hit = None
            for c in cands:
                for off in (0, 1, -1, 2, -2):
                    n = c + off
                    if resmap.get(n) and AA3.get(resmap[n]) == expect:
                        hit = n; break
                if hit is not None: break
            if hit is None: continue
            # is it the first observed residue of the chain?
            first_obs = min(resmap)
            key = str(hit)
            ra_c = cx.get(ch, {}).get(key); ra_i = iso.get(ch, {}).get(key)
            if ra_c is None or ra_i is None or ra_i.total <= 0: continue
            buried = 100.0 * (1 - ra_c.total / ra_i.total)
            # distances from this residue (and its alpha-amino N) to other chains
            resatoms = [a for a in atoms[ch] if a[0] == hit and a[1] == " "]
            other = allxyz[chain_lab != ch]
            if len(other) == 0: continue
            tree = cKDTree(other)
            d_res = float(tree.query(np.array([a[4] for a in resatoms]))[0].min())
            nat = [a for a in resatoms if a[3] == "N"]
            d_N = float(tree.query(np.array([nat[0][4]]))[0][0]) if nat else np.nan
            out.append(dict(pdb=pdb, chain=ch, acc=acc, gene=p["gene"],
                            n_chains=len(atoms), n_entities=t["n_entities"],
                            mature_start=ms, init_removed=p["init_removed"],
                            res1=p["res1"], res2=p["res2"], res3=p["res3"],
                            nt_acetyl=p["nt_acetyl"], has_signal=p["has_signal"],
                            has_propep=p["has_propep"], has_transit=p["has_transit"],
                            resname=resmap[hit], authnum=hit,
                            is_first_observed=(hit == first_obs),
                            sasa_complex=round(ra_c.total, 1), sasa_isolated=round(ra_i.total, 1),
                            buried_pct=round(buried, 1), min_dist_residue=round(d_res, 2),
                            min_dist_alphaamino_N=round(d_N, 2) if d_N == d_N else ""))
        return out
    except Exception as e:
        return []
    finally:
        if os.path.exists(tmp): os.remove(tmp)

KEYS = ['pdb','chain','acc','gene','n_chains','n_entities','mature_start','init_removed','res1','res2',
        'res3','nt_acetyl','has_signal','has_propep','has_transit','resname','authnum','is_first_observed',
        'sasa_complex','sasa_isolated','buried_pct','min_dist_residue','min_dist_alphaamino_N']
OUT = f"{WORK_DIR}/nterm_geometry.csv"
DONE = f"{WORK_DIR}/done_entries.txt"

if __name__ == "__main__":
    budget = int(sys.argv[1]) if len(sys.argv) > 1 else 250
    done = set(open(DONE).read().split()) if os.path.exists(DONE) else set()
    entries = [e for e in sorted(by_entry) if os.path.exists(f"{WORK_DIR}/pdb/{e}.pdb")
               and e not in done]
    new = entries[:budget]
    fresh = not os.path.exists(OUT)
    fh = open(OUT, "a", newline=""); w = csv.DictWriter(fh, fieldnames=KEYS)
    if fresh: w.writeheader()
    dfh = open(DONE, "a")
    n = 0
    for i, pdb in enumerate(new):
        rows = analyze(pdb)
        w.writerows(rows); n += len(rows)
        dfh.write(pdb + "\n")
        if i % 25 == 0:
            fh.flush(); dfh.flush()
            print(i, "/", len(new), "rows this run:", n, flush=True)
    fh.close(); dfh.close()
    print("BATCH DONE entries:", len(new), "rows:", n, "| remaining:", len(entries) - len(new))
```
