## Supplementary_Data_8a for "Human protein complex interfaces reveal candidate assembly roles for initiator methionine excision": step4_stats.py

```
"""RETIRED PILOT SCRIPT - not on the path to any reported number.

Kept for completeness only. Its per-protein collapse predates the v1.2.1/v1.2.2
provenance alignment, so the distance fields it prints are protein-level minima
that are not tied to the PDB entry printed beside them. Use step6_stats.py.
"""
import os as _os
BASE = _os.environ.get("NME_DIR") or _os.path.dirname(_os.path.dirname(_os.path.abspath(__file__)))
INPUT_DIR = _os.path.join(BASE, "inputs")
WORK_DIR = _os.environ.get("NME_WORK") or _os.path.join(BASE, "work")
_os.makedirs(WORK_DIR, exist_ok=True)
import csv, collections, math, json
import numpy as np
from scipy import stats

SMALL = set("AGSTCPV")            # MetAP-permissive P1'
PHI   = set("LFYWIVA")            # hydrophobic Phi -> M-Phi/N-degron on retention

rows = list(csv.DictReader(open(f"{WORK_DIR}/nterm_geometry.csv")))
for r in rows:
    r["buried_pct"] = float(r["buried_pct"])
    r["min_dist_residue"] = float(r["min_dist_residue"])
    r["dN"] = float(r["min_dist_alphaamino_N"]) if r["min_dist_alphaamino_N"] else np.nan
    r["init_removed"] = r["init_removed"] == "True"
    r["n_chains"] = int(r["n_chains"]); r["n_entities"] = int(r["n_entities"])
    r["other_proteolysis"] = any(r[k] == "True" for k in ("has_signal","has_propep","has_transit"))
print("observations:", len(rows), "| distinct proteins:", len({r['acc'] for r in rows}))

# ---- collapse to one record per protein: most interface-engaged observation ----
best = {}
for r in rows:
    k = r["acc"]
    if k not in best or r["buried_pct"] > best[k]["buried_pct"]:
        best[k] = r
prot = list(best.values())
# minimum alpha-amino N distance across all structures of that protein
mind = collections.defaultdict(lambda: np.inf)
for r in rows:
    if r["dN"] == r["dN"]: mind[r["acc"]] = min(mind[r["acc"]], r["dN"])
for p in prot: p["dN_min"] = mind[p["acc"]]

# ---- classification ----
def classify(p):
    if p["other_proteolysis"]:
        return "excluded: signal/propeptide/transit"
    if p["init_removed"]:
        return "NME-generated (annotated)"
    if p["res1"] == "M" and p["resname"] == "MET":
        return "Met-retained (annotated/observed)"
    return "unclassified"
for p in prot:
    p["cls"] = classify(p)
    p["pred"] = "predicted NME" if p["res2"] in SMALL else "predicted Met-retained"

print("\n=== class counts (one record per protein) ===")
for k, v in collections.Counter(p["cls"] for p in prot).most_common(): print(f"  {k}: {v}")

def summarize(group, label):
    b = np.array([p["buried_pct"] for p in group])
    d = np.array([p["dN_min"] for p in group if np.isfinite(p["dN_min"])])
    return dict(label=label, n=len(group),
                median_buried=round(float(np.median(b)), 1),
                mean_buried=round(float(b.mean()), 1),
                frac_buried_ge20=round(float((b >= 20).mean()), 3),
                frac_buried_ge50=round(float((b >= 50).mean()), 3),
                n_dist=len(d),
                median_dN=round(float(np.median(d)), 2),
                frac_dN_le5=round(float((d <= 5).mean()), 3),
                frac_dN_le4=round(float((d <= 4).mean()), 3))

def compare(A, B, la, lb, tag):
    print(f"\n=== {tag} ===")
    sA, sB = summarize(A, la), summarize(B, lb)
    for s in (sA, sB): print("  ", s)
    a = np.array([p["buried_pct"] for p in A]); b = np.array([p["buried_pct"] for p in B])
    u = stats.mannwhitneyu(a, b, alternative="less")
    print(f"  Mann-Whitney (burial, {la} < {lb}): U={u.statistic:.0f}  p={u.pvalue:.3g}")
    ct = [[(a >= 20).sum(), (a < 20).sum()], [(b >= 20).sum(), (b < 20).sum()]]
    orr, pf = stats.fisher_exact(ct)
    print(f"  Fisher (buried>=20%): OR={orr:.3f}  p={pf:.3g}   table={ct}")
    da = np.array([p['dN_min'] for p in A if np.isfinite(p['dN_min'])])
    db = np.array([p['dN_min'] for p in B if np.isfinite(p['dN_min'])])
    ct2 = [[(da <= 5).sum(), (da > 5).sum()], [(db <= 5).sum(), (db > 5).sum()]]
    or2, p2 = stats.fisher_exact(ct2)
    print(f"  Fisher (alpha-amino N within 5 A of another subunit): OR={or2:.3f}  p={p2:.3g}  table={ct2}")
    u2 = stats.mannwhitneyu(da, db, alternative="greater")
    print(f"  Mann-Whitney (dN, {la} > {lb}): p={u2.pvalue:.3g}")
    return sA, sB

out = {}
A = [p for p in prot if p["cls"] == "NME-generated (annotated)"]
B = [p for p in prot if p["cls"] == "Met-retained (annotated/observed)"]
out["annotated"] = compare(A, B, "NME-generated", "Met-retained",
    "TEST 1  annotation-based: are NME-generated N-termini depleted at subunit interfaces?")

# rule-based (larger N, independent of annotation completeness)
keep = [p for p in prot if not p["other_proteolysis"]]
A2 = [p for p in keep if p["pred"] == "predicted NME"]
B2 = [p for p in keep if p["pred"] == "predicted Met-retained"]
out["rulebased"] = compare(A2, B2, "predicted NME", "predicted Met-retained",
    "TEST 2  sequence-rule-based (P1' small vs large)")

# large machines only
big = [p for p in keep if p["n_chains"] >= 6]
A3 = [p for p in big if p["pred"] == "predicted NME"]
B3 = [p for p in big if p["pred"] == "predicted Met-retained"]
if len(A3) > 10 and len(B3) > 10:
    out["large"] = compare(A3, B3, "predicted NME", "predicted Met-retained",
        "TEST 3  restricted to large assemblies (>=6 chains)")

# degron-forming subset: Met-Ala termini that are normally cleaved
print("\n=== TEST 4  degron-forming potential among NME-dependent termini ===")
nme = [p for p in keep if p["pred"] == "predicted NME"]
phi = [p for p in nme if p["res2"] in PHI]
non = [p for p in nme if p["res2"] not in PHI]
print(f"  NME-dependent termini whose retention would create an MPhi/N-degron (P1'=Ala/Val): {len(phi)}")
print(f"  NME-dependent termini with no degron on retention (P1'=Ser/Gly/Thr/Cys/Pro): {len(non)}")
print("  ", summarize(phi, "degron-forming"))
print("  ", summarize(non, "non-degron-forming"))

# per-P1' residue breakdown
print("\n=== N-terminal burial by penultimate (P1') residue ===")
print(f"  {'P1':>3} {'n':>5} {'med.buried%':>12} {'%buried>=20':>12} {'med dN(A)':>10}")
for aa in "AGSTCPVMDELIFKNQRYWH":
    g = [p for p in keep if p["res2"] == aa]
    if len(g) < 8: continue
    s = summarize(g, aa)
    print(f"  {aa:>3} {s['n']:>5} {s['median_buried']:>12} {s['frac_buried_ge20']:>12} {s['median_dN']:>10}")

json.dump({k: v for k, v in out.items()}, open(f"{WORK_DIR}/stats_summary.json", "w"), default=str)

# per-protein master table
KEYS = ["acc","gene","cls","pred","res1","res2","res3","nt_acetyl","init_removed","resname",
        "pdb","chain","n_chains","n_entities","buried_pct","dN_min","min_dist_residue",
        "sasa_complex","sasa_isolated","is_first_observed"]
with open(f"{WORK_DIR}/per_protein_master.csv","w",newline="") as fh:
    w = csv.DictWriter(fh, fieldnames=KEYS, extrasaction="ignore"); w.writeheader()
    for p in sorted(prot, key=lambda x: -x["buried_pct"]): w.writerow(p)
print("\nwrote per_protein_master.csv")

# ---- TEST 5: the at-risk list ----
risk = [p for p in keep if p["pred"] == "predicted NME" and p["buried_pct"] >= 25]
risk.sort(key=lambda p: -p["buried_pct"])
print(f"\n=== TEST 5  NME-dependent N-termini buried >=25% on assembly: {len(risk)} proteins ===")
print(f"  {'gene':>10} {'P1':>3} {'buried%':>8} {'dN(A)':>7} {'chains':>7}  degron?  pdb")
for p in risk[:45]:
    print(f"  {str(p['gene'])[:10]:>10} {p['res2']:>3} {p['buried_pct']:>8} {p['dN_min']:>7.2f} "
          f"{p['n_chains']:>7}  {'YES' if p['res2'] in PHI else '-  '}     {p['pdb']}")
with open(f"{WORK_DIR}/at_risk_termini.csv","w",newline="") as fh:
    w = csv.DictWriter(fh, fieldnames=KEYS, extrasaction="ignore"); w.writeheader(); w.writerows(risk)
print(f"\n  fraction of NME-dependent termini that are interface-engaged (>=25% buried): "
      f"{len(risk)/len([p for p in keep if p['pred']=='predicted NME']):.3f}")

# ---- figure ----
import matplotlib
matplotlib.use("Agg")
import matplotlib.pyplot as plt
fig, ax = plt.subplots(1, 2, figsize=(11, 4.4))
groups = [("predicted NME", A2, "#2c7fb8"), ("predicted Met-retained", B2, "#d95f0e")]
for i, (lab, g, c) in enumerate(groups):
    v = np.array([p["buried_pct"] for p in g])
    x = np.random.normal(i, 0.07, len(v))
    ax[0].scatter(x, v, s=9, alpha=0.45, color=c, edgecolors="none")
    ax[0].hlines(np.median(v), i-0.25, i+0.25, color="k", lw=2)
    ax[0].hlines(v.mean(), i-0.18, i+0.18, color="k", lw=1, linestyle=":")
ax[0].set_xticks([0, 1]); ax[0].set_xticklabels([f"{g[0]}\n(n={len(g[1])})" for g in groups], fontsize=9)
ax[0].set_ylabel("N-terminal residue buried on assembly (%)")
ax[0].set_title("Interface engagement of the mature N-terminus", fontsize=10)
ax[0].axhline(25, color="grey", lw=0.6, ls="--")
HILITE = {"PSMA", "PSMB", "PSMC", "PSMD", "PSME", "PSMF", "PSMG", "KPNA", "KPNB", "NUP", "RAN", "TNPO"}
for lab, g, c in groups:
    gg = [p for p in g if np.isfinite(p["dN_min"])]
    b = np.array([p["buried_pct"] for p in gg])
    d = np.clip(np.array([p["dN_min"] for p in gg]), 0, 20)
    ax[1].scatter(d, b, s=11, alpha=0.5, color=c, label=f"{lab} (n={len(gg)})", edgecolors="none")
    for p, xx, yy in zip(gg, d, b):
        gname = str(p["gene"] or "")
        if any(gname.startswith(h) for h in HILITE) and yy > 15:
            ax[1].scatter([xx], [yy], s=46, facecolors="none", edgecolors="k", lw=0.8)
            ax[1].annotate(gname.split(";")[0], (xx, yy), fontsize=6.5,
                           xytext=(4, 3), textcoords="offset points")
ax[1].scatter([], [], s=46, facecolors="none", edgecolors="k", lw=0.8,
              label="proteasome / nuclear-transport subunit")
ax[1].set_xlabel(u"min distance, \u03b1-amino N to another subunit (\u00c5; clipped at 20)")
ax[1].set_ylabel("buried on assembly (%)")
ax[1].axvline(5, color="grey", lw=0.6, ls="--")
ax[1].legend(fontsize=8, frameon=False)
ax[1].set_title("Steric context of residue 1", fontsize=10)
plt.tight_layout(); plt.savefig(f"{WORK_DIR}/figure_nterm_interface.png", dpi=220)
print("wrote figure_nterm_interface.png")
```
