## Supplementary_Data_8a for "Human protein complex interfaces reveal candidate assembly roles for initiator methionine excision": step5_full.py

```
import os as _os
BASE = _os.environ.get("NME_DIR") or _os.path.dirname(_os.path.dirname(_os.path.abspath(__file__)))
INPUT_DIR = _os.path.join(BASE, "inputs")
WORK_DIR = _os.environ.get("NME_WORK") or _os.path.join(BASE, "work")
_os.makedirs(WORK_DIR, exist_ok=True)
"""Full-coverage rebuild on biological assemblies.

Task list: every human accession that has at least one multi-chain PDB entry.
Entries are ranked per accession (most distinct entities, then most chains) and
the global task list is interleaved by rank, so rank-1 entries for all 4,087
accessions are processed first and redundancy is added with remaining time.
Resumable: already-processed entries are read from the status log.
"""
import gzip, csv, json, os, sys, time, collections, multiprocessing as mp
sys.path.insert(0, _os.path.dirname(_os.path.abspath(__file__)))
import asm_lib as L

OUT = f"{WORK_DIR}/asm_rows.jsonl"
LOG = f"{WORK_DIR}/asm_status.tsv"
TIME_BUDGET = float(os.environ.get("BUDGET_MIN", 240)) * 60
NPROC = int(os.environ.get("NPROC", 6))

D = json.load(open(f"{WORK_DIR}/targets.json"))
prot = D["prot"]
targets = [t for t in D["targets"] if t["n_chains"] >= 2]

by_entry = collections.defaultdict(list)
for t in targets:
    by_entry[t["pdb"]].append(t)

# ---- rank entries per accession -------------------------------------------
ent_meta = {}
for pdb, ts in by_entry.items():
    ent_meta[pdb] = (ts[0].get("n_entities") or 0, ts[0]["n_chains"])
acc_entries = collections.defaultdict(set)
for t in targets:
    acc_entries[t["acc"]].add(t["pdb"])

ranked = collections.defaultdict(list)   # rank -> [pdb]
seen = set()
for acc, pdbs in acc_entries.items():
    order = sorted(pdbs, key=lambda p: (-int(ent_meta[p][0] or 0), -ent_meta[p][1], p))
    for i, p in enumerate(order):
        ranked[i].append(p)
task_list = []
for i in sorted(ranked):
    for p in ranked[i]:
        if p not in seen:
            seen.add(p)
            task_list.append(p)

done = set()
if os.path.exists(LOG):
    for ln in open(LOG):
        done.add(ln.split("\t")[0])
task_list = [p for p in task_list if p not in done]
print(f"accessions={len(acc_entries)} entries_total={len(by_entry)} queued={len(task_list)}", flush=True)

# ---- sequences and SIFTS segments -----------------------------------------
need = {t["acc"] for t in targets}
seqs = {}
with gzip.open(f"{INPUT_DIR}/human_nterm.tsv.gz", "rt") as fh:
    for r in csv.DictReader(fh, delimiter="\t"):
        if r["Entry"] in need:
            seqs[r["Entry"]] = r["Sequence"]
segs = collections.defaultdict(list)
with gzip.open(f"{INPUT_DIR}/pdb_chain_uniprot.tsv.gz", "rt") as fh:
    fh.readline()
    for r in csv.DictReader(fh, delimiter="\t"):
        pdb = r["PDB"].lower()
        if pdb not in by_entry:
            continue
        try:
            segs[(pdb, r["CHAIN"], r["SP_PRIMARY"])].append((int(r["SP_BEG"]), r["PDB_BEG"]))
        except ValueError:
            pass
segs_by_pdb = collections.defaultdict(dict)
for k, v in segs.items():
    segs_by_pdb[k[0]][k] = v
print(f"seqs={len(seqs)} sift_segments={len(segs)}", flush=True)

G = {}


def init():
    G["ok"] = True


def work(pdb):
    tl = by_entry[pdb]
    sub = segs_by_pdb.get(pdb, {})
    try:
        rows, status = L.analyze_entry(pdb, tl, prot, seqs, sub)
    except Exception as e:
        rows, status = [], f"fatal:{type(e).__name__}"
    return pdb, rows, status


if __name__ == "__main__":
    t0 = time.time()
    n = 0
    counts = collections.Counter()
    with open(OUT, "a") as fo, open(LOG, "a") as fl:
        with mp.Pool(NPROC, initializer=init, maxtasksperchild=25) as pool:
            for pdb, rows, status in pool.imap_unordered(work, task_list, chunksize=1):
                n += 1
                counts[status] += 1
                fl.write(f"{pdb}\t{status}\t{len(rows)}\n")
                for r in rows:
                    fo.write(json.dumps(r, default=float) + "\n")
                if n % 100 == 0:
                    fo.flush(); fl.flush()
                    el = time.time() - t0
                    print(f"{n}/{len(task_list)} {el/60:.1f} min "
                          f"({n/el*60:.0f} entries/min) {dict(counts)}", flush=True)
                if time.time() - t0 > TIME_BUDGET:
                    print("time budget reached", flush=True)
                    pool.terminate()
                    break
    print("DONE", n, dict(counts), flush=True)
```
