## Supplementary_Data_8a for "Human protein complex interfaces reveal candidate assembly roles for initiator methionine excision": step6_stats.py

```
import os as _os
BASE = _os.environ.get("NME_DIR") or _os.path.dirname(_os.path.dirname(_os.path.abspath(__file__)))
INPUT_DIR = _os.path.join(BASE, "inputs")
WORK_DIR = _os.environ.get("NME_WORK") or _os.path.join(BASE, "work")
_os.makedirs(WORK_DIR, exist_ok=True)
"""Statistics for the biological-assembly, full-coverage dataset."""
import json, csv, collections, os
import numpy as np
from scipy import stats

SMALL = set("AGSTCPV")
PHI = set("LFYWIVA")
ROWS = f"{WORK_DIR}/asm_rows.jsonl"
OUTDIR = f"{WORK_DIR}"

seen = set()
rows = []
with open(ROWS) as fh:
    for ln in fh:
        r = json.loads(ln)
        k = (r["pdb"], r["asm_chain"], r["acc"])
        if k in seen:
            continue
        seen.add(k)
        r["dN"] = float(r["min_dist_alphaamino_N"]) if r["min_dist_alphaamino_N"] not in ("", None) else np.nan
        r["other_proteolysis"] = bool(r["has_signal"] or r["has_propep"] or r["has_transit"])
        rows.append(r)

entries = {r["pdb"] for r in rows}
accs = {r["acc"] for r in rows}
print(f"observations (assembly chains): {len(rows)}")
print(f"distinct PDB entries: {len(entries)}")
print(f"distinct human proteins: {len(accs)}")
print(f"assembly chain counts: median {np.median([r['n_chains'] for r in rows]):.0f}, "
      f"max {max(r['n_chains'] for r in rows)}")

# ---------------- per-protein collapse, three rules ------------------------
by_acc = collections.defaultdict(list)
for r in rows:
    by_acc[r["acc"]].append(r)


def collapse(rule):
    out = []
    for acc, rs in by_acc.items():
        if rule == "max":
            p = dict(max(rs, key=lambda x: x["buried_pct"]))
        elif rule == "median":
            p = dict(max(rs, key=lambda x: x["buried_pct"]))
            p["buried_pct"] = float(np.median([x["buried_pct"] for x in rs]))
        elif rule == "largest":
            p = dict(max(rs, key=lambda x: (x["n_chains"], x["buried_pct"])))
        p["n_obs"] = len(rs)
        p["n_entries"] = len({x["pdb"] for x in rs})
        d = [x["dN"] for x in rs if x["dN"] == x["dN"]]
        # dN_min and dN_median are protein-level statistics over every observation
        # of this protein; they are not tied to the entry named in p["pdb"].
        p["dN_min"] = float(min(d)) if d else np.inf
        p["dN_median"] = float(np.median(d)) if d else np.inf
        # dN_at_max_burial is aligned to the single observation that supplies
        # pdb, asm_chain and buried_pct, and is the value quoted in the text.
        sel = max(rs, key=lambda x: x["buried_pct"])
        p["dN_at_max_burial"] = float(sel["dN"]) if sel["dN"] == sel["dN"] else ""
        p["buried_max"] = float(max(x["buried_pct"] for x in rs))
        p["buried_median"] = float(np.median([x["buried_pct"] for x in rs]))
        out.append(p)
    return out


def classify(p):
    if p["other_proteolysis"]:
        return "excluded: signal/propeptide/transit"
    if p["init_removed"]:
        return "NME-generated (annotated)"
    if p["res1"] == "M" and p["resname"] in ("MET", "MSE"):
        return "Met-retained (annotated/observed)"
    return "unclassified"


def summarize(group, label, field="buried_pct", dfield="dN_min"):
    b = np.array([p[field] for p in group], dtype=float)
    d = np.array([p[dfield] for p in group if np.isfinite(p[dfield])], dtype=float)
    return dict(label=label, n=len(group),
                median_buried=round(float(np.median(b)), 1),
                mean_buried=round(float(b.mean()), 1),
                frac_ge20=round(float((b >= 20).mean()), 3),
                frac_ge25=round(float((b >= 25).mean()), 3),
                frac_ge50=round(float((b >= 50).mean()), 3),
                n_dist=len(d),
                median_dN=round(float(np.median(d)), 2) if len(d) else None,
                frac_dN_le5=round(float((d <= 5).mean()), 3) if len(d) else None)


def compare(A, B, la, lb, tag, field="buried_pct", dfield="dN_min"):
    print(f"\n=== {tag} ===")
    sA, sB = summarize(A, la, field, dfield), summarize(B, lb, field, dfield)
    for s in (sA, sB):
        print("  ", s)
    a = np.array([p[field] for p in A], dtype=float)
    b = np.array([p[field] for p in B], dtype=float)
    u = stats.mannwhitneyu(a, b, alternative="less")
    print(f"  Mann-Whitney (burial, {la} < {lb}): U={u.statistic:.0f} p={u.pvalue:.4g}")
    u2s = stats.mannwhitneyu(a, b, alternative="two-sided")
    print(f"  Mann-Whitney two-sided p={u2s.pvalue:.4g}")
    ct = [[int((a >= 20).sum()), int((a < 20).sum())], [int((b >= 20).sum()), int((b < 20).sum())]]
    orr, pf = stats.fisher_exact(ct)
    print(f"  Fisher (>=20% buried): OR={orr:.3f} p={pf:.4g} table={ct}")
    ct25 = [[int((a >= 25).sum()), int((a < 25).sum())], [int((b >= 25).sum()), int((b < 25).sum())]]
    or25, p25 = stats.fisher_exact(ct25)
    print(f"  Fisher (>=25% buried): OR={or25:.3f} p={p25:.4g} table={ct25}")
    da = np.array([p[dfield] for p in A if np.isfinite(p[dfield])], dtype=float)
    db = np.array([p[dfield] for p in B if np.isfinite(p[dfield])], dtype=float)
    ct2 = [[int((da <= 5).sum()), int((da > 5).sum())], [int((db <= 5).sum()), int((db > 5).sum())]]
    or2, p2 = stats.fisher_exact(ct2)
    print(f"  Fisher (alpha-amino N within 5 A): OR={or2:.3f} p={p2:.4g} table={ct2}")
    print(f"  Mann-Whitney (dN, {la} > {lb}): p={stats.mannwhitneyu(da, db, alternative='greater').pvalue:.4g}")
    return dict(A=sA, B=sB, mw_p_less=float(u.pvalue), mw_p_two=float(u2s.pvalue),
                fisher20_or=float(orr), fisher20_p=float(pf),
                fisher25_or=float(or25), fisher25_p=float(p25),
                fisher_dN_or=float(or2), fisher_dN_p=float(p2), table20=ct, table25=ct25)


out = {}
for rule in ("max", "median", "largest"):
    prot = collapse(rule)
    for p in prot:
        p["cls"] = classify(p)
        p["pred"] = "predicted NME" if p["res2"] in SMALL else "predicted Met-retained"
    keep = [p for p in prot if not p["other_proteolysis"]]
    print(f"\n########## COLLAPSE RULE: {rule}  (proteins={len(prot)}, analysable={len(keep)}) ##########")
    if rule == "max":
        print("class counts:")
        for k, v in collections.Counter(p["cls"] for p in prot).most_common():
            print(f"   {k}: {v}")
    A = [p for p in prot if p["cls"] == "NME-generated (annotated)"]
    B = [p for p in prot if p["cls"] == "Met-retained (annotated/observed)"]
    out[f"{rule}_annotated"] = compare(A, B, "NME-generated", "Met-retained",
        f"TEST 1 [{rule}] annotation-based")
    A2 = [p for p in keep if p["pred"] == "predicted NME"]
    B2 = [p for p in keep if p["pred"] == "predicted Met-retained"]
    out[f"{rule}_rule"] = compare(A2, B2, "predicted NME", "predicted Met-retained",
        f"TEST 2 [{rule}] sequence-rule-based")
    if rule == "max":
        PROT = prot
        KEEP = keep
        A2M, B2M = A2, B2

prot, keep, A2, B2 = PROT, KEEP, A2M, B2M

# ---- stratification by assembly size -------------------------------------
print("\n=== TEST 3  stratified by assembly size (max-burial collapse) ===")
for lo, hi, lab in [(2, 3, "2-3 chains"), (4, 5, "4-5 chains"), (6, 11, "6-11 chains"),
                    (12, 10**6, ">=12 chains")]:
    A3 = [p for p in keep if p["pred"] == "predicted NME" and lo <= p["n_chains"] <= hi]
    B3 = [p for p in keep if p["pred"] == "predicted Met-retained" and lo <= p["n_chains"] <= hi]
    if len(A3) > 10 and len(B3) > 10:
        out[f"strat_{lab}"] = compare(A3, B3, "predicted NME", "predicted Met-retained",
            f"TEST 3 [{lab}]")

# ---- heteromeric only ----------------------------------------------------
het_keep = [p for p in keep if int(p.get("n_entities") or 0) >= 2]
A4 = [p for p in het_keep if p["pred"] == "predicted NME"]
B4 = [p for p in het_keep if p["pred"] == "predicted Met-retained"]
if len(A4) > 10 and len(B4) > 10:
    out["heteromeric"] = compare(A4, B4, "predicted NME", "predicted Met-retained",
        "TEST 3b heteromeric assemblies only (>=2 distinct entities)")

# ---- degron subset -------------------------------------------------------
print("\n=== TEST 4  degron-forming potential among NME-dependent termini ===")
nme = [p for p in keep if p["pred"] == "predicted NME"]
phi = [p for p in nme if p["res2"] in PHI]
non = [p for p in nme if p["res2"] not in PHI]
print("  degron-forming (P1' in Phi):", len(phi), "| non-degron:", len(non))
print("  ", summarize(phi, "degron-forming"))
print("  ", summarize(non, "non-degron-forming"))
out["degron"] = dict(phi=summarize(phi, "degron-forming"), non=summarize(non, "non-degron"))

# ---- P1' breakdown -------------------------------------------------------
print("\n=== burial by penultimate (P1') residue ===")
print(f"  {'P1':>3} {'n':>5} {'med%':>7} {'mean%':>7} {'>=20%':>7} {'meddN':>7}")
p1_tab = {}
for aa in "AGSTCPVMDELIFKNQRYWH":
    g = [p for p in keep if p["res2"] == aa]
    if len(g) < 8:
        continue
    s = summarize(g, aa)
    p1_tab[aa] = s
    print(f"  {aa:>3} {s['n']:>5} {s['median_buried']:>7} {s['mean_buried']:>7} "
          f"{s['frac_ge20']:>7} {str(s['median_dN']):>7}")
out["p1"] = p1_tab

# ---- at-risk list --------------------------------------------------------
risk = sorted([p for p in nme if p["buried_pct"] >= 25], key=lambda p: -p["buried_pct"])
print(f"\n=== TEST 5  NME-dependent termini buried >=25%: {len(risk)} / {len(nme)} "
      f"({len(risk)/len(nme):.3f}) ===")
for p in risk[:40]:
    print(f"  {str(p['gene'])[:12]:>12} {p['res2']:>2} {p['buried_pct']:>6.1f} "
          f"{p['dN_min']:>6.2f} {p['n_chains']:>4}  {'degron' if p['res2'] in PHI else '-'}  {p['pdb']}")

KEYS = ["acc", "gene", "cls", "pred", "res1", "res2", "res3", "nt_acetyl", "init_removed", "resname",
        "pdb", "asm_chain", "chain", "n_chains", "n_entities", "n_obs", "n_entries",
        "buried_pct", "buried_median", "dN_min", "dN_median", "min_dist_residue",
        "sasa_complex", "sasa_isolated", "is_first_observed", "dN_at_max_burial"]
with open(f"{OUTDIR}/asm_per_protein_master.csv", "w", newline="") as fh:
    w = csv.DictWriter(fh, fieldnames=KEYS, extrasaction="ignore")
    w.writeheader()
    for p in sorted(prot, key=lambda x: -x["buried_pct"]):
        w.writerow(p)
with open(f"{OUTDIR}/asm_at_risk_termini.csv", "w", newline="") as fh:
    w = csv.DictWriter(fh, fieldnames=KEYS, extrasaction="ignore")
    w.writeheader()
    w.writerows(risk)
with open(f"{OUTDIR}/asm_observations.csv", "w", newline="") as fh:
    OK = ["pdb", "asm_chain", "chain", "acc", "gene", "n_chains", "n_entities", "mature_start",
          "init_removed", "res1", "res2", "res3", "nt_acetyl", "resname", "authnum",
          "is_first_observed", "sasa_complex", "sasa_isolated", "buried_pct",
          "min_dist_residue", "min_dist_alphaamino_N", "n_atoms"]
    w = csv.DictWriter(fh, fieldnames=OK, extrasaction="ignore")
    w.writeheader()
    w.writerows(rows)
json.dump(out, open(f"{OUTDIR}/asm_stats_summary.json", "w"), default=str, indent=1)
print("\nwrote asm_per_protein_master.csv, asm_at_risk_termini.csv, asm_observations.csv, asm_stats_summary.json")
```
