## Supplementary_Data_8a for "Human protein complex interfaces reveal candidate assembly roles for initiator methionine excision": step7_targeted.py

```
import os as _os
BASE = _os.environ.get("NME_DIR") or _os.path.dirname(_os.path.dirname(_os.path.abspath(__file__)))
INPUT_DIR = _os.path.join(BASE, "inputs")
WORK_DIR = _os.environ.get("NME_WORK") or _os.path.join(BASE, "work")
_os.makedirs(WORK_DIR, exist_ok=True)
"""Targeted re-analysis of specific entries on biological assemblies (for Figs 3-4)."""
import gzip, csv, json, collections, sys, os
sys.path.insert(0, _os.path.dirname(_os.path.abspath(__file__)))
import asm_lib as L

L.MAX_ATOMS = 1500000
TEST = sys.argv[1:] or ["4r3o", "6rey", "8qyj", "9qoo", "1ibr", "5ijo", "3tj3", "5a9q", "9lwf", "1qgk"]
D = json.load(open(f"{WORK_DIR}/targets.json"))
prot = D["prot"]
sel = set(TEST)
by_entry = collections.defaultdict(list)
for t in D["targets"]:
    if t["pdb"] in sel:
        by_entry[t["pdb"]].append(t)
need = {t["acc"] for e in by_entry.values() for t in e}
seqs = {}
with gzip.open(f"{INPUT_DIR}/human_nterm.tsv.gz", "rt") as fh:
    for r in csv.DictReader(fh, delimiter="\t"):
        if r["Entry"] in need:
            seqs[r["Entry"]] = r["Sequence"]
segs = collections.defaultdict(list)
with gzip.open(f"{INPUT_DIR}/pdb_chain_uniprot.tsv.gz", "rt") as fh:
    fh.readline()
    for r in csv.DictReader(fh, delimiter="\t"):
        pdb = r["PDB"].lower()
        if pdb in sel:
            try:
                segs[(pdb, r["CHAIN"], r["SP_PRIMARY"])].append((int(r["SP_BEG"]), r["PDB_BEG"]))
            except ValueError:
                pass

allrows = []
for pdb in TEST:
    rows, status = L.analyze_entry(pdb, by_entry.get(pdb, []), prot, seqs, segs)
    print(f"== {pdb} {status} n={len(rows)}"
          f" chains={rows[0]['n_chains'] if rows else '-'} atoms={rows[0]['n_atoms'] if rows else '-'}")
    for r in sorted(rows, key=lambda x: (x["gene"], x["asm_chain"])):
        print(f"   {r['gene']:>10} ch {r['asm_chain']:>4} res1={r['res1']} P1'={r['res2']} "
              f"model={r['resname']}{r['authnum']} buried={r['buried_pct']:>5.1f}% "
              f"dN={r['min_dist_alphaamino_N']} dres={r['min_dist_residue']} "
              f"init_removed={r['init_removed']} firstobs={r['is_first_observed']}")
    allrows += rows
with open(f"{WORK_DIR}/asm_targeted.json", "w") as fh:
    json.dump(allrows, fh, default=float, indent=1)
print("wrote asm_targeted.json", len(allrows))
```
