## Supplementary_Data_8a for "Human protein complex interfaces reveal candidate assembly roles for initiator methionine excision": step8_cascade.py

```
import os as _os
BASE = _os.environ.get("NME_DIR") or _os.path.dirname(_os.path.dirname(_os.path.abspath(__file__)))
INPUT_DIR = _os.path.join(BASE, "inputs")
WORK_DIR = _os.environ.get("NME_WORK") or _os.path.join(BASE, "work")
_os.makedirs(WORK_DIR, exist_ok=True)
"""Fetch UniProt Nalpha-modification annotations (acetylation, lipidation, methylation)
and sequences for every protein in the assembly census, then test each downstream
branch of the modification cascade against interface geometry."""
import csv, json, time, urllib.request, urllib.parse, statistics as st
from scipy.stats import mannwhitneyu, fisher_exact

OUT = f"{WORK_DIR}/"
rows = [r for r in csv.DictReader(open(OUT + "asm_per_protein_master.csv"))]
accs = [r["acc"] for r in rows]
print("accessions:", len(accs))

FIELDS = "accession,sequence,ft_mod_res,ft_lipid,ft_init_met"
data = {}
for i in range(0, len(accs), 40):
    chunk = accs[i:i + 40]
    q = " OR ".join(f"accession:{a}" for a in chunk)
    url = "https://rest.uniprot.org/uniprotkb/search?" + urllib.parse.urlencode(
        {"query": q, "fields": FIELDS, "format": "json", "size": "500"})
    for attempt in range(4):
        try:
            d = json.load(urllib.request.urlopen(url, timeout=120))
            break
        except Exception as e:
            print("retry", i, e)
            time.sleep(4)
    else:
        continue
    for e in d.get("results", []):
        a = e["primaryAccession"]
        seq = e.get("sequence", {}).get("value", "")
        feats = e.get("features", [])
        def pos1(f):
            loc = f.get("location", {})
            return loc.get("start", {}).get("value")
        myr = met = ac = pal = False
        for f in feats:
            desc = (f.get("description") or "")
            if f["type"] == "Lipidation" and pos1(f) in (1, 2):
                if "yristoyl" in desc:
                    myr = True
                if "almitoyl" in desc:
                    pal = True
            if f["type"] == "Modified residue" and pos1(f) in (1, 2):
                if "ethyl" in desc:
                    met = True
                if "cetyl" in desc:
                    ac = True
        data[a] = dict(seq=seq, myr=myr, methyl=met, acetyl=ac, palm=pal)
    print("fetched", len(data), flush=True)

json.dump(data, open(OUT + "asm_uniprot_mods.json", "w"))

# ------------------------------------------------------------------ classify
PHI = set("LFYWIVA")
NME_P1 = set("AGSTCPV")
recs = []
for r in rows:
    a = r["acc"]
    if a not in data:
        continue
    d = data[a]
    seq = d["seq"]
    if r["cls"].startswith("excluded") or r["cls"].startswith("unclassified"):
        continue
    pred_nme = r["res2"] in NME_P1
    # mature sequence positions 1,2,3 = precursor 2,3,4
    m1, m2, m3 = (seq[1:2], seq[2:3], seq[3:4]) if len(seq) > 3 else ("", "", "")
    # canonical NTMT1 recognition motif on the mature terminus: [A/P/S/G]-P-K
    ntmt_motif = pred_nme and m1 in ("A", "P", "S", "G") and m2 == "P" and m3 == "K"
    # NMT recognition: mature Gly1, hydrophilic/permissive downstream, Lys at mature 6
    nmt_motif = pred_nme and m1 == "G"
    recs.append(dict(acc=a, gene=r["gene"].split(";")[0], p1=r["res2"],
                     buried=float(r["buried_pct"]),
                     dN=float(r["dN_min"]) if r["dN_min"] not in ("", "nan") else None,
                     nch=int(r["n_chains"]), pdb=r["pdb"], pred_nme=pred_nme,
                     degron=r["res2"] in PHI, myr=d["myr"], methyl=d["methyl"],
                     acetyl=d["acetyl"], palm=d["palm"],
                     ntmt_motif=ntmt_motif, nmt_motif=nmt_motif,
                     m1=m1, m2=m2, m3=m3))

json.dump(recs, open(OUT + "asm_cascade_records.json", "w"))
nme = [r for r in recs if r["pred_nme"]]
print(f"\nanalysable {len(recs)}, predicted NME {len(nme)}")


def desc(rs, lab):
    b = [r["buried"] for r in rs]
    d = [r["dN"] for r in rs if r["dN"] is not None]
    if not b:
        print(f"{lab:40s} n=0")
        return
    print(f"{lab:40s} n={len(b):4d} median={st.median(b):5.1f} mean={st.mean(b):5.1f} "
          f">=25%={100*sum(x>=25 for x in b)/len(b):5.1f}% "
          f"medianDN={st.median(d) if d else float('nan'):5.2f} "
          f"dN<=5A={100*sum(x<=5 for x in d)/len(d) if d else 0:5.1f}%")


def test(sub, rest, lab):
    b1 = [r["buried"] for r in sub]; b2 = [r["buried"] for r in rest]
    if len(b1) < 3:
        print(lab, "too few"); return
    p = mannwhitneyu(b1, b2, alternative="two-sided").pvalue
    tab = [[sum(x >= 25 for x in b1), sum(x < 25 for x in b1)],
           [sum(x >= 25 for x in b2), sum(x < 25 for x in b2)]]
    orr, pf = fisher_exact(tab)
    d1 = [r["dN"] for r in sub if r["dN"] is not None]
    d2 = [r["dN"] for r in rest if r["dN"] is not None]
    pd = mannwhitneyu(d1, d2, alternative="two-sided").pvalue if len(d1) >= 3 else float("nan")
    print(f"  {lab}: burial MW p={p:.4g} | >=25% OR={orr:.3f} p={pf:.4g} {tab} | dN MW p={pd:.4g}")


print("\n--- MYRISTOYLATION branch (mature Gly1 candidates)")
gly = [r for r in nme if r["p1"] == "G"]
desc([r for r in gly if r["myr"]], "Gly1, myristoylation annotated")
desc([r for r in gly if not r["myr"]], "Gly1, not annotated")
test([r for r in gly if r["myr"]], [r for r in gly if not r["myr"]], "within Gly1")
test([r for r in nme if r["myr"]], [r for r in nme if not r["myr"]], "myristoylated vs all other NME")
print("  genes:", sorted({r['gene'] for r in nme if r['myr']}))
for r in sorted([x for x in nme if x["myr"]], key=lambda r: -r["buried"])[:10]:
    print(f"    {r['gene']:8s} {r['buried']:5.1f}% dN={r['dN']} nch={r['nch']} {r['pdb']}")

print("\n--- METHYLATION branch")
desc([r for r in nme if r["ntmt_motif"]], "NTMT motif [A/P/S/G]-P-K")
desc([r for r in nme if not r["ntmt_motif"]], "other predicted NME")
test([r for r in nme if r["ntmt_motif"]], [r for r in nme if not r["ntmt_motif"]], "NTMT motif")
print("  motif genes:", [(r['gene'], r['m1'] + r['m2'] + r['m3'], round(r['buried'], 1),
                          r['dN'], r['nch'], r['pdb']) for r in
                         sorted([x for x in nme if x['ntmt_motif']], key=lambda r: -r['buried'])])
ann = [r for r in recs if r["methyl"]]
desc(ann, "annotated Nalpha-methylation")
print("  annotated-methyl genes:", [(r['gene'], r['p1'], round(r['buried'], 1), r['nch']) for r in
                                    sorted(ann, key=lambda r: -r['buried'])][:25])

print("\n--- PALMITOYLATION / ACETYLATION context")
desc([r for r in nme if r["palm"]], "palmitoylation annotated")
desc([r for r in nme if r["acetyl"]], "Nt-acetylation annotated (NME set)")
desc([r for r in nme if not r["acetyl"]], "no acetylation annotation (NME set)")
test([r for r in nme if r["acetyl"]], [r for r in nme if not r["acetyl"]], "acetylation annotated")

print("\n--- combined cascade classes among predicted-NME termini")
myr = [r for r in nme if r["myr"]]
mot = [r for r in nme if r["ntmt_motif"] and not r["myr"]]
oth = [r for r in nme if not r["myr"] and not r["ntmt_motif"]]
desc(myr, "lipid-anchored (myristoylated)")
desc(mot, "methylation-motif")
desc(oth, "remainder")
```
