## Supplementary_Data_8a for "Human protein complex interfaces reveal candidate assembly roles for initiator methionine excision": validate_and_summarize.py

```
"""Independent bookkeeping checks and supplementary evidence flags."""
from pathlib import Path
import hashlib, json, re
import pandas as pd, numpy as np

R=Path(__file__).resolve().parent
B=R/"frozen/nterm-census-1.3.0"
O=R/"results"
p=pd.read_csv(B/"data/asm_per_protein_master.csv")
o=pd.read_csv(B/"data/asm_observations.csv")
u=pd.read_csv(B/"inputs/human_nterm.tsv.gz",sep="\t").fillna("")
x=p.merge(u,left_on="acc",right_on="Entry",validate="one_to_one")
x=x[~x.cls.str.startswith("excluded")].copy()
rows=[]
for r in x.to_dict("records"):
    starts=[int(v) for v in re.findall(r"CHAIN (\d+)\.\.",r["Chain"])]
    rows.append(dict(acc=r["acc"],gene=r["gene"],classification=r["cls"],
        frozen_init_annotation=r["Initiator methionine"],
        terminal_acetylation_export=r["nt_acetyl"],
        alternate_removal_annotation="alternate" in r["Initiator methionine"],
        modeled_start_used=2 if r["init_removed"] else 1,
        annotated_chain_starts=";".join(map(str,starts)),
        multiple_chain_products_flag=any(v>2 for v in starts),
        chain_annotation=r["Chain"]))
ann=pd.DataFrame(rows)
ann.to_csv(O/"protein_processing_evidence_flags.csv",index=False)
checks=[]
def check(name,condition,detail):
    checks.append(dict(check=name,passed=bool(condition),detail=detail))

mx=o.groupby("acc").buried_pct.max()
check("master_maxima_equal_observation_maxima",
      np.allclose(p.set_index("acc").buried_pct.sort_index(),mx.sort_index()),
      "All 1,191 protein maxima checked against original observations.")
check("observation_keys_unique",not o.duplicated(["acc","pdb","asm_chain"]).any(),
      "Unique accession/PDB/assembly-chain keys.")
check("master_observation_counts",p.set_index("acc").n_obs.sort_index().equals(
    o.groupby("acc").size().sort_index().rename("n_obs")),
    "Per-protein observation counts.")
check("parser_INIT_assignments_match_frozen",
      (x.init_removed == x["Initiator methionine"].str.contains("Removed")).all(),
      "No mismatch between frozen feature presence and stored init_removed.")
geo=pd.read_csv(O/"focal_recomputed_geometry.csv")
matched=geo.dropna(subset=["original_census_buried_pct"])
check("focal_burial_reproduction",np.allclose(matched.buried_pct.round(1),
      matched.original_census_buried_pct),f"{len(matched)} previously exported measurements.")
check("focal_N_contact_reproduction",np.allclose(matched.target_N_contact_A.round(2),
      matched.original_census_N_contact_A),f"{len(matched)} previously exported measurements.")
sel=pd.read_csv(O/"figure5_residue_selection_audit.csv")
check("all_53_scan_targets_audited",len(sel)==53,"Identity match alone is not full residue-mapping validation.")
paths=B/"inputs/figure5_assemblies"
for line in (paths/"SHA256SUMS.txt").read_text().splitlines():
    expected,path=line.split()
    f=paths/Path(path).name
    check("frozen_coordinate_hash_"+f.name,
          hashlib.sha256(f.read_bytes()).hexdigest()==expected,"Compared with release manifest.")
submission=R.parent/"paper1_v8/pkg_v7"
for f in sorted(submission.glob("*Supplementary_Data*.xlsx")):
    name=f.name.split("_",1)[1]
    other=B/"data"/name
    if not other.exists():
        other=B/"data"/name.replace("Robustness_and","Robustness_analyses_and")
    check("submitted_workbook_hash_"+f.name,
          other.exists() and hashlib.sha256(f.read_bytes()).hexdigest()==hashlib.sha256(other.read_bytes()).hexdigest(),
          "Byte-identical submitted and frozen-release workbook; only filename may differ.")
pd.DataFrame(checks).to_csv(O/"validation_checks.csv",index=False)
summary=dict(retained_INIT_blank=int(((ann.classification.str.startswith("Met-retained"))&
    ann.frozen_init_annotation.eq("")).sum()),
    retained_with_N_acetylmethionine=int(((ann.classification.str.startswith("Met-retained"))&
    ann.terminal_acetylation_export.eq("N-acetylmethionine")).sum()),
    alternate_removal_count=int(ann.alternate_removal_annotation.sum()),
    multiple_chain_product_count=int(ann.multiple_chain_products_flag.sum()),
    validation_checks=len(checks),validation_passes=sum(c["passed"] for c in checks))
(O/"validation_summary.json").write_text(json.dumps(summary,indent=2)+"\n")
print(pd.DataFrame(checks).to_string(index=False))
print(json.dumps(summary,indent=2))
```
