## Supplementary_Data_8a for "Human protein complex interfaces reveal candidate assembly roles for initiator methionine excision": README.md

# Current figure presentation

This folder reproduces the 25 September 2026 figure presentation for “Human protein complex interfaces reveal candidate assembly roles for initiator methionine excision.” It changes figure typography and layout, not the underlying structural calculations or analytical cohorts.

Run `python rebuild_artwork.py` from this folder. The script detects the included `inputs` directory and writes the five main figures and three supplementary figures to `artwork` as editable SVG/PDF, 300-dpi PNG and losslessly compressed 600-dpi TIFF.

## Input provenance

- **Census tables:** Frozen `asm_per_protein_master.csv` and `asm_observations.csv` from release 1.3.0. The code retains the established 1,123-protein eligibility filter and original annotation-selected positional measurements.
- **Focal geometry:** Previously computed unchanged-model measurements. Figure 3 retains exactly 22 first-observed Met1/Ser2 observations for PSMA4, PSMA6 and PSMA7; no coordinate modeling is performed by this presentation script.
- **Partner tables:** Previously computed measurements from Supplementary Data 7. Figure 4 is checked against its supplied pair-burial matrix, and Supplementary Figure 3 is the identical image.
- **Supplementary source tables:** Previously computed corpus-sampling and annotation/motif values. No permutation or simulation is rerun.

The required environment is Python with NumPy, pandas, SciPy, Matplotlib and Pillow. The presentation was rendered with Matplotlib 3.11.2; the other analysis environment versions are recorded in the manuscript and existing supplementary provenance.

## Display conventions

All figures use a 6.5-inch canvas width, 9-point DejaVu Sans lettering, white backgrounds, and at least one-point plot lines. Panel letters are lowercase, and short panel headings identify the question or measurement rather than assert a causal conclusion.

The main manuscript legends define all denominators and evidence qualifications. The journal-facing TIFFs and Supplementary Information contain the current renderings; pre-existing image files elsewhere in the frozen archives are retained only as earlier presentation provenance.
