## Supplementary_Data_8a for "Human protein complex interfaces reveal candidate assembly roles for initiator methionine excision": rebuild_artwork.py

```
"""Reformat verified figure source values without changing analytical selection."""

from pathlib import Path
import json
import re
import shutil
import numpy as np
import pandas as pd
import matplotlib
matplotlib.use("Agg")
import matplotlib.pyplot as plt
from matplotlib.patches import Patch
from matplotlib.colors import LinearSegmentedColormap
from matplotlib.ticker import NullLocator, NullFormatter
from scipy.stats.contingency import odds_ratio
from PIL import Image

ROOT = Path(__file__).resolve().parent
WORKSPACE = ROOT.parents[1]
OUT = ROOT / "artwork"
OUT.mkdir(exist_ok=True)
INPUTS = ROOT / "inputs"
DATA = INPUTS / "census" if INPUTS.exists() else WORKSPACE / "paper1_audit/frozen/nterm-census-1.3.0/data"
PARTNERS = INPUTS / "partners" if INPUTS.exists() else WORKSPACE / "deliverables/paper1_partner_resolved"
SUPP = INPUTS / "supplement" if INPUTS.exists() else WORKSPACE / "supplement_build/outputs"
FOCAL = INPUTS / "focal_recomputed_geometry.csv" if INPUTS.exists() else WORKSPACE / "paper1_audit/results/focal_recomputed_geometry.csv"
W, FS = 6.5, 9
BLUE, ORANGE, TEAL, INK = "#3b6ea5", "#c9752b", "#20808D", "#202020"
SMALL = set("AGSTCPV")
plt.rcParams.update({
    "font.family": "DejaVu Sans", "font.size": FS, "axes.titlesize": FS,
    "axes.labelsize": FS, "xtick.labelsize": FS, "ytick.labelsize": FS,
    "legend.fontsize": FS, "text.color": INK, "axes.labelcolor": INK,
    "axes.linewidth": 1, "xtick.major.width": 1, "ytick.major.width": 1,
    "grid.linewidth": 1, "lines.linewidth": 1, "svg.fonttype": "none",
    "pdf.fonttype": 42, "ps.fonttype": 42, "savefig.facecolor": "white",
})
records = []


def save(fig, name, title):
    """Keep text editable in SVG/PDF and provide losslessly compressed TIFFs."""
    for ax in fig.axes:
        for spine in ax.spines.values():
            spine.set_linewidth(1)
        for line in ax.lines:
            if line.get_linewidth() > 0:
                line.set_linewidth(max(1, line.get_linewidth()))
    fig.canvas.draw()
    fig.savefig(OUT / f"{name}.svg")
    fig.savefig(OUT / f"{name}.pdf",
                metadata={"Title": title, "Author": "Perplexity Computer"})
    fig.savefig(OUT / f"{name}.png", dpi=300)
    fig.savefig(OUT / f"{name}.tif", dpi=600,
                pil_kwargs={"compression": "tiff_lzw"})
    records.append({"figure": name, "width_inches": fig.get_figwidth(),
                    "height_inches": fig.get_figheight(), "font_points": FS,
                    "tiff_dpi": 600})
    plt.close(fig)


def clean(ax, grid=False):
    for side in ["top", "right"]:
        ax.spines[side].set_visible(False)
    if grid:
        ax.grid(color="#E5E5E5")
        ax.set_axisbelow(True)


def heading(ax, letter, text):
    ax.set_title(f"{letter}  {text}", loc="left", fontweight="bold", pad=9)


p = pd.read_csv(DATA / "asm_per_protein_master.csv")
p = p[~p.cls.str.startswith("excluded") & (p.cls != "unclassified")].copy()
p["permissive"] = p.res2.isin(SMALL)
p["finite_N"] = pd.to_numeric(p.dN_min, errors="coerce").replace(
    [np.inf, -np.inf], np.nan)
assert len(p) == 1123 and p.permissive.sum() == 696
assert ((p.buried_pct >= 25) & p.permissive).sum() == 261

# Figure 1: preserve distributions, stratum estimates and original selections.
# Full-width residue panel keeps all twenty n labels at the same 9-point size.
fig = plt.figure(figsize=(W, 8.0), layout="constrained")
gs = fig.add_gridspec(3, 2, height_ratios=[1, 1, 1.08])
a = fig.add_subplot(gs[0, :])
b = fig.add_subplot(gs[1, :])
c = fig.add_subplot(gs[2, 0])
d = fig.add_subplot(gs[2, 1])
for group, color, label in [(True, BLUE, "Permissive"), (False, ORANGE, "Nonpermissive")]:
    sub = p[p.permissive == group]
    vals = np.sort(sub.buried_pct)
    a.step(vals, np.arange(1, len(vals) + 1) / len(vals), color=color, lw=1.8,
           label=f"{label} (n = {len(vals)})")
a.axvline(25, color="#707070", ls="--")
a.set(xlim=(-2, 102), ylim=(0, 1.02),
      xlabel="Selected-position burial (%)", ylabel="Cumulative fraction")
a.legend(frameon=False, loc="lower right")
heading(a, "a", "Burial distributions by sequence class")
order = list("ACGPSTVNQDEKRHFWYILM")
rng = np.random.default_rng(0)
for i, aa in enumerate(order):
    vals = p.loc[p.res2 == aa, "buried_pct"]
    b.scatter(rng.normal(i, .13, len(vals)), vals, s=7, alpha=.36,
              color=BLUE if aa in SMALL else ORANGE, edgecolors="none")
    b.plot([i - .27, i + .27], [vals.median()] * 2, color=INK, lw=1.5)
b.set(xticks=range(20), xticklabels=[f"{aa}\n{sum(p.res2 == aa)}" for aa in order],
      xlim=(-.6, 19.6), ylim=(-4, 106), ylabel="Burial (%)",
      xlabel="Residue-2 identity, with n below")
heading(b, "b", "Burial by residue-2 identity")
strata = [(2, 3, "2–3"), (4, 5, "4–5"), (6, 11, "6–11"),
          (12, 10**6, "≥12"), (2, 10**6, "All")]
for y, (lo, hi, label) in zip(range(4, -1, -1), strata):
    sub = p[p.n_chains.between(lo, hi)]
    x, z = sub[sub.permissive], sub[~sub.permissive]
    ka, kb = int((x.buried_pct >= 25).sum()), int((z.buried_pct >= 25).sum())
    est = odds_ratio([[ka, len(x) - ka], [kb, len(z) - kb]], kind="conditional")
    ci = est.confidence_interval(.95)
    color = INK if label == "All" else BLUE
    c.plot([ci.low, ci.high], [y, y], color=color, lw=1.4)
    c.plot(est.statistic, y, "o", color=color, ms=5)
    c.text(2.05, y, f"{est.statistic:.2f}\n[{ci.low:.2f}–{ci.high:.2f}]",
           va="center", linespacing=1.05)
c.axvline(1, color="#707070", ls="--")
c.set(xscale="log", xlim=(.28, 4.5), ylim=(-.6, 4.6),
      yticks=range(4, -1, -1), yticklabels=[x[2] for x in strata],
      xlabel="Odds ratio for ≥25% burial", ylabel="Assembly chains")
c.set_xticks([.3, .5, 1, 2], [".3", ".5", "1", "2"])
c.xaxis.set_minor_locator(NullLocator())
c.xaxis.set_minor_formatter(NullFormatter())
heading(c, "c", "Assembly-size strata")
for group, color, label in [(True, BLUE, "Permissive"), (False, ORANGE, "Nonpermissive")]:
    sub = p[(p.permissive == group) & p.finite_N.notna()]
    d.scatter(np.clip(sub.finite_N, 0, 20), sub.buried_pct, s=7,
              alpha=.4, color=color, edgecolors="none")
d.axhline(25, color="#707070", ls="--")
d.axvline(5, color="#707070", ls="--")
d.add_patch(plt.Rectangle((0, 25), 5, 75, facecolor=TEAL, alpha=.07))
d.set(xlim=(0, 20.5), ylim=(-4, 104), xlabel="Minimum measured-N distance\n(Å; clipped at 20)",
      ylabel="Maximum burial (%)")
heading(d, "d", "Burial and N distance")
for ax in [a, b, c, d]:
    clean(ax)
save(fig, "Figure_1", "Sequence class and positional interface engagement")


def machine(gene):
    g = gene.split(";")[0]
    for pattern, label in [
        (r"PSM[ABCDEFG]", "Proteasome / assembly"),
        (r"CCT\d|TCP1", "CCT/TRiC"),
        (r"MED\d", "Mediator"),
        (r"SF3[AB]|SNRP|PRPF|LSM|EFTUD|SNU|SART|BUD31|CDC5L", "Spliceosome"),
        (r"COPS|CUL\d|DDB1|DDA1|RBX1|SKP1|ELOB|ELOC|FBXW", "Cullin–RING / CSN"),
        (r"NDUF|UQCR|COX\d|SDH[ABCD]|ATP5|MT-", "OXPHOS"),
        (r"RPL\d|RPS\d|RPLP|FAU|RPSA|MRPL|MRPS", "Ribosomes (cyto + mito)"),
        (r"NUP\d|SEH1L|SEC13|AAAS|RAE1|NDC1|POM121|TPR$", "Nuclear pore scaffold"),
        (r"POLR[123]", "RNA polymerases"),
        (r"EIF3|EIF2", "eIF2 / eIF3"),
        (r"ACT[BGA]|TUB[ABG]|ARPC|ACTR", "Actin / tubulin"),
        (r"ANAPC|CDC2|CDK\d|CCN[A-Z]|MCM\d|GINS", "Cell cycle / replication"),
        (r"KPN[AB]|TNPO|RAN$|RANBP|IPO\d|XPO\d", "Soluble transport receptors"),
    ]:
        if re.match(pattern, g):
            return label
    return None


# Figure 2 uses exactly the original ordered membership rule.
p["machine"] = p.gene.map(machine)
rows = []
for name, sub in p.groupby("machine", sort=False):
    if len(sub) < 4:
        continue
    q = sub[sub.permissive]
    top = sub.loc[sub.buried_pct.idxmax()]
    rows.append((name, len(sub), 100 * len(q) / len(sub),
                 100 * (q.buried_pct >= 25).mean(), top.buried_pct,
                 100 * q.res2.isin(set("LFYWIVA")).mean(), top.gene.split(";")[0]))
rows.sort(key=lambda x: -x[3])
assert len(rows) == 12
matrix = np.array([r[2:6] for r in rows])
fig = plt.figure(figsize=(W, 5.4))
ax = fig.add_axes([.385, .24, .42, .70])
im = ax.imshow(matrix, cmap="YlOrRd", vmin=0, vmax=100, aspect="auto")
display_names = {"Ribosomes (cyto + mito)": "Ribosomes",
                 "Nuclear pore scaffold": "Nuclear pore"}
ax.set(xticks=range(4), xticklabels=["Permissive\nclass", "Engaged\npermissive",
                                    "Maximum\nburial", "Met-Φ\npotential"],
       yticks=range(12), yticklabels=[
           f"{display_names.get(r[0], r[0])} (n={r[1]})" for r in rows])
ax.tick_params(length=0, pad=6)
for i, r in enumerate(rows):
    for j in range(4):
        ax.text(j, i, f"{matrix[i, j]:.0f}", ha="center", va="center",
                color="white" if matrix[i, j] > 62 else INK)
    ax.text(3.72, i, r[6], va="center", clip_on=False)
ax.text(3.72, -1.03, "Most buried\nprotein", va="center")
for spine in ax.spines.values():
    spine.set_visible(False)
cax = fig.add_axes([.385, .092, .42, .022])
fig.colorbar(im, cax=cax, orientation="horizontal",
             ticks=[0, 25, 50, 75, 100], label="Percent")
save(fig, "Figure_2", "Descriptive prefix-group summaries")

# Figure 3 uses the established 22 observations, without rebuilding coordinates.
raw = pd.read_csv(FOCAL)
f = raw[(raw.gene.isin(["PSMA4", "PSMA6", "PSMA7"])) &
        (raw.scenario == "unchanged_deposited_model") &
        (raw.modeled_upstream_residues == 0) &
        (((raw.resname == "MET") & (raw.measured_authnum == 1)) |
         ((raw.resname == "SER") & (raw.measured_authnum == 2)))].copy()
assert len(f) == 22 and (f.resname == "SER").sum() == 17
contexts = {"4R3O": "20S", "5LN3": "26S", "6KWY": "PA200–20S",
            "7V5G": "20S + substrate", "8CVS": "PA200–20S + MG-132",
            "8TM4": "pre-13S", "8YIX": "half-proteasome"}
fig = plt.figure(figsize=(W, 7.2))
fig.text(.025, .97, "a  What terminal chemistry is measured?", weight="bold", va="top")
ax = fig.add_axes([.025, .755, .95, .18])
ax.axis("off")
table = ax.table(
    cellText=[["Independent MS", "Removed; Ac-Ser", "Removed; Ac-Ser", "Removed; Ac-Ser"],
              ["Same preparation?", "Not established", "Not established", "Not established"],
              ["Modeled starts", "Met1 and Ser2", "Met1 and Ser2", "Met1 and Ser2"]],
    colLabels=["Evidence", "PSMA4 / P25789", "PSMA6 / P60900", "PSMA7 / O14818"],
    colWidths=[.25, .25, .25, .25], cellLoc="left", colLoc="left", bbox=[0, 0, 1, 1])
table.auto_set_font_size(False)
table.set_fontsize(FS)
for (r, c), cell in table.get_celld().items():
    cell.set_linewidth(1)
    cell.set_edgecolor("#C0C0C0")
    if r == 0:
        cell.set_facecolor("#F0F3F3")
        cell.get_text().set_weight("bold")
fig.text(.025, .722, "b  How does engagement vary by context?", weight="bold")
fig.text(.025, .690, "● Ser2     ◇ Met1     Chain identifiers label individual observations")
for k, gene in enumerate(["PSMA4", "PSMA6", "PSMA7"]):
    ax = fig.add_axes([.30 + k * .225, .283, .19, .355])
    ax.set_title(gene, weight="bold", pad=9)
    for j, pdb in enumerate(contexts):
        z = f[(f.gene == gene) & (f.pdb == pdb)].sort_values("chain")
        offsets = [0] if len(z) == 1 else [-.13, .13]
        for off, (_, row) in zip(offsets, z.iterrows()):
            y, x = 6 - j + off, row.buried_pct
            ax.scatter(x, y, s=28, marker="o" if row.resname == "SER" else "D",
                       facecolor=TEAL if row.resname == "SER" else "white",
                       edgecolor=TEAL if row.resname == "SER" else INK, zorder=3)
            ax.annotate(row.chain, (x, y), xytext=(5 if x < 85 else -5, 0),
                        textcoords="offset points", ha="left" if x < 85 else "right",
                        va="center")
        if k == 0:
            ax.text(-.13, (6 - j + .6) / 7.3, f"{pdb}\n{contexts[pdb]}",
                    transform=ax.transAxes, ha="right", va="center")
    ax.axvline(25, color="#808080", ls="--")
    ax.set(xlim=(-5, 105), ylim=(-.6, 6.7), xticks=[0, 50, 100],
           yticks=range(7), yticklabels=[])
    ax.grid(axis="y", color="#E2E2E2")
    ax.tick_params(axis="y", length=0)
    clean(ax)
    ax.spines["left"].set_visible(False)
fig.text(.64, .232, "Resolved-residue burial (%)", ha="center")
fig.text(.025, .187, "c  Which tests separate chemistry from context?", weight="bold")
ax = fig.add_axes([.025, .037, .95, .126])
ax.axis("off")
table = ax.table(cellText=[
    ["Vary terminal\nchemistry", "Fix assembly\ncontext", "Verify proteoforms; measure\npartner binding and assembly"],
    ["Vary assembly\ncontext", "Fix terminal\nchemistry", "Compare interfaces;\nmeasure assembly and function"]],
    colWidths=[.27, .27, .46], cellLoc="left", bbox=[0, 0, 1, 1])
table.auto_set_font_size(False)
table.set_fontsize(FS)
for (r, c), cell in table.get_celld().items():
    cell.set_linewidth(1)
    cell.set_edgecolor("#C0C0C0")
    if c == 0:
        cell.get_text().set_weight("bold")
save(fig, "Figure_3", "Proteasome terminal evidence and assembly-context questions")

# Figure 4 retains every rounded value and contact dot in the established map.
s = pd.read_csv(PARTNERS / "observation_partner_summary.csv").fillna("")
partners = pd.read_csv(PARTNERS / "all_partner_measurements.csv").fillna("")
s = s.sort_values(["gene", "pdb", "target_chain"]).reset_index(drop=True)
genes = ["PSMA1", "PSMA2", "PSMA3", "PSMA4", "PSMA5", "PSMA6", "PSMA7", "PSMG1", "POMP"]
M = np.zeros((22, 9))
contact = np.zeros_like(M, dtype=bool)
for i, row in s.iterrows():
    for j, g in enumerate(genes):
        z = partners[(partners.observation == row.observation) & (partners.partner_genes == g)]
        if len(z):
            M[i, j] = z.pair_only_buried_pct.max()
            contact[i, j] = (z.atom_pairs_le_4_A > 0).any()
expected = pd.read_csv(PARTNERS / "contact_map_pair_burial_values.csv", index_col=0)
assert np.allclose(M, expected.loc[s.observation, genes].values)
fig = plt.figure(figsize=(W, 6.9))
ax = fig.add_axes([.365, .155, .59, .70])
cmap = LinearSegmentedColormap.from_list("pair", ["#FFFFFF", "#BCE2E7", TEAL, "#1B474D"])
im = ax.imshow(M, cmap=cmap, vmin=0, vmax=100, aspect="auto")
ax.set_xticks(range(9), genes, rotation=55, ha="left")
ax.xaxis.tick_top()
labels = [f"{r.gene}  {r.pdb}/{r.target_chain}  {r.target_resname.title()}{r.target_authnum}"
          + ("*" if r.target_missing_heavy_atoms else "") for r in s.itertuples()]
ax.set_yticks(range(22), labels)
ax.tick_params(length=0, pad=5)
ax.set_xticks(np.arange(-.5, 9, 1), minor=True)
ax.set_yticks(np.arange(-.5, 22, 1), minor=True)
ax.grid(which="minor", color="#D4D4D4", lw=1)
ax.tick_params(which="minor", length=0)
for i in range(22):
    for j in range(9):
        color = "white" if M[i, j] > 55 else INK
        if M[i, j] >= 1:
            ax.text(j, i, f"{M[i, j]:.0f}", ha="center", va="center", color=color)
        if contact[i, j]:
            ax.plot(j + .32, i - .28, "o", ms=2, color=color)
for y in [7.5, 12.5]:
    ax.axhline(y, color=INK, lw=1.25)
for spine in ax.spines.values():
    spine.set_visible(False)
cax = fig.add_axes([.365, .078, .59, .021])
fig.colorbar(im, cax=cax, orientation="horizontal",
             ticks=[0, 25, 50, 75, 100], label="Pair-only resolved-residue burial (%)")
fig.text(.025, .98, "22 chain observations · 3 proteins · 7 entries", va="top")
save(fig, "Figure_4", "Partner-resolved PSMA4 PSMA6 and PSMA7 contact map")
for ext in ["png", "svg", "pdf", "tif"]:
    shutil.copy2(OUT / f"Figure_4.{ext}", OUT / f"Supplementary_Figure_3.{ext}")

# Figure 5 retains the original same-observation burial/distance examples.
obs = pd.read_csv(DATA / "asm_observations.csv")
transport = [("KPNB1", "6n88"), ("RAN", "7mo1"), ("KPNA2", "9bfc"),
             ("NUP188", "5ijo"), ("NUP93", "5ijo"), ("SEH1L", "5a9q"), ("NUP50", "3tj3")]
tr = []
for gene, pdb in transport:
    z = obs[(obs.pdb == pdb) & (obs.gene.str.split(";").str[0] == gene)]
    tr.append(z.loc[z.buried_pct.idxmax()])
tr = pd.DataFrame(tr).sort_values("buried_pct", ascending=False)
fig = plt.figure(figsize=(W, 3.8))
a = fig.add_axes([.33, .25, .34, .65])
b = fig.add_axes([.715, .25, .26, .65])
for y, row in zip(range(6, -1, -1), tr.itertuples()):
    color = BLUE if row.res2 in SMALL else ORANGE
    a.barh(y, row.buried_pct, color=color, height=.58)
    a.text(row.buried_pct + 2, y, f"{row.buried_pct:.1f}%", va="center")
    b.scatter(row.min_dist_alphaamino_N, y, color=color, s=28)
    b.text(row.min_dist_alphaamino_N + 2.2, y,
           f"{row.min_dist_alphaamino_N:.2f}", va="center")
a.set(yticks=range(6, -1, -1),
      yticklabels=[f"{r.gene}  {r.pdb.upper()}, {r.n_chains} chains" for r in tr.itertuples()],
      xlim=(0, 112), ylim=(-.6, 6.6), xlabel="Selected-position burial (%)")
b.set(yticks=[], xlim=(0, 66), ylim=(-.6, 6.6), xticks=[0, 20, 40, 60],
      xlabel="Measured-N distance (Å)")
a.axvline(25, color="#707070", ls="--")
b.axvline(5, color="#707070", ls="--")
heading(a, "a", "Positional burial")
heading(b, "b", "N distance")
for ax in [a, b]:
    clean(ax)
    ax.tick_params(axis="y", length=0)
fig.legend(handles=[Patch(facecolor=BLUE, label="MetAP-permissive"),
                    Patch(facecolor=ORANGE, label="MetAP-nonpermissive")],
           frameon=False, ncol=2, loc="lower center", bbox_to_anchor=(.63, .015))
save(fig, "Figure_5", "Nuclear-transport positional geometry")

# Supplementary figures are re-rendered from already computed source tables.
rare = pd.read_csv(SUPP / "figure_S1_source.csv")
fig, axes = plt.subplots(3, 1, figsize=(W, 6.0), layout="constrained")
for k, (col, title, ylabel) in enumerate([
    ("protein_count", "Proteins represented", "Proteins"),
    ("percent_engaged", "Sampling dependence of maximum burial", "Engaged proteins (%)"),
    ("adjusted_odds_ratio", "Sequence-class association", "Adjusted odds ratio"),
]):
    z = rare.pivot(index="pdb_entries", columns="replicate", values=col)
    axes[k].fill_between(z.index, np.nanpercentile(z, 10, axis=1),
                         np.nanpercentile(z, 90, axis=1), color="#BCE2E7")
    axes[k].plot(z.index, np.nanmedian(z, axis=1), color=TEAL, lw=1.6)
    axes[k].set(xscale="log", xlim=(20, 6305), ylabel=ylabel)
    axes[k].set_xticks([20, 100, 1000, 6305], ["20", "100", "1,000", "6,305"])
    heading(axes[k], "abc"[k], title)
    clean(axes[k])
axes[0].set_ylim(0, 1200)
axes[1].set_ylim(0, 55)
axes[2].set_ylim(0, 2.5)
axes[2].axhline(1, color="#707070", ls="--")
axes[2].set_xlabel("PDB entries sampled (log scale)")
save(fig, "Supplementary_Figure_1", "Corpus-size sampling")

cascade = pd.read_csv(SUPP / "figure_S2_source.csv")
groups = ["Myristoylation annotation", "Strict NTMT motif",
          "N-terminal acetylation annotation", "None of these flags"]
labels = ["Myristoylation\nannotation", "Strict NTMT\nmotif",
          "N-acetylation\nannotation", "None of\nthese flags"]
fig, axes = plt.subplots(2, 1, figsize=(W, 7.1), layout="constrained",
                         gridspec_kw={"height_ratios": [1, 1.65]})
rng = np.random.default_rng(7)
for i, group in enumerate(groups):
    vals = cascade.loc[cascade.display_group == group, "buried"]
    axes[0].scatter(i + rng.uniform(-.19, .19, len(vals)), vals,
                    s=10, alpha=.45, color=TEAL)
    axes[0].plot([i - .3, i + .3], [vals.median()] * 2, color=INK, lw=1.6)
    axes[0].text(i, 104, f"n={len(vals)}", ha="center")
axes[0].set(xticks=range(4), xticklabels=labels, ylim=(-3, 115), ylabel="Burial (%)")
axes[0].axhline(25, color="#707070", ls="--")
heading(axes[0], "a", "Annotation/motif groups in the permissive class")
cases = pd.concat([cascade[cascade.display_group == g].sort_values("buried") for g in groups[:2]])
axes[1].barh(range(len(cases)), cases.buried,
             color=[TEAL if g == groups[0] else "#A84B2F" for g in cases.display_group])
axes[1].set(yticks=range(len(cases)), yticklabels=cases.gene, xlim=(0, 116),
            xlabel="Selected-position burial (%)")
for i, v in enumerate(cases.buried):
    axes[1].text(v + 1.5, i, f"{v:.1f}", va="center")
axes[1].axvline(25, color="#707070", ls="--")
heading(axes[1], "b", "Myristoylation-annotation and strict-motif examples")
for ax in axes:
    clean(ax)
save(fig, "Supplementary_Figure_2", "Annotation and motif groups")
(OUT / "artwork_checks.json").write_text(json.dumps(records, indent=2))
print(json.dumps(records, indent=2))
```
