## Supplementary_Data_8a for "Human protein complex interfaces reveal candidate assembly roles for initiator methionine excision": README.md

# Supplementary Data 8: audit, current numerical exports and figure source data

This archive supports the evidence-qualified Paper 1 supplement. It does not convert coordinate-model assignments into sample-validated native proteoforms.

## Authoritative current outputs

- `outputs/protein_census.csv`: 1,191 original protein rows, with corrected evidence labels and flags; 1,123 analyzable.
- `outputs/observations.csv`: 22,291 observations; 19,967 analyzable.
- `outputs/boundary_selected_845.csv`: First-observed observations filtered before protein-level maximum reselection.
- `outputs/candidates_original_261.csv` and `outputs/candidates_boundary_175.csv`: Distinct candidate definitions.
- `outputs/*tests.csv`, `current_models.csv`, `alternative_definitions.csv`, `assembly_strata.csv`, `class_summaries.csv`, `residue2_summary.csv`, `leave_one_residue_out.csv` and `sampling_sensitivities.csv`: Recomputed current-cohort numerical exports. Statistical test output is imported into the submission workbooks, not implemented as Excel formulas.
- `outputs/figure_S1_source.csv`: One row per replicate and sampling checkpoint (40 orderings × 44 checkpoints).
- `outputs/figure_S2_source.csv`: One row per plotted permissive-class protein (696 rows); operational annotation/motif categories, selected modeled identity and boundary flags.
- `audit_results/`: Observation mapping and processing flags, boundary transitions, association sensitivities, coordinate checks and focal geometry diagnostics.
- `outputs/reproduction_check.json` and validation files: Numerical checks of this build where available.

Distances are in angstroms, surface areas in square angstroms and burial in percent (0–100), not proportions. Statistical fractions use 0–1. Accession, PDB and chain identifiers are case-sensitive text. Blank/nonfinite values are unavailable, not zero.

## Recompute the current tables and Figs. 1–2

From the extracted archive root:

```sh
python -m pip install -r requirements.txt
python build_supplement_analysis.py
```

The script reads the included frozen input tables and audit exports. It recalculates the current cohort statistics and first-observed reselection, and regenerates the two figures and their source data. It does not refit the separately labeled supplied volume/hydropathy models.

The N-distance comparison excludes five nonfinite missing-value sentinels (four permissive and one nonpermissive). No such exclusions are applied to the all-protein burial comparison. The first-observed sensitivity preserves the frozen selected observation in maximum-burial ties when possible, then uses frozen observation order.

Figure 1 uses the frozen rarefaction implementation in `code/rebuild_supplementary_figure1.py` with revised artwork. Its tie convention selects the earliest observation-table row; the full endpoint reproduces the primary adjusted OR. Sampling bands are 10th–90th percentiles across random orderings, not confidence intervals.

Figure 2 uses exported annotation/motif flags with priority myristoylation, strict NTMT motif, acetylation, then none of these flags. They are not preparation-matched chemical measurements. The old distance-at-maximum presentation has been removed because the exported minimum distance may arise from another observation.

## Audit reproduction and provenance

The original targeted audit scripts, results, geometry library and 11 frozen assemblies are included under `paper1_audit/`. From the archive root:

```sh
python paper1_audit/audit.py
python paper1_audit/coordinate_followup.py
python paper1_audit/validate_and_summarize.py
```

These scripts use their original vocabulary internally to preserve provenance. Their output diagnostics do not validate every biological boundary. Geometry calculations can take longer than the table/figure rebuild.

The `paper1_audit/results/` copy and `audit_results/` snapshot describe the same audit release. The latter is the fixed input snapshot for the current-table build; rerunning the audit does not silently replace that snapshot.

Original submitted workbooks under `paper1_v8/`, original frozen workbooks under `paper1_audit/frozen/`, and CSVs under `historical_provenance/` are historical evidence only. They can contain superseded processing labels, 1,143-protein results, the Ran scan-target error and legacy manuscript wording. They are not the current submission workbooks.

The inherited recurrence code and JSON outputs are preserved to document the numerical origin of Data 6. The current workbook changes interpretation and labels, not those original recurrence numbers. The 261-candidate denominator must not be substituted with 175 without a new analysis.

The historical curated-enrichment source tables were not located in the supplied package, and this archive does not reproduce those tests. The frozen prefix map remains descriptive. The synchronized manuscript now reports a new, explicitly defined enrichment reconstruction in Supplementary Data 9, including curated membership, complete correction families and boundary sensitivities; that separate reconstruction does not restore the undocumented historical statistics.

## Current main-figure terminology

`code/relabel_main_figures.py` retains the frozen measurements for main Figures 1, 2 and 4 while changing labels to selected-position burial, sequence-class potential and measured-N geometry. To rebuild those figures from this archive root:

```sh
NME_DIR="$PWD/paper1_audit/frozen/nterm-census-1.3.0" \
NME_WORK="$PWD/paper1_audit/frozen/nterm-census-1.3.0/data" \
NME_FIGOUT="$PWD/main_figure_art" python code/relabel_main_figures.py
```

The inherited script also emits a historical alpha-ring Figure 3. That output is not a current manuscript figure and must not be substituted for the evidence-aware Figure 3. Current Figure 3 and the partner-resolved map are supplied in the package's figure-source folder; partner-map measurements and reproduction are in Supplementary Data 7. The current main-figure PNG/SVG exports are included under `main_figure_art/`.

## Integrity and validation limits

`SHA256SUMS.txt` covers all archive files except itself. Frozen coordinate hashes and audit validation are also included. Structural proximity is not a validated bond; pair-only burial is nonadditive; repeated chains and structures are not independent biological replicates.

Partner-resolved measurements and their seven-coordinate portable reproduction are supplied separately in Supplementary Data 7. Neither archive establishes processing causality or disease mechanism. No missing focal side-chain atoms were rebuilt.
