## Supplementary_Data_8a for "Human protein complex interfaces reveal candidate assembly roles for initiator methionine excision": TRANSFER_EXTRACTION_INSTRUCTIONS.md

# Supplementary Data 8: two-part delivery

Supplementary Data 8 is delivered as two ordinary ZIP archives so that each upload is below 50 MB. These are not split-volume ZIP files and do not require special archive software.

1. Extract Supplementary Data 8a (analysis and source data) into an empty directory.
2. Extract Supplementary Data 8b (frozen inputs) into the same directory.
3. Both archives use the same top-level `Supplementary_Data_8` folder. Merge the folders; do not create a second nested copy.
4. Read `Supplementary_Data_8/README.md` and run the existing commands from that folder.

Part 8a contains the analysis, exports, source values, code, documentation and historical provenance. Part 8b contains the frozen annotation, mapping and assembly inputs under `paper1_audit/frozen/nterm-census-1.3.0/inputs/`.

Every member of the preceding single-archive release is preserved byte-for-byte at the same relative path. The original checksum manifests still apply to those members after extraction. This delivery split does not alter any measurement, statistical result, code or input file.

The transfer manuscript presents the former Figure 3d contact map as Figure 4 and the former Figure 4 nuclear-transport artwork as Figure 5. Archived scripts and historical provenance retain their original internal file names; these do not override the current manuscript numbering. The current upload figure files are the authoritative numbered artwork.
