## Supplementary_Data_9 for "Human protein complex interfaces reveal candidate assembly roles for initiator methionine excision": analyze.py

```
"""Portable, outcome-independent membership reconstruction and enrichment tests."""
from pathlib import Path
import re, json, hashlib, ast
import numpy as np
import pandas as pd
from scipy.stats import fisher_exact, hypergeom
from scipy.stats.contingency import odds_ratio
from statsmodels.stats.multitest import multipletests

ROOT=Path(__file__).resolve().parent
I=ROOT/"inputs"; O=ROOT/"outputs"; O.mkdir(exist_ok=True)
cp=pd.read_csv(I/"9606_curated.tsv",sep="\t").fillna("")
p=pd.read_csv(I/"protein_census.csv")
b=pd.read_csv(I/"boundary_selected_845.csv")
observations=pd.read_csv(I/"observations.csv",low_memory=False)
eligible_observations=observations[observations.analyzable & observations.is_first_observed]
b["n_entries"]=b.acc.map(eligible_observations.groupby("acc").pdb.nunique())
assert b.n_entries.notna().all()
allp=p[~p.excluded_from_analysis].copy()
orig=allp[allp.sequence_class.eq("MetAP-permissive")].copy()
boundary=b[b.res2.isin(list("AGSTCPV"))].copy()
assert len(allp)==1123 and len(orig)==696 and len(boundary)==463
assert orig.acc.is_unique and boundary.acc.is_unique
cohorts={"original":orig,"first_observed":boundary}
uni=re.compile(r"^(?:[OPQ][0-9][A-Z0-9]{3}[0-9]|[A-NR-Z][0-9][A-Z][A-Z0-9]{2}[0-9](?:[A-Z][A-Z0-9]{2}[0-9])?)$")
members={}; strict={}; audits=[]
assert cp["Taxonomy identifier"].eq(9606).all()
assert not cp["Evidence Code"].str.contains("0007653|0008004").any()
for _,r in cp.iterrows():
    cid=r["#Complex ac"]; m=set(); s=set()
    for item in r["Expanded participant list"].split("|"):
        token=item.split("(")[0]
        alternative=token.startswith("[")
        for raw in token.strip("[]").split(","):
            base=raw.split("-")[0]
            accepted=bool(uni.fullmatch(base))
            audits.append(dict(complex_id=cid,raw_identifier=raw,canonical_accession=base if accepted else "",
                alternative_set=alternative,suffix_normalized=raw!=base,accepted=accepted))
            if accepted:
                m.add(base)
                if not alternative and raw==base:s.add(base)
    members[cid]=m;strict[cid]=s
pd.DataFrame(audits).to_csv(O/"participant_mapping_audit.csv",index=False)

# Extract only the literal frozen membership function, never execute plotting code.
tree=ast.parse((I/"frozen_figure_code.py").read_text())
func=next(n for n in tree.body if isinstance(n,ast.FunctionDef) and n.name=="machine")
space={"re":re};exec(compile(ast.Module(body=[func],type_ignores=[]),"<frozen machine function>","exec"),space)
allp["prefix_group"]=allp.gene.map(space["machine"])
counts=allp.prefix_group.value_counts()
prefix={n:set(allp.loc[allp.prefix_group.eq(n),"acc"]) for n,k in counts.items() if k>=4}
assert len(prefix)==12
prefix["Histone-symbol family"]=set(allp.loc[allp.gene.str.match(r"^(HIST|H[1234](?:[A-Z0-9-]|$))"),"acc"])
meta=cp.set_index("#Complex ac")
eligible=[cid for cid,m in members.items() if len(m & set(orig.acc))>=4]
catalogue={cid:members[cid] for cid in eligible}
focused_ids={
    "26S proteasome":["CPX-5993"],
    "CCT/TRiC":["CPX-6030"],
    "OXPHOS I–V":["CPX-560","CPX-561","CPX-577","CPX-6123","CPX-6151"],
    "Spliceosomal complexes":cp.loc[cp["Recommended name"].str.contains("spliceosom",case=False),"#Complex ac"].tolist(),
    "Core-histone membership":cp.loc[cp["Recommended name"].str.match(r"(?i)^(Nucleosome|CENP-A nucleosome)"),"#Complex ac"].tolist()
}
focused={name:set().union(*(members[c] for c in ids)) for name,ids in focused_ids.items()}
json.dump(focused_ids,open(O/"focused_complex_definitions.json","w"),indent=2)
definitions=[]
for family,groups in [("prefix13",prefix),("curated_catalogue",catalogue),("focused5",focused)]:
    for group,ms in groups.items():
        for acc in sorted(ms):
            definitions.append(dict(family=family,group=group,accession=acc,
                complex_name=meta.loc[group,"Recommended name"] if group in meta.index else "",
                source_complex_ids="|".join(focused_ids.get(group,[group] if group in meta.index else [])),
                original_eligible=acc in set(orig.acc),boundary_eligible=acc in set(boundary.acc)))
pd.DataFrame(definitions).to_csv(O/"group_memberships.csv",index=False)

def test(d,m,threshold=25):
    ing=d.acc.isin(m);hit=d.buried_pct.ge(threshold)
    a=int((ing&hit).sum()); bb=int((ing&~hit).sum())
    c=int((~ing&hit).sum());dd=int((~ing&~hit).sum())
    table=[[a,bb],[c,dd]]
    if a+bb==0 or c+dd==0:
        ratio=lo=hi=np.nan;pv=1.
    else:
        ratio,pv=fisher_exact(table)
        ci=odds_ratio(table,kind="conditional").confidence_interval()
        lo,hi=ci.low,ci.high
    return dict(member_engaged=a,member_unengaged=bb,nonmember_engaged=c,nonmember_unengaged=dd,
                member_n=a+bb,background_n=len(d),odds_ratio=ratio,ci_low=lo,ci_high=hi,p_two_sided=pv)

def tests(groups,family,thresholds=[25]):
    rows=[]
    for co,d in cohorts.items():
        for th in thresholds:
            for group,m in groups.items():
                rows.append(dict(family=family,cohort=co,threshold=th,group=group,
                    name=meta.loc[group,"Recommended name"] if group in meta.index else group,**test(d,m,th)))
    df=pd.DataFrame(rows)
    for _,inds in df.groupby(["cohort","threshold"]).groups.items():
        df.loc[inds,"q_BH"]=multipletests(df.loc[inds,"p_two_sided"],method="fdr_bh")[1]
        df.loc[inds,"q_BY"]=multipletests(df.loc[inds,"p_two_sided"],method="fdr_by")[1]
        df.loc[inds,"family_tests"]=len(inds)
    df.to_csv(O/f"{family}_tests.csv",index=False)
    return df

pref=tests(prefix,"prefix13")
cat=tests(catalogue,"curated_catalogue")
focus=tests(focused,"focused5",[20,25,30])
tests({k:v for k,v in prefix.items() if k!="Histone-symbol family"},"prefix12_sensitivity")
tests({cid:strict[cid] for cid in eligible},"curated_strict_identifiers")
tests({name:set().union(*(strict[c] for c in ids)) for name,ids in focused_ids.items()},"focused_strict_identifiers")
mt=set(allp.loc[allp.gene.str.startswith("MT-"),"acc"])
tests({"OXPHOS mitochondrially encoded":focused["OXPHOS I–V"]&mt,
       "OXPHOS nuclear encoded":focused["OXPHOS I–V"]-mt},"oxphos_encoding_sensitivity")

# Dependence from repeated overlapping curated memberships: deduplicate the exact
# original tested accession sets; do not count a family of variants as replication.
unique={}
for cid in eligible:
    key=tuple(sorted(members[cid]&set(orig.acc)))
    unique.setdefault(key,[]).append(cid)
dedup={"|".join(ids):set(key) for key,ids in unique.items()}
# Full membership union of equivalent original sets may differ in boundary only
# if backgrounds were not nested; assert nesting and use canonical full set.
assert set(boundary.acc)<=set(orig.acc)
tests(dedup,"curated_unique_tested_sets")

def conditional_stratified(d,m,threshold=25):
    d=d.copy()
    d["size_stratum"]=pd.cut(d.n_chains,[0,3,5,11,np.inf],labels=False)
    d["entry_stratum"]=pd.cut(d.n_entries,[0,1,4,np.inf],labels=False)
    d["member"]=d.acc.isin(m);d["engaged"]=d.buried_pct.ge(threshold)
    distribution=np.array([1.]);observed=0;expected=0.;informative=0
    for _,x in d.groupby(["size_stratum","entry_stratum"],observed=True):
        N=len(x);K=int(x.engaged.sum());n=int(x.member.sum())
        observed+=int((x.member&x.engaged).sum());expected+=n*K/N
        probs=hypergeom.pmf(np.arange(n+1),N,K,n)
        distribution=np.convolve(distribution,probs)
        informative+=int(0<n<N and 0<K<N)
    pv=float(distribution[observed:].sum())
    return dict(observed_engaged_members=observed,expected_engaged_members=expected,
                informative_strata=informative,p_stratified_enrichment=min(pv,1.))
strats=[]
for co,d in cohorts.items():
    for group,m in focused.items():
        strats.append(dict(cohort=co,group=group,**conditional_stratified(d,m)))
st=pd.DataFrame(strats)
for _,ix in st.groupby("cohort").groups.items():
    st.loc[ix,"q_BH_stratified5"]=multipletests(st.loc[ix,"p_stratified_enrichment"],method="fdr_bh")[1]
st.to_csv(O/"focused_stratified_tests.csv",index=False)
hist_strat=pd.DataFrame([dict(cohort=co,group="Histone-symbol family",
    **conditional_stratified(d,prefix["Histone-symbol family"])) for co,d in cohorts.items()])
hist_strat.to_csv(O/"histone_prefix_stratified_sensitivity.csv",index=False)

# Auditable protein-level selections used in every test.
coverage=[]
for co,d in cohorts.items():
    for family,groups in [("focused5",focused),("prefix13",prefix)]:
        for group,m in groups.items():
            for _,r in d[d.acc.isin(m)].iterrows():
                coverage.append(dict(cohort=co,family=family,group=group,accession=r.acc,gene=r.gene,
                    burial=r.buried_pct,engaged_25=r.buried_pct>=25,pdb=r.pdb,chain=r.chain,
                    n_chains=r.n_chains,n_entries=r.n_entries,is_first_observed=r.is_first_observed))
pd.DataFrame(coverage).to_csv(O/"tested_member_records.csv",index=False)
source={"complex_portal_url":"https://ftp.ebi.ac.uk/pub/databases/intact/complex/current/complextab/9606.tsv",
        "documentation_url":"https://www.ebi.ac.uk/complexportal/documentation",
        "retrieved_utc":"2026-09-25","catalogue_records":len(cp),"eligible_curated_tests":len(eligible),
        "unique_original_tested_sets":len(unique),"original_permissive_n":len(orig),
        "boundary_permissive_n":len(boundary),
        "input_sha256":{f.name:hashlib.sha256(f.read_bytes()).hexdigest() for f in I.iterdir() if f.is_file()}}
json.dump(source,open(O/"provenance.json","w"),indent=2)
print(json.dumps(source,indent=2))
print("\nFOCUSED 25%\n",focus[focus.threshold.eq(25)].to_string(index=False))
print("\nSTRATIFIED\n",st.to_string(index=False))
print("\nPREFIX\n",pref.to_string(index=False))
print("\nCATALOGUE TOP\n",cat.sort_values(["cohort","q_BH"]).groupby("cohort").head(12).to_string(index=False))
```
