## Supplementary_Data_9 for "Human protein complex interfaces reveal candidate assembly roles for initiator methionine excision": PROTOCOL.md

# Paper 1 enrichment reconstruction protocol

This is a new exploratory reconstruction, not recovery of the missing historical analysis. Membership and test-family rules below were written before calculating enrichment outcomes in this reconstruction. The biological targets were selected because the manuscript already discussed them, not as a prospective independent validation.

## Inputs and endpoint

Use the frozen 1,123-protein census and its 696 MetAP-permissive proteins as the original background. Use the previously audited first-observed sensitivity and its 463 permissive proteins as the boundary-qualified background. The outcome is selected-position burial of at least 25%, with one accession counted once. Filter observations before selecting the sensitivity maximum, using the existing audited table. Missing eligibility is not non-engagement.

## Membership and families

- **Prefix reconstruction:** Reuse the exact ordered gene-pattern assignment in the frozen Figure 2 code. Retain its twelve displayed groups, selected by at least four scored proteins in the original census. Add a clearly separate histone-prefix group (`HIST` or modern `H1`/`H2`/`H3`/`H4` histone symbols), for a 13-test family. Report historical twelve-group BH correction only as a sensitivity, never as a replacement for the 13-test correction.
- **Catalogue-wide curated screen:** Use all manually curated human Complex Portal records from the downloaded `9606.tsv`, excluding machine-learning predictions. Require at least four distinct permissive accessions represented in the original background, regardless of their burial. Fix this eligible family for the boundary sensitivity as well. Count accession membership only, ignoring stoichiometry. Parse the expanded participant list; normalize isoform/processed-chain suffixes to the canonical accession while preserving the original identifier and its normalization in an audit. Expand alternative sets as possible membership, flag them, and repeat a sensitivity excluding alternatives and suffix-normalized participants. Protein membership in a curated complex does not mean its maximum-burial structure depicts that complex.
- **Focused five-question reconstruction:** 26S proteasome CPX-5993; CCT/TRiC CPX-6030; OXPHOS union of CPX-560, CPX-561, CPX-577, CPX-6123 and CPX-6151; union of every recommended name containing `spliceosom`; and core-histone union of names beginning `Nucleosome` or `CENP-A nucleosome` (excluding nucleosome-remodeling complexes). These are five explicitly defined exploratory questions, not the missing twelve curated tests. Report BH over all five, and compare with the catalogue-wide screen. The nucleosome membership union tests histone-protein membership, not nucleosome-specific coordinates or histone-tail recognition chemistry.

## Statistics

For each group and background, report [[member engaged, member unengaged], [nonmember engaged, nonmember unengaged]], two-sided Fisher p, sample odds ratio, exact conditional odds-ratio confidence interval, BH q and dependence-conservative BY q. Also report an accession-level membership/coverage table and the eligible protein list with outcome.

For focused groups, repeat at 20% and 30% burial, retaining the same five-test family separately at each threshold. The 25% endpoint remains primary. Add exact conditional stratified upper-tail enrichment tests with fixed membership and outcome counts within assembly-size strata (2–3, 4–5, 6–11, >=12 chains) crossed with PDB-entry-count strata (1, 2–4, >=5). Use convolution of hypergeometric distributions, not simulations. These tests address those measured sampling covariates only. Their directionality and family correction must be labeled explicitly.

Boundary-qualified results are the stronger geometric sensitivity, but neither boundary filtering nor annotation membership proves native chemistry, processing causality, disease relevance or independence of homologous proteins. Repeated chains are not independent replicates. The screen remains conditional on the structurally observed census.

## Deliverables

Implementation correction: the initial modern-histone regular expression omitted the `B` in H2BC11. It was broadened to all histone-style H1/H2/H3/H4 symbols, restoring this unengaged record rather than selectively excluding it. This changes the prefix histone denominator from 10 to 11 in the original cohort. The correction and the initial result are recorded here to prevent silent outcome-dependent rule changes. Additional post-result diagnostics separate nuclear-encoded from mitochondrially encoded OXPHOS members and compute the prefix-histone sampling-stratified sensitivity; they are not new primary discovery tests.

Frozen source catalogue, source URLs and hashes; explicit memberships and normalization audit; all test tables and corrections; portable code; reconstruction report; manuscript-ready replacement language. Do not restore any historical q value merely because a new related comparison is significant.
