## Supplementary_Data_9 for "Human protein complex interfaces reveal candidate assembly roles for initiator methionine excision": README.md

# Supplementary Data 9: reconstructed enrichment

This archive supplies the exploratory enrichment reconstruction reported in the synchronized manuscript, Supplementary Note 6 and Supplementary Tables 3–4. The historical curated test definitions were not recovered; the new explicit test families must not be represented as reproduction of the old statistics.

## Reproduce

The recorded environment is Python 3.14.3; exact package versions are in `requirements.txt`. Run `python analyze.py`, followed by `python validate.py`, from an environment with those packages installed. All input data are included; neither script downloads data or requires authentication. The source catalogue, frozen census, audited boundary selection, observation-level sampling data and historical membership function are under `inputs/`. `SHA256SUMS.txt` records the packaged files for integrity checking.

The statistical protocol was written before calculating the new results. An implementation correction to the histone regular expression and explicitly post-result diagnostic sensitivities are recorded in `PROTOCOL.md`.

## Read the outputs correctly

- `prefix13_tests.csv`: twelve frozen prefix groups plus the histone-symbol family, corrected together.
- `prefix12_sensitivity_tests.csv`: the twelve historical display groups, without histones; not the main correction family.
- `curated_catalogue_tests.csv`: sixty-four eligible manually curated human complexes.
- `curated_unique_tested_sets_tests.csv`: forty-two distinct tested accession sets.
- `focused5_tests.csv`: five explicit questions, each at 20%, 25% and 30% burial.
- `curated_strict_identifiers_tests.csv` and `focused_strict_identifiers_tests.csv`: mapping sensitivities.
- `focused_stratified_tests.csv`: exact one-sided enrichment tests conditional on assembly-size and deposition-count strata, with focused-family BH correction.
- `oxphos_encoding_sensitivity_tests.csv`: post-result nuclear/mitochondrial-encoding split.
- `histone_prefix_stratified_sensitivity.csv`: post-result sampling-stratified histone diagnostic, not a new discovery family.
- `group_memberships.csv`, `participant_mapping_audit.csv` and `tested_member_records.csv`: auditable accession assignments, eligibility and selected structural observations.
- `focused_complex_definitions.json`: exact curated records entering the five focused comparisons.
- `provenance.json` and `validation.json`: source hashes, sample sizes and independent checks.

The primary threshold is 25%. Every q value belongs to a specific declared family and background. Small or zero cells produce wide or infinite estimates. Catalogue overlap, homologous proteins, structural ascertainment, terminal chemistry and construct identity remain limitations.

`Enrichment_Results.xlsx` is a browsable copy of the analysis outputs, not a spreadsheet recalculation of exact statistical tests. Complete CSV outputs retain full precision. Source rows for Supplementary Tables 3–4 are also supplied under `outputs/`; their display rounding does not change the calculations. The manuscript text, methods and legends distinguish original extraction, first-observed sensitivity, prefix membership, curated membership and focused questions.

The catalogue was downloaded from the [Complex Portal human ComplexTab endpoint](https://ftp.ebi.ac.uk/pub/databases/intact/complex/current/complextab/9606.tsv). The [database documentation](https://www.ebi.ac.uk/complexportal/documentation) describes manual curation, inferred evidence, and separately distributed predicted complexes. Manual curation is not equivalent to experimental proof for every record.
