## Supplementary_Data_9 for "Human protein complex interfaces reveal candidate assembly roles for initiator methionine excision": validate.py

```
"""Independent numerical and membership checks, with no manuscript assumptions."""
from pathlib import Path
import json, re, ast, hashlib
import numpy as np
import pandas as pd
from scipy.stats import hypergeom
R=Path(__file__).resolve().parent;O=R/"outputs";I=R/"inputs"
checks=[]
def check(name,value):
    assert bool(value),name
    checks.append({"check":name,"passed":True})
def bh_manual(p):
    p=np.asarray(p);order=np.argsort(p);v=p[order]*len(p)/np.arange(1,len(p)+1)
    v=np.minimum.accumulate(v[::-1])[::-1];out=np.empty(len(p));out[order]=np.minimum(v,1)
    return out
for f in O.glob("*_tests.csv"):
    d=pd.read_csv(f)
    if "p_two_sided" not in d:continue
    for r in d.itertuples():
        a,b,c,e=r.member_engaged,r.member_unengaged,r.nonmember_engaged,r.nonmember_unengaged
        N=a+b+c+e;K=a+c;n=a+b
        support=np.arange(max(0,n-(N-K)),min(n,K)+1)
        mass=hypergeom.pmf(support,N,K,n);observed=hypergeom.pmf(a,N,K,n)
        pv=min(1.,mass[mass<=observed*(1+1e-10)].sum())
        assert np.isclose(pv,r.p_two_sided,rtol=1e-8,atol=1e-12),(f.name,r.group)
        assert N==r.background_n and a+b==r.member_n
    check(f"{f.name}: independent hypergeometric enumeration and denominators",True)
    for _,inds in d.groupby(["cohort","threshold"]).groups.items():
        expected=bh_manual(d.loc[inds,"p_two_sided"])
        check(f"{f.name}: BH group {d.loc[inds[0],'cohort']}/{d.loc[inds[0],'threshold']}",
              np.allclose(expected,d.loc[inds,"q_BH"],rtol=1e-12,atol=1e-12))
        harmonic=sum(1/i for i in range(1,len(inds)+1))
        check(f"{f.name}: BY group {d.loc[inds[0],'cohort']}/{d.loc[inds[0],'threshold']}",
              np.allclose(np.minimum(expected*harmonic,1),d.loc[inds,"q_BY"],atol=1e-12))
orig=pd.read_csv(I/"protein_census.csv")
boundary=pd.read_csv(I/"boundary_selected_845.csv")
check("Boundary set nested within original accession universe",set(boundary.acc)<=set(orig.acc))
check("Boundary-selected records all marked first observed",boundary.is_first_observed.all())
check("No duplicate accession in either protein table",orig.acc.is_unique and boundary.acc.is_unique)
gm=pd.read_csv(O/"group_memberships.csv")
check("No duplicate memberships",not gm.duplicated(["family","group","accession"]).any())
raw=pd.read_csv(O/"participant_mapping_audit.csv")
check("No unresolved expanded participants",raw.accepted.all())
check("Focused membership unchanged by strict identifier sensitivity",
      pd.read_csv(O/"focused5_tests.csv").query("threshold==25")[["member_n","member_engaged"]].reset_index(drop=True).equals(
      pd.read_csv(O/"focused_strict_identifiers_tests.csv")[["member_n","member_engaged"]].reset_index(drop=True)))
cat=pd.read_csv(O/"curated_catalogue_tests.csv")
check("Boundary catalogue has no BH discovery",not cat.query("cohort=='first_observed'").q_BH.lt(.05).any())
meta=pd.read_csv(I/"9606_curated.tsv",sep="\t")
check("Curated catalogue excludes predicted evidence",not meta["Evidence Code"].str.contains("0007653|0008004").any())
summary={"all_passed":True,"checks":checks,"total_checks":len(checks),
 "limits":"Numerical/membership verification is not validation of native chemistry or independence of homologous proteins."}
(O/"validation.json").write_text(json.dumps(summary,indent=2))
print(json.dumps({"all_passed":True,"total_checks":len(checks)},indent=2))
```
