## Supplementary_Information for "Human protein complex interfaces reveal candidate assembly roles for initiator methionine excision"

Yie-Hwa Chang

Edward A. Doisy Department of Biochemistry and Molecular Biology, Saint Louis University School of Medicine, St. Louis, Missouri, USA

#### Scope and evidence convention

This supplement describes protein-interface engagement of annotation-selected N-terminal sequence positions in deposited human protein assemblies. Sequence-class associations, annotation evidence, modeled residue identity, coordinate boundary and chemical state are kept distinct. Neither modeled Met1 nor a first-observed residue alone establishes the processing state of the structural preparation.

The frozen export contains 1,191 protein records and 22,291 chain observations. After excluding 48 signal/propeptide/transit records and 20 immunoglobulin/T-cell-receptor constant-region records, the primary analysis contains 1,123 proteins, 19,967 observations and 6,305 PDB entries. The first-observed sensitivity retains 845 proteins after filtering observations and reselecting maxima.

#### Contents

Supplementary Figs. 1–3: corpus sampling, annotation/motif groups and partner-resolved proteasome geometry.

**Supplementary Notes: grouping, Ran boundary, experimental design, proteasome evidence, audit methods and reconstructed enrichment.**

**Supplementary Tables 1–4: robustness, experiment priorities, thirteen-group tests and curated/focused comparisons.**

Supplementary Data 1–9: six standalone workbooks and four archives for partner-resolved geometry, audit/source data (Data 8a and 8b) and reconstructed enrichment.

“Engaged” denotes at least 25% protein-only residue burial unless otherwise specified. It is an operational geometric threshold, not a validated interaction or functional state. Distances refer to the measured residue N atom, not necessarily a free terminal amino group.

**Supplementary Fig. 1**

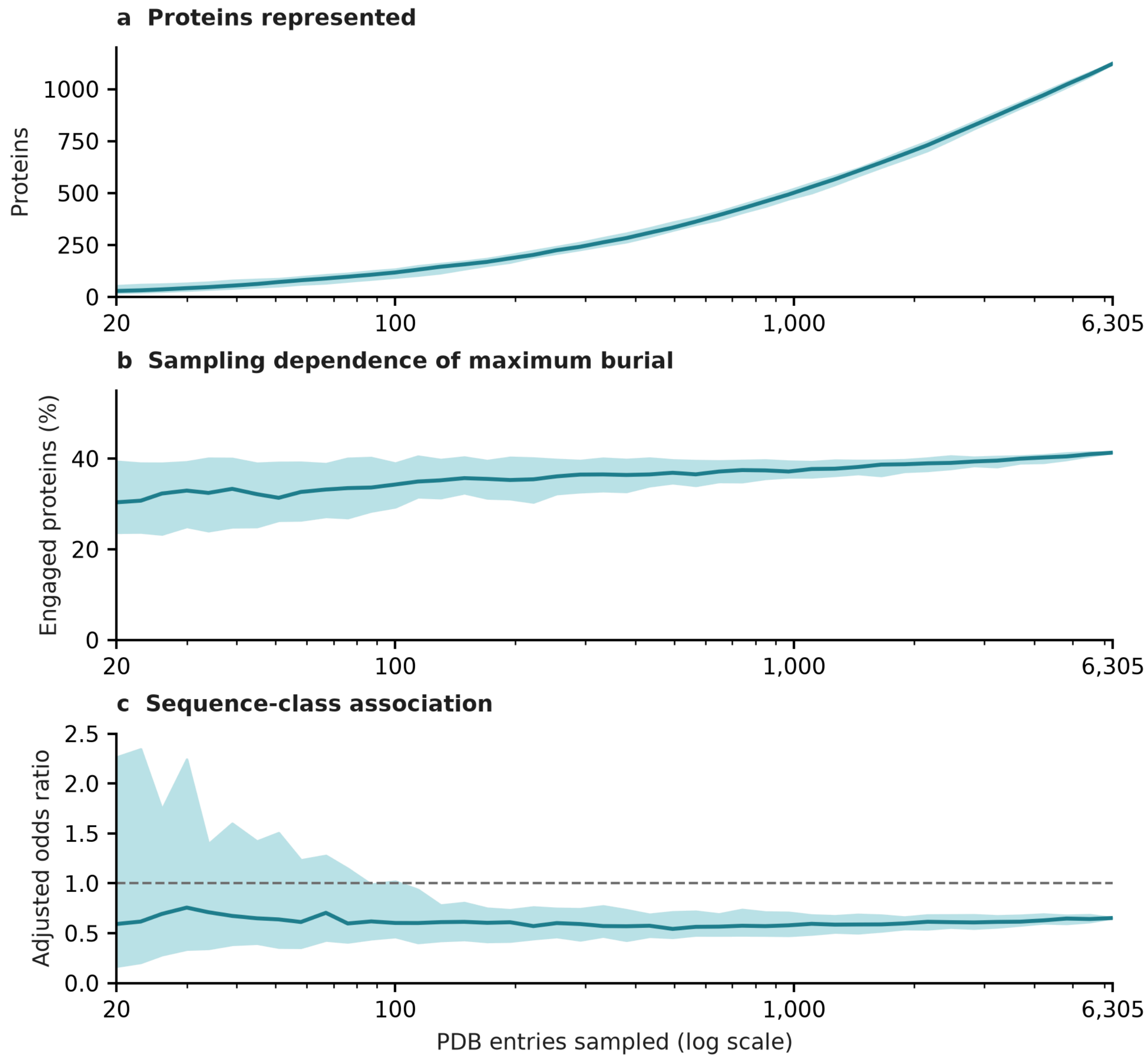

Supplementary Fig. 1 | Corpus-size sampling of the original extraction-selected cohort. a, Number of represented proteins. b, Percentage of represented proteins whose largest observed burial is at least 25%. c, Sequence-class odds ratio adjusted for natural-log chain count in the selected maximum-burial assembly. The comparison is MetAP-permissive versus nonpermissive residue-2 classes. PDB entries were sampled without replacement in 40 random orderings using seed 20250823, at 44 logarithmically spaced checkpoints from 20 to 6,305 entries. At each checkpoint, the maximum-burial observation was selected for each represented protein. Ties use the earliest row in the frozen observation table. Lines show medians and bands show the 10th–90th percentiles across orderings; these bands are not confidence intervals or biological replicate variation. At the full-corpus endpoint, 1,123 proteins are represented, 41.32% reach the burial threshold, and the adjusted class odds ratio is 0.652. The increasing engaged fraction illustrates the sampling dependence of maximum selection. Apparent numerical stability does not establish saturation of native proteoforms or remove sampling and residue-chemistry confounding. The analysis uses the original extraction-selected cohort, not the 845-protein first-observed sensitivity. Chain count is taken from the selected observation. Supplementary Data 8 includes every replicate/checkpoint value, the frozen inputs and the calculation and plotting code.

**Supplementary Fig. 2**

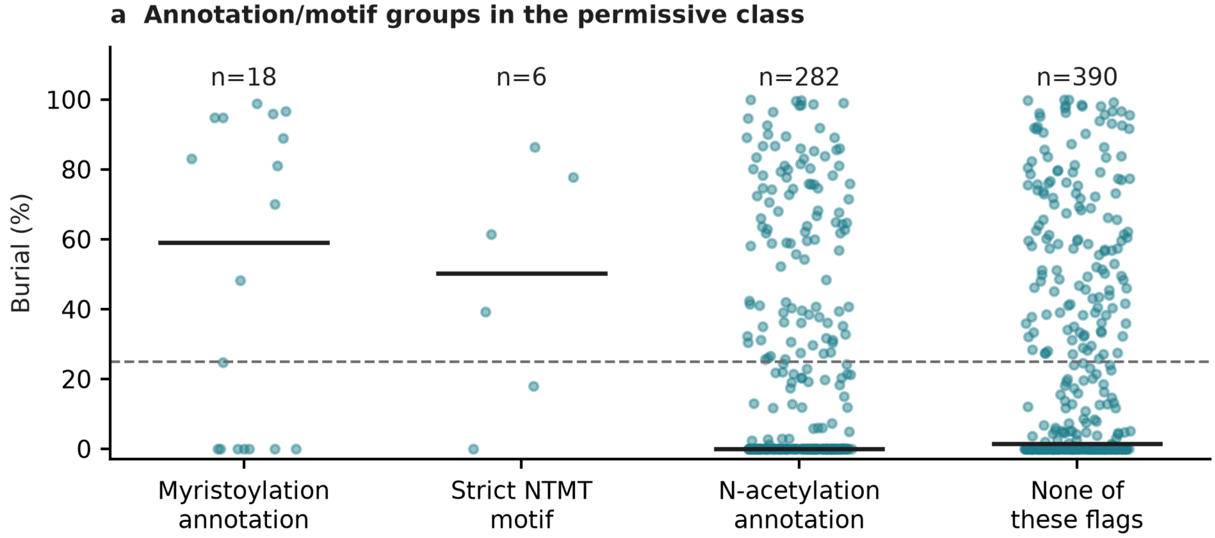


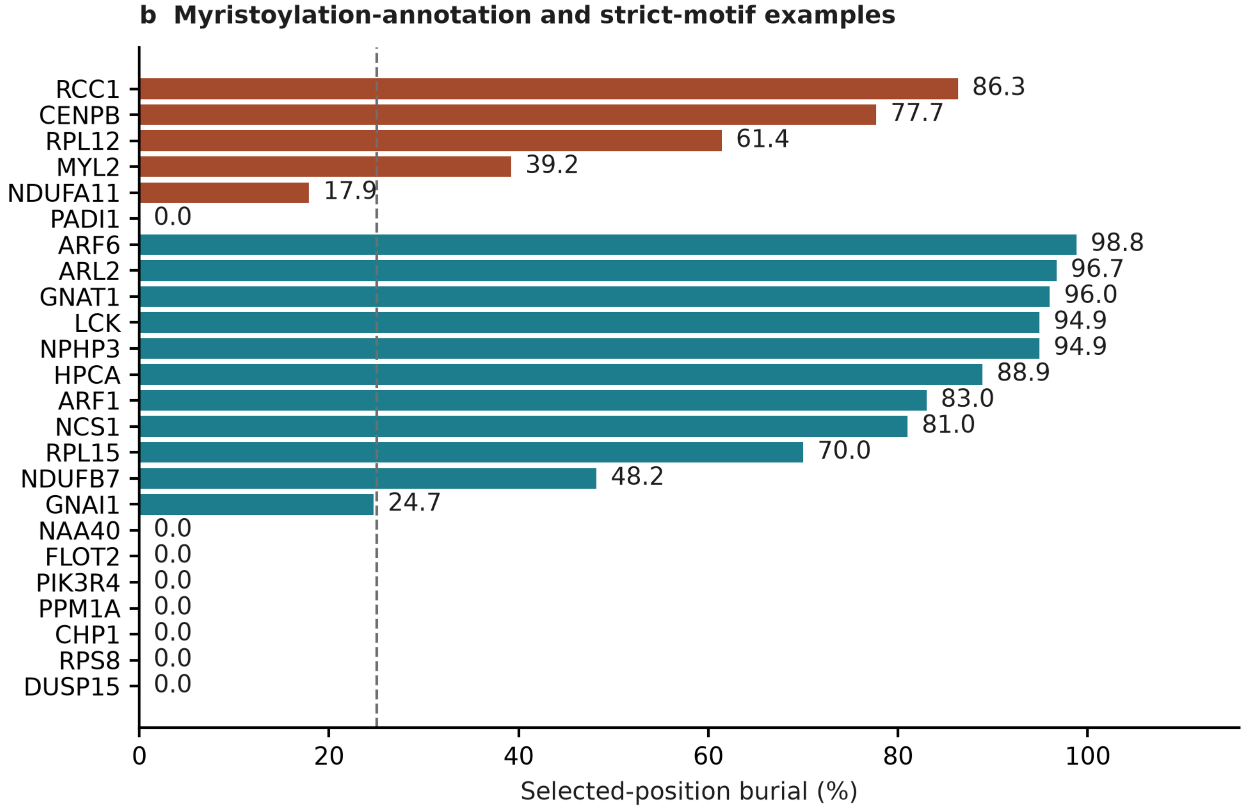

Supplementary Fig. 2 | Exported annotation/motif groups within the 696-protein MetAP-permissive sequence class. a, Maximum burial of the annotation-selected residue, with one point per protein and a horizontal median. Dashed lines mark 25% burial. Display groups are assigned exclusively in the following priority: myristoylation annotation (n = 18), strict NTMT motif without the preceding flag (n = 6), N-terminal acetylation annotation without either preceding flag (n = 282), and none of these flags (n = 390). Their median burial values are 59.1%, 50.3%, 0.0% and 1.55%, respectively. b, Individual members of the myristoylation-annotation and strict-motif groups. Bars and printed values show burial percentages only. The myristoylation-annotation examples occupy the lower group and the strict-motif examples occupy the upper group; all genes are directly labeled. The underlying strict motif is [A/P/S/G]-P-K at the annotation-selected start. It is an operational sequence screen, not exhaustive enzyme-substrate assignment or proof of methylation. These are inherited accession-level annotation and motif flags, not measurements of modification in each structural preparation. Group exclusivity is a display rule and does not mean modifications are biologically mutually exclusive. “None of these flags” does not mean an unmodified N terminus. Modeled position and first-observed flags are supplied with every plotted record in Supplementary Data 8. Burial includes protein chains only; lipid, nucleic-acid and ligand occlusion are not measured. Missing upstream coordinates, construct boundaries and capped or internal N atoms limit biological interpretation. This panel is descriptive, not a test that a modification causes interface engagement. Minimum N distances across all observations are not plotted beside maximum-burial examples, because those measurements can originate from different structures.

### Supplementary Fig. 3

22 chain observations · 3 proteins · 7 entries

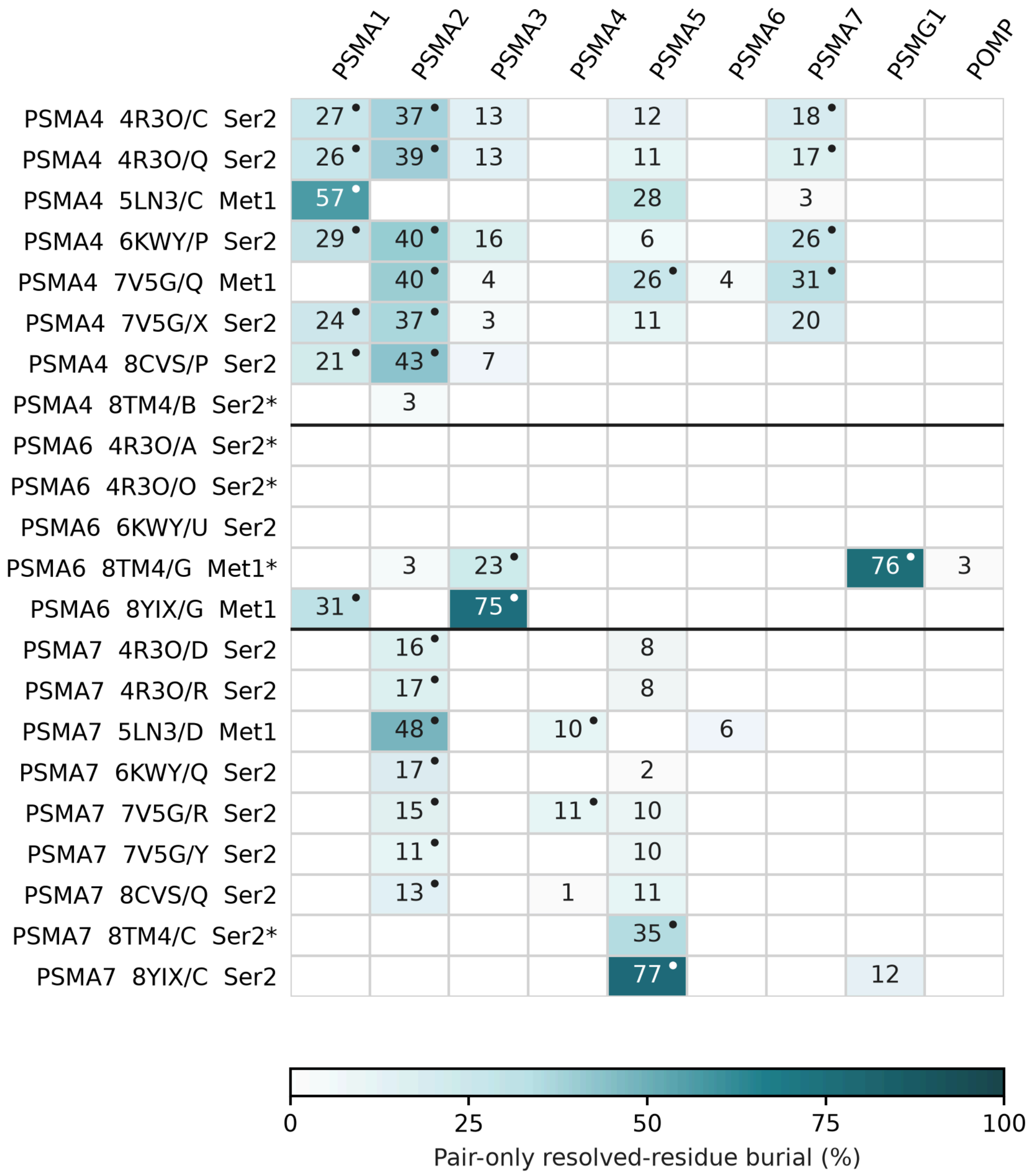

**Supplementary Fig. 3 | Partner-resolved geometry of selected PSMA4, PSMA6 and PSMA7 N-terminal positions.** Rows show 22 first-modeled Met1 or Ser2 observations from seven entries; columns identify partner genes. Shading and values ≥1% report burial of resolved target-residue atoms with one partner chain present. Dots mark heavy-atom approaches ≤4 Å. Pair-only burial uses target-residue SASA in the isolated target chain as denominator; values are nonadditive, not fractions of total assembly burial. The largest chain-specific value is displayed per partner gene; no row contains multiple informative chains of the same gene at the displayed thresholds. Blanks mean <1% burial or no eligible partner, not absence of interaction. Pair SASA was calculated for partners within 8 Å; more distant pairs are marked not calculated in Supplementary Data 7. The color scale is 0–100%. Asterisks mark missing OG in PSMA6 Ser2 (4R3O/A, 4R3O/O), PSMA4 Ser2 (8TM4/B) and PSMA7 Ser2 (8TM4/C), and missing CG, SD and CE in PSMA6 Met1 (8TM4/G). No atoms were rebuilt. Met1 and Ser2 are distinct positions, not sample-validated processing states. Proximity does not establish bonds, energies or function; chains and entries are not independent biological replicates. PSMA7 Ser2 occupies PSMA2-facing or PSMA5-facing environments, PSMA6 includes a PSMG1-facing Met1 backbone, and PSMA4 illustrates multi-partner packing. These comparisons do not establish temporal exchange, processing causality or disease mechanisms. Supplementary Data 7 supplies measurements, contacts, backbone sensitivities, chain assignments, code and frozen inputs. Main Fig. 4 and Supplementary Fig. 3 show the same map, not independent analyses. This selected comparison is distinct from group-level tests in Supplementary Tables 3–4 and Supplementary Data 9 and does not independently confirm proteasome-wide enrichment.

### Supplementary notes

#### Supplementary Note | Machine membership and scope

Prefix-defined machine summaries are descriptive groupings of the frozen extraction, not experimentally validated complexes or boundary-qualified enrichment estimates. Membership rules differ between the frozen machine map and the recurrence analysis; the two denominators must not be interchanged. The relevant original rules and code are retained in Supplementary Data 8.

The frozen map uses gene-symbol patterns for proteasome/assembly, CCT/TRiC, Mediator, spliceosome, cullin–RING/CSN, oxidative phosphorylation, ribosomes, nuclear-pore scaffold, RNA polymerases, eIF2/eIF3, actin/tubulin and cell-cycle/replication groups. A group is displayed when at least four scored members are present. The recurrence workbook separately groups only members of the 261-candidate subset.

The frozen Figure 2 map remains descriptive. The new enrichment reconstruction separately tests thirteen prefix-defined groups and explicitly curated memberships, each against the remaining eligible permissive proteins. Its correction families, boundary sensitivities and complete outcomes are reported in Supplementary Note, “Reconstructed enrichment”, Tables 3–4 and Data 9. Prefix summaries cannot validate a different curated membership analysis; neither analysis establishes processing causality or native-terminal persistence.

#### Supplementary Note | Ran is a terminal-boundary counterexample

In deposited 7MO1 chain A, the modeled prefix is Ser0–Met1–Ala2–Ala3. The measured Ala2 N lies 1.331 Å from the preceding Met1 carbonyl C, consistent with its being an internal backbone N. The original Ala2 burial of 66.8% and interchain N distance of 2.72 Å reproduce as positional measurements, but they do not measure a free native processed Ran terminus.

Static deletion of Ser0 and Met1 changes Ala2 burial to 54.1%. This is a diagnostic perturbation of the deposited coordinates, not a corrected native proteoform: the model is not relaxed, acetylated or experimentally validated. No effect on transport-factor competition, residence time, nucleotide exchange or turnover is established by this exercise.

The superseded conformer scan selected Ser0 rather than the intended Ala2 in this structure. Its Ran interpretation is not retained here. Supplementary Data 8 preserves the target-selection diagnostic for transparency; the old conformer artwork and structural atlas are not part of this supplementary figure set.

#### Supplementary Note | Processing-to-function experiments

The atlas is a starting point for biological questions, not an assignment of the proteoform created by a global processing perturbation. For nuclear-transport candidates, test the actual N-terminal sequence, initiator-methionine state, acetylation or other capping and construct boundary in the preparation used for structural or functional measurements. A Met-Ala sequence alone does not establish a specific degron or acetyltransferase-dependent fate.

A useful experiment has three linked readouts: preparation-matched terminal proteoform measurement, partner engagement or local structural context, and a pathway-relevant functional endpoint. Manipulations should preserve expression and stoichiometry where possible, include an appropriate rescue and distinguish loss of protein abundance from changes in partner binding. A static deletion model is not a substitute for a verified processed construct.

Disease- or stress-relevant settings can be introduced after the molecular comparison is defined. For proteasome candidates, compare native assembly and substrate degradation along with the terminal and partner readouts; for transport candidates, compare transport-related function with matched abundance controls. A disease mechanism should be claimed only if a defined terminal state is connected experimentally to both an interaction change and a phenotype.

**Supplementary Note | Proteasome evidence and state-dependent geometry**

Independent top-down analysis of human proteasome preparations reports initiator-methionine removal and N-acetyl-serine for accessions P25789 (PSMA4), P60900 (PSMA6) and O14818 (PSMA7). These observations supply independent proteoform evidence, not sample-matched proof for each structural preparation analyzed here.[1]

The partner-resolved analysis measures first-modeled Met1 or Ser2 in a fixed 22-observation subset from seven structural entries. It does not reclassify every proteasome structure. PSMA7 Ser2 examples distinguish PSMA2-facing from PSMA5-facing environments; PSMA6 includes a PSMG1-facing modeled Met1 backbone; and PSMA4 illustrates packing against multiple partners. These are context-associated comparisons, not a temporal handoff or proof that processing drives partner exchange.

Five focal residues are incomplete: four Ser2 models lack OG, and PSMA6 Met1 in 8TM4/G lacks CG, SD and CE. No atoms were rebuilt. Backbone-only sensitivities and chain-specific measurements accompany the map in Supplementary Data 7. Pair-only burial is nonadditive and must not be summed into a partition of total burial.

**Supplementary Note | Audit, selection and statistical scope**

The primary maximum-burial table retains the frozen model-selected values while correcting evidence labels. All 735 analyzable proteins previously labeled as retained have no removal feature in the frozen annotation table; 108 separately have an exported N-acetylmethionine annotation. Of 388 proteins with removal annotations, 21 include alternate removal. Missing removal annotation is not evidence of retention, and alternate annotation is not an exclusive proteoform assignment.

In the analyzable observation table, 5,329 of 19,967 rows are not first observed. The maximum-burial record is nonfirst for 396 of 1,123 proteins. The sensitivity removes nonfirst observations, then selects each remaining protein’s largest burial, preserving the original selected row in ties when it remains eligible; otherwise frozen observation order breaks ties. This retains 845 proteins, including 175 engaged permissive-class proteins among 463 and 178 engaged nonpermissive-class proteins among 382.

First-observed filtering is a coordinate-boundary sensitivity, not native-terminal validation. Upstream sequence can be unresolved, removed by construct design or chemically capped. Loss

[1] Lakshmanan et al., Proteomics (2014), Table 1 and sample preparation; doi:10.1002/pmic.201300339. Full text: https://escholarship.org/content/qt2x63q3r2/qt2x63q3r2_noSplash_d09b51f49c3c303fadc3a7ea0da41c18.pdf

of a candidate after filtering means that the retained subset no longer supports its threshold assignment; it does not disprove biological engagement.

Endpoint-offset mapping checks flag five analyzable observations with a one-residue shift: ACTG1 in 9SMX, MRPL13 in 6ZM5, 6ZM6 and 7PO4, and GSTP1 in 5L6X. Agreement with endpoint arithmetic is not a full residue-level alignment. Mapped-span flags identify 368 observations below 30 residues and 1,155 below 25% of canonical length; these overlapping exploratory flags are not direct coordinate-completeness measures.

Direct inspection of the 72 analyzable observations in 11 frozen assemblies identifies seven measured N atoms with upstream C–N distances of 1.0–1.8 Å. The seven include Ran and examples from PSMA2, PSMA3, PSMD11 and PSMG1. These coordinate checks cover a selected subset, not the entire atlas.

The primary binary outcome is residue burial ≥25%; sequence-class odds ratios compare permissive with nonpermissive proteins. Fisher and Mann–Whitney tests are two-sided. Logistic confidence intervals are Wald intervals, with natural-log chain count as the primary adjustment. Sequence-class sensitivity tests are exploratory, unadjusted and not independent replications. Group-enrichment odds ratios instead compare group members with nonmembers within the permissive class and use the explicit multiplicity corrections and directional stratified sensitivities in the Supplementary Note, “Reconstructed enrichment”. Sampling, residue chemistry and sequence-class membership remain correlated.

The continuous N-distance comparison excludes five proteins without a finite exported minimum N distance (four permissive and one nonpermissive), rather than ranking infinite missing-value sentinels as measurements. It therefore uses 692 permissive and 426 nonpermissive proteins and gives a two-sided Mann–Whitney p value of 0.004983. The all-protein burial comparison uses all 1,123 proteins.

Supplementary Data 4–5 separate recomputed current-cohort results from supplied chemistry-model estimates at exported precision. Historical mixed-cohort output is isolated in the provenance directory of Supplementary Data 8. The latter is not an alternative current result set.

**Supplementary Note | Reconstructed enrichment**

This is a new exploratory reconstruction, not recovery of the undocumented historical twelve-curated-comparison analysis. One accession contributes one record. The original background contains 696 permissive proteins, with 261 engaged at the primary 25% threshold. The first-observed background contains 463, with 175 engaged, after observation-level filtering and maximum reselection. Missing eligibility is not non-engagement, and the two backgrounds are not independent experiments.

The prefix family reuses the ordered assignment function in the frozen figure code, retaining the twelve groups with at least four scored proteins in the full original cohort and adding the histone-symbol family. Its exact symbol pattern is ^(HIST|H[1234](?:[A-Z0-9-]|$)). All thirteen tests are reported, including nonsignificant comparisons; the original twelve-group correction is a separately labeled sensitivity.

Curated membership uses the human Complex Portal ComplexTab file retrieved on 25 September 2026 UTC. It contains 2,498 manually curated entries; separately distributed predicted-complex files were not used. Manual curation includes inferred evidence and does not imply direct experimental proof for every entry.[2]

Expanded participant lists were normalized to canonical UniProt accessions, preserving isoform/processed-chain suffixes and alternative-set mappings in an audit. A strict sensitivity excluded suffix-normalized and alternative-set members. Sixty-four complexes with at least four original-background permissive accessions defined the outcome-independent eligible family, retained unchanged for the boundary analysis. Deduplication of identical original tested accession sets yielded 42 comparisons. Overlapping memberships are not independent confirmations.

Five focused questions used CPX-5993 (26S proteasome), CPX-6030 (CCT/TRiC), CPX-560/561/577/6123/6151 (OXPHOS I–V union), all seventeen recommended names containing “spliceosom”, and eight names beginning “Nucleosome” or “CENP-A nucleosome” (core-histone membership). These are selected biological questions, not prospective independent validation. Catalogue membership does not identify the structure supplying a protein’s maximum burial.

**Supplementary Note | Reconstructed enrichment: statistics and interpretation**

Tables are [[member engaged, member unengaged], [nonmember engaged, nonmember unengaged]]. Two-sided Fisher tests accompany sample odds ratios and exact conditional odds-ratio confidence intervals. BH and dependence-conservative BY q values are calculated separately for each declared family, cohort and threshold. Infinite sample odds ratios arise from zero cells, not infinite biological effects. Complete intervals, counts and all outcomes are in Data 9.

The 25% endpoint is primary; 20% and 30% are focused-family sensitivities. Exact one-sided upper-tail enrichment tests condition on fixed margins within assembly-size strata (2–3, 4–5, 6–11 and ≥12 chains) crossed with PDB-entry-count strata (one, two to four and ≥five). Hypergeometric probabilities are convolved without Monte Carlo sampling. Boundary deposition counts use eligible observations only. BH correction covers five focused stratified tests per cohort. Nuclear- versus mitochondrial-encoded OXPHOS tests and the histone-prefix stratified diagnostic are labeled post-result analyses.

Proteasome enrichment is present in the original extraction but is sensitive to boundary eligibility and multiplicity treatment. CPX-5993 has original catalogue-wide BH $q = 0.0309$, changing to 0.0507 after deduplicating tested membership sets and to 0.481 after boundary filtering. No boundary-qualified catalogue comparison passes BH correction. The PSMA4/6/7 partner-resolved observations in Data 7 remain a separate, candidate-specific evidence layer.

The histone-symbol association passes thirteen-test BH correction originally and after filtering ($q = 0.0216$ and 0.00180). The boundary result also passes BY correction ($q = 0.00571$), but

[2]Complex Portal human ComplexTab (retrieved 25 September 2026 UTC): https://ftp.ebi.ac.uk/pub/databases/intact/complex/current/complextab/9606.tsv
Curation and evidence documentation: https://www.ebi.ac.uk/complexportal/documentation

strengthens partly because two unengaged records lose eligibility. Selected maxima often concern histone-peptide recognition rather than intact nucleosomes. Related variants are not independent biological replications, and neither coordinate boundary nor catalogue membership validates native chemistry.

CCT/TRiC and OXPHOS remain exploratory. OXPHOS preserves 11/16 engaged records across eligibility schemes, but its boundary significance differs between the five-question and thirteen-prefix corrections ($q = 0.0389$ and 0.101) and does not pass the five-question sampling-stratified correction ($q = 0.0947$). The broader curated spliceosomal union is not enriched. Smaller selected correction families must not be used to override nonsignificant broader-screen results.

### Supplementary Table 1 | Robustness analyses

| Analysis | N / denominator | Odds ratio | 95% CI | Two-sided p |
|---|---|---|---|---|
| Primary Fisher comparison | 1,123 | 0.662 | Not shown | 0.000948 |
| Class + log chain count | 1,123 | 0.652 | 0.509–0.837 | 0.000764 |
| Plus log PDB-entry count | 1,123 | 0.686 | 0.532–0.885 | 0.003694 |
| First observed, reselected maximum: Fisher | 845 | 0.696 | Not shown | 0.011602 |
| First observed, reselected maximum: adjusted | 845 | 0.669 | 0.506–0.886 | 0.005030 |
| Exclude PSMA4/6/7 and Ran: Fisher | 1,119 | 0.652 | Not shown | 0.000724 |
| Exclude permissive modeled MET/MSE: Fisher | 381 vs 427 | 0.693 | Not shown | 0.010582 |
| Leave one residue-2 identity out | 20 tests | 0.578–0.716 | Not shown | Maximum 0.010330 |
| At least two PDB entries: Fisher | 739 | 0.668 | Not shown | 0.008655 |
| At least five PDB entries: Fisher | 371 | 0.695 | Not shown | 0.108593 |
| Alternative class boundaries | 1,123 | 0.663–0.731 | Not shown | 0.000974–0.023274 |
| Median / largest / heteromeric selection | 1,123 / 1,123 / 772 | 0.554 / 0.663 / 0.625 | Not shown | 0.000018 / 0.001518 / 0.001846 |

The first-observed analysis removes observations before reselection. Primary engagement counts are 261/696 versus 203/427; first-observed counts are 175/463 versus 178/382. Odds-ratio direction is permissive versus nonpermissive. Confidence intervals shown are adjusted-model Wald intervals; “not shown” does not indicate that an interval cannot be calculated.

Alternative boundaries reassign C to nonpermissive or add N/Q, D/N or D/N/Q to the permissive set. Largest-assembly selection maximizes chain count then burial; the heteromeric sensitivity retains proteins whose original selected maximum occurs in an assembly with at least two entities. Exact definitions, counts and full-precision results are in Supplementary Data 4–5.

### Supplementary Table 2 | Candidate-to-experiment priorities

| Question / candidates | Evidence-aware experiment | Interpretation controls |
|---|---|---|
| PSMA7: partner context | Compare verified Ser-starting preparations in PSMA2-facing and PSMA5-facing structural contexts; measure partner engagement and activity. | Match assembly context, abundance and terminal chemistry; no temporal exchange inferred. |
| PSMA6: chaperone versus mature context | Resolve terminal chemistry and backbone contacts in PSMG1-associated versus mature-complex preparations. | Do not interpret incomplete Met1 side-chain burial as a fully resolved interaction. |
| PSMA4: multi-partner packing | Test whether defined terminal changes redistribute partner contacts and alter native assembly or substrate degradation. | Pair-only burial is nonadditive; assay multiple partners rather than one presumed contact. |
| CCT8, CCT2, CCT5; CCT6A as boundary contrast | Test verified terminal states against complex abundance and client folding under matched conditions. | Boundary burial is 96.4%, 71.3% and 75.8% for the retained candidates; CCT6A falls to 0.0%. No robust complex-wide enrichment. |
| ARF6, LCK, GNAT1, NPHP3, NCS1 | Measure actual terminal proteoforms, myristoylation, localization and stability in the same preparation. | Accession annotation is not preparation-matched modification evidence. |
| RCC1, CENPB: motif candidates | Verify terminal chemistry and methylation, then test chromatin-related interactions and function. | Strict motif membership is a screen, not a demonstrated substrate assignment. |
| ALDOB, IMPDH2 | Test verified terminal states against oligomerization and catalytic readouts. | Separate abundance, stability and specific activity. |
| GTF2H5, RAD51D, MED21 | Measure terminal state, partner engagement and pathway-specific function. | Include construct, abundance and rescue controls; no disease causality inferred. |

These are prospective tests, not reported functional findings. The original selected measurements and first-observed eligibility flags are paired in Supplementary Data 5. Graded MetAP perturbation, where used, requires direct measurement of the focal proteoform and controls for broader changes in the cell.

### Supplementary Table 3 | Thirteen prefix-group comparisons

#### Original extraction

Background: 696 permissive proteins, 261 engaged. Each group is compared with the remaining permissive proteins; engaged means ≥25% burial.

| Prefix group | Engaged / eligible | Sample OR | Fisher p | BH q (13) | BY q (13) |
|---|---|---|---|---|---|
| Ribosomes (cyto + mito) | 20/45 | 1.36 | 0.342 | 0.555 | 1 |
| OXPHOS complexes | 11/16 | 3.78 | 0.0157 | 0.0563 | 0.179 |
| 26S proteasome / assembly | 12/15 | 6.94 | 0.000879 | 0.0114 | 0.0363 |
| Cell cycle / replication | 5/7 | 4.23 | 0.11 | 0.237 | 0.755 |
| Actin / tubulin systems | 4/8 | 1.68 | 0.482 | 0.626 | 1 |
| Spliceosome | 9/13 | 3.85 | 0.0216 | 0.0563 | 0.179 |
| RNA polymerases | 2/4 | 1.67 | 0.633 | 0.748 | 1 |
| Cullin–RING / CSN | 2/8 | 0.552 | 0.717 | 0.776 | 1 |
| eIF2 / eIF3 | 1/6 | 0.331 | 0.419 | 0.605 | 1 |
| TRiC/CCT chaperonin | 4/4 | Inf | 0.0195 | 0.0563 | 0.179 |
| Mediator | 3/4 | 5.05 | 0.151 | 0.28 | 0.891 |
| Nuclear pore scaffold | 1/4 | 0.554 | 1 | 1 | 1 |
| Histone-symbol family | 9/11 | 7.73 | 0.00332 | 0.0216 | 0.0687 |

Two-sided Fisher tests; BH, Benjamini–Hochberg; BY, Benjamini–Yekutieli. Inf denotes a zero unengaged-member cell. The count denominators are eligible permissive members, not all scored proteins or all curated subunits. Exact conditional confidence intervals and full-precision values are in Supplementary Data 9 (prefix13_tests.csv).

### Supplementary Table 3 | Thirteen prefix-group comparisons (continued)

#### First-observed sensitivity

Background: 463 permissive proteins, 175 engaged. Observations are filtered before maximum reselection. The same thirteen-group correction family is retained.

| Prefix group | Engaged / eligible | Sample OR | Fisher p | BH q (13) | BY q (13) |
|---|---|---|---|---|---|
| Ribosomes (cyto + mito) | 16/40 | 1.11 | 0.865 | 1 | 1 |
| OXPHOS complexes | 11/16 | 3.8 | 0.0155 | 0.101 | 0.321 |
| 26S proteasome / assembly | 7/10 | 3.96 | 0.0466 | 0.202 | 0.643 |
| Cell cycle / replication | 4/5 | 6.71 | 0.0701 | 0.228 | 0.725 |
| Actin / tubulin systems | 3/4 | 5.01 | 0.154 | 0.286 | 0.908 |
| Spliceosome | 4/7 | 2.22 | 0.434 | 0.564 | 1 |
| RNA polymerases | 1/1 | Inf | 0.378 | 0.546 | 1 |
| Cullin–RING / CSN | 2/5 | 1.1 | 1 | 1 | 1 |
| eIF2 / eIF3 | 0/3 | 0 | 0.293 | 0.476 | 1 |
| TRiC/CCT chaperonin | 3/4 | 5.01 | 0.154 | 0.286 | 0.908 |
| Mediator | 3/4 | 5.01 | 0.154 | 0.286 | 0.908 |
| Nuclear pore scaffold | 0/1 | 0 | 1 | 1 | 1 |
| Histone-symbol family | 9/9 | Inf | 0.000138 | 0.0018 | 0.00571 |

Two-sided Fisher tests; BH, Benjamini–Hochberg; BY, Benjamini–Yekutieli. Inf denotes a zero unengaged-member cell. The count denominators are eligible permissive members, not all scored proteins or all curated subunits. Exact conditional confidence intervals and full-precision values are in Supplementary Data 9 (prefix13_tests.csv).

### Supplementary Table 4 | Curated and focused comparisons

#### A. Selected catalogue comparisons

| Complex / cohort | Engaged / eligible | Sample OR | Fisher p | BH q (64) | BY q (64) |
|---|---|---|---|---|---|
| CPX-5993 Original | 10/13 | 5.74 | 0.00628 | 0.0309 | 0.147 |
| CPX-5993 Boundary | 7/10 | 3.96 | 0.0466 | 0.481 | 1 |
| CPX-6030 Original | 4/4 | Inf | 0.0195 | 0.0734 | 0.348 |
| CPX-6030 Boundary | 3/4 | 5.01 | 0.154 | 0.481 | 1 |

CPX-5993: 26S proteasome. CPX-6030: CCT/TRiC. These rows are selected for discussion from the complete 64-complex family, not a separate four-test correction. All 64 outcomes are supplied in curated_catalogue_tests.csv. No boundary-qualified complex passes catalogue-wide BH correction. Deduplicating original tested accession sets yields 42 comparisons and changes original CPX-5993 BH q from 0.0309 to 0.0507.

#### B. Five focused questions: original extraction

| Focused group | Engaged / eligible | Sample OR | Fisher p | BH q (5) | BY q (5) | Stratified BH q (5) |
|---|---|---|---|---|---|---|
| 26S proteasome | 10/13 | 5.74 | 0.00628 | 0.0314 | 0.0717 | 0.091 |
| CCT/TRiC | 4/4 | Inf | 0.0195 | 0.0325 | 0.0742 | 0.0947 |
| OXPHOS I–V | 11/16 | 3.78 | 0.0157 | 0.0325 | 0.0742 | 0.091 |
| Spliceosomal complexes | 11/22 | 1.7 | 0.264 | 0.264 | 0.603 | 0.421 |
| Core-histone membership | 5/7 | 4.23 | 0.11 | 0.137 | 0.313 | 0.091 |

Original background: 696 permissive proteins, 261 engaged. Focused questions were selected from the manuscript’s biological themes; they are not independent validation. Two-sided Fisher p values are contrasted with the separately labeled exact one-sided sampling-stratified enrichment tests. Complete intervals and all thresholds are in Data 9.

**Supplementary Table 4 | Focused boundary sensitivity (continued)**

**C. Five focused questions: first-observed sensitivity**

| Focused group | Engaged / eligible | Sample OR | Fisher p | BH q (5) | BY q (5) | Stratified BH q (5) |
|---|---|---|---|---|---|---|
| 26S proteasome | 7/10 | 3.96 | 0.0466 | 0.0777 | 0.177 | 0.161 |
| CCT/TRiC | 3/4 | 5.01 | 0.154 | 0.192 | 0.439 | 0.291 |
| OXPHOS I–V | 11/16 | 3.8 | 0.0155 | 0.0389 | 0.0887 | 0.0947 |
| Spliceosomal complexes | 4/13 | 0.725 | 0.774 | 0.774 | 1 | 0.91 |
| Core-histone membership | 5/5 | Inf | 0.00744 | 0.0372 | 0.085 | 0.0135 |

Boundary background: 463 permissive proteins, 175 engaged. The same five questions are retained. “Stratified” conditions on assembly-chain-count and eligible PDB-entry-count strata as defined in the Supplementary Note, “Reconstructed enrichment”; it adjusts measured sampling covariates only. Full stratum-test p values, expected counts and informative-stratum counts are in focused_stratified_tests.csv.

The core-histone membership union (5/7 original, 5/5 boundary) differs from the histone-symbol family in Table 3 (9/11 and 9/9). It excludes some variants and does not establish that the maximum-burial structures are nucleosomes. The 26S curated membership likewise differs from the proteasome/assembly prefix set. Do not interchange denominators or q values across these definitions.

In the OXPHOS encoding diagnostic, five of six mitochondrially encoded members and six of ten nuclear-encoded members are engaged in both cohorts. Neither subgroup passes correction across the two subgroup tests. Threshold, strict-identifier, deduplicated-membership and encoding diagnostics are fully reported in Data 9; they do not change the designated 25% primary endpoint.

### Supplementary data inventory

| Data | Format | Contents and scope |
|---|---|---|
| 1 | XLSX | 1,191 protein records with corrected annotation/model labels, exclusion and boundary flags; separate 845-protein first-observed subset. |
| 2 | XLSX | 22,291 observations with analyzable, boundary, mapping and construct-span flags; 19,967 analyzable observations. |
| 3 | XLSX | Separate original 261-candidate and first-observed 175-candidate tables, with eligibility transitions. |
| 4 | XLSX | Current-cohort statistical summaries, tests, models and sensitivities; supplied chemistry-model estimates clearly identified; descriptive frozen prefix map. |
| 5 | XLSX | Current robustness calculations and evidence-aware candidate-to-experiment map. |
| 6 | XLSX | Conditional recurrence of the original 261 candidates; chain-weighted and entry-weighted bins explicitly distinguished; no claim of proteoform-complete persistence. |
| 7 | ZIP | 22-observation partner-resolved proteasome analysis: measurements, contacts, atom-completeness and backbone sensitivities, chain assignments, code and seven frozen assemblies. |
| 8 | ZIP | Two ZIP files, 8a and 8b: audit results, current CSV exports, source data for Supplementary Figs. 1–2, frozen inputs, code and historical provenance. Extract both into the same directory to restore the complete archive. |
| 9 | ZIP | Reconstructed enrichment: browsable results workbook, all test tables, curated memberships and mapping audit, frozen inputs, code, environment versions, validation and source values for Tables 3–4. |

Supplementary Data 7–9 comprise four ZIP files: Data 7, Data 8a, Data 8b and Data 9. Data 8a and 8b are two parts of the same dataset; extract both into one directory, preserving their shared Supplementary_Data_8 folder. Original numerical records are retained. Data 8a additionally supplies the current figure-presentation code and its source-table inputs in presentation_20260925; earlier renderings remain provenance, not alternative analyses. Data 9 contains its own results workbook, not a separately numbered Data 10. Supplementary Fig. 3 repeats main Fig. 4 and is not an independent analysis; its source values are in Supplementary Data 7.

The superseded structural-atlas PDF and legacy conformer-scan artwork are excluded: their original terminal-state interpretations are not carried forward. Superseded statistical tables are retained only as labeled historical provenance, not as current conclusions. The current results and their provenance are distinguished throughout this inventory.
